# Discovery of AVI-6451, a Potent and Selective Inhibitor of the SARS-CoV-2 ADP-Ribosylhydrolase Mac1 with Oral Efficacy in vivo

**DOI:** 10.1101/2025.10.11.681833

**Authors:** Priyadarshini Jaishankar, Galen J. Correy, Yusuke Matsui, Takaya Togo, Moira M. Rachman, Maisie G.V. Stevens, Eric R. Hantz, Jeffrey Zheng, Morgan E. Diolaiti, Mauricio Montano, Taha Y. Taha, Julia Rosecrans, Julius Pampel, Nevan J. Krogan, Brian K. Shoichet, Alan Ashworth, Melanie Ott, James S. Fraser, Adam R. Renslo

## Abstract

The COVID-19 pandemic made plain the need for effective antivirals acting on novel antiviral targets, among which viral macrodomains have attracted considerable attention. We recently described AVI-4206 (**1**), a potent and selective inhibitor of the SARS-CoV-2 ADP-ribosylhydrolase Mac1 based on a 9*H*-pyrimido[4,5-*b*]indole core, the first Mac1 inhibitor to demonstrate antiviral efficacy in mouse models of SARS-CoV-2 infection, but requiring IP administration and frequent dosing. Herein we describe an extensive, structurally enabled medicinal chemistry effort to identify orally bioavailable Mac1 inhibitors by addressing permeability and efflux liabilities of **1** and many of its analogs. Multiple strategies were pursued to overcome these issues, including replacing a urea function to reduce hydrogen bond donor count. While heterocyclic urea mimetics could deliver analogs like AVI-6318 (**3**) with potencies and ADME profiles similar to **1**, abrogation of the P-gp liability was finally achieved with entirely non-polar substituents in place of urea. Thus, AVI-6451 (**4**) is a potent Mac1 inhibitor lead with low intrinsic clearance, high oral bioavailability, and antiviral efficacy with once-daily oral administration in a mouse model of SARS-CoV-2 infection.

## INTRODUCTION

The COVID pandemic emphasized the urgent need for effective antiviral agents acting by novel mechanisms.^1,2^ The SARS-CoV-2 macrodomain Mac1 which is part of the non-structural protein 3 (NSP3) has emerged as an intriguing and novel target for the development of new antiviral agents.^3^ Much of this enthusiasm has been based on compelling genetic evidence in which SARS-CoV-2 virus bearing deletion or inactivating mutations in Mac1 are found to have dramatically reduced pathogenesis in mice.^4,5^ As an ADP-ribosylhydrolase, the function of Mac 1 is to blunt the host immune response by removing ADP-ribose marks installed by host PARPs to mediate interferon signalling.^6,7^ The discovery of Mac1 inhibitors has been pursued by our combined research groups and several others since the beginning of the COVID-19 pandemic.^7–11^ Nevertheless, the development of potent inhibitors with drug-like properties suitable for *in vivo* validation of Mac1 as an antiviral target have remained elusive. Recently, we reported on the discovery of the AVI-4206 (**1**), a potent (Mac1 IC_50_ = 20 nM) and selective Mac1 inhibitor with an ADME/PK profile suitable for in vivo pharmacodynamic evaluation by the intraperitoneal (IP) route.^12^ Gratifyingly, in a lethal infection model involving K18-hACE2 mice infected with the WA1 strain of SARS-CoV-2, IP dosing of **1** at 100 mg/kg BID for five days produced both a survival benefit and significantly reduced viral titers in lung, thus providing the first pharmacological validation of Mac1 an antiviral drug target.

Although its potency and selectivity profile make **1** a vital in vivo tool compound, its poor permeability and a high efflux in Caco2 assays unsurprisingly translated to limited (<5%) oral bioavailability in mouse. We attributed this to an abundance of hydrogen bond donors, including two in the C8 urea function that engages Asp22 of Mac1. Empirical SAR led us to suspect the urea function as also contributing to P-gp recognition. To find alternate Mac1 inhibitor chemotypes, we employed shape-based screening (FrankenROCS) using fused fragments as queries, to efficiently search across billions of tangible fragment- and lead-sized molecules.^13^ Among the most promising fragments from this effort was the deazapurine (7*H*-pyrrolo[2,3-*d*]pyrimidine) analog **2** (AVI-1504, Figure 1), which exhibited excellent Caco2 permeability without efflux, combined with good ADME properties, and good ligand efficiency (L.E. = 0.43). Thus, from two distinct chemical starting point, we sought to develop potent, selective, and orally bioavailable Mac1 inhibitor leads.

**Figure 1.**
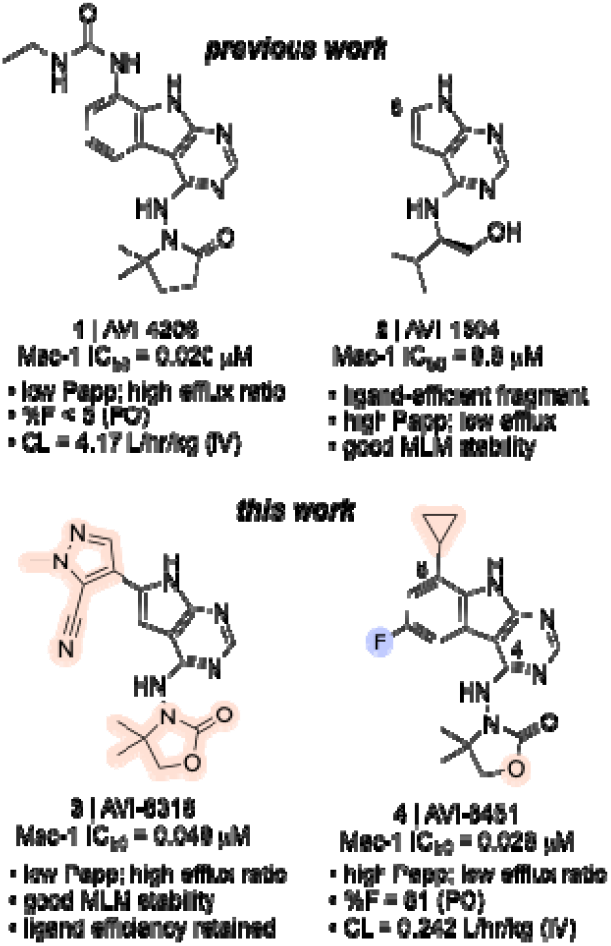
Structures and properties (top) of Mac1 inhibitor AVI-4206 (**1**) and a ligand-efficient deazapurine fragment AVI-1504 (**2**), described by our groups previously. Structures and properties (bottom) of optimized leads **3** (AVI-6318) and **4** (AVI-6451), the discovery of which is detailed herein.

Herein, we describe the evolution of fragment **2** to the potent and ligand efficient lead **3** (AVI-6318; L.E. = 0.39, LipE = 6.0), by combining a heterocyclic urea surrogate at C6 with the introduction of an optimized oxazolidinone sidechain at C4 (Figure 1). We also describe an extensive, and ultimately successful effort to resolve the poor permeability and P-gp mediated efflux of **1**, ultimately arriving at the potent (Mac1 IC_50_ = 28 nM) and orally bioavailable (61 %F) lead **4** (AVI-6451). In mice, compound **4** maintains free plasma exposure above its Mac1 IC_50_ for a full 24 hours after a single 50 mg/kg PO dose. Its excellent PK profile and potent Mac1 inhibition enabled oral efficacy in SARS-CoV-2 infected mice with once-daily administration. The studies outlined herein provide a case-study in addressing a challenging ADME liability, while ultimately delivering a potent, orally bioavailable Mac1 inhibitor as a therapeutic lead, and as an ideal tool compound for cellular studies of macrodomains and their role in altering the innate immune response to viral infection.

## RESULTS

The complex structures of **1** and ADP-ribose bound to Mac1 reveal the key polar and hydrophobic interactions formed by substrate and inhibitor in the Mac1 active site (Figure 2). These include aromatic (F156) and aliphatic (L126) amino acid side chains that flank the adenine and pyrimidoindole heterocycles, backbone amides (F156, D157), and hydrophobic side chains (V155, L160) in the ‘southern’ ribose binding site, and a ‘northern’ region comprising Asp22 that hydrogen bonds with the heteroaryl N–H and urea functions of substrate and inhibitor. Notably, polar interactions with the F156/D156 backbone is mediated by a water molecule in the case of ADP-ribose while the pyrrolidinone carbonyl of **1** makes direct hydrogen bonding contact with these backbone amides. A tunnel that leads to the catalytic machinery, and which binds the triphosphate of ADP-ribose is unoccupied by **1** and indeed yielded few hits in two prior fragnement screening efforts.

**Figure 2.**
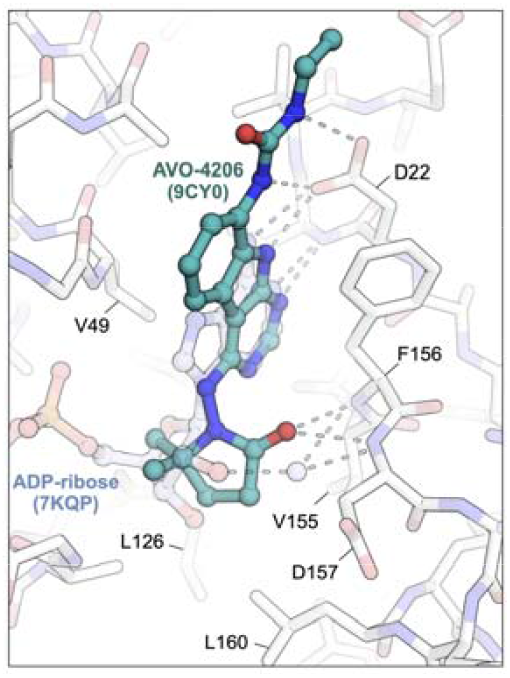
Superposition of the complex structures of **1 (9C**Y0) and ADP-ribose (7KQP) bound to Mac1 illustrating im**po**rtant polar contacts made with the backbone amides of F156/D**15**7 in the ribose binding subsite and multiple hydrogen bond**s** with Asp22. Major hydrophobic contacts in the adenine site in**clu**ding F156, L126, L160, and V155 are indicated. Note the str**uct**ured water that mediates hydrogen bonding between substrate and backbone residues in the ribose binding site.

In our initial SAR studies leading to **1**, we found that an alkyl urea function at C8 formed productive H-bonding contacts with Asp22 and afforded ≥10-fold gains in potency as compared to C8 acetamide or carbamate functionality (Table 1). Of additional note is a hydrophobic cleft lying above **and** behind Asp22 (as viewed in Figure 2) that tolerates large hydrophobes like the cyclopentane of urea analog AVI-**4**057, the most potent of these early leads (Mac1 IC_50_ = 42 nM). While aqueous solubility and mouse microsome stabilities of the urea analogs were favorable, efflux in Caco2 monolayers expressing P-gp was high (efflux ratio; ER >10). At the southern ribose binding site, we found the introduction of gem-dimethyl substitution on the C4 pyrrolidione ring improved potencies up to 5-fold across the series, but unfortunately did not mitigate efflux in Caco2 assays.

**Table 1.** Structure-Activity Relationships in early 9*H*-pyrimidoindole analogs bearing C4 pyrrolidinones combined with C8 urea or related side chains^a^.

| compound | R = |
| --- | --- |
| AVI-1499 | NH<sub>2</sub> |
| AVI-1501 | NHAc |
| AVI-1500 | NHC(=O)NHEt |
| AVI-3367 | NHC(=O)OEt |
| AVI-3766 | NHSO<sub>2</sub>NHMe |
| AVI-4051 | NHC(=O)NH-cyclopropyl |
| AVI-4057 | NHC(=O)NH-cyclopentyl |

| compound | Mac1 IC<sub>50</sub><br>(μM) | MLM T<sub>1/2</sub><br>(min) | Caco2<br>ER | Ksolb<br>(μM) |
| --- | --- | --- | --- | --- |
| AVI-1499 | 0.63 | >120 | — | 294 |
| AVI-1501 | 1.6 | >120 | 39.2 | 355 |
| AVI-1500 | 0.083 | >120 | 51.6 | 418 |
| AVI-3367 | 3.1 | — | — | — |
| AVI-3766 | 2.0 | — | — | — |
| AVI-4051 | 0.19 | — | — | — |
| AVI-4057 | 0.042 | >120 | 311 | 315 |

<sup>a</sup>Mac1 IC<sub>50</sub> in HTRF assay; MLM = mouse liver microsome stability; ER = efflux ratio BtoA/AtoB; Ksolb = kinetic solubility in pH 7.4 PBS. See Figure S1 for dose-response curves.

Our FrankenROCS screening campaign^13^ identified various C4 side chains for the ‘southern’ ribose binding site of Mac1, many of which could be applied beneficially in either the deazapurine (e.g. **2**) or tricyclic 9*H*-pyrimido[4,5-*b*]indole (e.g. **1**) series. For example, the C4 valinol side chain present in **2** was observed to form hydrophobic and polar contacts that mimic, respectively, those of the gem-dimethyl and carbonyl functions of the pyrrolidinone side chain in **1**. Being fragment-sized, **2** formed limited contacts with the northern site, although notably formed a hydrogen bond with Asp22. As noted above, fragment **2** was of considerable interest given its promising ADME profile, which included high permeability (P_app A-B_ = 14.9 × 10^-6^ cm/s) and lack of efflux (ER = 0.**79**) in Caco2 monolayers.

To elaborate **2** towards more potent analogs, we sought to target Asp22 and the surrounding hydrophobic cleft that binds distal alkyl substituent in urea analogs like **1**. Tactically, this involved the introduction of heterocycles at C6 were judged to mimic an amide or urea (i.e., with N–H or C–H donors) while also contacting the hydrophobic cleft (Chart 2 and Table 2). Also, modification of the C6 position in **2** was much more tractable than analogous modification of C8 position in **1**. Hence, in contrast to a ∼6-10 step synthesis required for **1** and analogs, a streamlined 2-3 step synthetic route was feasible for the desired aryl and heteroaryl C6 analogs of **2** (Scheme 1). This route, which proceeds through sequential SNAr and Suzuki coupling reactions, enabled rapid exploration of (hetero)aryl side chains.

**Table 2.** In vitro ADME data for AVI-1504 and selected C6-substituted analogs from Chart 1.^a^.

| Compound | Mac1 HTRF IC<sub>50</sub> (μM) | MLM CL (μL/min/mg) | MLM T<sub>1/2</sub> (min) | Caco2 Papp A to B (x 10<sup>-6</sup> cm/s) | Caco2 Papp B to A (x 10<sup>-6</sup> cm/s) | Kinetic Solubility in pH 7.4 PBS (μM) |
| --- | --- | --- | --- | --- | --- | --- |
| AVI-1504 (2) | 9.8 | <11.6 | >120 | 14.9 | 11.3 | 370 |
| AVI-4678 | 1.9 | 611 | 2.3 | 17.4 | 13.2 | 479 |
| AVI-4334 | 1.1 | 841 | 1.65 | 10.2 | 16.5 | 498 |
| AVI-4684 | 0.27 | 354 | 3.92 | 2.12 | 16.7 | 476 |
| AVI-4683 | 0.40 | 198 | 7.00 | 3.82 | 17.6 | 457 |
| AVI-4266 | 0.77 | 29.1 | 47.6 | 0.2 | 15.3 | 508 |
| AVI-4054 | 0.33 | 52.3 | 26.5 | 0.56 | 16.3 | 549 |

<sup>a</sup>Mac1 IC<sub>50</sub> in HTRF assay; MLM = mouse liver microsome clearance; Caco2 apparent permeability in apical to basolateral (A to B) and basolateral to apical (B to A) directions; kinetic solubility in pH 7.4 PBS. See Figure S1 for dose-response curves.

A survey of more than twenty C6 (hetero)aryl side chains was thus undertaken, as summarized below (Chart 1 and Table 2), with complex structures for representative analogs confirming a conserved binding mode across the series (Figure 3). We found that aryl substitution at C6 was tolerated and generally produced a 2–4-fold improvement over **2** in Mac1 IC_50_ values, as determined in a homogenous time-resolved fluorescence (HTRF) ligand displacement assay.^7^ Analogs with electron withdrawing ortho substituents (F, CN) were moderately but consistently more potent than comparators (Cl, CH_3_) possibly due to more favorable interaction of ortho C–H functions of the aryl ring with Asp22. This was supported by the complex structure of ortho-F analog AVI-4678 in which the opposing ortho C–H bond is directed toward Asp22, although the closest H-bonding contact is in fact formed with the deazapurine N–H function (Figure 3).

**Figure 3.**
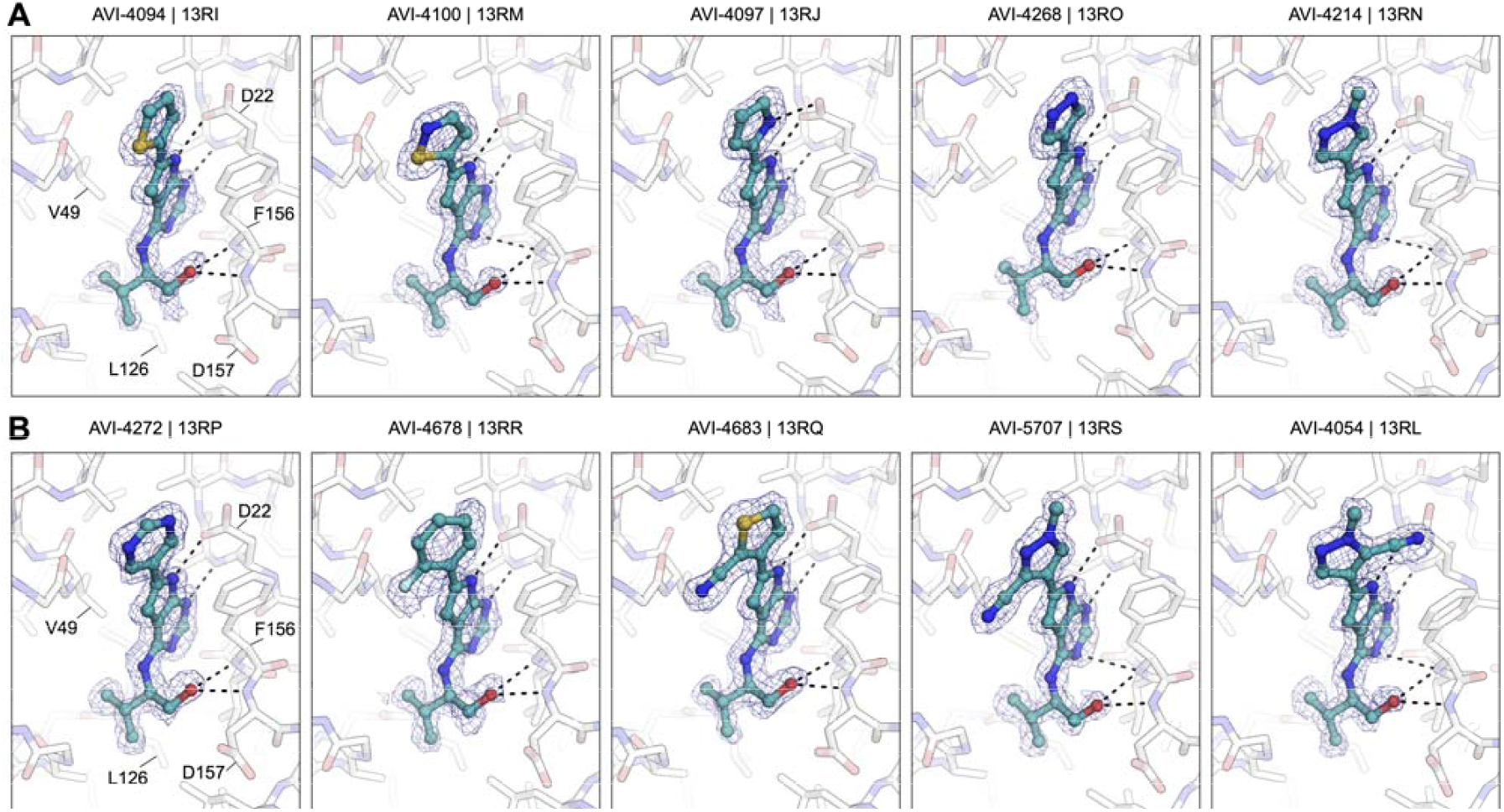
Complex structures of analogs from Chart 1 bound to Mac1. Panel A) from left to right, AVI-4094, AVI-4100, AVI-4097, AVI-4268, AVI-4214. Panel B) left to right, AVI-4272, AVI-4678, AVI-4683, AVI-5707, AVI-4054. PanDDA event maps are contoured around ligands at 2 σ (blue mesh). Hydrogen bonds are shown with dashed black lines.

Next, we explored 6 and 5-membered heterocycles at C6, including ring systems capable of presenting an N–H, acidic C– H, or a heterocyclic S atom (σ-hole) for interaction with Asp22. We found that pyridin-3-yl (AVI-4271), pyrimid-3-yl (AVI-4272), thiophen-2-yl (AVI-4094), thiophen-3-yl (AVI4334), furan-2-yl (AVI-4267), pyrrol-2-yl (AVI-4097)), 1*H*pyrazol-5-yl (AVI-4099), 1*H*-pyrazol-4-yl (AVI-4268), and isothiazol-5-yl (AVI-4100) analogs were either equipotent or at most two-fold more potent than unsubstituted aryl comparator AVI-4211. This revealed broad tolerance for flat (hetero)aryl substitution at C6, but also suggested that these substituents did not form strong interactions with Asp22. In particular, we noted that analogs seemingly best equipped present an N–H hydrogen bond donor (AVI-4099 and AVI-4097), or an empty σ orbital (AVI-4100) for interaction with Asp22 were no more potent than analogs that could not. Indeed, while the pyrrole N–H of AVI-4097 did appear to interact with Asp22 in the complex structure, the S atom of AVI-4100 was oriented away from Asp22 (Figure 3). Also, the introduction of aryl or thiophene rings at C6 was associated with reduced stability in liver microsomes, while good aqueous solubility was retained (Table 2). Caco2 permeability varied widely across the various (hetero)aryl C6 analogs, from high in fluoroaryl (AVI-4678) to low for pyrazoles generally (e.g., AVI-4268).

**Scheme 1.**
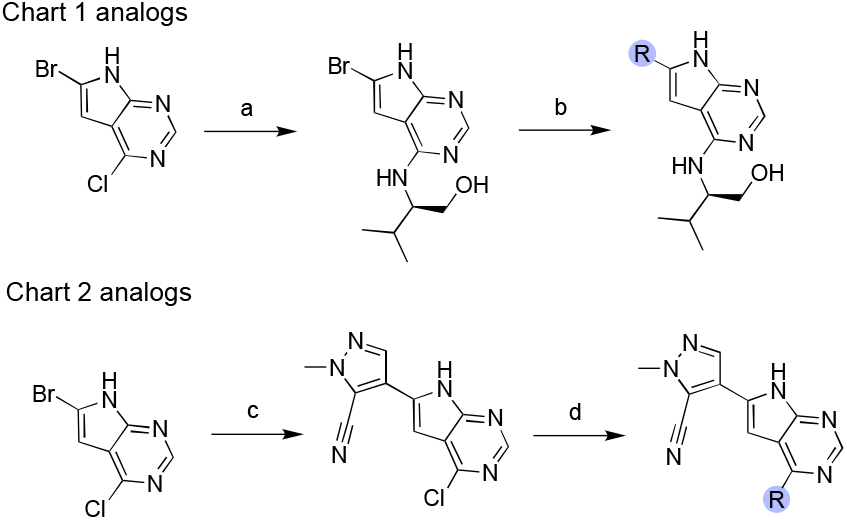
Synthesis of deazapurine analogs described below in Chart 1 (top scheme) and Chart 2 (bottom scheme). Conditions: (a) (*R*)-2-amino-3-methylbutan-1-ol, Et_3_N, DMSO, 110°C, 16 h, 45%. (b) R-B(OH)_2_, Pd(dppf)Cl_2_, Cs_2_CO_3_, dioxane-water (10:1), 110 °C; (c) 1-methyl-4-(pinacolborane)-1H-pyrazole-5-carbonitrile, dioxane-water (10:1), Pd(dppf)Cl_2_, K_2_CO_3_, 95 °C, 2 h, 72%; (d) R-NH_2_, iPrOH, aq. HCl, 100 °C.

Since ortho withdrawing group (F or CN) proven beneficial in the aryl series, we explored analogous substitution in the heteroarene analogs. Gratifyingly, an ortho CN afforded enhanced potencies in both thiophen-3-yl analogs (AVI-4683 and AVI-4684) and pyrazol-4-yl analogs (AVI-4054 and AVI5707), with Mac1 IC_50_ values improved ∼3–4-fold into the mid-nM regime (0.27–0.40 μM). The complex structures of C6 analogs showed an overall similar binding mode with a C– H bond preferentially oriented toward Asp22 (Figure 3). This suggests a weak C–H--O interaction with Asp22, whereby the effect of the CN is likely to polarize the C–H bond, making for a better H-bond donor. However, the distinct orientations of the cyanopyrazole ring observed in complex structures of regioisomers AVI-4054 and AVI-5705 (Figure 3B) could indicate that favorable disruption of water networks (i.e., entropic effects) in the northern region could be as important as contact with Asp22, at least in the case of the C6 (hetero)aryl series of analogs.

**Chart 1.**
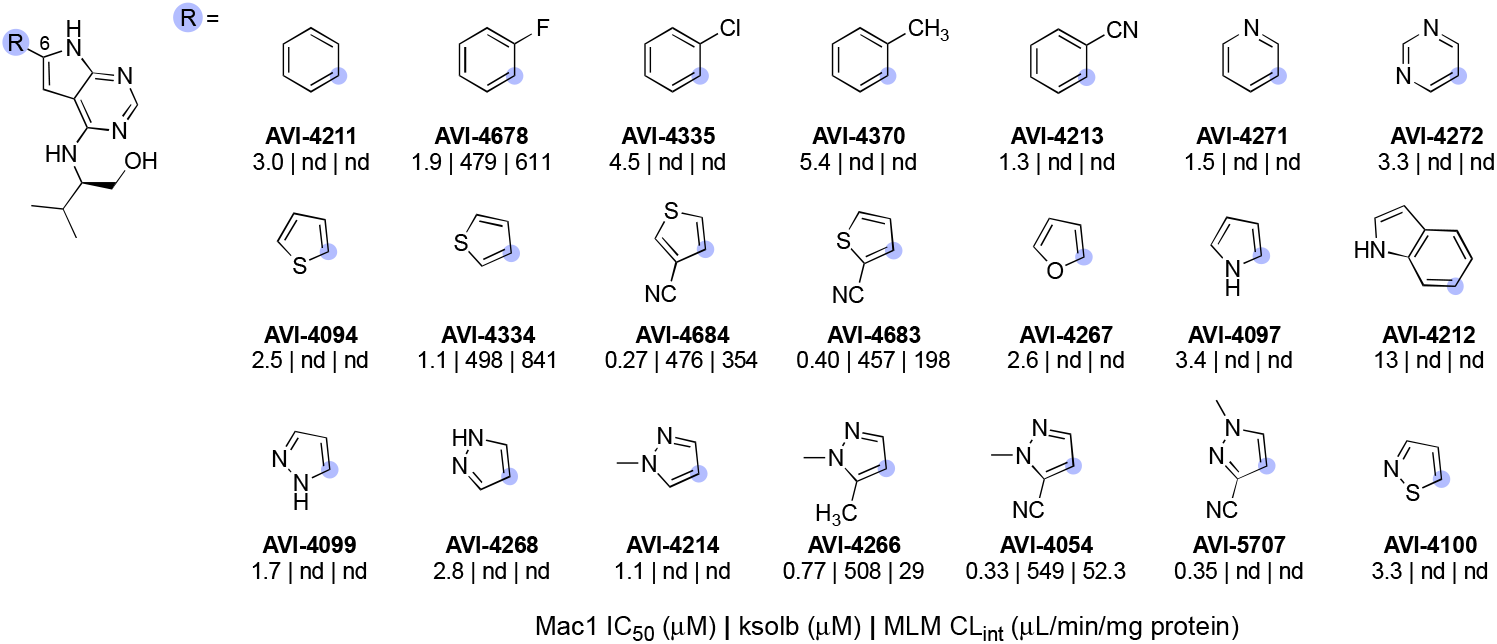
Novel at C6 of the 5-deazapurine ring. Mac1 HTRF IC_50_, kinetic solubility and mouse liver microsome clearance values are indicated below the AVI-ID. See Figure S1 for dose-response curves.

In summary, a survey of C6 heteroaryl substituents yielded analogs up to ∼30-fold more potent than **2**, confirming the northern site as a fruitful one for potency enhancement, as predicted. Conversely, the addition of C6 aryl or heteroaryl rings in **2** introduced metabolic instability to varying degrees in mouse liver microsomes (Table 2). Continuing optimization from the C6 cyanopyrazole AVI-4054, we next explored various C4 side chains to engage the ribose binding site and phosphate tunnel of Mac 1. A representative sampling of such analogs is detailed here (Chart 2) and were prepared by an analogous synthetic route wherein the cyanopyrazole C6 substituent was introduced by Suzuki coupling, followed by SNAr reactions with amine-bearing C4 sidechains (Scheme 1 and Supporting information).

**Chart 2.**
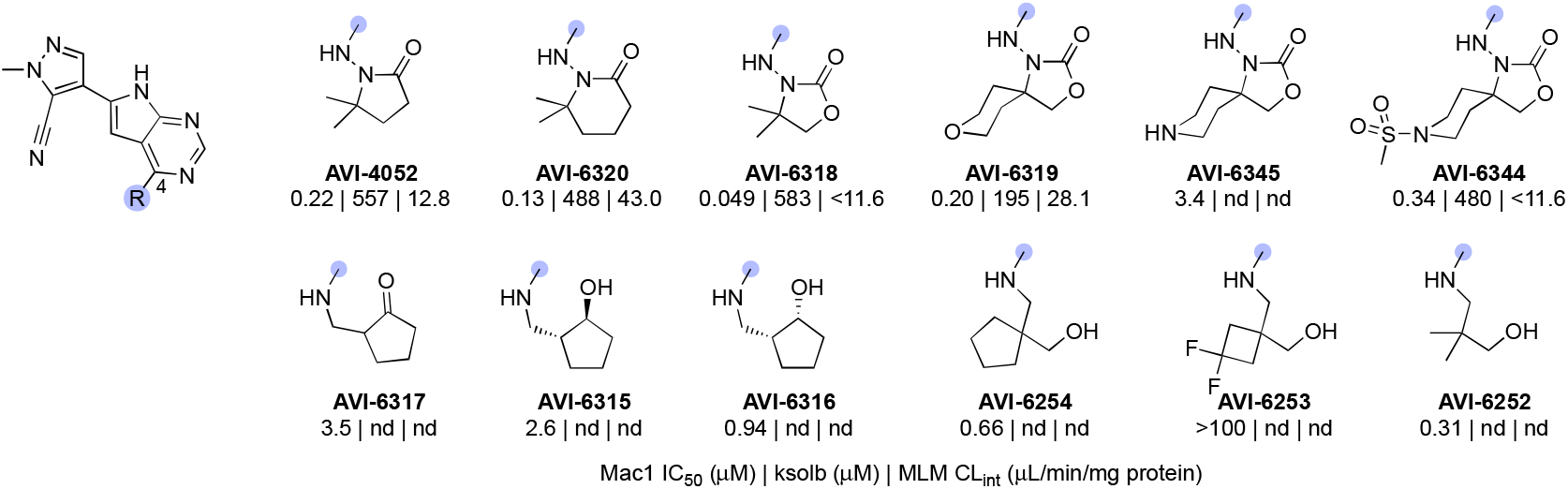
Novel analogs bearing cyanopyrazole substitution at C6 and diverse substitutions at C4 of the 5-deazapurine ring. Mac1 HTRF IC_50_, kinetic solubility values are indicated below the AVI-ID. See Figure S1 for dose-response curves.

Many of these new C4 analogs incorporated the gemdimethyl pyrrolidinone of **1**, (AVI-4052) or related piperidin-2-one (AVI-6320) or oxazolidinone sidechains (AVI-6318), including examples with spirocyclic tetrahydropyran (AVI-6319) or piperidine (AVI-63**45** and AVI-6344) rings that we predicted should bind the phosphate tunnel (Chart 2, top row). Encouragingly, these new analogs were 2–5-fold more potent that AVI-4054, with AVI**-**6318 (**3**) exhibiting an excellent Mac1 IC_50_ of 49 nM, combined with excellent solubility and mouse liver microsome stability (Table 3). The complex crystal structures of AVI-4052 and **3** revealed an analogous binding pose, the C–H bond of the pyrazole ring oriented toward Asp22 as expected, and the carbonyl of the pyrrolidinone or oxazolidinone rings forming H-bonding contact with the backbone N–H groups of Phe156 and Asp157 (Figure 4). The ∼4-fold improved potency of oxazolidinone **3** vs. AVI-4052 was gratifying and is consistent with a more basic carbonyl acceptor leading to stronger/shorter hydrogen bonds, as borne out in the respective structures (Figure 4). While less potent than **3**, spirocyclic analogs AVI-6319 and AVI-6344, as hoped, projected their distal ring systems into the phosphate tunnel of the Mac1 binding site (Figure 4). Most compelling was the piperidine sulfonamide AVI-6344, in which the sulfonamide function formed H-bond contacts to backbone residues that form similar contacts with the α phosphate of ADP-ribose substrate. This provocative binding mode of AVI-6344 hints at the potential for more productive engagement of the phosphate tunnel with further elaboration of this C4 chemotype.

**Table 3.**
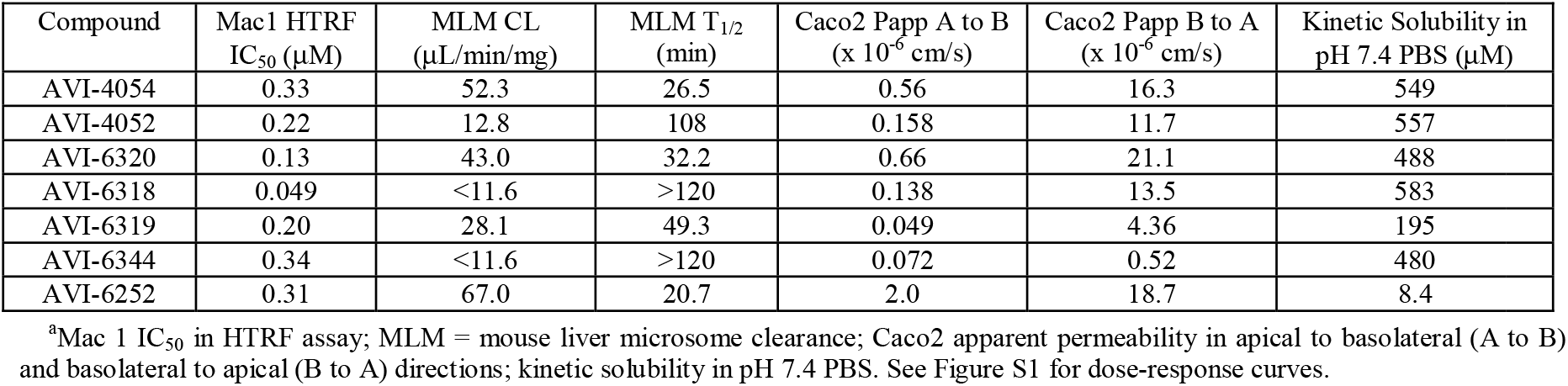
In vitro ADME data for AVI-4054 and selected additional C4-substituted analogs from Chart 2.^a^.

**Figure 4.**
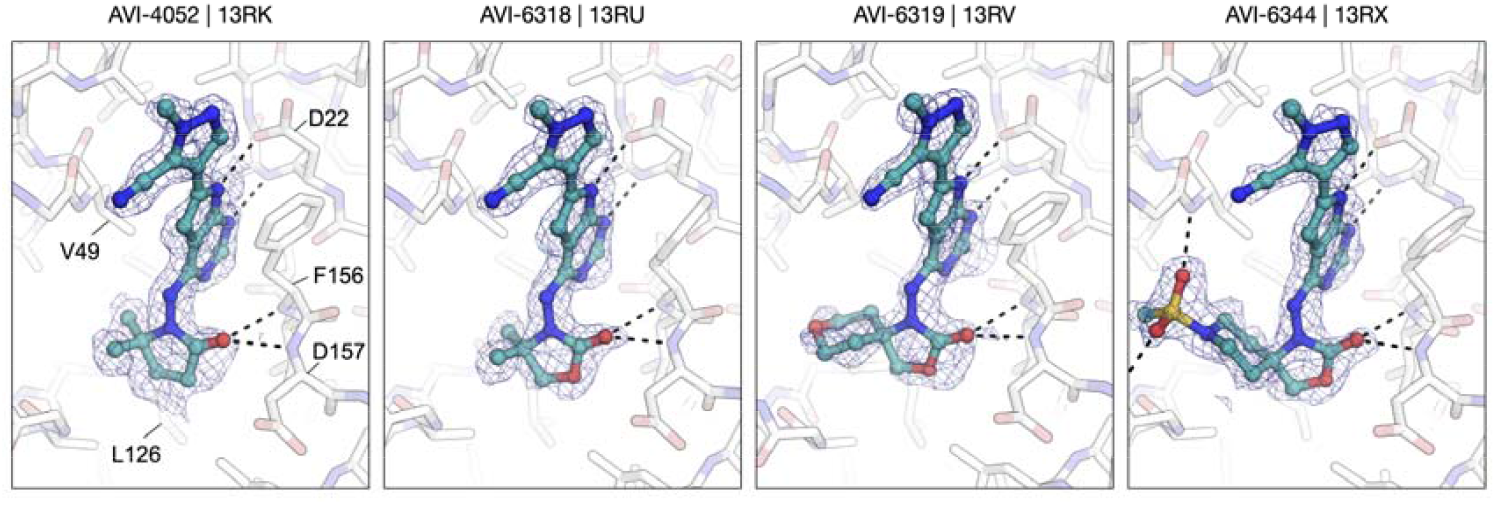
Complex structures of analogs from Chart 2 bound to Mac1. From left to right, AVI-4052, AVI-6318, AVI-6319, and AVI-6344. PanDDA event maps are contoured around ligands at 2 σ (blue mesh). Hydrogen bonds are shown with dashed black lines.

In terms of their ADME properties, oxazolidinone analogs **3**, AVI-6319, and AVI-6344 all showed notable improvements in microsome stability when compared to the progenitor C4-valinol derivative AVI-4054. On the other hand, all four compounds retained concerning Caco2 efflux ratios (ER ≥ 10), indicating that a key ADME liability of **1** was not overcome with the deazapurine scaffold, and further suggesting that recognition by P-gp was not solely associated with a C8 urea. To further differentiate the deazapurines from **1** structurally, we next explored a diverse array of C4 side chains that were either inspired or taken directly from our FrankenROCS screening effort. Among such novel C4 groups were aliphatic amines possessing small-medium ring hydrophobes combined with carbonyl or hydroxyl functions to engage the backbone amides of the ribose binding site (Chart 2, bottom row). These interactions were indeed confirmed in the complex structures of selected analogs (e.g. AVI-6317, Figure 4). The acyclic analog AVI-6252 exhibited Mac1 potency that was comparable to pyrrolidinone AVI-4052, albeit at the expense of reduced metabolic stability and aqueous solubility. While the permeability of this analog was improved over the oxazolidinones, efflux remained higher than desired (ER ∼9), and this discouraged further efforts in this vein.

In a pivotal early study in the program, we evaluated the PK profile of **1** and AVI-4052 in CD1 mice (10 mg/kg IP dose) with the goal of selecting one compound for PD studies. Consistent with their similar in vitro ADME profiles and plasma protein binding (61% and 72%, respectively), the plasma exposure profiles of the two compounds were remarably similar (Figure 5). Although we could estimate that a 100 mg/kg IP dose of AVI-4052 (blue dotted line, Figure 5) would be sufficient to retain free plasma exposure near or above Mac1 IC_50_ values for several hours, compound **1** was ultimately selected for more extensive in vivo PD studies, in large part due to its ∼10-fold greater biochemical potency. Although more potent analogs of AVI-4052 were subsequently identified (Chart 1 and 2, as detailed above), their overall similar ADME profiles, and efflux liability, discouraged the further evaluation of deazapurine analogs in vivo. Accordingly, our focus returned to the 9*H*-pyrimido[4,5-*b*]indole scaffold and lead compound **1**, with the goal identifying an orally bioavailable lead.

**Figure 5.**
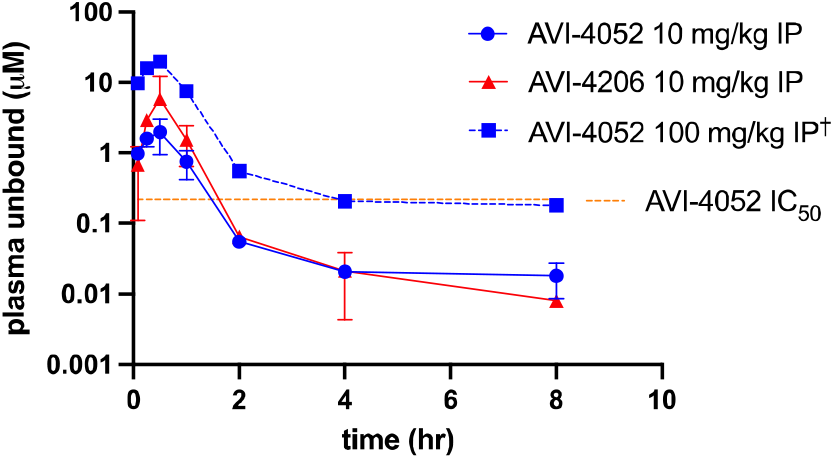
Unbound plasma exposure of **1** (AVI-4206) and AVI-4052 following a single IP dose of 10 mg/kg in mice. ^†^Extrapolation of the free plasma exposure of AVI-4052 to a 10-fold higher dose, assuming linear pharmacokinetics. The Mac1 IC_50_ value of AVI-4052 is shown as an orange dotted line.

Initial SAR leading to **1** showed that even large alkyl ureas were well tolerated as C8 substituents (e.g. cyclopentyl urea AVI-4057; Mac1 IC_50_ = 42 nM, Table 1). We considered whether further modification of the urea might abrogate recognition by P-gp. However, analogs with six-membered oxygen-bearing heterocycles (AVI-6188), or those with an extended methoxyethyl urea (AVI-6187) failed to notably reduce efflux, despite excellent potencies (IC_50_ ∼20 nM) and microsome clearance values (Table 4).

**Table 4.** Potency and in vitro ADME data for C8 urea isosteric analogs bearing pyrrolidinone or oxazolidinone at C4.^a^.

| compound | Mac1 $IC_{50}$ (μM) | MLM $T_{1/2}$ (min) | Caco2 ER | Ksolb (μM) |
| --- | --- | --- | --- | --- |
| AVI-4206 | 0.020 | >120 | 241 | 505 |
| AVI-6187 | 0.051 | >120 | 54.9 | 468 |
| AVI-6188 | 0.013 | >120 | 71.5 | 542 |
| AVI-6347 | 0.020 | >120 | 100 | 498 |
| AVI-6414 | 0.24 | 42 | 46.1 | 232 |
| AVI-6249 | 0.45 | 73.1 | 102 | 518 |
| AVI-6425 | 0.026 | 42.9 | 99.0 | 501 |

<sup>a</sup>Mac 1 $IC_{50}$ in HTRF assay; MLM = mouse liver microsome stability; ER = efflux ratio BtoA/AtoB; Ksolb = kinetic solubility in pH 7.4 PBS. See Figure S1 for dose-response curves.

We next explored urea bioisosteres in which the distal N–H function of the urea was replaced by a sulfur atom as in thiocarbamate AVI-6414, or a polarized C–H bond as in difluoroacetamide AVI-6249. These analogs were prepared with the optimized C4 oxazolidinone moiety, after we first confirmed this modification of **1** (i.e., oxazolidinone AVI-6347) retained excellent potency and microsome stability. Unfortunately, both ethyl thiocarbamate (AVI-6414) and difluoroacetamide (AVI-6249) analogs were ∼10-fold less potent, with still unacceptable efflux ratios and moderately reduced microsome stability (Table 4). Importantly however, a difluoroethylaniline analog (AVI-6425) was equipotent to ethyl ureas **1** and AVI-6347, with an IC_50_ = 26 nM. While this analog still suffered from high efflux, it nevertheless demonstrated that a C8 urea was non-essential for low nM potency.

This important finding encouraged a reevaluation of other small aliphatic C8 substituents lacking an N–H donor or carbonyl function. A pivotal set of analogs included C8 methyl, trifluoromethyl, and cyclopropyl analogs, prepared with either C4 pyrrolidinone or oxazolidinone side chains (Chart 4). We observed C8 cyclopropyl analogs AVI-6354 and AVI-6373 to be ≥10-fold more potent than isosteric C8 trifluoromethyl analogs AVI-6371 and AVI-6372 (Table 5). This implied a key difference between the more acidic C–H bonds of the cyclopropyl ring and the inversely polarized C– F bonds of the CF_3_ group. An alternate explanation involving electronic effects on the aryl π system were ruled out by similar (and 10-fold weaker) potency of C6 cyclopropyl (AVI-4769) and trifluoromethyl (AVI-3865) comparators (Chart 4 and Table 5).

**Table 5.**
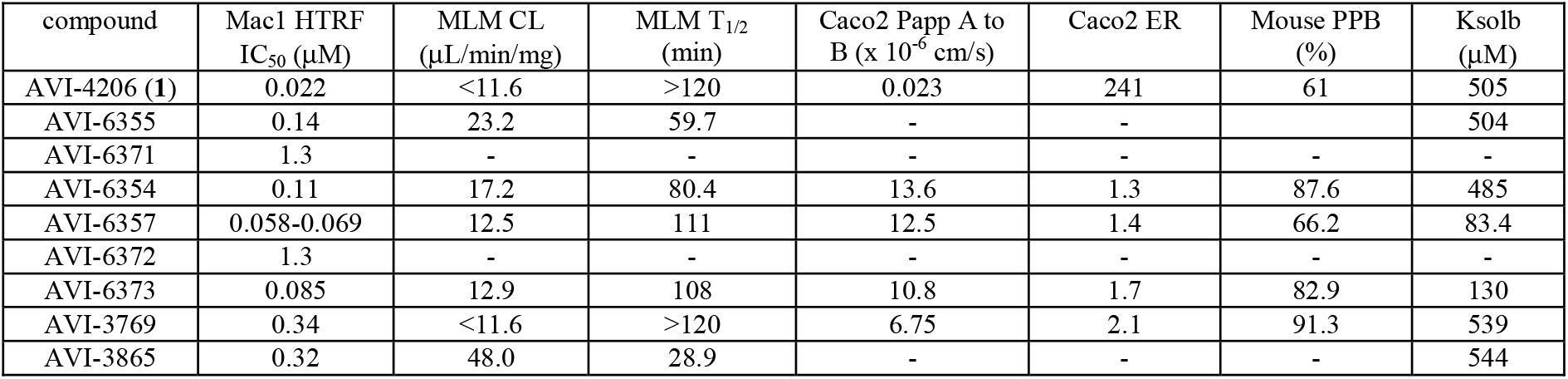
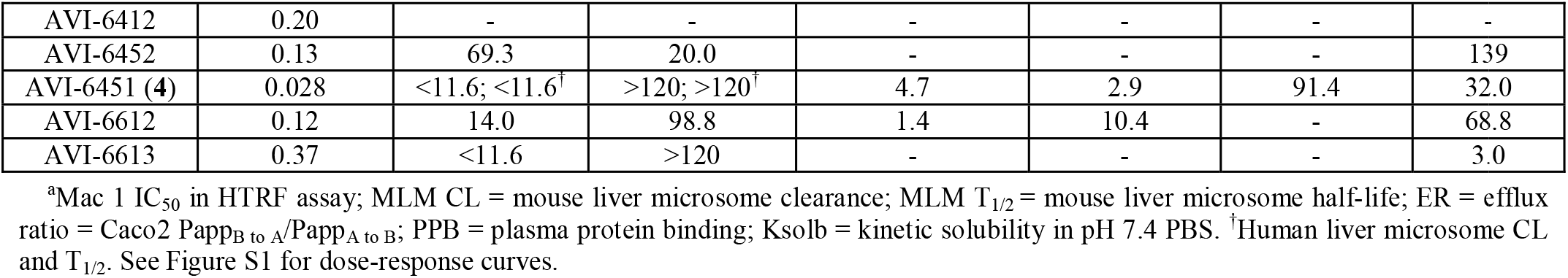
Potency and in vitro ADME data for compounds from Chart 4.^a^.

In summary, C8-cyclopropyl leads AVI-6354 and AVI-6373 exhibited potency only marginally weaker than **1**, whilst eliminating two hydrogen bond donors and reducing calculated topological polar surface area (tPSA) from 119 Å^2^ to just 69 Å^2^ (for AVI-6354). Most importantly, permeability in Caco2 monolayers was improved from 0.023 × 10^-6^ cm/s for **1** to 13.6 × 10^-6^ cm/s for AVI-6354, with the efflux ratio plummeting from >200 to 1.3 (Table 5). To improve potency and reverse a slight increase in microsome clearance, we introduced fluorine substitution at C6 or C7, yielding new benchmark analogs AVI-6451 (**4**) and AVI-6452. Interestingly, C6 fluorination in **4** produced an improved IC_50_ of ∼28 nM, essentially equipotent to **1**, while C7 fluorination in AVI-6452 had the opposite effect on potency and significantly worsened microsome stability (compared to AVI-6373). Compound **4** demonstrated low clearance (<11.6 μL/min/mg) in both mouse and human liver microsomes and good Caco2 permeability with an acceptable efflux ratio of 2.9 (Table 5). The potency and improved ADME properties of **4** led to its selection for mouse PK studies, where gratifyingly, the compound exhibited very low clearance (0.242 L/hr/kg, ∼3% of liver blood flow), a favorable mean residence time and T_1/2_ of ∼ 4 hrs, and high oral bioavailability at 61 %F (Table 6). By comparison, C7-F comparator AVI-6452 showed higher clearance, shorter MRT and T_1/2_ values that were consistent with its reduced microsome stability as compared to **4**.

**Table 6.**
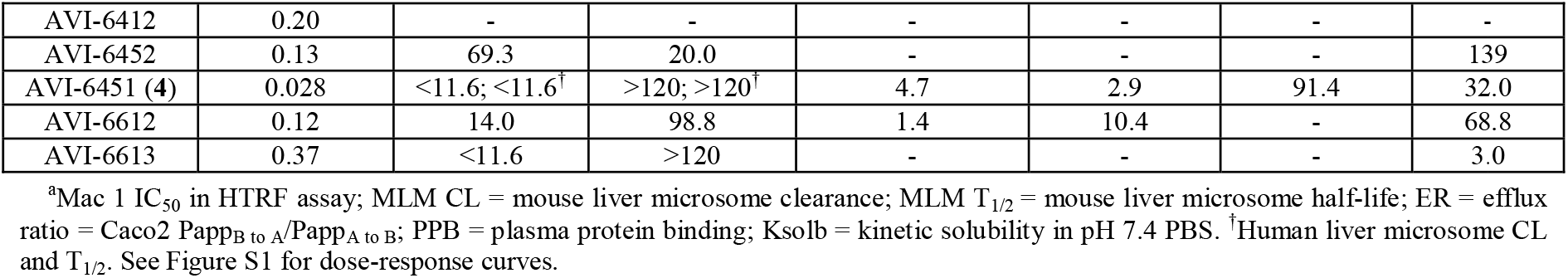
Selected PK parameters for Mac1 inhibitors following a single dose in male CD-1 mice.^a^.

To better understand the SAR of urea mimetic and non-polar C8 groups, we turned to X-ray crystallography and isothermal titration calorimetry (ITC). As noted above, the complex structure of **1** with Mac1 shows close hydrogen bonding contact between the urea function and the oxygen atoms of Asp22 in a seemingly important interaction (Figure 1). Interestingly then, in C8 difluoroacetamide AVI-6249, the amide function occupies a rotameric state in which the carbonyl π system is orthogonal to the aromatic π system, with the N–H function pointing away from Asp22, despite their being no apparent impediment to adopting a rotamer that allows contact with Asp22 (Figure 6). This suggested that H-bonding with Asp22 was perhaps not as enthalpically favorable as supposed. Furthermore, in comparing the structures of cyclopropane analog AVI-6354 and trifluoromethyl congener AVI-6372 we observed a nearly identical binding mode in which Asp22 hydrogen bonds to the 9*H*-pyrimido[4,5-*b*]indole N–H and presents its carbonyl π surface toward the C8 substituent (Figure 6). The ∼10-fold difference in IC_50_ values likely reflects a favorable electrostatic interaction of the cyclopropane C–H bonds with the π electrons of Asp22, and/or a corresponding repulsive interaction in the case of the CF_3_ substituent. Notably, the orientation of Asp22 is similar in the complex structure of AVI-6451 and distinct from that of **1**, where the carboxylate turns to face and directly engage the urea N–H donors. Thus, the crystal structures of urea, cyclopropyl, and trifluoromethyl analogs provided at least one possible rationale for C8 SAR based on purely enthalpic considerations.

**Chart 4.**
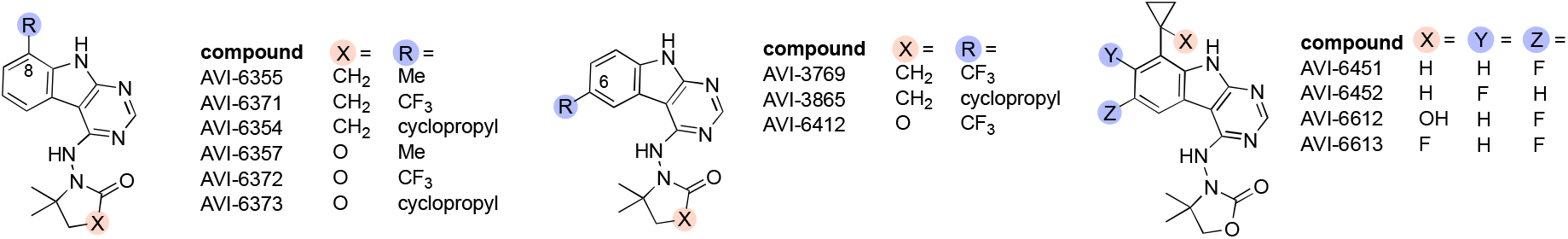
Structure of 9*H*-pyrimidoindole leads bearing optimized C4 heterocycles and non-urea substituents at C8. See Figure S1 for dose-response curves.

**Figure 6.**
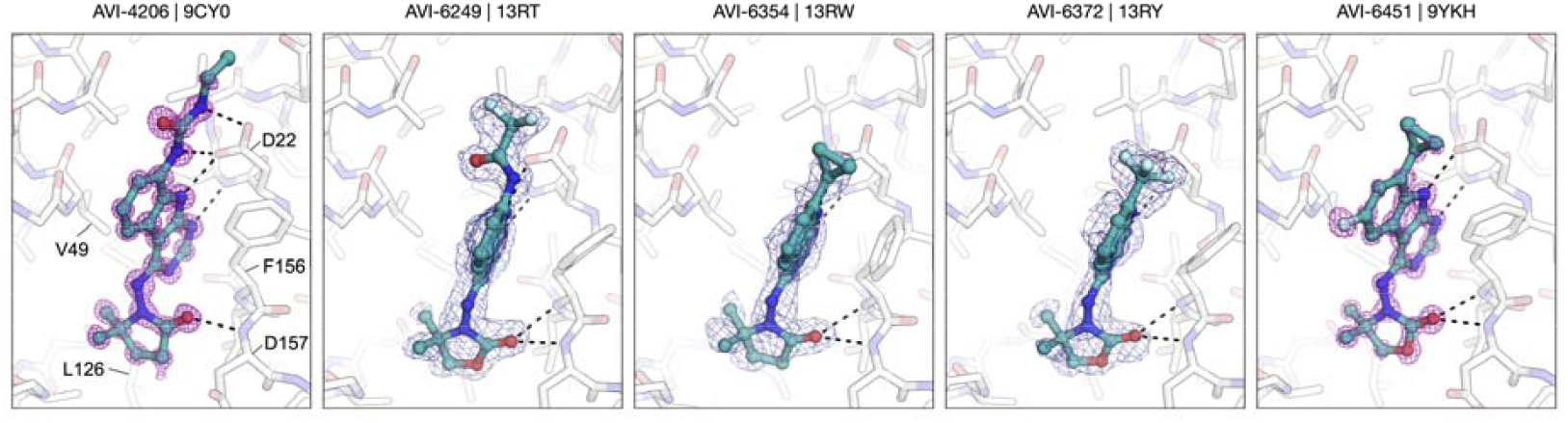
Complex structures of C8 urea and non-urea analogs bound to Mac1. From left to right, AVI-4206, AVI-6249, AVI-6354, AVI-6372, and AVI-6451 (**4**). PanDDA event maps are contoured around ligands at 2 σ (blue mesh) and F_O_-F_C_ difference electron density maps calcucalculated prior to ligand placement are contoured at 4 σ (purple mesh). Hydrogen bonds are shown with dashed black lines.

We next turned to isothermal titration calorimetry (ITC) to determine experimentally the enthalpic and entropic contributions to binding of analogs **1** and **4**; the possibly confounding effect of ring fluorination was excluded by also including cyclopropyl analog AVI-6373, which lacks the C6-fluorination of **4**. While it is most common in ITC experiments to titrate ligand into protein, we found it necessary in this case to employ reverse titration of Mac1 protein into a solution of ligands, due to a significant heat of solvation observed in control experiments wherein **4** was added to the ITC buffer without Mac1. Having demonstrated no such complications with reverse addition of protein to ligand, we were able to measure the heat released upon binding and could record accurate thermodynamic parameters for all three compounds.

We found that ITC-determined Kd values for **1** and **4** were comparable within the standard error of the measurement (44 ± 18 nM and 55 ± 11 nM respectively), and in good agreement with their Mac1 IC_50_ values determined in the HTRF assay (Table 7). The favorable effect of the C6-F substituent observed in HTRF assays was confirmed by ITC, with a ∼2-fold improvement in the measured Kd for **4** vs. des-F congener AVI-6373. The fluoro and des-fluoro congeners showed similar ΔH and ΔS values, indicating that ring fluorination did not affect enthalpic or entropic contributions to binding. Interestingly, while both **1** and **4** showed favorable ΔH and ΔS of binding, their relative contributions were distinct. Thus, C8 urea **1** exhibited a smaller enthalpy of binding that was compensated for by a significantly greater entropy of binding, as compared to **4** (Table 7). This result was some-what counter-intuitive given that the urea–Asp22 inte**rac**tion requires precise positioning of functional groups, and **in** isolation should be entropically costly. However, the pre organization of water networks in the apo vs. bound state, **as** well as desolvation energies of the ligands are also important factors to consider, if less easy to predict.

**Table 7.** Thermodynamic parameters of Mac1 binding as determined by ITC of three analogs.

| compound | K<sub>d</sub> (nM) | ΔH (kJ/mol) | ΔS (J/K•mol) |
| --- | --- | --- | --- |
| AVI-4206 (**1**) | 44 ± 18 | -14.02 ± 0.41 | 94.6 ± 4.3 |
| AVI-6451 (**4**) | 55 ± 11 | -27.00 ± 0.92 | 48.5 ± 3.7 |
| AVI-6373 | 83 ± 10 | -24.29 ± 0.65 | 54.0 ± 3.5 |

**Table 8.** Calculated thermodynamic parameters of small molecule desolvation, determined with DFT wB97X-D|def2SVP and SMD(water)-wB97X-D|def2SVP level of theory.

| compound | $\Delta G$<br>(kJ/mol) | $\Delta H$<br>(kJ/mol) | $T\Delta S$<br>(kJ/mol) |
| --- | --- | --- | --- |
| AVI-4206 (**1**) | 117.11 | 115.60 | -1.51 |
| AVI-6451 (**4**) | 78.29 | 75.79 | -2.51 |

To gain further insight, we employed density function**al** theory (DFT) to calculate the energetics of desolvation for compounds **1** and **4** using the w**B9**7X-D/def2SVP//SMD(water)-wB97X-D/def2SVP level of theory. The results indicated that desolvation of **1** is enthalpically much more costly than for **4**, presumably due enthalpically favorable solvent interactions with **1**, particularly at the polar urea function. This penalty is ultimately overcome by favorable enthalpic interactions with Mac1 upon binding (including with Asp22) and by the entropic gains of net release of water to bulk solvent. Together the ITC and DFT data suggest that while binding of **1** and **4** to Mac 1 is downhill both enthalpically *and* entropically, the binding of **1** is driven more by the entropic term (and likely by desolvation of the active site), while binding of **4** is more enthalpically driven, and likely with fewer net water molecules released to the bulk solvent.

Previously, we described safety screening data for **1** against a panel of kinases, proteases, and enzyme/receptors, and found that at a concentration of 10 μM, **1** showed minimal interaction (<50% response) with *any* of the ∼300 off-targets.^12^ Here, we present new data for **4** alongside the previously reported results for **1** in the enzyme/receptor panel specifically (SafetyScreen44, performed by Eurofins Panlabs, Inc.). At the 10 μM concentration employed, compound **4** showed ≥50% response at 10 μM for only three offtargets, namely phosphodiesterase PDE4D2 (51%), and the adenosine A_2A_ (56%) and serotonin 5-HT_2B_ receptors (56%) while overall retaining a favorable safety profile (Figure 7A). Against a panel of CYP enzymes, compound **4** at 10 μM did show more potent inhibition of CYP2C8 and CYP2C19 compared to **1**, identifying an area for further optimization (Figure 7B).

**Figure 7.**
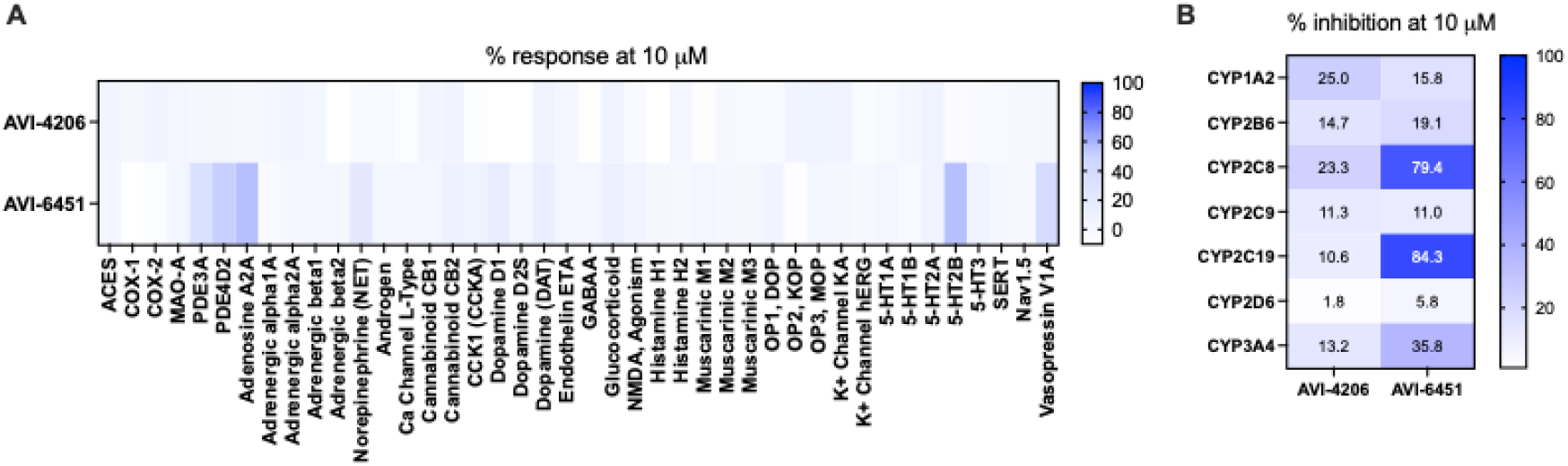
Heatmaps showing the activity of 10 μM AVI-4206 and AVI-6451 in A) a Eurofins SafetySceen 44 panel of potential enzyme and receptor off targets and B) a panel of CYP isoforms.

Having identified a permeable and orally bioavailable Mac1 inhibitor with excellent potency and selectivity, we set about to explore the efficacy of **4** in cellular and in vivo mouse models. As we and others have noted, the antiviral effects of Mac1 inhibitors is highly dependent on the cell model, with human airway organoid (HAO) and monocyte-derived macrophage (MDM) models showing the strongest Mac1 phenotype in our program.^12^ Here we employed the MDM model to demonstrate the superior cellular potency of **4**, which was consistent with its higher cellular permeability in Caco2 assays. Thus, SARS-CoV-2 infection experiments were performed using MDM cells differentiated from CD14-positive monocytes isolated from human peripheral blood mononuclear cells (PBMCs). In these experiments, the presence of infectious viral particles is evaluated by plaque assay using Vero-TMPRSS2 cells. We found that both **1** and **4** exhibited antiviral effects in MDMs, but whereas **1** required μM concentrations to show antiviral effects in both the HAO and MDM models,^12^ compound **4** was effective at mid-low nM concentrations (Figure 8C).

**Figure 8.**
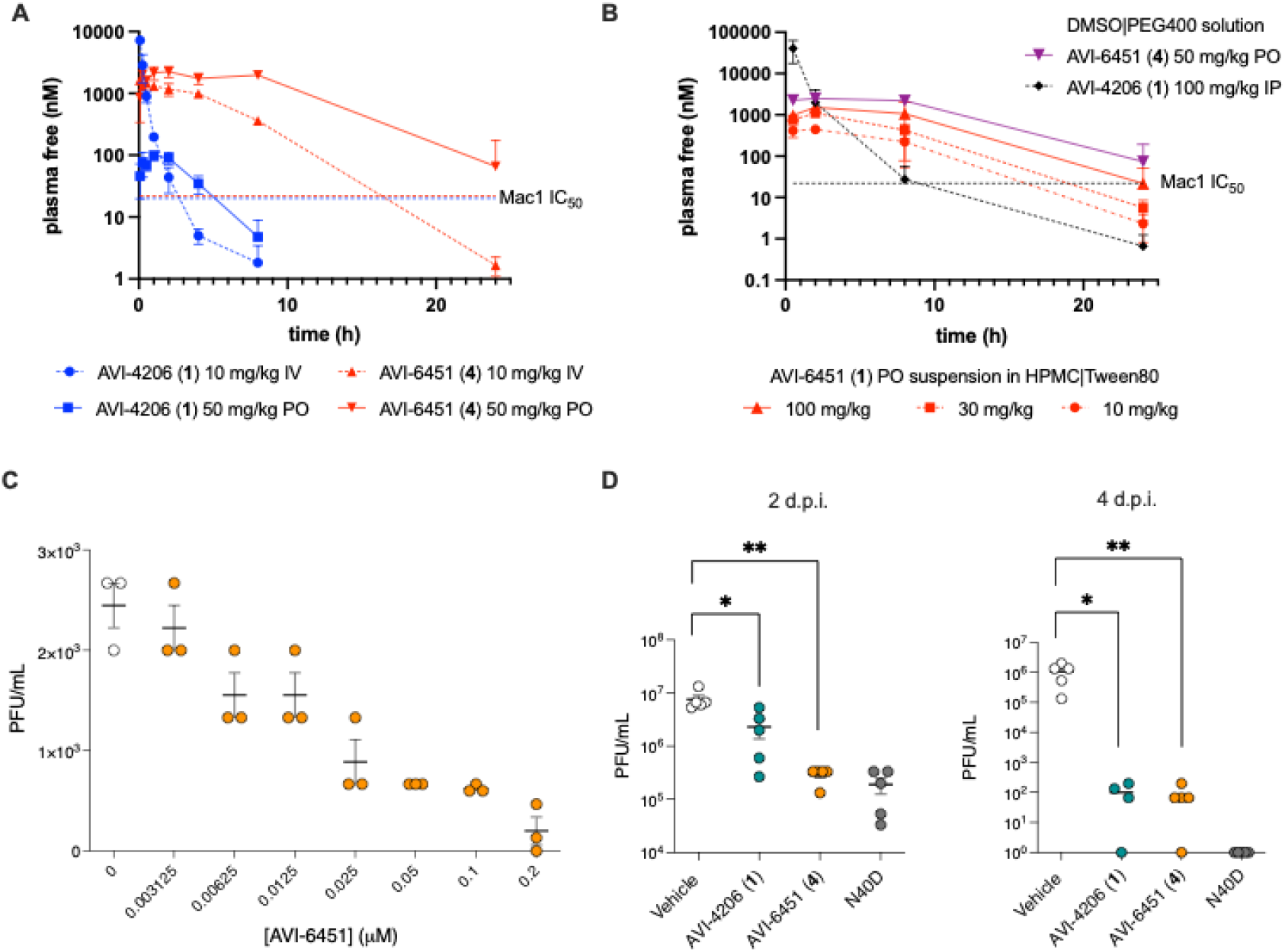
AVI-6451 shows enhanced PK and cellular and in vivo antiviral efficacy compared to AVI-4206. Panel **A.** Single-dose mouse PK experiments demonstrate the reduced clearance and higher oral exposure of AVI-6451 (**4**, in red) compared to AVI-4206 (**1**, in blue). Compounds formulated in 10% DMSO|50% PEG400|40% (20% HP-β-CD in water). Panel **B**. Dose escalation study of **4** formulated as a suspension in 1% (hydroxypropyl)methyl cellulose|1% Tween 80 in water. Data from prior studies with **1** and **4** in DMSO|PEG400|HP-β-CD are shown for comparison. **C**. Dose-dependent antiviral activity of AVI-6451 (**4**, in orange) in monocyte derived macrophages infected with SARS-CoV-2 MA-WA1. panel **D**. Viral load in the lungs of wild-type mice (N=10/group) at days 2 and 4 post-infection with 1 × 10^4^ PFU of SARS-CoV-2 MA-WA1. Mice were treated from day −1 to day 3 with vehicle, with 100 mg/kg AVI-4206 IP BID, or 100 mg/kg AVI-6451 PO QD. Viral load of mice infected with SARS CoV-2 MA-WA1 Mac1 N40D (mutant lacking Mac1 activity) are shown for comparison.

As noted above, the superior oral PK profile of **4** results in >100-fold higher plasma exposure than **1** following a single oral dose (Table 6). Indeed, the free unbound plasma concentration of **4** remained well above its Mac1 IC_50_ a full 24 hrs after a 50 mg/kg oral dose (Figure 8A), suggesting the potential for once-daily oral administration. To select dosing levels for PD models, a dose escalation PK study of **4** in mice was performed in an oral suspension comprising 1% (hydroxypropyl)methyl cellulose and 1% Tween 80 in water. At the highest 100 mg/kg oral dose in this new formulation, free plasma concentrations remained above Mac1 IC_50_ values for nearly 24 hours and was well tolerated (Figure 8B). On this basis, we selected once-daily oral administration of **4** at 100 mg/kg for evaluation in a SARS-CoV-2 infection model in wild type mice. As a positive control, we chose **1** at 100 mg/kg BID by the IP route as this dose had previously proven effective in multiple in vivo studies of **1** using a lethal K18-hACE2 mouse model. A separate group of wild-type mice were infected with SARS-CoV-2 MA-WA1 Mac1 N40D, a mutant virus that lacks Mac1 enzymatic activity.

As noted above, the superior oral PK profile of **4** results in >100-fold higher plasma exposure than **1** following a single oral dose (Table 6). Indeed, the free unbound plasma concentration of **4** remained well above its Mac1 IC_50_ a full 24 hrs after a 50 mg/kg oral dose (Figure 8A), suggesting the potential for once-daily oral administration. To select dosing levels for PD models, a dose escalation PK study of **4** in mice was performed in an oral suspension comprising 1% (hydroxypropyl)methyl cellulose and 1% Tween 80 in water. At the highest 100 mg/kg oral dose in this new formulation, free plasma concentrations remained above Mac1 IC_50_ values for nearly 24 hours and was well tolerated (Figure 8B). On this basis, we selected once-daily oral administration of **4** at 100 mg/kg for evaluation in a SARS-CoV-2 infection model in wild type mice. As a positive control, we chose **1** at 100 mg/kg BID by the IP route as this dose had previously proven effective in multiple in vivo studies of **1** using a lethal K18-hACE2 mouse model. A separate group of wild-type mice were infected with SARS-CoV-2 MA-WA1 Mac1 N40D, a mutant virus that lacks Mac1 enzymatic activity In the experiment, wild-type mice (N=10 per group) were infected with 1 × 10^4^ PFU of SARS-CoV-2 mouse-adapted (MA-WA1) strain and treated from day –1 to day 3 with either vehicle, compound **1** BID at 100 mg/kg via the IP route (DMSO|PEG|HP-βCD formulation) or with compound **4** QD at 100 mg/kg via oral gavage (as a suspension in 1%HPMC|1%Tween80). Viral load in lungs was assessed at 2 and 4 days post-infection (dpi) by plaque assay. Consistent with our prior studies of **1** in the K18-hACE2 mouse model, compound **1-**treated mice showed significantly reduced viral titers in lung compared to vehicle controls at 2 dpi and more dramatically at 4 dpi (Figure 8D). Notably, at roughly half the total dose, and with PO rather than IP administration, compound **4** demonstrated a more significant reduction in viral load over control at 2 dpi, and an excellent ∼4-log reduction in viral titers over control by 4 dpi. Also of note, the effect of once-daily PO administration of **4** in this model was comparable at 2 dpi to infection with the N40D Mac1 mutant virus, the first time we have observed small molecule inhibition of Mac1 to equal the pharmacodynamic effect of the control virus with inactivated Mac1.

In summary, two chemical series were investigated in an effort to discover what is, to our knowledge, the first orally bioavailable and efficacious inhibitor of the SARS-CoV-2 ribosylhydrolase Mac1. As an improved *in vivo* tool compound and drug lead, compound **4** presents several advantages over **1** with regard to future studies of macrodomains as antiviral drug targets. These include suitability for once-daily oral administration in mice as an oral suspension. This will enable more convenient dosing regimens in further studies of **4**, alone or in combination with other clinical or pre-clinical antiviral agents, both in SARS-CoV-2 and potentially other coronavirus infection models. Moreover, the improved cellular permeability and potent antiviral effects of **4** revealed in the monocyte-derived macrophage assay nominate the compound for future cellular studies aimed an untangling the role of Mac1 in the innate immune response to viral infection, an area of active inquiry. With regard to the further development of antiviral drug leads from this scaffold, efforts are well underway and will be reported in due course.

## EXPERIMENTAL PROCEDURES

### Homogenous time-resolved fluorescence (HTRF) Assay of Mac1 Inhibition

Binding of AVI analogs to Mac1 was assessed by the displacement of an ADPr conjugated biotin peptide from His6-tagged protein using an HTRF-technology based screening assay Mac1 protein expressed and purified as previously described.^12,14^ Compounds were dispensed into ProxiPlate-384 Plus (PerkinElmer) assay plates using an Echo 650 Liquid Handler (Beckman Coulter). Binding assays were conducted in a final volume of 16 μl with 12.5 nM NSP3 Mac1 protein, 200 nM peptide ARTK(Bio)QTARK(Aoa-RADP)S (Cambridge Peptides), 1:20000 Anti-His6-Eu^3+^ cryptate (HTRF donor, PerkinElmer AD0402) and 1:500 Streptavidin-XL665 (HTRF acceptor, PerkinElmer 610SAXLB) in assay buffer (25 mM 4-(2-hydroxyethyl)-1-piperazine-1-ethanesulfonic acid (HEPES) pH 7.0, 20 mM NaCl, 0.05% bovine serum albumin and 0.05% Tween-20). Assay reagents were dispensed manually into plates using an electronic multichannel pipette. Mac1 protein and peptide were dispensed and preincubated for 30 min at room temperature before HTRF reagents were added. Fluorescence was measured following a 1-hour incubation at room temperature using a Perkin Elmer EnVision 2105-0010 Dual Detector Multimode microplate reader with dual emission protocol (A = excitation of 320 nm, emission of 665 nm, and B = excitation of 320 nm, emission of 620 nm). Compounds were tested in triplicate in a 14-point dose response. Raw data were processed to give an HTRF ratio (channel A/B × 10,000), which was used to generate IC_50_ curves. The IC_50_ values were determined by nonlinear regression using GraphPad Prism v.10.0.2 (GraphPad Software, CA, USA). Data presented as mean ± SD of three technical replicates

### Macrophage Infection Assay

Macrophages were generated by culturing monocyte fractions isolated from peripheral blood mononuclear cells (PBMCs) (Vitalant, CA, USA) using CD14 negative selection (STEMCELL Technologies, Vancouver, Canada) in serum-free medium supplemented with 50ng/mL M-CSF for 8 days. The resulting monocyte-derived macrophages (MDMs) were pre-treated with specified concentrations of AVI-6451 in the presence of 10 ng/mL IFN-γ for 2 hours, followed by infection with SARS-CoV-2 isolate USA-WA1/2020 at a multiplicity of infection (MOI) of 2 for 2 hours. After viral exposure, cells were thoroughly washed three times with phosphate-buffered saline (PBS) to remove unbound virus. The cells were then maintained in medium containing the respective compounds and 10 ng/mL IFN-γ for an additional 18 hours. Viral particle production was subsequently quantified by plaque assay. Data analysis was carried out using GraphPad Prism version 10.2.0 (GraphPad Software, CA, USA).

### Pharmacokinetic Studies

Procedures and data for PK studies of AVI-4052 and AVI-4206 (**1**) were reported previously. Mouse pharmacokinetic studies of AVI-6451 (**4**) and AVI-6452 with IV (10 mg/kg) or PO (50 mg/kg) dosing (Table 6, Figure 8, and Supplementary Tables S1 and S2) were performed in male CD1 mice (n = 9 per group) with a formulation of 10% DMSO/50% PEG400/40% of a 20% HP-β-CD solution in water. Dose escalation studies with **4** (Figure 8C) were performed in CD1 mice (n = 3) with administration by oral gavage using a suspension formulation of 1% (hydroxypropyl)methyl cellulose/1% Tween 80 at 1, 3, or 10 mg/mL for the 10, 30, and 100 mg/kg dosing levels, respectively. Animals were restrained manually at designated timepoints and ca. 110 μL of blood (30 μL in dose-escalation study with fewer mice/samples) was taken into K_2_EDTA tubes via facial vein. The blood samples were collected on ice and centrifuged (2000 g, 5 min) to obtain plasma samples within 15 minutes post sampling. Three blood samples were collected from each mouse; three samples were collected at each time point. Plasma samples (20 μL) were diluted into 200 μL acetonitrile with internal standard (glipizide, 60 ng/mL) were analyzed by LC-MS/MS (triple quad 6500+) and quantified by comparison with a standard curve (1.0-3000 ng/mL) generated from authentic analyte. Data was processed by Phoenix WinNonlin (version 8.3); samples below limit of quantitation were excluded in the PK parameters and mean concentration calculation.

### Pharmacodynamic Study in Mice

C57BL/6J wild-type mice were housed in the Gladstone Animal Facility. Mice aged 6–8 weeks were randomLy assigned to experimental groups and transferred to the ABSL-3 facility at the Gladstone Institutes. Mouse-adapted (MA) SARS-CoV-2 strains, MA-SARS-CoV-2 (Spike: Q498Y/P499T) and MA-SARS-CoV-2 Mac1 N40D, were generated using pGLUE, as previously described^15^. Mice were infected intranasally with MA-SARS-CoV-2 (1 × 10□PFU in 40 μL) under anesthesia induced by intraperitoneal injection of ketamine (150 mg/kg) and xylazine (10 mg/kg), and assigned to the following treatment groups: vehicle (PO, once daily), AVI-4206 (100 mg/kg, IP, twice daily), or AVI-6451 (100 mg/kg, PO, once daily). AVI-4206 was formulated in with a formulation of 10% DMSO/50% PEG400/40% of a 20% HP-β-CD solution in water for IP injection while AVI-6451 was administered by oral gavage as a suspension in 1% (hydroxypropyl)methyl cellulose and 1%Tween 80. A control group was infected with MA-SARS-CoV-2 Mac1 N40D under identical conditions. Compound administration began one day prior to infection (day −1) and continued through 3 days post-infection. Mice were euthanized at 2 and 4 days post-infection. Lungs were harvested and homogenized using a bead homogenizer (zirconium bead tubes pre-filled). Viral titers in lung homogenates were quantified via plaque assay. VeroE6-ACE2-TMPRSS2 cells were seeded in 12-well plates and incubated overnight at 37°C with 5% CO□. Serial dilutions (10□^1^ to 10□) of lung homogenates were prepared in serum-free DMEM and added to the cells for 1 hour to allow viral adsorption. After adsorption, cells were overlaid with 2.5% Avicel (DuPont) in complete DMEM and incubated for 72 hours. The overlay was then removed, cells were fixed with 10% neutral-buffered formalin for 1 hour, stained with crystal violet, and plaques were counted to determine plaque-forming units (PFU).

### Isothermal Titration Calorimetry

All ITC titrations were performed on an Affinity ITC Low Volume instrument (TA Instruments, WatersTM). Reactions were performed at 25°C in 20 mM HEPES (pH 7.5) and 150 mM NaCl. Titrations were performed with 25 µM of ligand in the sample cell and 300 µM Mac1 in the injection syringe using 20 injections of 2.5 µL each at 200-second intervals with a 300-second initial baseline and 200-second post-injection baseline. Data was analyzed using the NanoAnalyze software v4.0.2.0 (TA Instruments, WatersTM). Thermograms were integrated and the normalized binding enthalpies fitted to an independent binding model (4-variable nonlinear least squares fit). Thermodynamic parameters were then reported as the average and standard deviation obtained through five independent titrations.

### Crystallography

The P43 crystals of SARS-CoV-2 Mac1 were grown from a solution containing 28% PEG 3000 and 100 mM CHES pH 9.5 as described previously.^8,13^ Ligands were soaked into crystals to a final concentration of 10 mM using acoustic dispensing with an Echo 650 liquid handler.^16^ Crystals were vitrified in liquid nitrogen without additional cryoprotection and X-ray diffraction data were collected at beamline 8.3.1 of the Advanced Light Source (ALS) and beam lines 9-2 and 12-2 of the Stanford Synchrotron Radiation Lightsource (SSRL). Data were indexed, integrated and scaled with XDS^17^ and merged with Aimless^18^. Data collection statistics are provided in Table S3. Because several ligands bound with relatively low occupancy (<50%), we modeled ligands using PanDDA^19^ event maps as described previously^8,13^ using coordinates and restraints generated with phenix.elbow^20^. Coordinates were refined as a multi-state model with both apo and ligand-bound states using phenix.refine^21^ as described previously.^8,13^ As we previously reported for AVI-4206^12^, low nanomolar Mac1 ligands dissolved the P43 crystals, presumably because a crystal contact is displaced when ligands bind to the adenosine site of chain B. To obtain a higher occupancy structure of AVI-6451, we co-crystallized Mac1 using an alternative construct that crystallizes in P21 or C2.^22^ The Mac1-AVI-6451 complex was prepared by incubating protein (30 mg/ml, 1.6 mM) and compound (3.2 mM from a 100 mM stock in DMSO) at room temperature for five minutes. Co-crystals were grown using sitting drop vapor diffusion with drops containing 200 nl protein:AVI-6451 and 200 nl reservoir solution (200 mM lithium acetate and 20% PEG 3350). Diffraction data were collected at ALS and processed as described above. Phases were obtained by molecular replacement using Phaser^23^ and chain A of 7KQO as the search model. The initial model was improved by cycles of model building with COOT^24^ and refinement with phenix.refine. Data collection and refinement statistics for the Mac1:AVI-6451 complex are reported in Supplemtary Dataset S1. Crystallographic coordinates and structure factor intensities have been deposited in the PDB with the following accessing codes: 13RI, 13RJ, 13RK, 13RL, 13RM, 13RN, 13RO, 13RP, 13RQ, 13RR, 13RS, 13RT, 13RU, 13RV, 13RW, 13RX, 13RY and 9YKH.

### Computational Density Functional Theory (DFT) Calculations

All computations were performed using Gaussian16 (Revision C.01). Geometry optimizations, force constants, and subsequent vibrational frequencies for compounds AVI-4206 and AVI-6451 were performed using the wB97X-D functional^25^ and def2SVP basis set.^26^ Gas-phase calculations were carried out using this level of theory. Solvation effects in water (epsilon=78.3553) were evaluated using the same level of theory in conjunction with the SMD implicit solvation model^27^ (SMD(water)-wB97X-D|def2SVP). All calculations were performed at the default temperature (298K) and pressure (1 atm). The thermodynamic parameters were determined by the difference in electron energy (EE) with the appropriate correction term between the gas phase and implicit solvent model systems (= EE + Thermal Free Energy correction, = EE + Thermal Enthalpy Correction, = Entropy). The desolvation terms were determined based on the following equations:

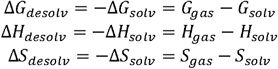

All calculations were performed utilizing 30 CPUs, made available through UCSF’s Wynton HPC cluster.

### ADME Assays

In vitro ADME assays were performed at Quintara Discovery (Hayward, CA) using their standard experimental procedures and controls. Additional detail is provided in the Supporting Information File.

### Safety Panel

The SafetyScreen44 receptor and ion channel offtarget screen was performed by Eurofins Panlabs, Inc. Additional detail is provided in the Supporting Information File.

### Synthesis

Unless otherwise noted all chemical reagents and solvents used are commercially available. Air and/or moisture sensitive reactions were carried out under an argon atmosphere in oven-dried glassware using anhydrous solvents from commercial suppliers. Solvent removal was accomplished with a rotary evaporator at ca. 10-50 Torr. Microwave reactions were carried out in a CEM Discover microwave reactor. Chromatography was carried out using Isolera Four or CombiFlash NextGen 300 flash chromatography system with SiliaSep silica gel cartridges from Silicycle. Reverse phase chromatography was carried out on either, (1) Waters 2535 Separation module with Waters 2998 Photodiode Array Detector. Separations were carried out on XBridge Preparative C18, 19 x 50 mm column at ambient temperature using a mobile phase of water-acetonitrile containing a constant 0.1% formic acid or (2) Gilson GX-281 instrument, separations using Xtimate Prep C18, 21.2 x 250mm, 150Å, 10μm particle size column at ambient temperature using a mobile phase of acetonitrile-water with 10 mM ammonium bicarbonate. LC/MS data were acquired on either (1) Waters Acquity UPLC QDa mass spectrometer equipped with Quaternary Solvent Manager, Photodiode Array Detector and Evaporative Light Scattering Detector. Separations were carried out with Acquity UPLC® BEH C18 1.7 μm, 2.1 x 50 mm column at 25 oC, using a mobile phase of water-acetonitrile containing a constant 0.1 % formic acid or (2) Agilent 1200 Infinity LC with an Agilent 1956 single quadrupole MS using electrospray ionization: Column: SunFire C18 (4.6x 50 mm, 3.5 μm), Mobile phase: H_2_O (10 mmol NH_4_HCO_3_) (A) / CH_3_CN (B) at 50 ºC with Photodiode Array Detector. Chemical shifts are reported in δ units (ppm). NMR spectra were referenced relative to residual NMR solvent peaks. Coupling constants (*J*) are re orted in hertz (Hz). NMR spectra were recorded either on Bruker 500 MHz or Bruker Avance III HD 400 MHz spectrometer.

### Synthesis of 1 8-amino-9H-pyrimido[4,5-b]indol-4-yl)amino)pyrrolidin-2-one (AVI-1499)

A mixture of 4-chloro-9H-pyrimido[4,5-b]indol-8-amine (28 mg, 0.13 mmol) and 1-aminopyrrolidin-2-one hydrochloride (35 mg, 0.26 mmol) in isopropanol/water (10: 1, 1.1 mL) were heated to 100 oC for 18 h. The reaction mixture was filtered, the residue was washed with ethyl acetate and dried to obtain 28 mg (77%) of AVI-1499 as brown solid. ^1^H NMR (DMSO-d6, 400 MHz) δ 12.99 (br s, 1H), 8.62 (s, 1H), 7.92 (br d, 1H, *J*=7.5 Hz), 7.27 (t, 1H, *J*=7.9 Hz), 7.05 (br d, 1H, *J*=7.5 Hz), 3.70 (br t, 2H, *J*=6.9 Hz), 2.44-2.53 (m, 2H), 2.20 (br t, 2H, *J*=7.4 Hz). LCMS (ESI): m/z= 283 (M+H)+

### Synthesis o -(4-((2-oxopyrrolidin-1-yl)amino)-9H-pyrimido[4,5-b]indol-8-yl)acetamide (AVI-1501)

To a solution of 1-((8-amino-9H-pyrimido[4,5-b]indol-4-yl)amino)pyrrolidin-2-one (15 mg, 0.053 mmol) and triethylamine (0.015 mL, 0.11 mmol) in THF (1 mL), was added acetyl chloride (0.004 mL, 0.056 mmol). After stirring at 65 °C for 3 h, the reaction mixture was purified by reverse phase chromatography (water/acetonitrile/0.1% formic acid) to obtain 9 mg (50%) **AVI-1501** formate, as a white solid. ^1^H NMR (METHANOL-d_4_, 400 MHz) δ 8.39 (s, 1H), 7.90 (d, 1H, *J*=7.8 Hz), 7.45 (d, 1H, *J*=7.8 Hz), 7.19-7.21 (m, 1H), 3.81-3.85 (m, 2H), 2.59-2.63 (m, 2H), 2.27-2.31 (m, 5H). LCMS (ESI): m/z= 325 (M+H)^+^

### Synthesis of 1-ethyl-3-(4-((2-oxopyrrolidin-1-yl)amino)-9H-pyrimido[4,5-b]indol-8-yl)urea formic acid salt (AVI-1500)

To a solution of 4-chloro-9H-pyrimido[4,5-b]indol-8-amine (50 mg, 0.23 mmol) and triethylamine (0.064 mL, 0.46 mmol) in THF (2 mL), was added ethyl isocyanate (0.018 mL, 0.23 mmol). After stirring at 65 °C for 18 h, the reaction mixture was filtered. The residue was washed with ethyl acetate and dried to obtain 50 mg of 1-(4-chloro-9H-pyrimido[4,5-b]indol-8-yl)-3-ethylurea as a white solid that was used in the next step without further purification. ^1^H NMR (DMSO-d_6_, 400 MHz) δ 12.39 (br s, 1H), 8.80 (s, 1H), 8.43 (s, 1H), 7.96 (d, 1H, *J*=7.6 Hz), 7.72 (d, 1H, *J*=7.8 Hz), 7.35 (t, 1H, *J*=7.9 Hz), 6.38 (s, 1H), 3.18-3.21 (m, 2H), 1.12 (t, 3H, *J*=7.2 Hz). LCMS (ESI): m/z= 290, 292 (M+H)^+^

A mixture of 1-(4-chloro-9H-pyrimido[4,5-b]indol-8-yl)-3-ethylurea (26 mg, 0.09 mmol) and 1-aminopyrrolidin-2-one hydrochloride (25 mg, 0.18 mmol) in isopropanol/water (10:1, 1.1 mL) were heated to 100 °C for 18 h. The reaction mixture was purified by reverse phase chromatography (water/acetonitrile/0.1% formic acid) to obtain 8 mg (20%) **AVI-1500** formate as a white solid. ^1^H NMR (DMSO-d_6_, 400 MHz) δ 11.75 (br s, 1H), 9.31 (s, 1H), 8.53 (s, 1H), 8.41 (s, 1H), 7.97 (d, 1H, *J*=7.8 Hz), 7.63 (d, 1H, *J*=7.8 Hz), 7.20 (t, 1H, *J*=7.9 Hz), 6.37 (br s, 1H), 3.70 (t, 2H, *J*=7.1 Hz), 3.17-3.20 (m, 2H), 2.39-2.41 (m, 2H), 2.12-2.16 (m, 2H), 1.09-1.13 (m, 3H). ^13^C NMR (DMSO-d_6_, 100 MHz) δ 173.5, 156.2, 155.9, 155.5, 155.0, 128.5, 125.5, 121.3, 120.3, 117.1, 116.7, 96.4, 48.4, 34.8, 28.9, 16.7, 15.9. LCMS (ESI): m/z= 354 (M+H)^+^

### Synthesis of ethyl (4-((2-oxopyrrolidin-1-yl)amino)-9H-pyrimido[4,5-b]indol-8-yl)carbamate formic acid salt (AVI-3367)

To a solution of 1-((8-amino-9H-pyrimido[4,5-b]indol-4-yl)amino)pyrrolidin-2-one (15 mg, 0.053 mmol) and triethylamine (0.015 mL, 0.11 mmol) in THF (1 mL), was added ethyl chloroformate (0.005 mL, 0.056 mmol). After stirring at 65 °C for 18 h, the reaction mixture was purified by reverse phase chromatography (water/acetonitrile/0.1% formic acid) to obtain 2.7 mg (13%) **AVI-3367** formate as tan solid. ^1^H NMR (METHANOL-d_4_, 400 MHz) δ 8.42 (s, 1H), 7.94 (d, 1H, *J*=7.8 Hz), 7.59 (br s, 1H), 7.28 (t, 1H, *J*=7.9 Hz), 4.1-4.26-4.30 (m, 2H), 3.84 (t, 2H, *J*=7.1 Hz), 2.60 (t, 2H, *J*=8.0 Hz), 2.30-2.33 (m, 2H), 1.36-1.39 (m, 3H). LCMS (ESI): m/z= 355 (M+H)^+^

### Synthesis of methyl (4-((2-oxopyrrolidin-1-yl)amino)-9H-pyrimido[4,5-b]indol-8-yl)sulfonamide formic acid salt (AVI-3766)

To a solution of 1-((8-amino-9H-pyrimido[4,5-b]indol-4-yl)amino)pyrrolidin-2-one (10 mg, 0.035 mmol), DMAP (2.2 mg, 0.18 mmol) and triethylamine (0.010 mL, 0.071 mmol) in DMA (0.5 mL), was added methylsulfamoyl chloride (15 mg, 0.11 mmol). After stirring at 65 °C for 90 h, the reaction mixture was purified by reverse phase chromatography (water/acetonitrile/0.1% formic acid) to obtain 2.6 mg (17%) **AVI-3766** formate as brown solid. ^1^H NMR (METHANOL-d_4_, 400 MHz) δ 8.44 (s, 1H), 8.01 (d, 1H, *J*=7.8 Hz), 7.51 (d, 1H, *J*=7.8 Hz), 7.34 (t, 1H, *J*=7.9 Hz), 3.84 (t, 2H, *J*=7.2 Hz), 2.68 (s, 3H), 2.60 (t, 2H, *J*=8.2 Hz), 2.28-2.32 (m, 2H). LCMS (ESI): m/z= 376 (M+H)^+^

### Synthesis of 1-cyclopropyl-3-(4-((2-oxopyrrolidin-1-yl)amino)-9H-pyrimido[4,5-b]indol-8-yl)urea formic acid salt (AVI-4051)

To a solution of 1-((8-amino-9H-pyrimido[4,5-b]indol-4-yl)amino)pyrrolidin-2-one (20 mg, 0.071 mmol) and triethylamine (0.040 mL, 0.28 mmol) in THF (1 mL), was added cyclopropyl isocyanate (24 mg, 0.28 mmol). After stirring at 65 °C for 48 h, the reaction mixture was purified by reverse phase chromatography (water/acetonitrile/0.1% formic acid) to obtain 12 mg (41%) of **AVI-4051** formate as a white solid. ^1^H NMR (DMSO-d_6_, 400 MHz) δ 11.81 (br s, 1H), 9.31 (s, 1H), 8.50 (br s, 1H), 8.41 (s, 1H), 7.99 (d, 1H, *J*=7.8 Hz), 7.63 (d, 1H, *J*=7.8 Hz), 7.21 (t, 1H, *J*=7.9 Hz), 6.74 (br s, 1H), 3.70 (br t, 2H, *J*=7.1 Hz), 3.12-3.17 (m, 2H), 2.54-2.65 (m, 1H), 2.39-2.43 (m, 2H), 0.98 (t, 2H, *J*=7.1 Hz), 0.68-0.70 (m, 2H). LCMS (ESI): m/z= 366 (M+H)^+^.

### Synthesis of 1-cyclopentyl-3-(4-((2-oxopyrrolidin-1-yl)amino)-9H-pyrimido[4,5-b]indol-8-yl)urea (AVI-4057)

To a solution of 1-((8-amino-9H-pyrimido[4,5-b]indol-4-yl)amino)pyrrolidin-2-one (20 mg, 0.071 mmol) and triethylamine (0.040 mL, 0.28 mmol) in THF (1 mL), was added cyclopentyl isocyanate (0.016 mL, 0.14 mmol). After stirring at 65 °C for 18 h, the reaction mixture was purified by reverse phase chromatography (water/acetonitrile/0.1% formic acid) to obtain 6 mg (20%) of **AVI-4057** formate as a white solid. ^1^H NMR (DMSO-d_6_, 400 MHz) δ 11.74 (br s, 1H), 9.31 (s, 1H), 8.41 (s, 1H), 7.97 (d, 1H, *J*=7.8 Hz), 7.58 (d, 1H, *J*=7.8 Hz), 7.35 (s, 1H), 7.20 (t, 1H, *J*=7.9 Hz), 6.47 (br d, 1H, *J*=7.1 Hz), 4.03-4.05 (m, 1H), 3.70 (t, 2H, *J*=7.1 Hz), 2.41 (t, 2H, *J*=8.0 Hz), 2.13 (br dd, 2H, *J*=7.8, 15.6 Hz), 1.89 (br dd, 2H, *J*=5.5, 12.1 Hz), 1.67-1.69 (m, 2H), 1.46-1.58 (m, 4H). LCMS (ESI): m/z= 394 (M+H)^+^

### Synthesis of (R)-3-methyl-2-((6-phenyl-7H-pyrrolo[2,3-d]pyrimidin-4-yl)amino)butan-1-ol (AVI-4211)

A mixture of (R)-2-((6-bromo-7*H*-pyrrolo[2,3-*d*]pyrimidin-4-yl)amino)-3-methylbutan-1-ol (15.0 mg, 50.1 μmol), phenylboronic acid (12.2 mg, 100.0 μmol), Pd(dppf)Cl_2_ (3.7 mg, 5.01 μmol) and Cs_2_CO_3_ (40.8 mg, 125 μmol) in 0.22 mL of dioxane/H_2_O (10:1) was stirred at 110 °C for 17 hours. The residue was purified by prep-HPLC (0%-70% water/ACN with 0.1% formic acid) to give **AVI-4211** formate as a white solid (9.7 mg, yield: 57%). 1H NMR (400 MHz, MeOD) δ 8.41 (brs, 1H), 8.11 (s, 1H), 7.79 (brd, 1H, *J* = 8.0 Hz), 7.45 (brdd, 2H, *J* = 8.0, 7.5 Hz), 7.33 (brt, 1H, *J* = 7.5 Hz), 7.03 (s, 1H), 4.16-4.12 (m, 1H), 3.85-3.75 (m, 2H), 2.15-2.08 (m, 1H), 1.08-1.04 (m, 6H). LCMS (ESI): m/z=297 (M+H)^+^

### Synthesis of (R)-2-((6-(2-fluorophenyl)-7H-pyrrolo[2,3-d]pyrimidin-4-yl)amino)-3-methylbutan-1-ol (AVI-4678)

A mixture of (R)-2-((6-bromo-7*H*-pyrrolo[2,3-*d*]pyrimidin-4-yl)amino)-3-methylbutan-1-ol (15.0 mg, 50.1 μmol), (2-fluorophenyl)boronic acid (14.0 mg, 100.0 μmol), Pd(dppf)Cl_2_ (3.7 mg, 5.01 μmol) and Cs_2_CO_3_ (40.8 mg, 125 μmol) in 0.22 mL of of dioxane/H_2_O (10:1) was stirred at 110 °C for 2 hours. The residue was purified by prep-HPLC (0%-70% water/ACN with 0.1% formic acid) to give **AVI-4678** formate as a white solid (10.9 mg, yield: 60%). 1H NMR (400 MHz, MeOD) δ 8.39 (brs, 1H), 8.13 (s, 1H), 7.83-7.78 (m, 1H), 7.37-7.20 (m, 3H), 7.16 (s, 1H), 4.17-4.12 (m, 1H), 3.86-3.75 (m, 2H), 2.17-2.07 (m, 1H), 1.08-1.03 (m, 6H). LCMS (ESI): m/z=315 (M+H)+.

### Synthesis f (R)-2-((6-(2-chlorophenyl)-7H-pyrrolo[2,3-d]pyrimidin-4-yl)amino)-3-methylbutan-1-ol (AVI-4335)

A mixture of (R)-2-((6-bromo-7*H*-pyrrolo[2,3-*d*]pyrimidin-4-yl)amino)-3-methylbutan-1-ol (15.0 mg, 50.1 μmol), (2-chlorophenyl)boronic acid (15.7 mg, 100.0 μmol), Pd(dppf)Cl_2_ (3.7 mg, 5.01 μmol) and Cs_2_CO_3_ (40.8 mg, 125 μmol) in 0.22 mL of of dioxane/H_2_O (10:1) was stirred at 110 °C for 1 hour. The residue was purified by prep-HPLC (0%-50% water/ACN with 0.1% formic acid) to give **AVI-4335** formate as a white solid (8.3 mg, yield: 44%). 1H NMR (400 MHz, MeOD) δ 8.16 (brs, 1H), 7.66 (brd, 1H, *J* = 7.7 Hz), 7.55 (brd, 1H, *J* = 7.9 Hz), 7.45-7.35 (m, 2H), 7.13 (s, 1H), 4.15-4.11 (m, 1H), 3.86-3.75 (m, 2H), 2.15-2.07 (m, 1H), 1.08-1.04 (m, 6H). LCMS (ESI): m/z=331 (M+H)+

### Synthesis of (R)-3-methyl-2-((6-(o-tolyl)-7H-pyrrolo[2,3-d]pyrimidin-4-yl)amino)butan-1-ol (AVI-4370)

A mixture of (R)-2-((6-iodo-7H-pyrrolo[2,3-d]pyrimidin-4-yl)amino)-3-methylbutan-1-ol (15.0 mg, 43.3 μmol), *o*-tolylboronic acid (11.8 mg, 86.7 μmol), Pd(dppf)Cl_2_ (3.2 mg, 4.3 μmol) and Cs_2_CO_3_ (35.3 mg, 108 μmol) in 0.22 mL of of dioxane/H_2_O (10:1) was stirred at 110 °C for 1 hour. The residue was purified by prep-HPLC (0%-50% water/ACN with 0.1% formic acid) to give **AVI-4370** formate as a white solid (5.4 mg, yield: 35%). 1H NMR (400 MHz, MeOD) δ 8.35 (brs, 1H), 8.13 (s, 1H), 7.51-7.49 (m, 1H), 7.34-7.27 (m, 3H), 7.16-7.14 (brs, 1H), 6.68 (brs, 1H), 4.13-4.11 (m, 1H), 3.86-3.75 (m, 2H), 2.50 (s, 3H), 2.14-2.09 (m, 1H), 1.08-1.04 (m, 6H). LCMS (ESI): m/z=311 (M+H)^+^ Synthesis of (R)-2-(4-((1-hydroxy-3-methylbutan-2-yl)amino)-7H-pyrrolo[2,3-d]pyrimidin-6-yl)benzonitrile (AVI-4213). A mixture of (R)-2-((6-bromo-7*H*-pyrrolo[2,3-*d*]pyrimidin-4-yl)amino)-3-methylbutan-1-ol (15.0 mg, 50.1 μmol), (2-cyanophenyl)boronic acid (14.7 mg, 100.0 μmol), Pd(dppf)Cl_2_ (3.7 mg, 5.01 μmol) and Cs_2_CO_3_ (40.8 mg, 125 μmol) in 0.22 mL of of dioxane/H_2_O (10:1) was stirred at 110 °C for 4 hours. The residue was purified by prep-HPLC (0%-50% water/ACN) to give **AVI-4213** as a white solid (1.2 mg, yield: 7%). 1H NMR (400 MHz, MeOD) δ 8.52 (brs, 1H), 8.16 (s, 1H), 7.88-7.75 (m, 3H), 7.52 (ddd, 1H, *J* = 7.6, 7.6, 1.4 Hz), 7.33 (s, 1H), 4.21-4.14 (m, 1H), 3.85-3.75 (m, 2H), 2.14-2.09 (m, 1H), 1.08-1.04 (m, 6H). LCMS (ESI): m/z=322 (M+H)^+^

### Synthesis of)-3-methyl-2-((6-(pyridin-3-yl)-7H-pyrrolo[2,3-d]pyrimidin-4-yl)amino)butan-1-ol (AVI-4271)

A mixture of (R)-2-((6-bromo-7*H*-pyrrolo[2,3-*d*]pyrimidin-4-yl)amino)-3-methylbutan-1-ol (15.0 mg, 50.1 μmol), pyridin-3-ylboronic acid (18.5 mg, 150.0 μmol), Pd(dppf)Cl_2_ (7.4 mg, 10.0 μmol) and Cs_2_CO_3_ (40.8 mg, 125 μmol) in 0.22 mL of of dioxane/H_2_O (10:1)was stirred at 110 °C for 24 hours. The residue was purified by prep-HPLC (0%-30% water/ACN with 0.1% formic acid) to give **AVI-4271** formate as a white solid (3.7 mg, yield: 22%). 1H NMR (400 MHz, MeOD) δ 8.98 (s, 1H), 8.48 (d, 1H, *J* = 4.8 Hz), 8.20 (d, 1H, *J* = 8.1 Hz), 8.13 (s, 1H), 7.52 (dd, 1H, *J* = 8.1, 4.7 Hz), 7.16 (brs, 1H), 4.18 (ddd, 1H, *J* = 7.6, 5.8, 4.1 Hz), 3.85-3.76 (m, 2H), 2.16-2.07 (m, 1H), 1.08-1.04 (m, 6H). LCMS (ESI): m/z=298 (M+H)^+^

### Synthesis of (R)-3-methyl-2-((6-(pyrimidin-5-yl)-7H-pyrrolo[2,3-d]pyrimidin-4-yl)amino)butan-1-ol (AVI-4272)

A mixture of (R)-2-((6-bromo-7*H*-pyrrolo[2,3-*d*]pyrimidin-4-yl)amino)-3-methylbutan-1-ol (15.0 mg, 50.1 μmol), pyrimidin-5-ylboronic acid (24.8 mg, 200.0 μmol), Pd(dppf)Cl_2_ (18.4 mg, 25.1 μmol) and Cs_2_CO_3_ (40.8 mg, 125 μmol) in 0.22 mL of of dioxane/H_2_O (10:1)was stirred at 110 °C for 4 days. The residue was purified by prep-HPLC (0%-30% water/ACN with 0.1% formic acid) to give **AVI-4272** formate as a white solid (1.0 mg, yield: 6%). 1H NMR (400 MHz, MeOD) δ 9.19 (s, 2H), 9.08 (s, 1H), 8.48 (brs, 2H), 8.16 (s, 1H), 7.26 (s, 1H), 4.22-4.18 (m, 1H), 3.85-3.75 (m, 2H), 2.14-2.07 (m, 1H), 1.08-1.04 (m, 6H). LCMS (ESI): m/z=299 (M+H)^+^

### Synthesis of (R)-3-methyl-2-((6-(thiophen-2-yl)-7H-pyrrolo[2,3-d]pyrimidin-4-yl)amino)butan-1-ol (AVI-4094)

A mixture of (R)-2-((6-bromo-7*H*-pyrrolo[2,3-*d*]pyrimidin-4-yl)amino)-3-methylbutan-1-ol (15 mg, 50.1 μmol), thiophen-2-ylboronic acid (12.8 mg, 100.0 μmol), Pd(dppf)Cl_2_ (3.7 mg, 5.01 μmol) and Cs_2_CO_3_ (40.8 mg, 125 μmol) in 0.22 mL of dioxane/H_2_O (10:1) was stirred at 110 °C for 10 min. The residue was purified by prep-HPLC (0%-50% water/ACN with 0.1% formic acid) to give **AVI-4094** formate as a white solid (13.2 mg, yield: 87%). 1H NMR (400 MHz, MeOD) δ 8.30 (brs, 1H), 8.11 (s, 1H), 7.43 (dd, 1H, *J* = 3.6, 1.4 Hz), 7.39 (dd, 1H, *J* = 5.1, 1.4 Hz), 7.12-7.09 (m, 1H), 6.89 (s, 1H), 4.14-4.04 (m, 1H), 3.85-3.74 (m, 2H), 2.14-2.06 (m, 1H), 1.07-1.03 (m, 6H). LCMS (ESI): m/z=303 (M+H)^+^

### Synthesis of (R)-3-methyl-2-((6-(thiophen-3-yl)-7H-pyrrolo[2,3-d]pyrimidin-4-yl)amino)butan-1-ol (AVI-4334)

A mixture of (R)-2-((6-bromo-7*H*-pyrrolo[2,3-*d*]pyrimidin-4-yl)amino)-3-methylbutan-1-ol (15 mg, 50.1 μmol), thiophen-3-ylboronic acid (12.8 mg, 100.0 μmol), Pd(dppf)Cl_2_ (3.7 mg, 5.01 μmol) and Cs_2_CO_3_ (40.8 mg, 125 μmol) in 0.22 mL of dioxane/H_2_O (10:1)was stirred at 110 °C for 2 hours. The residue was purified by prep-HPLC (0%-60% water/ACN with 0.1% formic acid) to give **AVI-4334** formate as a white solid (10.5 mg, yield: 60%). 1H NMR (400 MHz, MeOD) δ 8.10 (s, 1H), 7.72-7.70 (m, 1H), 7.52-7.50 (m, 2H), 6.87 (brs, 1H), 4.13 (ddd, 1H, *J* = 7.5, 5.7, 4.1 Hz), 3.84-3.75 (m, 2H), 2.14-2.09 (m, 1H), 1.08-1.03 (m, 6H). LCMS (ESI): m/z=303 (M+H)^+^

### Synthesis of (R)-4-(4-((1-hydroxy-3-methylbutan-2-yl)amino)-7H-pyrrolo[2,3-d]pyrimidin-6-yl)thiophene-3-carbonitrile (AVI-4684)

A mixture of (R)-2-((6-iodo-7H-pyrrolo[2,3-d]pyrimidin-4-yl)amino)-3-methylbutan-1-ol (15.0 mg, 43.3 μmol), 4-(4,4,5,5-tetramethyl-1,3,2-dioxaborolan-2-yl)thiophene-3-carbonitrile (16.3 mg, 69.3 μmol), Pd(dppf)Cl_2_ (3.2 mg, 4.33 μmol) and Cs_2_CO_3_ (35.3 mg, 108 μmol) in 0.22 mL of dioxane/H_2_O (10:1) was stirred at 110 °C for 1 hour. The residue was purified by prep-HPLC (0%-80% water/ACN) to give **AVI-4684** as a white solid (4.5 mg, yield: 32%). 1H NMR (400 MHz, *d*-DMSO) δ 12.0 (brs, 1H), 8.71 (d, 1H, *J* = 3.1 Hz), 8.11 (s, 1H), 8.02 (d, 1H, *J* = 3.1 Hz), 7.43 (brd, 1H, *J* = 8.6 Hz), 7.28 (s, 1H), 4.65-4.62 (m, 1H), 4.25-4.18 (m, 1H), 3.60-3.57 (m, 2H), 2.07-1.99 (m, 1H), 0.96-0.92 (m, 6H). LCMS (ESI): m/z=328 (M+H)^+^

### Synthesis of (R)-3-(4-((1-hydroxy-3-methylbutan-2-yl)amino)-7H-pyrrolo[2,3-d]pyrimidin-6-yl)thiophene-2-carbonitrile (AVI-4683)

A mixture of (R)-2-((6-iodo-7H-pyrrolo[2,3-d]pyrimidin-4-yl)amino)-3-methylbutan-1-ol (15.0 mg, 43.3 μmol), 3-(4,4,5,5-tetramethyl-1,3,2-dioxaborolan-2-yl)thiophene-2-carbonitrile (16.3 mg, 69.3 μmol), Pd(dppf)Cl_2_ (3.2 mg, 4.33 μmol) and Cs_2_CO_3_ (35.3 mg, 108 μmol) in 0.22 mL of dioxane/H_2_O (10:1) was stirred at 110 °C for 1 hour. The residue was purified by prep-HPLC (0%-60% water/ACN) to give **AVI-4683** as a white solid (5.3 mg, yield: 37%). 1H NMR (400 MHz, *d*-DMSO) δ 12.2 (brs, 1H), 8.15-8.12 (m, 2H), 7.74 (d, 1H, *J* = 5.2 Hz), 7.57 (brd, 1H, *J* = 8.7 Hz), 7.50 (s, 1H), 4.65-4.64 (m, 1H), 4.25-4.17 (m, 1H), 3.61-3.58 (m, 2H), 2.06-2.01 (m, 1H), 0.96-0.93 (m, 6H). LCMS (ESI): m/z=328 (M+H)^+^

### Synthesis of (R)-2-((6-(furan-2-yl)-7H-pyrrolo[2,3-d]pyrimidin-4-yl)amino)-3-methylbutan-1-ol (AVI-4267)

A mixture of (R)-2-((6-bromo-7*H*-pyrrolo[2,3-*d*]pyrimidin-4-yl)amino)-3-methylbutan-1-ol (15.0 mg, 50.1 μmol), 2-(furan-2-yl)-4,4,5,5-tetramethyl-1,3,2-dioxaborolane (19.5 mg, 100.0 μmol), Pd(dppf)Cl_2_ (3.7 mg, 5.0 μmol) and Cs_2_CO_3_ (40.8 mg, 125 μmol) in 0.22 mL of dioxane/H_2_O (10:1) was stirred at 110 °C for 5 minutes. The residue was purified by prep-HPLC (0%-50% water/ACN with 0.1% formic acid) to give **AVI-4267** formate as a white solid (6.7 mg, yield: 40%). 1H NMR (400 MHz, MeOD) δ 8.11 (brs, 1H), 7.59-7.57 (m, 1H), 6.90 (m, 1H), 6.75 (m, 1H), 6.57-6.54 (m, 1H), 4.15-4.11 (m, 1H), 3.84-3.74 (m, 1H), 2.15-2.06 (m, 1H), 1.07-1.02 (m, 6H). LCMS (ESI): m/z=287 (M+H)^+^

### Synthesis of (R)-2-((6-(1H-pyrrol-2-yl)-7H-pyrrolo[2,3-d]pyrimidin-4-yl)amino)-3-methylbutan-1-ol (AVI-4097)

A mixture of (R)-2-((6-bromo-7*H*-pyrrolo[2,3-*d*]pyrimidin-4-yl)amino)-3-methylbutan-1-ol (15.0 mg, 50.1 μmol), (1-(tert-butoxycarbonyl)-1*H*-pyrrol-2-yl)boronic acid (21.2 mg, 100.0 μmol), Pd(dppf)Cl_2_ (3.67 mg, 5.0 μmol) and Cs_2_CO_3_ (40.8 mg, 125 μmol) in 0.22 mL of dioxane/H_2_O (10:1) was stirred at 110 °C for 24 hours. The residue was purified by NP-column (CH_2_CL_2_/MeOH 95:5 with NH_3_) to give tert-butyl (*R*)-2-(4-((1-hydroxy-3-methylbutan-2-yl)amino)-7*H*-pyrrolo[2,3-*d*]pyrimidin-6-yl)-1*H*-pyrrole-1-carboxylate as a brown solid (11.9 mg, yield: 62%). LCMS (ESI): m/z=386 (M+H)^+^ Add solution of tert-butyl (*R*)-2-(4-((1-hydroxy-3-methylbutan-2-yl)amino)-7*H*-pyrrolo[2,3-*d*]pyrimidin-6-yl)-1*H*-pyrrole-1-carboxylate (11.9 mg, 30.9 μmol) in 0.20 mL of methanol dropwise at 0 °C to a solution of acetyl chloride (44.0 μL, 618 mmol) and the reaction mixture stirred 2 days at room temperature. The reaction mixture was concentrated and the residue washed with 7M NH_3_ in MeOH and evaporated three times repeatedly. The residue was purified by prep-HPLC (0%-50% water/ACN with 0.1% formic acid) to give **AVI-4097** formate as a white solid (5.8 mg, yield: 66%). 1H NMR (400 MHz, MeOD) δ 8.31 (brs, 1H), 8.07 (brs, 1H), 6.87-6.86 (m, 1H), 6.68-6.66 (m, 1H), 6.57-6.55 (m, 1H), 6.21-6.19 (m, 1H), 4.10-4.04 (m, 1H), 3.85-3.75 (m, 2H), 2.17-2.04 (m, 1H), 1.08-1.04 (m, 6H). LCMS (ESI): m/z=286 (M+H)+

### Synthesis of (R)-2-((6-(1H-indol-6-yl)-7H-pyrrolo[2,3-d]pyrimidin-4-yl)amino)-3-methylbutan-1-ol (AVI-4212)

A mixture of (R)-2-((6-bromo-7*H*-pyrrolo[2,3-*d*]pyrimidin-4-yl)amino)-3-methylbutan-1-ol (15.0 mg, 50.1 μmol), (1*H*-indol-6-yl)boronic acid (16.1 mg, 200.0 μmol), Pd(dppf)Cl_2_ (3.7 mg, 5.01 μmol) and Cs_2_CO_3_ (40.8 mg, 125 μmol) in 0.22 mL of dioxane/H_2_O (10:1)was stirred at 110 °C for 17 hours. The residue was purified by prep-HPLC (0%-50% water/ACN with 0.1% formic acid) to give **AVI-4212** formate as a white solid (8.1 mg, yield: 42%). 1H NMR (400 MHz, MeOD) δ 8.43 (brs, 1H), 8.10 (s, 1H), 7.81 (s, 1H), 7.62 (d, 1H, *J* = 8.3 Hz), 7.48 (d, 1H, *J* = 8.3 Hz), 7.30 (d, 1H, *J* = 3.1 Hz), 6.97 (s, 1H), 6.48 (d, 1H, *J* = 3.1 Hz), 4.14-4.10 (m, 1H), 3.87-3.76 (m, 2H), 2.17-2.08 (m, 1H), 1.09-1.05 (m, 6H). LCMS (ESI): m/z=336 (M+H)^+^

### Synthesis of (R)-2-((6-(1H-pyrazol-5-yl)-7H-pyrrolo[2,3-d]pyrimidin-4-yl)amino)-3-methylbutan-1-ol (AVI-4099)

A mixture of (R)-2-((6-bromo-7*H*-pyrrolo[2,3-*d*]pyrimidin-4-yl)amino)-3-methylbutan-1-ol (10.0 mg, 33.4 μmol), 5-(4,4,5,5-tetramethyl-1,3,2-dioxaborolan-2-yl)-1*H*-pyrazole (13.0 mg, 66.9 μmol), Pd(dppf)Cl_2_ (4.9 mg, 6.7 μmol) and CsOH (12.5 mg, 83.6 μmol) in 0.25 mL of ^*n*^BuOH/H_2_O (4:1) was stirred at 130 °C for 20 minutes with microwave. The residue was purified by prep-HPLC (0%-30% water/ACN with 0.1% formic acid) to give **AVI-4099** formate as a white solid (3.7 mg, yield: 39%). 1H NMR (400 MHz, MeOD) δ 8.42 (brs, 1H), 8.12 (brs, 1H), 7.73 (d, 1H, *J* = 2.3 Hz), 6.97 (s, 1H), 6.72 (d, 1H, *J* = 2.3 Hz), 4.16-4.11 (m, 1H), 3.84-3.75 (m, 2H), 2.16-2.06 (m, 1H), 1.09-1.02 (m, 6H). LCMS (ESI): m/z=287 (M+H)^+^

### Synthesis of (R)-2-((6-(1H-pyrazol-4-yl)-7H-pyrrolo[2,3-d]pyrimidin-4-yl)amino)-3-methylbutan-1-ol (AVI-4268)

A mixture of (R)-2-((6-bromo-7*H*-pyrrolo[2,3-*d*]pyrimidin-4-yl)amino)-3-methylbutan-1-ol (15.0 mg, 50.1 μmol), (1*H*-pyrazol-4-yl)boronic acid (22.4 mg, 200.0 μmol), Pd(dppf)Cl_2_ (7.3 mg, 10.0 μmol) and Cs_2_CO_3_ (40.8 mg, 125 μmol) in 0.22 mL of dioxane/H_2_O (10:1)was stirred at 110 °C for 4 days. The residue was purified by prep-HPLC (0%-40% water/ACN with 0.1% formic acid) to give **AVI-4268** formate as a white solid (2.7 mg, yield: 16%). 1H NMR (400 MHz, MeOD) δ 8.48 (brs, 1H), 8.15-8.14 (m, 2H), 7.77 (m, 1H), 6.71 (s, 1H), 6.57 (m, 1H), 4.18-4.14 (m, 1H), 3.84-3.74 (m, 2H), 2.16-2.01 (m, 1H), 1.07-1.03 (m, 6H). LCMS (ESI): m/z=287 (M+H)^+^

### Synthesis of (R)-3-methyl-2-((6-(1-methyl-1H-pyrazol-4-yl)-7H-pyrrolo[2,3-d]pyrimidin-4-yl)amino)butan-1-ol (AVI-4214)

A mixture of (R)-2-((6-bromo-7*H*-pyrrolo[2,3-*d*]pyrimidin-4-yl)amino)-3-methylbutan-1-ol (15.0 mg, 50.1 μmol), (1-methyl-1*H*-pyrazol-4-yl)boronic acid (12.6 mg, 100.0 μmol), Pd(dppf)Cl_2_ (3.7 mg, 5.01 μmol) and Cs_2_CO_3_ (40.8 mg, 125 μmol) in 0.22 mL of dioxane/H_2_O (10:1) was stirred at 110 °C for 4 hours. The residue was purified by prep-HPLC (0%-50% water/ACN with 0.1% formic acid) to give **AVI-4214** formate as a white solid (11.1 mg, yield: 64%). 1H NMR (400 MHz, MeOD) δ 8.29 (brs, 1H), 8.08 (s, 1H), 7.97 (s, 1H), 7.86 (s, 1H), 6.75 (brs, 1H), 4.10-4.05 (m, 1H), 3.96 (s, 3H), 3.86-3.74 (m, 2H), 2.14-2.06 (m, 1H), 1.09-1.03 (m, 6H). LCMS (ESI): m/z=301 (M+H)^+^

### Synthesis of (R)-2-((6-(1,5-dimethyl-1H-pyrazol-4-yl)-7H-pyrrolo[2,3-d]pyrimidin-4-yl)amino)-3-methylbutan-1-ol (AVI-4266)

A mixture of (R)-2-((6-bromo-7*H*-pyrrolo[2,3-*d*]pyrimidin-4-yl)amino)-3-methylbutan-1-ol (15.0 mg, 50.1 μmol), 1,5-dimethyl-4-(4,4,5,5-tetramethyl-1,3,2-dioxaborolan-2-yl)-1*H*-pyrazole (44.6 mg, 200.0 μmol), Pd(dppf)Cl_2_ (7.3 mg, 10.0 μmol) and Cs_2_CO_3_ (40.8 mg, 125 μmol) in 0.22 mL of dioxane/H_2_O (10:1) was stirred at 110 °C for 2 days. The residue was purified by prep-HPLC (0%-50% water/ACN with 0.1% formic acid) to give **AVI-4266** formate as a white solid (4.1 mg, yield: 23%). 1H NMR (400 MHz, MeOD) δ 8.52 (brs, 1H), 8.08 (s, 1H), 7.74 (s, 1H), 6.67 (s, 1H), 4.15-4.10 (m, 1H), 3.88-3.87 (m, 3H), 3.84-3.74 (m, 2H), 2.53-2.52 (m, 3H), 2.14-2.07 (m, 1H), 1.07-1.03 (m, 6H). LCMS (ESI): m/z=315 (M+H)^+^

### Synthesis of (R)-4-(4-((1-hydroxy-3-methylbutan-2-yl)amino)-7H-pyrrolo[2,3-d]pyrimidin-6-yl)-1-methyl-1H-pyrazole-5-carbonitrile (AVI-4054)

To a solution of 4-(4-chloro-7H-pyrrolo[2,3-d]pyrimidin-6-yl)-1-methyl-1H-pyrazole-5-carbonitrile (70 mg, 0.27 mmol) in dry DMSO (2 mL) was added (R)-2-amino-3-methylbutan-1-ol (42.23 mg, 0.41 mmol) and triethylamine (72.4 mg, 0.81 mmol). After stirring at 100oC for 16 h, the mixture was diluted with ethyl acetate (40.0 mL) and washed with water (5.0 mL) and brine (5.0 mL). The organic layer was dried over Na_2_SO_4_ and concentrated under reduced pressure. The residue was purified by prep-HPLC (10%-100% 0.1% NH_4_HCO_3_ in water/ACN) to give **AVI-4054** as a white solid (25.4 mg, yield: 28.8%). 1H NMR (500 MHz, MeOD) δ 8.09 (s, 1H), 7.98 (s, 1H), 7.13 (s, 1H), 4.14 (s, 1H), 4.09 (s, 3H), 3.77 (dd, *J* = 13.4, 5.0 Hz, 2H), 2.08 (dt, *J* = 13.8, 6.9 Hz, 1H), 1.02 (dd, *J* = 13.2, 6.8 Hz, 6H). LCMS (ESI): m/z 326.1 (M+H)+

### (R)-4-(4-((1-hydroxy-3-methylbutan-2-yl)amino)-7H-pyrrolo[2,3-d]pyrimidin-6-yl)-1-methyl-1H-pyrazole-3-carbonitrile (AVI-5707)

A mixture of (R)-2-((6-iodo-7H-pyrrolo[2,3-d]pyrimidin-4-yl)amino)-3-methylbutan-1-ol (15 mg, 43.3 umol), 1-methyl-4-(4,4,5,5-tetramethyl-1,3,2-dioxaborolan-2-yl)-1H-pyrazole-3-carbonitrile (16.2 mg, 69.3 umol), Pd(dppf)Cl_2_ (3.2 mg, 4.33 umol) and CsCO_3_ (35.3 mg, 108 mmol) in 0.22 mL of dioxane/H_2_O (10:1) was stirred at 110 °C for 1 hour. Subsequently, the mixture was extracted thrice with EtOAc, the combined organic phase was washed with brine, dried over Na2SO4 and concentrated in vacuo. The residue was purified by silica gel chromatography (20-40%EtOAc in hexane and prep-HPLC, 2.1 mg (13%) of **AVI-5707** formate as a white solid. 1H NMR (DMSO-d6, 400 MHz) δ 11.96-12.03 (m, 1H), 8.44 (br s, 1H), 8.28-8.30 (m, 1H), 8.08 (s, 1H), 7.40 (br d, 1H, J=8.0 Hz), 7.14 (s, 1H), 4.19 (br s, 1H), 4.00 (s, 3H), 3.57-3.61 (m, 2H), 2.02 (qd, 1H, J=6.7, 13.6 Hz), 1.24 (s, 1H), 0.94 (t, 6H, **J**=6.5 Hz). LCMS (ESI): m/z=326 (M+H)+

### Synthesis of (R)-2-((6-(isothiazol-5-yl)-7H-pyrrolo[2,3-d]pyrimidin-4-yl)amino)-3-methylbutan-1-ol (AVI-4100)

A mixture of (R)-2-((6-bromo-7*H*-pyrrolo[2,3-*d*]pyrimidin-4-yl)amino)-3-methylbutan-1-ol (10.0 mg, 33.4 μmol), 5-(4,4,5,5-tetramethyl-1,3,2-dioxaborolan-2-yl)isothiazole (14.1 mg, 66.9 μmol), Pd(PPh_3_)_4_ (7.7 mg, 6.7 μmol) and Na_2_CO_3_ (7.09 mg, 66.9 μmol) in 0.22 mL of dioxane/H_2_O (10:1) was stirred at 110 °C for 24 hours. The residue was purified by prep-HPLC (0%-40% water/ACN with 0.1% formic acid) to give **AVI-4100** formate as a white solid (1.2 mg, yield: 10%). 1H NMR (400 MHz, MeOD) δ 8.55 (brs, 1H), 8.49 (d, 1H, *J* = 1.9 Hz), 8.15 (s, 1H), 7.63 (d, 1H, *J* = 1.9 Hz), 7.14 (s, 1H), 4.19-4.18 (m, 1H), 3.83-3.74 (m, 2H), 2.13-2.08 (m, 1H), 1.07-1.03 (m, 6H). LCMS (ESI): m/z=304 (M+H)^+^

### Synthesis of 4-(4-((2,2-dimethyl-5-oxopyrrolidin-1-yl)amino)-7H-pyrrolo[2,3-d]pyrimidin-6-yl)-1-methyl-1H-pyrazole-5-carbonitrile (AVI-4052)

A solution of 4-(4-chloro-7H-pyrrolo[2,3-d]pyrimidin-6-yl)-1-methyl-1H-pyrazole-5-carbonitrile (70 mg, 0.27 mmol) and 1-amino-5,5-dimethylpyrrolidin-2-one (63.76 mg, 0.41 mmol) in iPrOH (2 mL) and HCl (1 drop) was stirred at 100 °C for 16 hours. The reaction mixture was suspended in water (30 mL x 3) and extracted with ethyl acetate (3 x 60 mL). The organic layer was dried over Na_2_SO_4_ and concentrated under reduced pressure. The residue was purified by prep-HPLC (10%-100% 0.1% NH_4_HCO_3_ in water/ACN) to **AVI-4052** as a white solid (25.1 mg, yield: 26.4%). 1H NMR (500 MHz, MeOD) δ 8.32 (s, 1H), 8.04 (s, 1H), 7.34 (s, 1H), 4.12 (s, 3H), 2.60 (d, J = 7.7 Hz, 2H), 2.22 (s, 2H), 1.37 (s, 6H). LCMS (ESI): m/z 351.1 (M+H)^+^

### Synthesis of 4-(4-((2,2-dimethyl-6-oxopiperidin-1-yl)amino)-7H-pyrrolo[2,3-d]pyrimidin-6-yl)-1-methyl-1H-pyrazole-5-carbonitrile (AVI-6320)

To a solution of 4-(4-chloro-7H-pyrrolo[2,3-d]pyrimidin-6-yl)-1-methyl-1H-pyrazole-5-carbonitrile (120 mg, 0.46 mmol) in IPA (5 mL) was added 1-amino-6,6-dimethylpiperidin-2-one (99 mg, 0.70 mmol) and conc. HCl (0.04 mL). After stirring at 100°C for 24 hours, aqueous, saturated sodium bicarbonate (5 mL) was added to the reaction mixture and extracted with _DCM_ (20 mL×3), organic layers were separated and concentrated in vacuo, and purified by preprarative HPLC (10%-40% 0.1% NH_4_HCO_3_ in water/ACN) to obtain **AVI-6320** as a white solid (24mg, Yield: 14.45%). ^1^H NMR (400 MHz, DMSO) δ 12.11 (s, 1H), 9.15 (s, 1H), 8.15 (d, J = 12.7 Hz, 2H), 6.92 (s, 1H), 4.04 (s, 3H), 2.68 – 2.52 (m, 1H), 2.39 – 2.25 (m, 1H), 2.05 (s, 1H), 1.84 (d, J = 5.5 Hz, 3H), 1.30 (d, J = 26.5 Hz, 6H). LCMS (ESI): m/z= 365.3 (M+H)^+^

### Synthesis of 4-(4-((4,4-dimethyl-2-oxooxazolidin-3-yl)amino)-7H-pyrrolo[2,3-d]pyrimidin-6-yl)-1-methyl-1H-pyrazole-5-carbonitrile (AVI-6318)

To a solution of 4-(4-chloro-7-((2-(trimethylsilyl)ethoxy)methyl)-7H-pyrrolo[2,3-d]pyrimidin-6-yl)-1-methyl-1H-pyrazole-5-carbonitrile (200 mg, 0.52 mmol) in IPA (5 mL) were added 3-amino-4,4-dimethyloxazolidin-2-one (98 mg, 0.75 mmol) and conc. HCl (0.04 mL). After stirring at 100°C for 24 hours, aqueous saturated sodium bicarbonate (5 mL) was added to the reaction mixture and extracted with DCM (20 mL×3), organic layers were separated and concentrated in vacuo and purified by silica gel chromatography (dichloromethane/methanol, 10:1) to obtain 4-(4-((4,4-dimethyl-2-oxooxazolidin-3-yl)amino)-7-((2-(trimethylsilyl)ethoxy)methyl)-7H-pyrrolo[2,3-d]pyrimidin-6-yl)-1-methyl-1H-pyrazole-5-carbonitrile (55 mg, Yield:21.9%) as a white solid. LCMS (ESI): m/z= 483.2 (M+H)^+^

To a solution of the intermediate above (55 mg, 0.11mmol) in DCM (2.0 mL) was added TFA (0.5 mL). After stirring at 20 °C for 4 h, the reaction mixture was concentrated to dryness, the resulting residue was suspended in aqueous saturated NaHCO3 and extracted with DCM (10 mL × 3). The organic layers were washed with brine, concentrated under reduced pressure and purified by preprarative HPLC (10%-50% 0.1% NH_4_HCO_3_ in water/ACN) to obtain **AVI-6318** as a white solid (20 mg, Yield: 29.8%). ^1^H NMR (500 MHz, DMSO) δ 12.44 (s, 1H), 9.76 (s, 1H), 8.25 (s, 1H), 8.17 (s, 1H), 7.08 (s, 1H), 4.41 – 4.15 (m, 2H), 4.07 (s, 3H), 1.31 (d, *J* = 11.9 Hz, 6H). LCMS (ESI): m/z= 353.1 (M+H)^+^

### Synthesis of 1-methyl-4-(4-((2-oxo-3,8-dioxa-1-azaspiro[4.5]decan-1-yl)amino)-7H-pyrrolo[2,3-d]pyrimidin-6-yl)-1H-pyrazole-5-carbonitrile (AVI-6319)

To a solution of 4-(4-chloro-7-((2-(trimethylsilyl)ethoxy)methyl)-7H-pyrrolo[2,3-d]pyrimidin-6-yl)-1-methyl-1H-pyrazole-5-carbonitrile (200 mg, 0.52 mmol) in IPA(5 mL) were added 1-amino-3,8-dioxa-1-azaspiro[4.5]decan-2-one (129 mg, 0.75 mmol) and conc. HCl (0.04 mL). After stirring at 100°C for 24 hours, aqueous saturated sodium bicarbonate (5 mL) was added to the reaction mixture and extracted with DCM (20 mL×3), organic layers were separated and concentrated in vacuo and purified by silica gel chromatography (dichloromethane/methanol, 10:1) to obtain 1-methyl-4-(4-((2-oxo-3,8-dioxa-1-azaspiro[4.5]decan-1-yl)amino)-7-((2-(trimethylsilyl)ethoxy)methyl)-7H-pyrrolo[2,3-d]pyrimidin-6-yl)-1H-pyrazole-5-carbonitrile (60 mg, Yield:22%) as a white solid. LCMS (ESI): m/z= 525.3 (M+H)^+^

To a solution of the intermediate above (60 mg, 0.11mmol) in DCM (2.0 mL) was added TFA (0.5 mL). After stirring at 20 °C for 4 h, the reaction mixture was concentrated to dryness, the resulting residue was suspended in aqueous saturated NaHCO3 and extracted with DCM (10 mL × 3). The organic layers were washed with brine, concentrated under reduced pressure and purified by preprarative HPLC (10%-50% 0.1% NH_4_HCO_3_ in water/ACN) to obtain **AVI-6319** as a white solid (27 mg, yield: 62.3%). ^1^H NMR (500 MHz, DMSO) δ 12.47 (s, 1H), 9.88 (s, 1H), 8.25 (s, 1H), 8.18 (s, 1H), 7.13 (s, 1H), 4.50 (d, *J* = 76.4 Hz, 2H), 4.07 (s, 3H), 3.91 (dd, *J* = 11.8, 4.2 Hz, 1H), 3.78 (d, *J* = 9.5 Hz, 1H), 3.46 (s, 1H), 3.26 (d, *J* = 12.2 Hz, 1H), 1.96 (td, *J* = 12.9, 4.7 Hz, 1H), 1.85 (s, 1H), 1.76 – 1.56 (m, 2H). LCMS (ESI): m/z= 395.1 (M+H)^+^

### Synthesis of 1-methyl-4-(4-((2-oxo-3-oxa-1,8-diazaspiro[4.5]decan-1-yl)amino)-7H-pyrrolo[2,3-d]pyrimidinyl)-1H-pyrazole-5-carbonitrile (AVI-6345)

To a solution of 4- (4-chloro-7H-pyrrolo[2,3-d]pyrimidin-6-yl)-1-methyl-1H-pyrazole-5-carbonitrile (500 mg, 1.93 mmol) in IPA (15 mL) were added benzyl 1-amino-2-oxo-3-oxa-1,8-diazaspiro[4.5]decane-8-carboxylate (707 mg, 2.32 mmol) and conc. HCl (0.06 mL). After stirring at 100°C for 24 hours, aqueous saturated sodium bicarbonate (5 mL) was added to the reaction mixture and extracted with DCM (20 mL×3), organic layers were separated and concentrated in vacuo and purified by silica gel chromatography (dichloromethane/methanol, 10:1) to obtain benzyl 1-((6-(5-cyano-1-methyl-1H-pyrazol-4-yl)-7H-pyrrolo[2,3-d]pyrimidin-4-yl)amino)-2-oxo-3-oxa-1,8-diazaspiro[4.5]decane-8-carboxylate (30 mg, Yield: 2.9%) as a white solid. LCMS (ESI): m/z= 528.1 (M+H)^+^

To a solution of the intermediate above (30 mg, 0.057mmol) in acetonitrile (2.0 mL) was added TMSI (23 mg, 0.114 mmol). After stirring at 20 °C for 4 h, the reaction mixture was concentrated to dryness, the resulting residue was suspended in aqueous saturated NaHCO3 and extracted with DCM (10 mL × 3). The organic layers were washed with brine, concentrated under reduced pressure and purified by prep-TLC (DCM/MeOH=10/1) to obtain **AVI-6345** as a white solid (6 mg, Yield: 26.8%). 1H NMR (500 MHz, DMSO) δ 12.44 (s, 1H), 9.85 (s, 1H), 8.24 (s, 1H), 8.17 (s, 1H), 7.13 (s, 1H), 4.39 (d, J = 69.5 Hz, 2H), 4.06 (s, 3H), 2.95 (d, J = 12.4 Hz, 1H), 2.82 (d, J = 12.5 Hz, 1H), 2.64-2.62 (m, 1H), 2.54-2.32 (m, 2H), 1.92 – 1.70 (m, 2H), 1.60-1.52 (m, 2H). LCMS (ESI): m/z= 394.1 (M+H)^+^

### Synthesis of 1-methyl-4-(4-((8-(methylsulfonyl)-2-oxo-3-oxa-1,8-diazaspiro[4.5]decan-1-yl)amino)-7H-pyrrolo[2,3-d]pyrimidin-6-yl)-1H-pyrazole-5-carbonitrile (AVI-6344)

To a solution of 4-(4-chloro-7H-pyrrolo[2,3-d]pyrimidin-6-yl)-1-methyl-1H-pyrazole-5-carbonitrile (100 mg, 0.39 mmol) in IPA (5 mL) were added 1-amino-8-(methylsulfonyl)-3-oxa-1,8-diazaspiro[4.5]decan-2-one (149 mg, 0.60 mmol) and conc. HCl (0.04 mL). After stirring at 100°C for 24 hours, aqueous saturated sodium bicarbonate (5 mL) was added to the reaction mixture and extracted with DCM (20 mL × 3), organic layers were separated and concentrated in vacuo and purified by prep-HPLC (10%-50% 0.1% NH4HCO3 in water/ACN) to obtain **AVI-6344** as a white solid (22 mg, yield: 12.0%). 1H NMR (500 MHz, DMSO) δ 12.47 (s, 1H), 9.88 (s, 1H), 8.25 (s, 1H), 8.18 (s, 1H), 7.02 (d, J = 100.1 Hz, 1H), 4.78 – 4.20 (m, 2H), 4.07 (s, 3H), 3.63 (d, J = 12.0 Hz, 1H), 3.50 (d, J = 12.2 Hz, 1H), 2.93 (s, 1H), 2.86 (s, 3H), 2.76 (dd, J = 22.8, 12.0 Hz, 1H), 2.16 – 1.89 (m, 3H), 1.76 (d, J = 65.1 Hz, 2H). LCMS (ESI): m/z= 472.0 (M+H)+

### Synthesis of 4-(4-((((*trans*)-2-hydroxycyclopentyl)methyl)amino)-7H-pyrrolo[2,3-d]pyrimidin-6-yl)-1-methyl-1H-pyrazole-5-carbonitrile (AVI-6315)

To a solution of 4-(4-chloro-7-((2- (trimethylsilyl)ethoxy)methyl)-7H-pyrrolo[2,3-d]pyrimidin-6-yl)-1-methyl-1H-pyrazole-5-carbonitrile (500 mg, 1.29 mmol) in dry DMSO (10 mL) were added 2-(aminomethyl)cyclopentan-1-ol (223 mg, 1.93 mmol) and TEA (651mg, 6.44mmol) After stirring at 110°C for 3 hours, the reaction mixture was diluted with ethyl acetate (100.0 mL) and washed with water (10.0 mL), brine (10.0 mL). The organic layers were dried over Na_2_SO_4_ and concentrated under reduced pressure. The residue was purified by prep-HPLC (10%-60% 10mmol NH_4_HCO_3_ in water/ACN) to obtain trans and cis diastereomers 4-(4-((((*trans*)-2-hydroxycyclopentyl)methyl)amino)-7-((2-(trimethylsilyl)ethoxy)methyl)-7H-pyrrolo[2,3-d]pyrimidin-6-yl)-1-methyl-1H-pyrazole-5-carbonitrile as a white solid (200mg, yield: 33.2%). ^1^H NMR (500 MHz, DMSO) δ 8.32 (s, 1H), 8.20 (s, 1H), 7.97 (s, 1H), 7.18 (s, 1H), 5.69 (d, *J* = 11.3 Hz, 2H), 4.66 (d, *J* = 4.3 Hz, 1H), 4.21 (s, 3H), 3.99 – 3.91 (m, 1H), 3.66 (dd, *J* = 14.3, 6.3

Hz, 2H), 3.64 – 3.58 (m, 1H), 3.52 – 3.46 (m, 1H), 2.23 – 2.15 (m, 1H), 1.99 – 1.86 (m, 2H), 1.81 – 1.55 (m, 3H), 1.40 (td, *J* = 15.1, 7.2 Hz, 1H), 0.98 – 0.91 (m, 2H), −0.00 (d, *J* = 3.1 Hz, 9H). LCMS (ESI): m/z= 468.2 (M+H)^+^. 4-(4-((((*cis*)-2-hydroxycyclopentyl)methyl)amino)-7-((2-(trimethylsilyl)ethoxy)methyl)-7H-pyrrolo[2,3-d]pyrimidin-6-yl)-1-methyl-1H-pyrazole-5-carbonitrile as a white solid (120mg, yield: 20%). ^1^H NMR (500 MHz, DMSO) δ 8.32 (s, 1H), 8.20 (s, 1H), 8.06 (s, 1H), 7.17 (s, 1H), 5.72 – 5.65 (m, 2H), 5.03 (s, 1H), 4.22 (s, 3H), 4.09 (d, *J* = 20.0 Hz, 1H), 3.78 – 3.70 (m, 1H), 3.66 (t, *J* = 7.9 Hz, 2H), 3.60 – 3.51 (m, 1H), 2.18 – 2.07 (m, 1H), 1.90 – 1.74 (m, 3H), 1.74 – 1.49 (m, 3H), 0.98 – 0.91 (m, 2H), −0.01 (d, *J* = 5.7 Hz, 9H). LCMS (ESI): m/z= 468.1 (M+H)^+^

To a solution of the *trans* diastereomer above (80 mg, 0.17 mmol) in DCM (3 mL) was added TFA (1 mL). After stirring at 20°C for 4h, the reaction mixture was then concentrated to dryness. To the resulting residue was added saturated aqueous NaHCO_3_ (5 mL) and extracted with DCM (30 mL × 2). The organic layers were washed with brine and concentrated under reduced pressure. The crude was purified by prep-HPLC (10%-60% 10mmol NH_4_HCO_3_ in water /ACN) to obtain **AVI-6315** as a white solid (30 mg, yield: 51.9%). ^1^H NMR (500 MHz, DMSO) δ 12.09 (s, 1H), 8.13 (d, *J* = 2.0 Hz, 2H), 7.75 (s, 1H), 7.06 (s, 1H), 4.54 (d, *J* = 4.3 Hz, 1H), 4.05 (s, 3H), 3.87 – 3.76 (m, 1H), 3.50 – 3.43 (m, 1H), 3.37 (d, *J* = 6.5 Hz, 1H), 2.15 – 1.99 (m, 1H), 1.89 – 1.71 (m, 2H), 1.70 – 1.43 (m, 3H), 1.28 (dt, *J* = 15.2, 7.2 Hz, 1H). LCMS (ESI): m/z= 338.1 (M+H)^+^

### Synthesis of 4-(4-((((*cis*)-2-hydroxycyclopentyl)methyl)amino)-7H-pyrrolo[2,3-d]pyrimidin-6-yl)-1-methyl-1H-pyrazole-5-carbonitrile (AVI-6316)

To a solution of the *cis* diastereomer (80 mg, 0.17 mmol) in DCM (3.0 mL) was added TFA (1.0 mL). After stirring at 20°C for 4h, the reaction mixture was then concentrated to dryness. To the resulting residue was added saturated aqueous NaHCO3 (5 mL) and extracted with DCM (30 mL × 2). The organic layers were washed with brine and concentrated under reduced pressure. The crude was purified by prep-HPLC (10%-60% 10mmol NH_4_HCO_3_ in water/ACN) to obtain **AVI-6316** as a white solid (28 mg, yield: 48.5%). ^1^H NMR (500 MHz, DMSO) δ 12.14 (s, 1H), 8.12 (d, *J* = 7.4 Hz, 2H), 7.84 (s, 1H), 7.05 (s, 1H), 5.00 (s, 1H), 4.05 (s, 3H), 3.97 (s, 1H), 3.68 – 3.56 (m, 1H), 3.47 – 3.36 (m, 1H), 2.05 – 1.92 (m, 1H), 1.82 – 1.62 (m, 3H), 1.62 – 1.33 (m, 3H). LCMS (ESI): m/z= 338.1 (M+H)^+^

### Synthesis of 1-methyl-4-(4-(((2-oxocyclopentyl)methyl)amino)-7H-pyrrolo[2,3-d]pyrimidin-6-yl)-1H-pyrazole-5-carbonitrile (AVI-6317)

To a cooled (−78°C) solution of oxalyl chloride (82mg, 0.64mmol) in DCM (3mL) was added DMSO (82 mg, 1.07 mmol) in DCM (1mL) dropwise while stirring at under nitrogen gas. After 30 min, 4-(4-((((*trans*)-2-hydroxycyclopentyl)methyl)amino)-7-((2-(trimethylsilyl)ethoxy)methyl)-7H-pyrrolo[2,3-d]pyrimidin-6-yl)-1-methyl-1H-pyrazole-5-carbonitrile (200 mg, 0.43 mmol) in DCM (1mL) was added dropwise and the mixture was stirred for an additional 90 min at −78°C. Triethylamine (433mg, 4.3mmol) was then added dropwise and the reaction mixture was allowed to warm to room temperature. To this mixture was added water (10 mL) and extracted with DCM (10 mL × 3). The organic layers were washed with water, dried over Na_2_SO_4_, filtered, and concentrated under reduced pressure. The residue was purified by prep-HPLC (10mmol NH_4_HCO_3_ in water, 10%-60% MeCN) to obtain 1-methyl-4-(4-(((2-oxocyclopentyl)methyl)amino)-7-((2-(trimethylsilyl)ethoxy)methyl)-7H-pyrrolo[2,3-d]pyrimidin-6-yl)-1H-pyrazole-5-carbonitrile (90 mg, Yield: 45.2%) as a white solid. LCMS (ESI): m/z= 466 (M+H)^+^

To a solution of the intermediate above (80 mg, 0.19 mmol) in DCM (3 mL) was added TFA (1 mL). After stirring at 20°C for 4h, the reaction mixture was concentrated to dryness. To the resulting residue was added saturated aqueous NaHCO_3_ (5 mL) and extracted with DCM (30 mL × 2). The organic layers were washed with brine and concentrated under reduced pressure. The crude was purified by prep-HPLC (10%-100% 10mmol NH_4_HCO_3_ in water/ ACN) to obtain **AVI-6317** as a white solid (28 mg, yield: 40.7%). ^1^H NMR (500 MHz, DMSO) δ 12.07 (s, 1H), 8.14 (d, *J* = 3.9 Hz, 2H), 7.89 – 7.74 (m, 1H), 7.06 (s, 1H), 4.05 (s, 3H), 3.85 (dt, *J* = 13.2, 5.4 Hz, 1H), 3.37 (dd, *J* = 8.1, 5.7 Hz, 1H), 2.67 – 2.52 (m, 1H), 2.28 – 2.05 (m, 3H), 1.98 – 1.85 (m, 1H), 1.83 – 1.55 (m, 2H). LCMS (ESI): m/z= 336.5 (M+H)^+^

### Synthesis of 4-(4-(((1-(hydroxymethyl)cyclopentyl)methyl)amino)-7H-pyrrolo[2,3-d]pyrimidin-6-yl)-1-methyl-1H-pyrazole-5-carbonitrile (AVI-6254)

To a solution of 4-(4-chloro-7-((2-(trimethylsilyl)ethoxy)methyl)-7H-pyrrolo[2,3-d]pyrimidin-6-yl)-1-methyl-1H-pyrazole-5-carbonitrile (100.0 mg, 0.25 mmol) in DMSO (2 mL) was added (1-(aminomethyl)cyclopentyl)methanol (49.9 mg, 0.38 mmol) and TEA (52.1 mg, 0.51 mmol). After stirred at 100°C for 1 hour, the reaction mixture was extracted with ethyl acetate (3 x 20 mL), washed with brine (10 mL) and dried over Na_2_SO_4_. The organic layers were concentrated and the residue was purified by silica gel chromatography (DCM/MeOH, 20:1) to obtain 4-(4-(((1-(hydroxymethyl)cyclopentyl)methyl)amino)-7-((2-(trimethylsilyl)ethoxy)methyl)-7H-pyrrolo[2,3-d]pyrimidin-6-yl)-1-methyl-1H-pyrazole-5-carbonitrile as a yellow solid (120 mg, Yield: 97%). LCMS (ESI): m/z= 482.4 (M+H)^+^

To a solution of the above intermediate (120 mg, 0.25 mmol) in DCM (2 mL) was added TFA (1.0 mL). After stirring at 20°C for 3 h, the reaction mixture was concentrated to dryness. To the resulting residue was added saturated aqueous NaHCO3 (5 mL) and extracted with DCM (30 mL × 2). The organic layers were washed with brine and concentrated under reduced pressure. The crude was purified by prep-HPLC (10%-50% 0.1% NH_4_HCO_3_ in water/ ACN) to obtain **AVI-6254** as a white solid (29 mg, Yield: 33.33%). 1H NMR (500 MHz, DMSO-d6) δ 12.19 (s, 1H), 8.12 (d, J = 12.0 Hz, 2H), 7.85 (s, 1H), 7.11 (s, 1H), 5.41 (s, 1H), 4.05 (s, 3H), 3.44 (d, J = 6.4 Hz, 2H), 3.14 (d, J = 6.6 Hz, 2H), 1.58 (dd, J = 17.5, 6.9 Hz, 4H), 1.49 – 1.43 (m, 2H), 1.40 – 1.34 (m, 2H). LCMS (ESI): m/z= 352.4 (M+H)^+^

### Synthesis of 4-(4-(((3,3-difluoro-1-(hydroxymethyl)cyclobutyl)methyl)amino)-7H-pyrrolo[2,3-d]pyrimidin-6-yl)-1-methyl-1H-pyrazole-5-carbonitrile (AVI-6253)

To a solution of 4-(4-chloro-7-((2-(trimethylsilyl)ethoxy)methyl)-7H-pyrrolo[2,3-d]pyrimidin-6-yl)-1-methyl-1H-pyrazole-5-carbonitrile (120 mg, 0.31 mmol) in dry DMSO (5 mL) was added (1-(aminomethyl)-3,3-difluorocyclobutyl)methanol (70 mg, 0.46 mmol) and TEA (156 mg, 1.55 mmol). After stirring at 110°C for 3 h, the reaction mixture was diluted with ethyl acetate (50.0 mL), washed with water (10.0 mL) and brine (10.0 mL). The organic layer was dried over Na_2_SO_4_ and concentrated under reduced pressure. The residue was purified by column chromatography on silica gel (DCM/MeOH, 10:1)) to obtain 4-(4-(((3,3-difluoro-1-(hydroxymethyl)cyclobutyl)methyl)amino)-7-((2-(trimethylsilyl)ethoxy)methyl)-7H-pyrrolo[2,3-d]pyrimidin-6-yl)-1-methyl-1H-pyrazole-5-carbonitrile (100 mg, Yield: 64.3%) as a white solid. LCMS (ESI): m/z= 504.3 (M+H)^+^

To a solution of the above intermediate (100 mg, 0.2 mmol) in DCM (3 mL) was added TFA (1 mL). After stirring at 20°C for 4 h, the reaction mixture was concentrated to dryness. To the resulting residue was added saturated aqueous NaHCO3 (5 mL) and extracted with DCM (30 mL × 2). The organic layers were washed with brine and concentrated under reduced pressure. The crude was purified by prep-HPLC (10%-100% 10mmol NH_4_HCO_3_ in water/ACN) to obtain **AVI-6253** as a white solid (27 mg, yield: 36.4%).

### Synthesis of 4-(4-((3-hydroxy-2,2-dimethylpropyl)amino)-7H-pyrrolo[2,3-d]pyrimidin-6-yl)-1-methyl-1H-pyrazole-5-carbonitrile (AVI-6252)

To a solution of 4-(4-chloro-7-((2-(trimethylsilyl)ethoxy)methyl)-7H-pyrrolo[2,3-d]pyrimidin-6-yl)-1-methyl-1H-pyrazole-5-carbonitrile (200 mg, 0.52 mmol) in DMSO (2 mL) was added 3-amino-2,2-dimethylpropan-1-ol (77 mg, 0.75 mmol) and TEA (158 mg,1.56 mmol). After stirring at 100°C for 4 h, the reaction mixture was diluted with dichloromethane (5 mL) and washed with water (10.0 mL x 3). The organic layer was dried over Na_2_SO_4_ and concentrated under reduced pressure. The residue was purified by silica gel chromatography (DCM/MeOH, 10:1) to obtain 4-(4-((3-hydroxy-2,2-dimethylpropyl)amino)-7-((2-(trimethylsilyl)ethoxy)methyl)-7H-pyrrolo[2,3-d]pyrimidin-6-yl)-1-methyl-1H-pyrazole-5-carbonitrile (90 mg, Yield: 38.0%) as a white solid. LCMS (ESI): m/z= 456.2 (M+H)^+^

To a solution of the above intermediate (90 mg, 0.20 mmol) in DCM(2.0 mL) was added TFA (0.5 mL). After stirring at 20 °C for 4 h, the reaction mixture was concentrated to dryness. To the resulting residue was added saturated aqueous NaHCO_3_ (5 mL) and extracted with DCM (30 mL × 2). The organic layers were washed with brine and concentrated under reduced pressure. The crude was purified by prep-HPLC (10%-50% 0.1% NH_4_HCO_3_ in water/MeCN) to obtain **AVI-6252** as a white solid (29 mg, yield: 44.6%). ^1^H NMR (500 MHz, DMSO) δ 12.19 (s, 1H), 8.12 (d, *J* = 11.9 Hz, 2H), 7.83 (s, 1H), 7.14 (d, *J* = 1.4 Hz, 1H), 5.27 (s, 1H), 4.05 (s, 3H), 3.34 (d, *J* = 6.4 Hz, 2H), 3.07 (s, 2H), 0.87 (s, 6H). LCMS (ESI): m/z= 325.3 (M+H)^+^

### Synthesis of 1-(4-((2,2-dimethyl-5-oxopyrrolidin-1-yl)amino)-9H-pyrimido[4,5-b]indol-8-yl)-3-ethylurea (AVI-6347)

To a solution of 1-(4-chloro-9H-pyrimido[4,5-b]indol-8-yl)-3-ethylurea (100 mg, 0.35 mmol) in dry DMSO (3.0 mL) was added 3-amino-4,4-dimethyloxazolidin-2-one (67 mg, 0.52 mmol), Pd_2_(dba)_3_ (32 mg, 0.03 mmol), tri-*tert*-butylphosphine tetrafluoroborate (15 mg, 0.05 mmol) and t-BuONa(83 mg, 0.86 mmol). After stirring at 100 °C for 2 h, the reaction mixture was filtered and the filtrate was purified by reversed phase chromatography (10-32% acetonitrile/water/0.1%TFA). Further purification by silica gel column chromatography (dicholoromethane/methanol 10:1) afforded **AVI-6347** (20 mg, yield: 15%) as a white solid. ^1^H NMR (500 MHz, DMSO-*d*6) δ 11.74 (s, 1H), 9.28 (s, 1H), 8.45 (d, J = 10.6 Hz, 1H), 8.38 (s, 1H), 8.07 (d, J = 7.8 Hz, 1H), 7.61 (d, J = 7.8 Hz, 1H), 7.22 (t, J = 7.9 Hz, 1H), 6.29 (t, J = 5.5 Hz, 1H), 4.29 (s, 2H), 3.22 – 3.16 (m, 2H), 1.34 (s, 6H), 1.11 (t, J = 7.2 Hz, 3H). LCMS (ESI): m/z=384.3 (M+H)^+^

### Synthesis of 1-(4-((2,2-dimethyl-5-oxopyrrolidin-1-yl)amino)-9H-pyrimido[4,5-b]indol-8-yl)-3-(2-methoxyethyl)urea (AVI-6187)

To a solution of 1-((8-amino-9H-pyrido[2,3-b]indol-4-yl)amino)-5,5-dimethylpyrrolidin-2-one (20 mg, 0.064 mmol) and triethylamine (0.026 mL, 0.26 mmol) in THF (1 mL), was added 1-isocyanato-2-methoxyethane (0.013 mg, 0.13 mmol). After stirring at 55 °C for 18 h, the reaction mixture was purified by reverse phase chromatography (water/acetonitrile) to obtain **AVI-6187** as a white solid (4.3 mg; Yield: 16%). 1H NMR (METHANOL-d4, 400 MHz) δ 8.27 (s, 1H), 7.86 (d, 1H, *J*=8.0 Hz), 7.29 (d, 1H, *J*=7.8 Hz), 7.14 (t, 1H, *J*=7.5 Hz), 3.57-3.60 (m, 2H), 3.49-3.51 (m, 2H), 3.44 (s, 3H), 2.61 (t, 2H, *J*=8.0 Hz), 2.21 (br s, 2H), 1.37 (s, 6H). LCMS (ESI): m/z= 412 (M+H)^+^

### Synthesis of 1-(4-((2,2-dimethyl-5-oxopyrrolidin-1-yl)amino)-9H-pyrimido[4,5-b]indol-8-yl)-3-(tetrahydro-2H-pyran-4-yl)urea (AVI-6188)

To a solution of 1-((8-amino-9H-pyrido[2,3-b]indol-4-yl)amino)-5,5-dimethylpyrrolidin-2-one (20 mg, 0.064 mmol) and triethylamine (0.026 mL, 0.26 mmol) in THF (1 mL), was added 4-isocyanatotetrahydro-2H-pyran (0.016 mg, 0.13 mmol). After stirring at 55 °C for 18 h, the reaction mixture was purified by reverse phase chromatography (water/acetonitrile) to obtain **AVI-6188** as a white solid (3.6 mg; Yield: 13%). ^1^H NMR (METHANOL-d_4_, 400 MHz) δ 8.33 (s, 1H), 7.94 (dd, 1H, *J*=1.0, 7.8 Hz), 7.32 (dd, 1H, *J*=0.7, 7.8 Hz), 7.23 (t, 1H, *J*=7.8 Hz), 3.88-4.01 (m, 3H), 3.52-3.59 (m, 2H), 2.60 (t, 2H, *J*=7.9 Hz), 2.20 (br s, 2H), 1.98 (br d, 2H, *J*=2.2 Hz), 1.60 (br s, 2H), 1.39 (s, 6H). LCMS (ESI): m/z= 438 (M+H)^+^

### Synthesis of 5,5-dimethyl-1-((8-(trifluoromethyl)-9H-pyrimido[4,5-b]indol-4-yl)amino)pyrrolidin-2-one (AVI-6371)

To a solution of 4-chloro-8-(trifluoromethyl)-9H-pyrimido[4,5-b]indole (200 mg, 0.74 mmol) in dry DMSO (4 mL) was added 1-amino-5,5-dimethylpyrrolidin-2-one (142 mg, 1.1 mmol) and t-BuOK (414 mg, 3.69 mmol). The mixture was stirred at 100°C for 45 minutes, diluted with ethyl acetate (50.0 mL) and washed with water (10.0 mL), brine (10.0 mL). The organic layers was dried over Na_2_SO_4_ and concentrated under reduced pressure. The residue was purified by reverse phase chromatography (10mM NH_4_HCO_3_ in water, 5%-60% ACN) to give **AVI-6371** as a white solid (25 mg, yield: 9.33%). ^1^H NMR (500 MHz, DMSO-*d*6) δ 12.57 (s, 1H), 9.33 (s, 1H), 8.73 (d, *J* = 7.8 Hz, 1H), 8.49 (s, 1H), 7.77 (d, *J* = 7.7 Hz, 1H), 7.48 (t, *J* = 7.8 Hz, 1H), 2.45 (t, *J* = 7.8 Hz, 2H), 2.04 (t, *J* = 7.7 Hz, 2H), 1.30 (s, 6H). LCMS (ESI): m/z=364.3 (M+H)^+^.

### Synthesis of 4,4-dimethyl-3-((8-(trifluoromethyl)-9H-pyrimido[4,5-b]indol-4-yl)amino)oxazolidin-2-one (AVI-6372)

To a solution of 4-chloro-8-(trifluoromethyl)-9H-pyrimido[4,5-b]indole (50 mg, 0.18 mmol) and 3-amino-4,4-dimethyloxazolidin-2-one (36 mg, 0.27 mmol) in i-PrOH (0.8 mL) was added 1N aqueous HCl (0.2 mL). The reaction mixture was stirred at 100 °C for 18 h, concentrated and the residue was purified by reverse phase chromatography (water/10-100% MeCN) to obtain 2.1 mg (3%) of **AVI-6372** as a white colored solid (2.1 mg; Yield: 3%). 1H NMR (METHANOL-d4, 400 MHz) δ 8.50-8.57 (m, 2H), 7.79 (d, 1H, *J*=7.8 Hz), 7.49 (t, 1H, *J*=7.8 Hz), 4.41 (br s, 2H), 1.47 (br s, 6H). LCMS (ESI): m/z= 366 (M+H)^+^

### Synthesis of 5,5-dimethyl-1-((8-methyl-9H-pyrimido[4,5-b]indol-4-yl)amino)pyrrolidin-2-one (AVI-6355)

To a solution of 8-bromo-4-chloro-9H-pyrimido[4,5-b]indole (4 g, 14.23 mmol) in dry DMSO (50 mL) was added 1-amino-5,5-dimethylpyrrolidin-2-one (2.73 g, 21.35 mmol) and t-BuOK (3.99 g, 35.58 mmol). The mixture was stirred at 100°C for 45 minutes, diluted with ethyl acetate (80.0 mL) and washed with water (100.0 mL), brine (100.0 mL). The organic layers were dried over Na_2_SO_4_ and concentrated under reduced pressure. The crude product was purified by silica gel chromatography (dicholoromethane/methanol 10:1) to obtain 1-((8-bromo-9H-pyrimido[4,5-b]indol-4-yl)amino)-5,5-dimethylpyrrolidin-2-one as a brown solid (2 g, yield: 37.66%). ^1^H NMR (500 MHz, DMSO) δ 12.65 (s, 1H), 9.55 (br s, 1H), 8.50 (s, 1H), 8.46 (d, J = 7.8 Hz, 1H), 7.69 (d, J = 7.7 Hz, 1H), 7.29 (t, J = 7.9 Hz, 1H), 2.45 (t, J = 7.9 Hz, 2H), 2.05 (t, J = 7.8 Hz, 2H), 1.29 (s, 6H). LCMS (ESI): m/z=374.1 (M+H) ^+^.

To a solution of the above intermediate (120 mg, 0.32 mmol) in dioxane (2 mL) and H_2_O (0.2 mL) was added methylboronic acid (38 mg, 0.64 mmol), K_2_CO_3_ (110 mg, 0.80 mmol) and Pd(dppf)Cl_2_ (35 mg, 0.05 mmol). The mixture was stirred at 95 °C for 16 hours under N_2_. The reaction mixture was diluted with water (50 mL) and extracted with DCM (3 x 50 mL). The organic layer was dried over Na_2_SO_4_ and concentrated under reduced pressure. The residue was purified by reverse phase chromatography (10mM NH_4_HCO_3_ in water, 5%-35% MeCN) to obtain **AVI-6355** as a white solid (25 mg, yield: 14.63%). ^1^H NMR (500 MHz, DMSO-*d*6) δ 12.08 (s, 1H), 9.05 (s, 1H), 8.39 (s, 1H), 8.23 (t, J = 16.0 Hz, 1H), 7.32 – 7.10 (m, 2H), 2.56 (s, 3H), 2.42 (t, J = 7.8 Hz, 2H), 2.03 (t, J = 7.8 Hz, 2H), 1.27 (d, J = 21.2 Hz, 6H). LCMS (ESI): m/z=310.4 (M+H)^+^.

### Synthesis of 4,4-dimethyl-3-((8-methyl-9H-pyrimido[4,5-b]indol-4-yl)amino)oxazolidin-2-one (AVI-6357)

To a solution of 4-chloro-8-methyl-9H-pyrimido[4,5-b]indole(50 mg, 0.23 mmol) and 3-amino-4,4-dimethyloxazolidin-2-one (45 mg, 0.34 mmol) in i-PrOH (0.8 mL) was added 1N aqueous HCl (0.2 mL). The reaction mixture was stirred at100 °C for 18 h, concentrated and the residue was purified by reverse phase chromatography (water/10-100% MeCN) to obtain **AVI-6357** (18 mg, Yield: 25%) as a cream colored solid. ^1^H NMR (DMSO-d_6_, 400 MHz) δ 12.15 (s, 1H), 9.27 (s, 1H), 8.46 (s, 1H), 8.24 (br d, 1H, *J*=7.3 Hz), 7.22-7.27 (m, 2H), 4.29 (s, 2H), 2.57 (s, 3H), 1.35 (br s, 6H). LCMS (ESI): m/z= 312 (M+H)^+^

### Synthesis of 1-((8-cyclopropyl-9H-pyrimido[4,5-b]indol-4-yl)amino)-5,5-dimethylpyrrolidin-2-one (AVI-6354)

To a solution of 1-((8-bromo-9H-pyrimido[4,5-b]indol-4-yl)amino)-5,5-dimethylpyrrolidin-2-one (200 mg, 0.53 mmol) in dioxane (5 mL) and water (0.5 mL) was added cyclopropylboronic acid (184 mg, 2.14 mmol), K_2_CO_3_ (185 mg, 1.34 mmol) and Pd(dppf)Cl_2_ (58 mg, 0.08 mmol). The mixture was stirred at 95 °C for 16 hours under N_2_. The reaction mixture was then diluted with water (50 mL) and extracted with DCM (3 x 50 mL). The organic layers were dried over Na_2_SO_4_ and concentrated under reduced pressure. The residue was purified by reverse phase chromatography (10mM NH_4_HCO_3_ in water, 5%-50% ACN) to give **AVI-6354** as a white solid (25 mg, yield: 13.91%). ^1^H NMR (500 MHz, DMSO-*d*6) δ 12.22 (s, 1H), 9.05 (s, 1H), 8.40 (s, 1H), 8.22 (d, J = 7.8 Hz, 1H), 7.19 (t, J = 7.7 Hz, 1H), 6.97 (d, J = 7.5 Hz, 1H), 2.43-2.39 (m, 3H), 2.03 (t, J = 7.8 Hz, 2H), 1.29 (s, 6H), 1.08 – 1.02 (m, 2H), 0.80 – 0.75 (m, 2H). LCMS (ESI): m/z=336.3 (M+H)^+^.

### Synthesis of 4,4-dimethyl-3-((8-(cyclopropyl)-9H-pyrimido[4,5-b]indol-4-yl)amino)oxazolidin-2-one (AVI-6373)

To a solution of 4-chloro-8-(cyclopropyl)-9H-pyrimido[4,5-b]indole (50 mg, 0.21 mmol) and 3-amino-4,4-dimethyloxazolidin-2-one (40 mg, 0.31 mmol) in i-PrOH (0.8 mL) was added 1N aqueous HCl (0.2 mL). The reaction mixture was stirred at100 °C for 18 h, concentrated and the residue was purified by reverse phase chromatography (water/10-100% MeCN) to obtain of **AVI-6373** as a white colored solid (2.2 mg; Yield: 3%). ^1^H NMR (METHANOL-d_4_, 400 MHz) δ 8.45 (s, 1H), 8.05 (d, 1H, *J*=7.8 Hz), 7.27 (t, 1H, *J*=7.7 Hz), 7.12 (d, 1H, *J*=7.3 Hz), 4.40 (br s, 2H), 2.28-2.35 (m, 1H), 1.45 (br s, 6H), 1.12 (dd, 2H, *J*=1.8, 8.4 Hz), 0.83 (dd, 2H, *J*=1.7, 5.1 Hz). LCMS (ESI): m/z= 338 (M+H)^+^

### Synthesis of 5,5-dimethyl-1-((6-(trifluoromethyl)-9H-pyrimido[4,5-b]indol-4-yl)amino)pyrrolidin-2-one (AVI-3769)

A solution of 4-chloro-6-(trifluoromethyl)-9H-pyrimido[4,5-b]indole (900 mg, 3.3 mmol) and 1-amino-5,5-dimethylpyrrolidin-2-one hydrochloride (637.6 mg, 4.98 mmol,) in isopropanol (10 mL) and concentrated HCl (1 drop) was stirred at 100°C for 16 hours. The reaction mixture was concentrated under reduced pressure and the residue was purified by reverse phase chromatography ((0.1% NH_4_HCO_3_ in water, 10-100%ACN) to obtain of **AVI-3769** as a white solid (102 mg; Yield: 9.9%). 1H NMR (500 MHz, DMSO) δ 12.55 (s, 1H), 9.47 (s, 1H), 8.93 (s, 1H), 8.46 (s, 1H), 7.75 (d, 1H, J = 8.5 Hz), 7.68 (d, 1H, *J* = 8.5 Hz), 2.45 (t, 2H, *J* = 7.8 Hz), 2.04 (d, 2H, *J* = 7.5 Hz), 1.30 (s, 6H). LCMS (ESI): m/z 364 (M+H)^+^

### Synthesis of-dimethyl-3-((6-(trifluoromethyl)-9H-pyrimido[4,5-b]indol-4-yl)amino)oxazolidin-2-one (AVI-3865)

To a solution of 4-chloro-6-(trifluoromethyl)-9H-pyrimido[4,5-b]indole (50 mg, 0.18 mmol) and 3-amino-4,4-dimethyloxazolidin-2-one (36 mg, 0.27 mmol) in i-PrOH (0.8 mL) was added 1N aqueous HCl (0.2 mL). The reaction mixture was stirred at100 °C for 18 h, concentrated and the residue was purified by reverse phase chromatography (water/10-100% MeCN) to obtain of **AVI-6412** as a white colored solid (6 mg; Yield: 9%). ^1^H NMR (METHANOL-d_4_, 400 MHz) δ 8.68 (s, 1H), 8.47 (s, 1H), 7.69-7.77 (m, 2H), 4.41 (br s, 2H), 1.46 (br s, 6H). LCMS (ESI): m/z= 366 (M+H)^+^

### Synthesis of 1-((6-cyclopropyl-9H-pyrimido[4,5-b]indol-4-yl)amino)-5,5-dimethylpyrrolidin-2-one (AVI-3865)

A neat mixture of 4-chloro-6-cyclopropyl-9H-pyrimido[4,5-b]indole (260 mg, 1.07 mmol) and 1-amino-5,5-dimethylpyrrolidin-2-one hydrochloride (351 mg, 2.14 mmol) was heated at 115°C for 16 h. The reaction mixture was taken in water (10 mL) and extracted with ethyl acetate (3 x 20 mL). The organic layer was dried over Na_2_SO_4_ and concentrated. The residue was purified by reverse phase chromatography (0.1% NH_4_HCO_3_ in water/MeCN) to obtain of **AVI-3865** as a white solid (26 mg; Yield: 7.3%). ^1^H NMR (500 MHz, DMSO) δ 11.94 (s, 1H), 9.10 (s, 1H), 8.34 (s, 1H), 8.08 (s, 1H), 7.38 (d, J = 8.3 Hz, 1H), 7.20 (d, J = 8.3 Hz, 1H), 2.47 – 2.35 (m, 2H), 2.06 (ddd, J = 16.3, 11.1, 6.6 Hz, 3H), 1.27 (d, J = 19.4 Hz, 6H), 1.01 – 0.93 (m, 2H), 0.85 – 0.74 (m, 2H). LCMS (ESI): m/z= 336 (M+H)^+^

### Synthesis of 3-((8-cyclopropyl-6-fluoro-9H-pyrimido[4,5-b]indol-4-yl)amino)-4,4-dimethyloxazolidin-2-one (AVI-6451)

To a solution of 4-chloro-8-cyclopropyl-6-fluoro-9H-pyrimido[4,5-b]indole (900 mg, 3.45 mmol) in i-PrOH (20 mL) was added 3-amino-4,4-dimethyloxazolidin-2-one (897mg, 6.9mmol) and conc. HCl (0.1 mL). After stirring at 100°C for 72 h, the reaction mixture was concentrated and the residue was purified by reverse phase chromatography (0.1% NH4HCO_3_ in water, 10-100%ACN) to obtain **AVI-6451** as a white solid (220 mg, Yield: 17.9%). 1H NMR (500 MHz, DMSO-d6) δ 12.39 (s, 1H), 9.31 (s, 1H), 8.47 (s, 1H), 8.13 (d, *J* = 8.2 Hz, 1H), 6.84 (dd, *J* = 10.8, 2.2 Hz, 1H), 4.29 (s, 2H), 2.48 – 2.42 (m, 1H), 1.46 – 1.27 (m, 6H), 1.09 (dd, J = 8.3, 1.7 Hz, 2H), 0.85 (d, *J* = 4.1 Hz, 2H). LCMS (ESI): m/z 356.1 (M+H)^+^

### Synthesis of 3-((8-cyclopropyl-7-fluoro-9H-pyrimido[4,5-b]indol-4-yl)amino)-4,4-dimethyloxazolidin-2-one (AVI-6452)

To a solution of 4-chloro-8-cyclopropyl-7-fluoro-9H-pyrimido[4,5-b]indole (900mg, 3.45mmol) in isopropanol (20 mL) was added 3-amino-4,4-dimethyloxazolidin-2-one (897mg, 6.9mmol) and conc. HCl (0.1 mL). The mixture was stirred at 100°C for 72 hours, concentrated under reduced pressure and the residue was purified by reverse phase chromatography (0.1% NH_4_HCO_3_ in water, 10-100% MeCN) to obtain **AVI-6452** as a white solid (240 mg, yield: 19.6%). ^1^H NMR (400 MHz, DMSO) δ 12.31 (s, 1H), 9.32 (s, 1H), 8.46 (s, 1H), 8.24 (dt, *J* = 13.8, 6.9 Hz, 1H), 7.10 (dd, *J* = 11.7, 8.7 Hz, 1H), 4.29 (s, 2H), 2.08 (tt, *J* = 8.6, 5.4 Hz, 1H), 1.30 (t, *J* = 26.8 Hz, 6H), 1.18 – 0.98 (m, 2H), 0.92 – 0.76 (m, 2H). LCMS (ESI): m/z 356.2 (M+H)^+^.

### Synthesis of 3-((6-fluoro-8-(1-hydroxycyclopropyl)-9H-pyrimido[4,5-b]indol-4-yl)amino)-4,4-dimethyloxazolidin-2-one (AVI-6612)

To solution of (4-chloro-6-fluoro-9H-pyrimido[4,5-b]indol-8-yl)cyclopropan-1-ol (100 mg, 0.36 mmol) in *i*-PrOH (6 mL) was added 3-amino-4,4-dimethyloxazolidin-2-one (94 mg, 0.72 mmol) and conc. HCl (0.1 mL). After stirring at 100°C for 16 hours, the reaction mixture was concentrated and the residue was purified by reverse phase chromatography (10 mmol NH_4_HCO_3_ in water, 10-100% ACN) to obtain **AVI-6612** as a white solid (15 mg, yield: 11.2%). ^1^H NMR (400 MHz, DMSO) δ 11.73 (d, *J* = 24.3 Hz, 1H), 9.36 (s, 1H), 8.49 (s, 1H), 8.25 (dd, *J* = 9.7, 1.9 Hz, 1H), 7.15 (dd, *J* = 10.2, 2.4 Hz, 1H), 6.10 (d, *J* = 5.1 Hz, 1H), 4.29 (s, 2H), 1.30 (dd, *J* = 31.3, 27.0 Hz, 6H), 1.14 (d, *J* = 13.9 Hz, 2H), 1.08 – 0.89 (m, 2H). LCMS (ESI): m/z 372.1 (M+H)^+^.

### Synthesis of 3-((6-fluoro-8-(1-fluorocyclopropyl)-9H-pyrimido[4,5-b]indol-4-yl)amino)-4,4-dimethyloxazolidin-2-one (AVI-6613)

To a solution of 1-(4-chloro-6-fluoro-9H-pyrimido[4,5-b]indol-8-yl)cyclopropan-1-ol (200 mg, 0.72 mmol) in dichloromethane (10 mL) was added DAST (581 mg, 3.61 mmol) and the mixture was stirred at room temperature for an hour under nitrogen gas. The mixture was then quenched with saturated aqueous NaHCO_3_. The layers were separated and aqueous layer was extracted with ethyl acetate (3 x 30 mL), The combined organic layers were dried over anhydrous Na_2_SO_4_, filtered and concentrated under reduced pressure. The crude residue was then purified by silica gel chromatography (3:1 petroleum ether/ ethyl acetate) to afford 4-chloro-6-fluoro-8-(1-fluorocyclopropyl)-9H-pyrimido[4,5-b]indole as a white solid (150 mg, Yield: 74.5%). LCMS (ESI): m/z=280.2 (M+H)^+^.

To solution of the above intermediate (100 mg, 0.36 mmol) in *i*-PrOH (6 mL) was added 3-amino-4,4-dimethyloxazolidin-2-one (94 mg, 0.72 mmol) and conc. HCl (0.1 mL). The mixture was stirred at 100°C for 16 h, concentrated and the residue was purified by reverse phase chromatography (10 mmol NH_4_HCO_3_ in water, 10-100% ACN) to obtain **AVI-6613** as a white solid (13 mg, yield: 9.76%). ^1^H NMR (400 MHz, DMSO) δ 12.41 (d, *J* = 36.2 Hz, 1H), 9.42 (s, 1H), 8.51 (s, 1H), 8.42 (d, *J* = 9.5 Hz, 1H), 7.40 (d, *J* = 9.7 Hz, 1H), 4.30 (s, 2H), 1.55 (d, *J* = 18.6 Hz, 2H), 1.34 (d, *J* = 33.5 Hz, 6H), 1.20 (t, *J* = 15.1 Hz, 2H). LCMS (ESI): m/z 374.2 (M+H)^+^.

## ASSOCIATED CONTENT

### Supporting Information

The Supporting Information is available free of charge on the ACS Publications website.

Supplementary Figures and Tables. Tables of crystallographic data and refinement statistics. Supplementary synthetic schemes and procedures for the preparation of key intermediates and final analogs not presented in the main text. Additional experimental procedures.

## AUTHOR INFORMATION

### Author Contributions

† P.J., G. J. C., Y. M, and T.T. and contributed equally to this work

### Notes

P.J., G.J.C., T.T., M.M.R., E.R.H., M. D., N.J.K., B.K.S, A.A, M.O., J.S.F, and A.R.R. are listed as inventors on a patent application describing compounds described herein. M.O. and T.Y.T. are listed as inventors on a patent filed by the Gladstone Institutes that covers the use of pGLUE to generate SARS-CoV-2 infectious clones and replicons.

## ACKNOWLEDGMENTS

This work was supported by the National Institutes of Health NIAID Antiviral Drug Discovery (AViDD) grant U19AI171110. M.O. received support from the Roddenberry Foundation, from P. and E. Taft, and the Gladstone Institutes. M.O. is a Chan Zuckerberg Biohub – San Francisco Investigator. The ALS, a U.S. DOE Office of Science User Facility under contract no. DE-AC02-05CH11231, is supported in part by the ALS-ENABLE program funded by the NIH, National Institute of General Medical Sciences, grant P30 GM124169. Use of the SSRL, SLAC National Accelerator Laboratory, is supported by the U.S. Department of Energy, Office of Science, Office of Basic Energy Sciences under Contract No. DE-AC02-76SF00515. The SSRL Structural Molecular Biology Program is supported by the DOE Office of Biological and Environmental Research, and by the National Institutes of Health, National Institute of General Medical Sciences (P30GM133894).Portions of this work were performed on the Wynton HPC Co-Op cluster, which is supported by UCSF research faculty and UCSF institutional funds. The authors wish to thank the UCSF Wynton team for their ongoing technical support of the Wynton environment.

## ABBREVIATIONS

ACN: acetonitrile
DCM: dichloromethane
HPMC: (hydroxypropyl)methyl cellulose
PBMC: peripheral blood mononuclear cells
MDM: monocyte-derived macrophages
TMPRSS2: transmembrane protease, serine 2
HTRF: homogeneous time-resolved fluorescence
HP-βCD: hydroxypropyl-β-cyclodextrin
QD: once daily
BID: twice daily
IP: intraperitoneal
PO: oral administration
DFT: density functional theory
ITC: isothermal titration calorimetry
PARP: Poly(ADP-ribose) polymerase
P-gp: P-glycoprotein
ER: efflux ratio
MLM: mouse liver microsomes
IFN-γ: interferon gamma
M-CSF: macrophage colony-stimulating factor
PBS: phosphate buffered saline
PFU: plaque forming units
hACE2: human angiotensin converting enzyme 2
PK: pharmacokinetic
PD: pharmacodynamic
ADME: absorption distribution metabolism and execretion.

## SYNOPSIS TOC

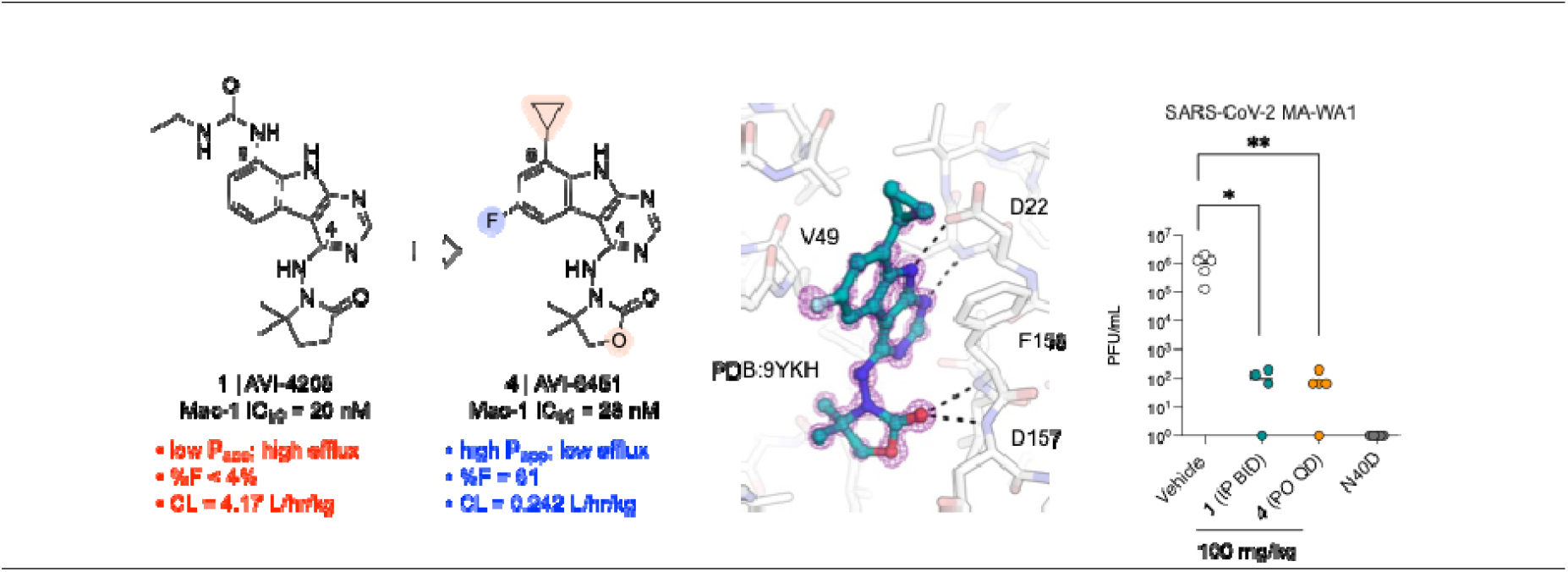

