## Supplementary Information for "Discovery of AVI-6451, a Potent and Selective Inhibitor of the SARS-CoV-2 ADP-Ribosylhydrolase Mac1 with Oral Efficacy in vivo"

**Table of Contents**

Supplementary Figures and Tables……………………………………………S2
Supplementary Experimental Procedures…………………………….…………..S13
Supplementary Synthetic Schemes and Procedures……………..…………....S15

### Supporting Figures and Tables

**Figure S1.** Dose response curves for Mac1 inhibitors as evaluated in the HTRF assay.

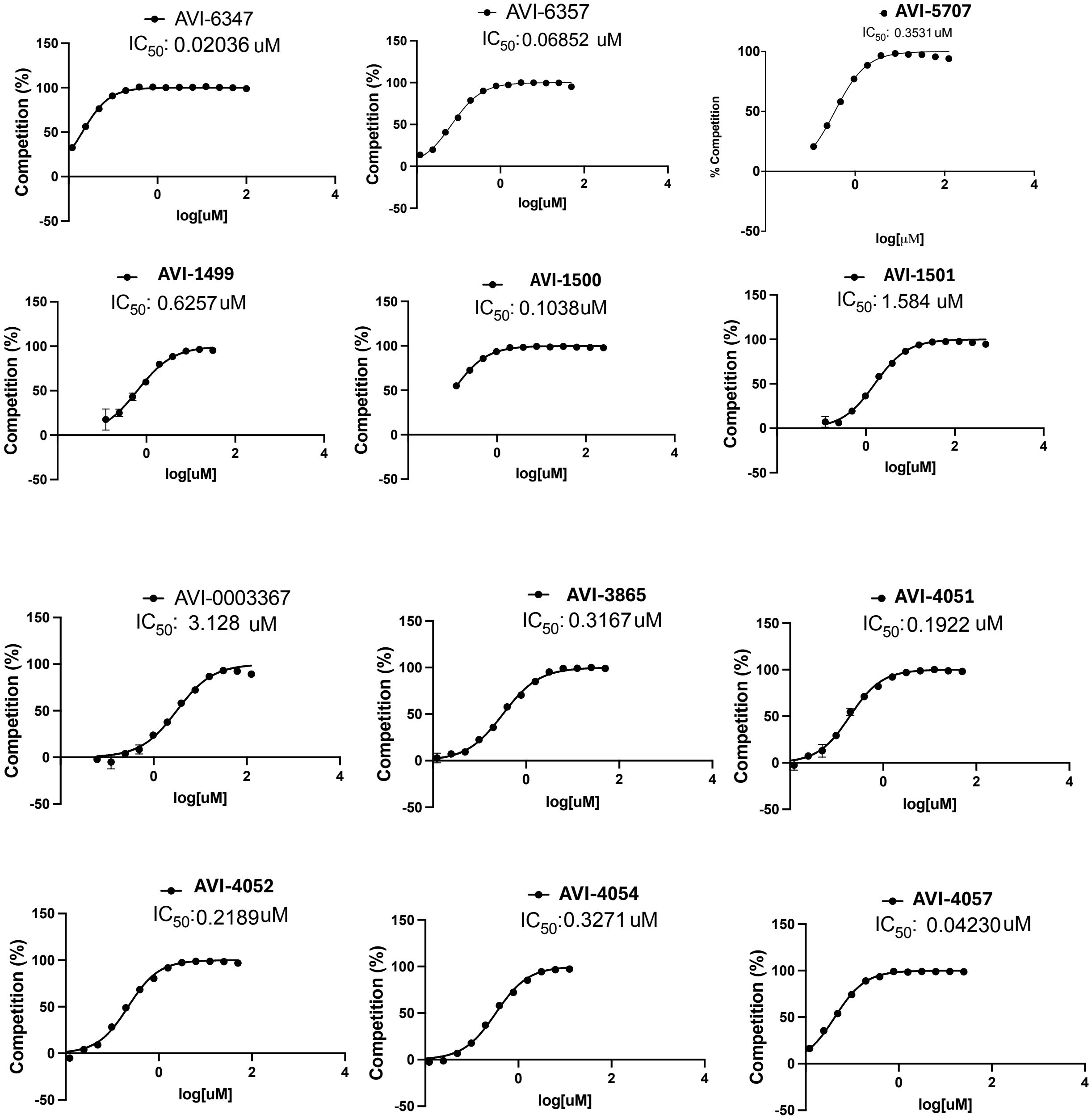

**Figure S1.** continued

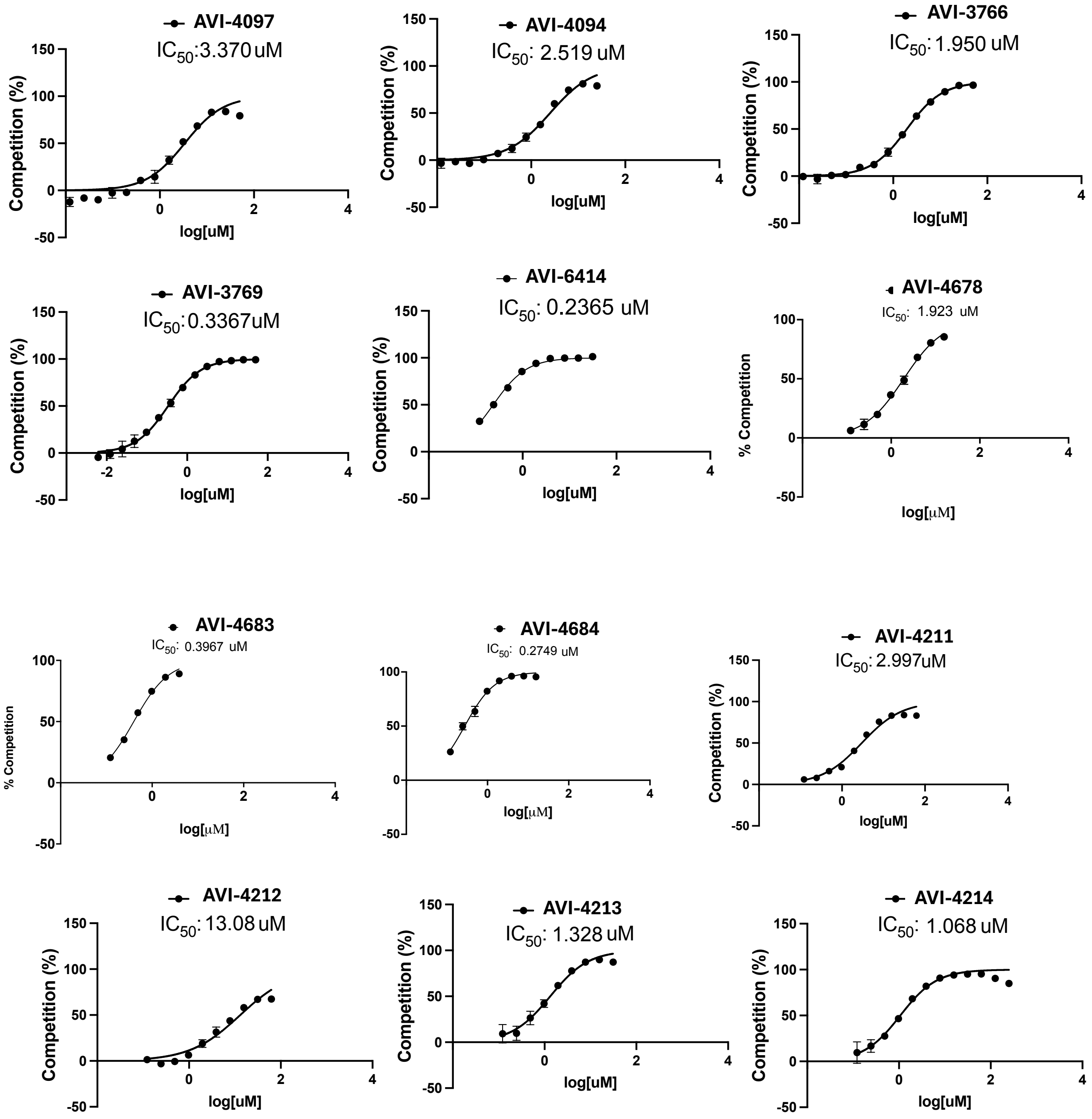

**Figure S1.** continued

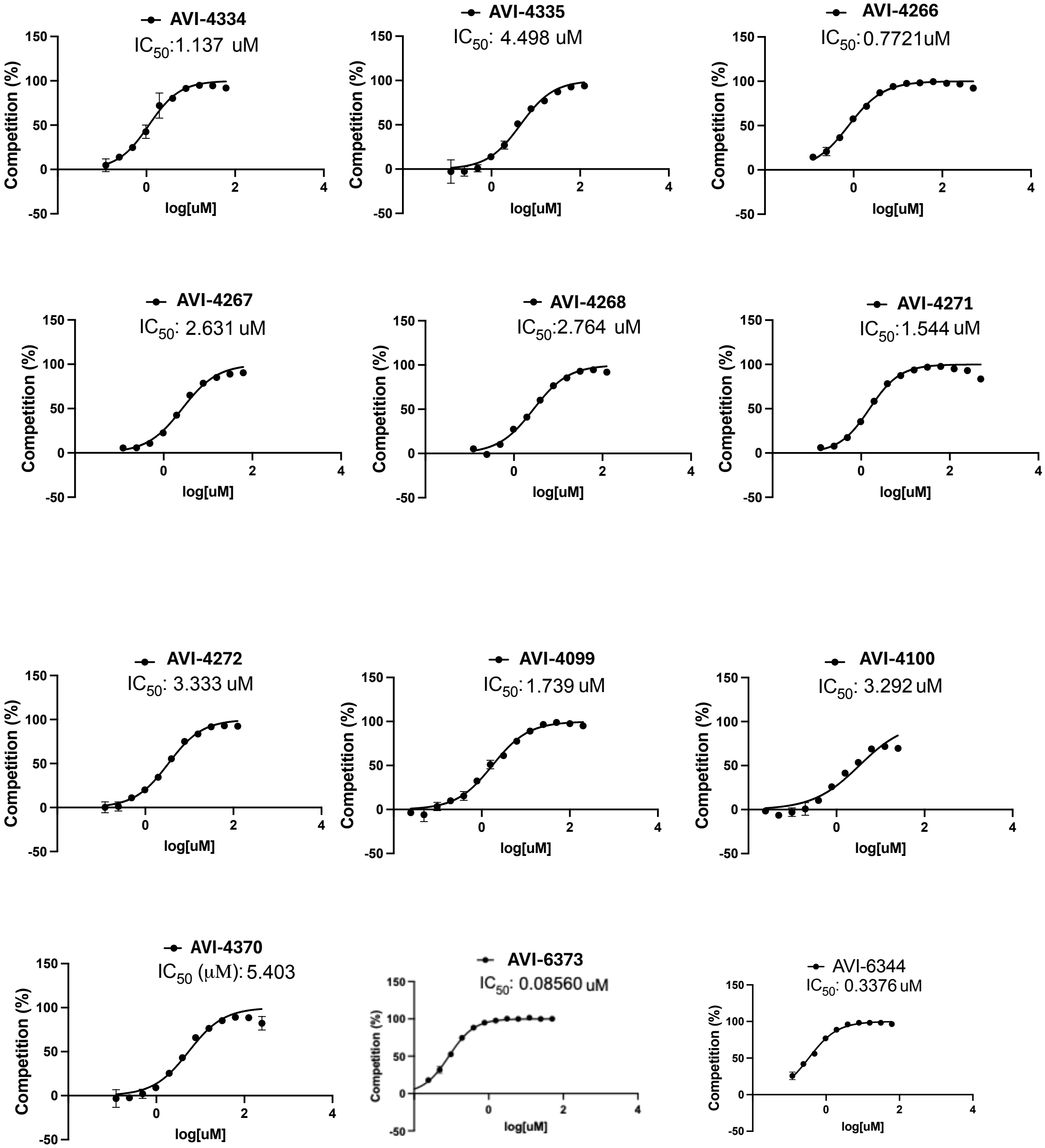

**Figure S1.** continued

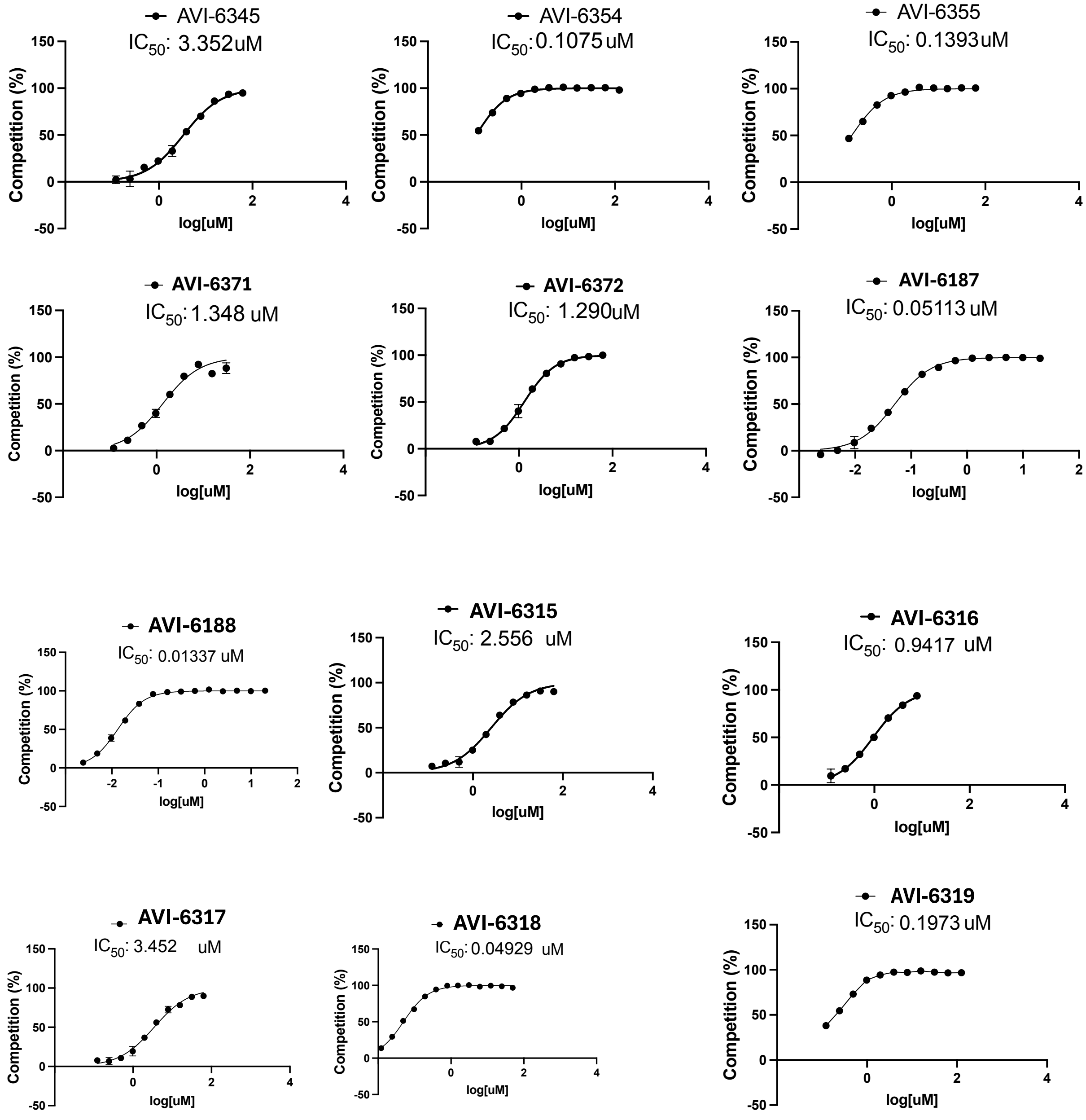

**Figure S1.** continued

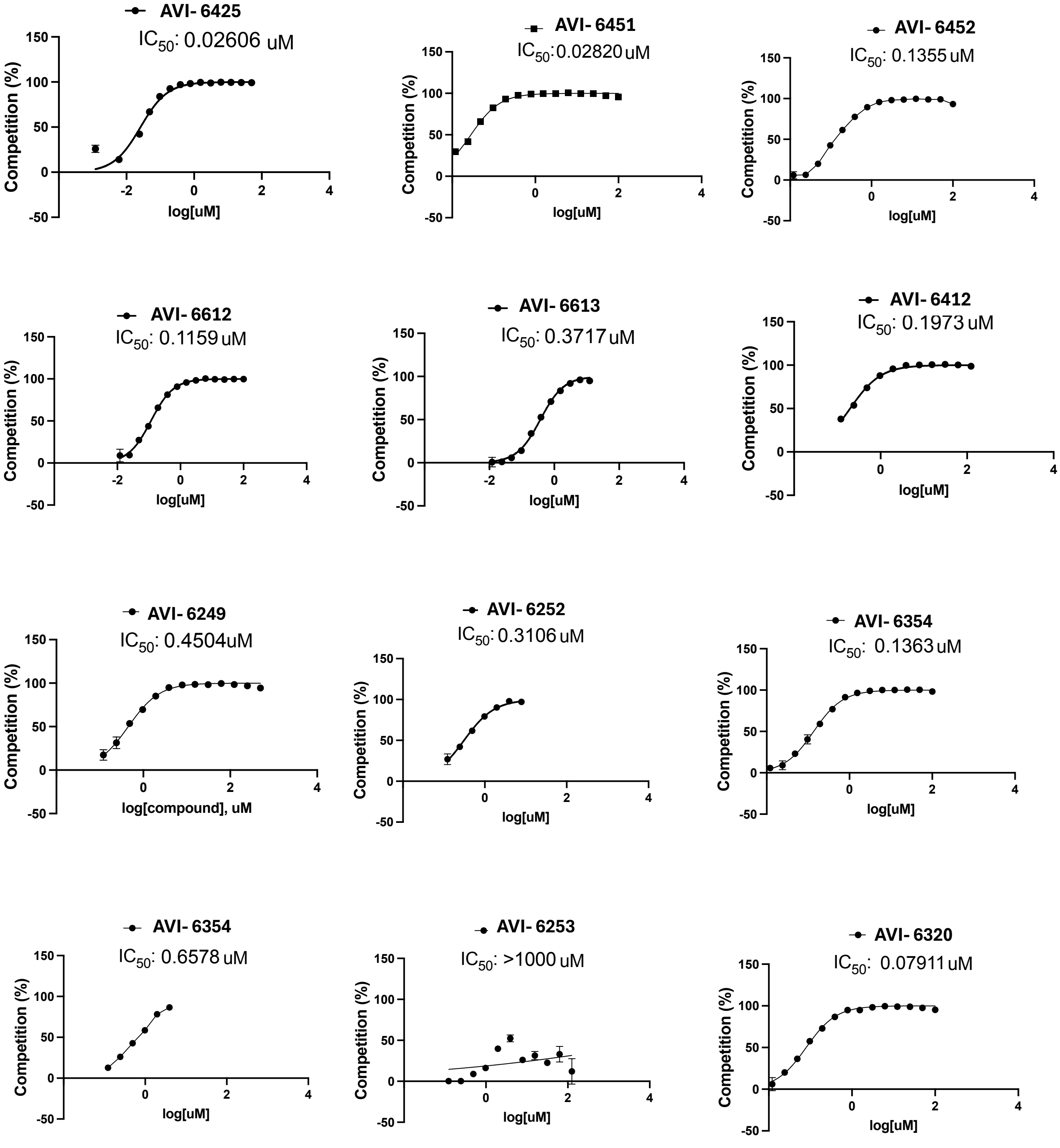

**Table S1A**. Pharmacokinetic parameters for AVI-6451 (**4**) following IV (10 mg/kg) and PO (50 mg/kg) doses in male CD1 mice (n = 9 per group; n = 3 plasma samples per timepoint).

| IV | | | PO | | |
|---|---|---|---|---|---|
| Parameter | units | value | Parameter | units | value |
| CL | L/hr/kg | 0.242 | $T_{max}$ | hr | 2.00 |
| $V_{ss}$ | L/kg | 1.07 | $C_{max}$ | ng/mL | 8863 |
| $T_{1/2}$ | hr | 2.16 | $T_{1/2}$ | hr | 3.91 |
| $AUC_{last}$ | hr*ng/mL | 41350 | $AUC_{last}$ | hr*ng/mL | 124756 |
| $AUC_{INF}$ | hr*ng/mL | 41372 | $AUC_{INF}$ | hr*ng/mL | 126226 |
| $MRT_{INF}$ | hr | 4.42 | F | % | 61.0 |

Data was processed by Phoenix WinNonlin 8.3, and samples below limit of quantitation excluded in the calculation of PK parameters and mean concentration. Liver blood flow (mouse) = 7.2 L/hr/kg

**Table S1B**. Individual sample plasma concentrations of AVI-6451 (**4**) following IV (10 mg/kg) in male CD1 mice (n = 9 per group; n = 3 plasma samples per timepoint).

| Dose (mg/kg) | Dose route | Sampling time (hr) | Concentration (ng/mL) | | | Mean (ng/mL) | SD |
|---|---|---|---|---|---|---|---|
| | | | Individual | | | | |
| 10 | IV | Pre-dose | BQL | BQL | BQL | BQL | NA |
| | | 0.083 | 6570 | 7150 | 6060 | 6593 | 545 |
| | | 0.25 | 5170 | 5030 | 5200 | 5133 | 90.7 |
| LLOQ=1.00 ng/mL | | 0.5 | 5700 | 5640 | 6010 | 5783 | 199 |
| BQL＜LLOQ | | 1 | 5140 | 5450 | 4980 | 5190 | 239 |
| | | 2 | 3920 | 4120 | 5850 | 4630 | 1061 |
| | | 4 | 3540 | 4450 | 3760 | 3917 | 475 |
| | | 8 | 1290 | 1450 | 1560 | 1433 | 136 |
| | | 24 | 7.12 | 5.84 | 8.18 | 7.05 | 1.17 |

**Table S1C**. Individual sample plasma concentrations of AVI-6451 (**4**) following PO (50 mg/kg) in male CD1 mice (n = 9 per group; n = 3 plasma samples per timepoint).

| Dose (mg/kg) | Dose route | Sampling time (hr) | Concentration (ng/mL) | | | Mean (ng/mL) | SD |
|---|---|---|---|---|---|---|---|
| | | | Individual | | | | |
| 50 | PO | Pre-dose | BQL | BQL | BQL | BQL | NA |
| | | 0.083 | 4920 | 1060 | 4170 | 3383 | 2047 |
| | | 0.25 | 8450 | 6790 | 8820 | 8020 | 1081 |
| LLOQ=1.00 ng/mL | | 0.5 | 5700 | 7580 | 6020 | 5570 | 6390 |
| BQL＜LLOQ | | 1 | 5140 | 10400 | 6330 | 8670 | 8467 |
| | | 2 | 7920 | 7770 | 10900 | 8863 | 1765 |
| | | 4 | 6830 | 5490 | 8280 | 6867 | 1395 |
| | | 8 | 7170 | 8630 | 7580 | 7793 | 753 |
| | | 24 | 752 | 14.0 | 16.4 | 261 | 425 |

**Figure S2**. Total mean plasma exposure for AVI-6451 (**4**) following IV (10 mg/kg) and PO (50 mg/kg) doses in male CD1 mice (n = 9 per group; n = 3 plasma samples per timepoint).

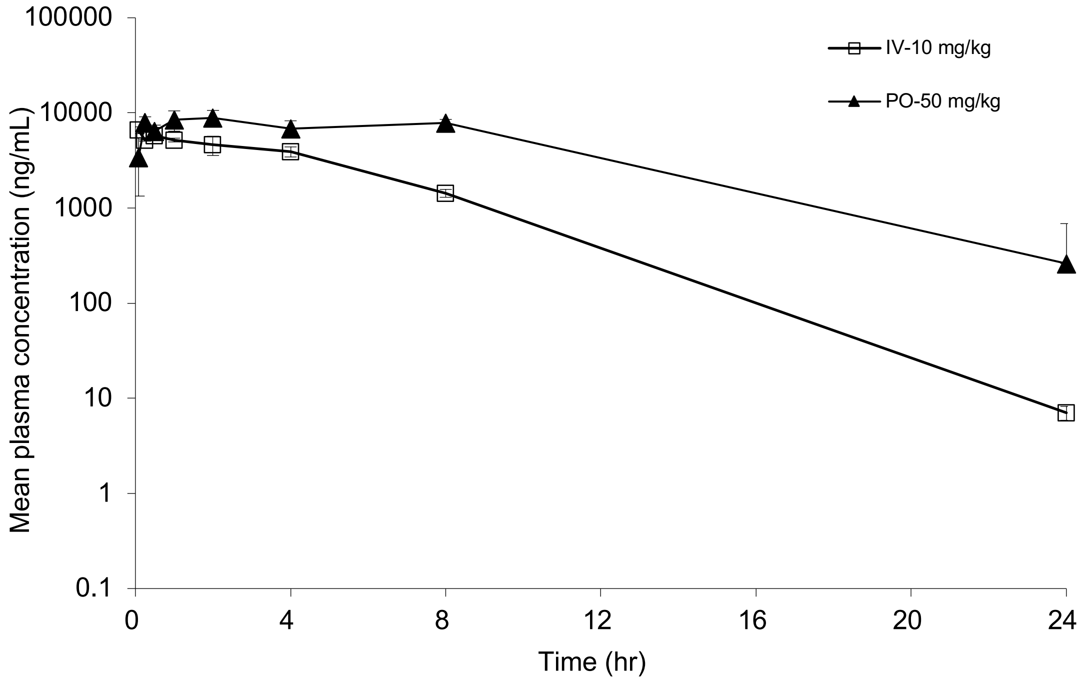

**Table S2A**. Pharmacokinetic parameters for AVI-6452 following IV (10 mg/kg) and PO (50 mg/kg) doses in male CD1 mice (n = 9 per group; n = 3 plasma samples per timepoint).

| IV | | | PO | | |
|---|---|---|---|---|---|
| Parameter | units | value | Parameter | units | mean |
| CL | L/hr/kg | 2.15 | $T_{max}$ | hr | 0.500 |
| $V_{ss}$ | L/kg | 2.09 | $C_{max}$ | ng/mL | 4317 |
| $T_{1/2}$ | hr | 0.870 | $T_{1/2}$ | hr | 1.87 |
| $AUC_{last}$ | hr*ng/mL | 4645 | $AUC_{last}$ | hr*ng/mL | 31356 |
| $AUC_{INF}$ | hr*ng/mL | 4657 | $AUC_{INF}$ | hr*ng/mL | 31362 |
| $MRT_{INF}$ | hr | 0.974 | F | % | 135 |

Data was processed by Phoenix WinNonlin 8.3, and samples below limit of quantitation excluded in the calculation of PK parameters and mean concentration. Liver blood flow (mouse) = 7.2 L/hr/kg

**Table S2B**. Individual sample plasma concentrations of AVI-6452 following IV (10 mg/kg) in male CD1 mice (n = 9 per group; n = 3 plasma samples per timepoint).

| Dose (mg/kg) | Dose route | Sampling time (hr) | Concentration (ng/mL) Individual | | | Mean (ng/mL) | SD |
|---|---|---|---|---|---|---|---|
| 10 | IV | Pre-dose | BQL | BQL | BQL | BQL | N/A |
| | | 0.083 | 4100 | 3990 | 4800 | 4297 | 439 |
| | | 0.25 | 2970 | 3400 | 2850 | 3073 | 289 |
| LLOQ=1.00 ng/mL | | 0.5 | 5700 | 2240 | 2180 | 2800 | 2407 |
| BQL<LLOQ | | 1 | 5140 | 1490 | 1880 | 1940 | 1770 |
| | | 2 | 435 | 905 | 409 | 583 | 279 |
| | | 4 | 24.9 | 55.5 | 51.5 | 44.0 | 16.6 |
| | | 8 | 3.87 | 14.7 | 11.4 | 9.99 | 5.55 |
| | | 24 | BQL | BQL | BQL | BQL | N/A |

**Table S2C**. Individual sample plasma concentrations of AVI-6452 following PO (50 mg/kg) in male CD1 mice (n = 9 per group; n = 3 plasma samples per timepoint).

| Dose (mg/kg) | Dose route | Sampling time (hr) | Concentration (ng/mL) Individual | | | Mean (ng/mL) | SD |
|---|---|---|---|---|---|---|---|
| 50 | PO | Pre-dose | BQL | BQL | BQL | BQL | N/A |
| | | 0.083 | 601 | 1840 | 3230 | 1890 | 1315 |
| | | 0.25 | 2690 | 1750 | 2730 | 2390 | 555 |
| LLOQ=1.00 ng/mL | | 0.5 | 5700 | 3190 | 6150 | 3610 | 4317 |
| BQL<LLOQ | | 1 | 5140 | 3120 | 4850 | 3060 | 3677 |
| | | 2 | 2080 | 1150 | 1820 | 1683 | 480 |
| | | 4 | 1450 | 4510 | 1830 | 2597 | 1668 |
| | | 8 | 2560 | 1580 | 634 | 1591 | 963 |
| | | 24 | 2.35 | 1.94 | BQL | 2.15 | N/A |

**Figure S3**. Total mean plasma exposure for AVI-6452 following IV (10 mg/kg) and PO (50 mg/kg) doses in male CD1 mice (n = 9 per group; n = 3 plasma samples per timepoint).

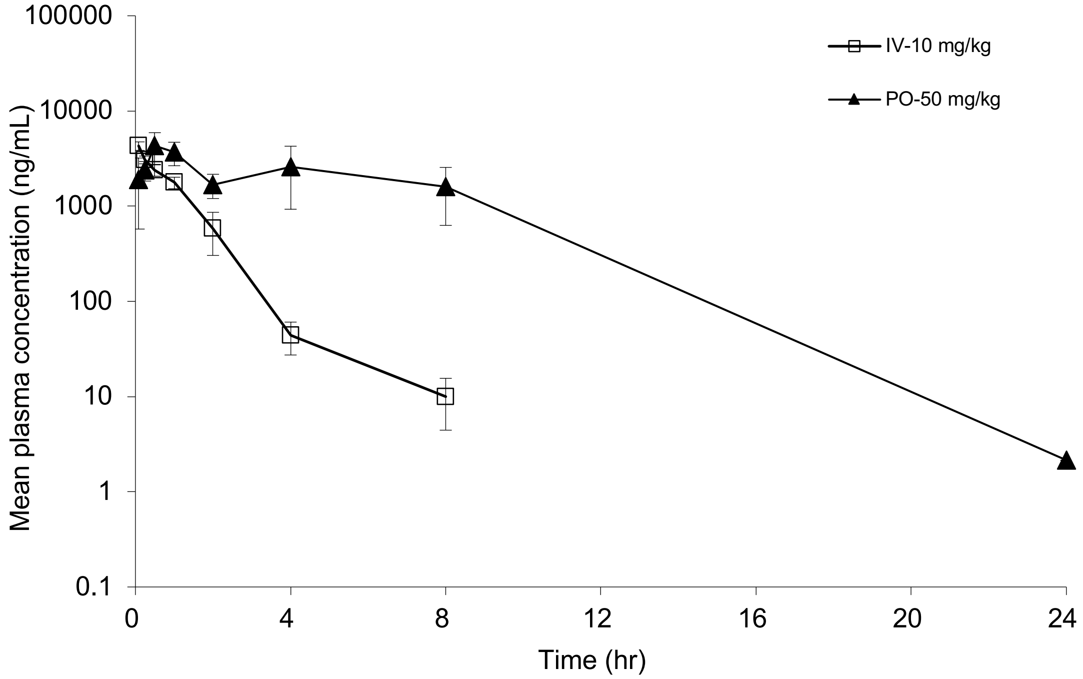

**Table S3.** X-ray data collection and refinement statistics**.**

| Compound | AVI-4094 | AVI-4097 | AVI-4052 | AVI-4054 | AVI-4100 | AVI-4214 | AVI-4268 | AVI-4272 | AVI-4683 |
|---|---|---|---|---|---|---|---|---|---|
| PDB code | 13RI | 13RJ | 13RK | 13RL | 13RM | 13RN | 13RO | 13RP | 13RQ |
| Ligand occupancy (%) | 28 | 24 | 46 | 38 | 74 | 78 | 54 | 60 | 36 |
| Beam line | ALS 8.3.1 | ALS 8.3.1 | ALS 8.3.1 | ALS 8.3.1 | SSRL BL9-2 | SSRL BL9-2 | SSRL BL12-2 | SSRL BL12-2 | ALS 8.3.1 |
| Wavelength (Å) | 0.88557 | 0.88557 | 0.88557 | 0.88557 | 0.88557 | 0.88557 | 0.77488 | 0.77488 | 0.88557 |
| Resolution range | 44.37 - 1.02 (1.057 - 1.02) | 39.68 - 1.0 (1.036 - 1.0) | 39.7 - 1.03 (1.067 - 1.03) | 39.73 - 1.02 (1.056 - 1.02) | 39.73 - 1.01 (1.046 - 1.01) | 39.7 - 1.01 (1.046 - 1.01) | 28.02 - 0.95 (0.984 - 0.95) | 27.99 - 0.95 (0.984 - 0.95) | 39.68 - 0.99 (1.025 - 0.99) |
| Space group | P 43 | P 43 | P 43 | P 43 | P 43 | P 43 | P 43 | P 43 | P 43 |
| Unit cell | 88.734 88.734 39.531 90 90 90 | 88.726 88.726 39.437 90 90 90 | 88.763 88.763 39.705 90 90 90 | 88.85 88.85 39.563 90 90 90 | 88.83 88.83 39.42 90 90 90 | 88.771 88.771 39.407 90 90 90 | 88.622 88.622 39.608 90 90 90 | 88.674 88.674 39.519 90 90 90 | 88.724 88.724 39.642 90 90 90 |
| Total reflections | 1012166 (95535) | 1059181 (85008) | 990983 (95734) | 1017121 (95231) | 1047394 (100527) | 1029083 (99512) | 1299063 (116810) | 1293635 (116113) | 1077438 (78169) |
| Unique reflections | 156326 (15489) | 165156 (16053) | 152764 (15169) | 156871 (15499) | 159518 (15821) | 158934 (15710) | 191840 (18476) | 191803 (18531) | 170401 (16064) |
| Multiplicity | 6.5 (6.2) | 6.4 (5.3) | 6.5 (6.3) | 6.5 (6.1) | 6.6 (6.4) | 6.5 (6.3) | 6.8 (6.3) | 6.7 (6.3) | 6.3 (4.9) |
| Completeness (%) | 99.87 (99.60) | 99.68 (97.24) | 99.87 (99.43) | 99.85 (99.24) | 99.03 (98.86) | 98.80 (98.35) | 99.20 (96.35) | 99.26 (96.47) | 99.31 (94.31) |
| Mean I/sigma(I) | 16.41 (2.03) | 16.26 (1.28) | 13.40 (0.75) | 16.58 (1.61) | 21.47 (2.48) | 18.33 (1.65) | 13.15 (1.07) | 12.05 (1.02) | 11.25 (0.93) |
| Wilson B-factor | 11.07 | 11.63 | 13.02 | 10.44 | 9.44 | 9.62 | 11.39 | 10.9 | 11.52 |
| R-merge | 0.04566 (0.6894) | 0.04365 (1.142) | 0.05084 (2.283) | 0.04964 (1.093) | 0.04024 (0.6721) | 0.04631 (0.9398) | 0.06054 (1.472) | 0.06764 (1.2) | 0.06615 (1.345) |
| R-meas | 0.04972 (0.7539) | 0.04751 (1.267) | 0.05536 (2.489) | 0.05401 (1.195) | 0.04372 (0.7325) | 0.05041 (1.025) | 0.06568 (1.606) | 0.07324 (1.31) | 0.07217 (1.507) |
| R-pim | 0.01944 (0.3002) | 0.01858 (0.5391) | 0.02171 (0.9818) | 0.02105 (0.4752) | 0.01686 (0.2871) | 0.0196 (0.4035) | 0.02511 (0.631) | 0.02778 (0.5166) | 0.02848 (0.6619) |
| CC1/2 | 0.999 (0.884) | 0.999 (0.724) | 0.999 (0.528) | 0.999 (0.772) | 1 (0.857) | 1 (0.723) | 0.997 (0.679) | 0.997 (0.636) | 0.998 (0.612) |
| CC* | 1 (0.969) | 1 (0.916) | 1 (0.831) | 1 (0.933) | 1 (0.961) | 1 (0.916) | 0.999 (0.899) | 0.999 (0.882) | 0.999 (0.872) |
| Reflections used in refinement | 156294 (15489) | 165109 (16053) | 152646 (15170) | 156834 (15501) | 159504 (15821) | 158906 (15710) | 191782 (18476) | 191762 (18531) | 170337 (16064) |
| Reflections used for R-free | 7544 (764) | 7969 (752) | 7390 (808) | 7568 (753) | 7692 (735) | 7658 (731) | 9255 (931) | 9246 (919) | 8227 (760) |
| R-work | 0.1400 (0.2322) | 0.1520 (0.3438) | 0.1644 (0.4521) | 0.1431 (0.2737) | 0.1356 (0.1983) | 0.1418 (0.2532) | 0.1459 (0.3463) | 0.1407 (0.3319) | 0.1461 (0.3826) |
| R-free | 0.1549 (0.2324) | 0.1690 (0.3367) | 0.1856 (0.4458) | 0.1606 (0.2716) | 0.1493 (0.2111) | 0.1555 (0.2666) | 0.1619 (0.3410) | 0.1553 (0.3306) | 0.1610 (0.3758) |
| CC(work) | 0.970 (0.949) | 0.970 (0.884) | 0.971 (0.794) | 0.971 (0.911) | 0.969 (0.940) | 0.967 (0.878) | 0.967 (0.871) | 0.969 (0.837) | 0.971 (0.834) |
| CC(free) | 0.960 (0.950) | 0.959 (0.894) | 0.958 (0.761) | 0.961 (0.906) | 0.965 (0.925) | 0.967 (0.884) | 0.950 (0.875) | 0.962 (0.868) | 0.961 (0.846) |
| Number of non-hydrogen atoms | 3669 | 3668 | 3803 | 3794 | 3820 | 3916 | 3792 | 3771 | 3726 |
| - macromolecules | 3175 | 3188 | 3341 | 3298 | 3346 | 3435 | 3313 | 3300 | 3232 |
| - ligands | 39 | 40 | 44 | 44 | 39 | 43 | 40 | 40 | 41 |
| - solvent | 473 | 459 | 436 | 471 | 452 | 458 | 457 | 449 | 470 |
| Protein residues | 336 | 336 | 336 | 336 | 336 | 336 | 336 | 336 | 336 |
| RMS (bonds) (Å) | 0.005 | 0.004 | 0.005 | 0.004 | 0.005 | 0.004 | 0.004 | 0.004 | 0.005 |
| RMS (angles) (°) | 0.76 | 0.76 | 0.77 | 0.78 | 0.81 | 0.8 | 0.76 | 0.77 | 0.78 |
| Ramachandran favored (%) | 99.4 | 99.4 | 99.4 | 99.4 | 99.1 | 99.1 | 99.1 | 98.49 | 99.4 |
| Ramachandran allowed (%) | 0.6 | 0.6 | 0.6 | 0.6 | 0.9 | 0.9 | 0.9 | 1.51 | 0.6 |
| Ramachandran outliers (%) | 0 | 0 | 0 | 0 | 0 | 0 | 0 | 0 | 0 |
| Rotamer outliers (%) | 0.87 | 2.02 | 2.47 | 1.66 | 1.36 | 1.33 | 1.1 | 1.11 | 1.13 |
| Clashscore | 1.4 | 1.4 | 2.66 | 0.9 | 0.88 | 1.29 | 1.78 | 1.19 | 0.92 |
| Average B-factor | 17.09 | 18.04 | 20.65 | 15.93 | 14.15 | 14.14 | 16.81 | 16.05 | 16.58 |
| - macromolecules | 15.4 | 16.47 | 19.12 | 14.35 | 12.86 | 12.87 | 15.21 | 14.54 | 14.85 |
| - ligands | 11.24 | 11.21 | 17.2 | 9.05 | 7.97 | 8.14 | 11.69 | 8.75 | 11.17 |
| - solvent | 28.67 | 29.24 | 32.59 | 27.36 | 23.96 | 23.99 | 28.65 | 27.56 | 28.76 |

**Table S3 (cont.).** X-ray data collection and refinement statistics.

| Compound | AVI-4678 | AVI-5707 | AVI-6249 | AVI-6318 | AVI-6319 | AVI-6354 | AVI-6344 | AVI-6372 | AVI-6451 |
|---|---|---|---|---|---|---|---|---|---|
| PDB code | 13RR | 13RS | 13RT | 13RU | 13RV | 13RW | 13RX | 13RY | 9YKH |
| Ligand occupancy (%) | 14 | 54 | 44 | 56 | 16 | 32 | 32 | 20 | 100 |
| Beam line | ALS 8.3.1 | SSRL BL12-2 | ALS 8.3.1 | ALS 8.3.1 | ALS 8.3.1 | ALS 8.3.1 | ALS 8.3.1 | ALS 8.3.1 | ALS 8.3.1 |
| Wavelength (Å) | 0.88557 | 0.77488 | 0.88557 | 0.88557 | 0.88557 | 0.88557 | 0.88557 | 0.88557 | 0.8856 |
| Resolution range | 44.4 - 0.99 (1.025 - 0.99) | 29.54 - 0.94 (0.9736 - 0.94) | 44.38 - 1.02 (1.056 - 1.02) | 36.25 - 1.11 (1.15 - 1.11) | 39.66 - 1.0 (1.036 - 1.0) | 36.15 - 1.08 (1.119 - 1.08) | 44.33 - 1.04 (1.077 - 1.04) | 39.67 - 1.01 (1.046 - 1.01) | 39.44 - 1.02 (1.056 - 1.02) |
| Space group | P 43 | P 43 | P 43 | P 43 | P 43 | P 43 | P 43 | P 43 | C 1 2 1 |
| Unit cell | 88.795 88.795 39.49 90 90 90 | 88.891 88.891 39.528 90 90 90 | 88.755 88.755 39.554 90 90 90 | 88.709 88.709 39.716 90 90 90 | 88.682 88.682 39.516 90 90 90 | 88.754 88.754 39.576 90 90 90 | 88.655 88.655 39.63 90 90 90 | 88.701 88.701 39.523 90 90 90 | 129.389 30.339 39.438 90 90.14 90 |
| Total reflections | 1070623 (77609) | 1322402 (107570) | 1009737 (94875) | 798770 (75031) | 1049171 (84890) | 856900 (80124) | 953704 (93723) | 1036225 (92521) | 474741 (40869) |
| Unique reflections | 170644 (16410) | 199694 (19367) | 156414 (15369) | 122108 (12001) | 165335 (16033) | 132067 (12973) | 147774 (14636) | 160786 (15710) | 78093 (7694) |
| Multiplicity | 6.3 (4.7) | 6.6 (5.6) | 6.5 (6.2) | 6.5 (6.2) | 6.3 (5.3) | 6.5 (6.2) | 6.5 (6.4) | 6.4 (5.9) | 6.1 (5.3) |
| Completeness (%) | 99.64 (96.73) | 99.64 (97.27) | 99.77 (98.70) | 99.82 (98.86) | 99.66 (97.12) | 99.83 (98.98) | 99.91 (99.52) | 99.74 (98.04) | 99.71 (99.18) |
| Mean I/sigma(I) | 9.92 (0.69) | 19.02 (0.94) | 15.52 (0.77) | 13.39 (0.68) | 13.80 (0.98) | 11.98 (0.75) | 12.42 (0.76) | 14.92 (0.71) | 11.64 (3.10) |
| Wilson B-factor | 11.76 | 10.48 | 12.7 | 13.5 | 12.05 | 12.49 | 12.37 | 12.07 | 7.1 |
| R-merge | 0.06947 (1.409) | 0.03637 (1.438) | 0.04446 (2.074) | 0.06272 (2.41) | 0.04729 (1.266) | 0.0703 (2.161) | 0.05882 (2.184) | 0.04818 (2.176) | 0.08503 (0.3812) |
| R-meas | 0.07566 (1.585) | 0.03946 (1.587) | 0.04841 (2.267) | 0.0682 (2.633) | 0.05151 (1.406) | 0.07654 (2.36) | 0.06405 (2.378) | 0.05243 (2.389) | 0.09289 (0.4223) |
| R-pim | 0.02969 (0.7121) | 0.01515 (0.66) | 0.01891 (0.8999) | 0.02657 (1.051) | 0.02023 (0.599) | 0.02997 (0.9376) | 0.02513 (0.9329) | 0.02048 (0.9707) | 0.03688 (0.1781) |
| CC1/2 | 0.998 (0.521) | 0.999 (0.482) | 1 (0.482) | 1 (0.425) | 0.999 (0.659) | 0.999 (0.44) | 1 (0.491) | 1 (0.466) | 0.997 (0.909) |
| CC* | 1 (0.827) | 1 (0.806) | 1 (0.806) | 1 (0.773) | 1 (0.891) | 1 (0.782) | 1 (0.812) | 1 (0.797) | 0.999 (0.976) |
| Reflections used in refinement | 170539 (16410) | 199670 (19367) | 156332 (15369) | 121936 (12002) | 165249 (16032) | 132000 (12973) | 147718 (14636) | 160633 (15710) | 78089 (7694) |
| Reflections used for R-free | 8216 (761) | 9569 (866) | 7543 (758) | 5875 (562) | 7968 (743) | 6336 (628) | 7137 (742) | 7741 (736) | 3968 (374) |
| R-work | 0.1482 (0.3887) | 0.1440 (0.3327) | 0.1520 (0.3860) | 0.1728 (0.4499) | 0.1539 (0.3904) | 0.1581 (0.4034) | 0.1593 (0.4530) | 0.1575 (0.4074) | 0.2165 (0.2287) |
| R-free | 0.1645 (0.3631) | 0.1591 (0.3256) | 0.1736 (0.3764) | 0.2007 (0.4819) | 0.1717 (0.3829) | 0.1817 (0.3954) | 0.1823 (0.4662) | 0.1777 (0.4092) | 0.2393 (0.2647) |
| CC(work) | 0.970 (0.801) | 0.970 (0.768) | 0.971 (0.782) | 0.969 (0.750) | 0.970 (0.868) | 0.971 (0.755) | 0.972 (0.788) | 0.970 (0.763) | 0.926 (0.874) |
| CC(free) | 0.959 (0.796) | 0.962 (0.804) | 0.960 (0.805) | 0.958 (0.707) | 0.958 (0.873) | 0.956 (0.756) | 0.958 (0.770) | 0.958 (0.745) | 0.924 (0.809) |
| Number of non-hydrogen atoms | 3749 | 3812 | 3788 | 3771 | 3812 | 3819 | 3806 | 3772 | 1469 |
| - macromolecules | 3266 | 3303 | 3303 | 3328 | 3320 | 3335 | 3313 | 3297 | 1248 |
| - ligands | 43 | 44 | 45 | 42 | 47 | 47 | 54 | 41 | 44 |
| - solvent | 459 | 484 | 456 | 417 | 463 | 458 | 460 | 448 | 195 |
| Protein residues | 336 | 336 | 336 | 336 | 336 | 336 | 336 | 336 | 165 |
| RMS (bonds) (Å) | 0.004 | 0.004 | 0.005 | 0.005 | 0.004 | 0.005 | 0.005 | 0.004 | 0.006 |
| RMS (angles) (°) | 0.75 | 0.78 | 0.76 | 0.79 | 0.76 | 0.75 | 0.75 | 0.74 | 0.86 |
| Ramachandran favored (%) | 99.4 | 99.4 | 99.4 | 99.4 | 99.4 | 99.4 | 99.4 | 99.4 | 98.77 |
| Ramachandran allowed (%) | 0.6 | 0.6 | 0.6 | 0.6 | 0.6 | 0.6 | 0.6 | 0.6 | 1.23 |
| Ramachandran outliers (%) | 0 | 0 | 0 | 0 | 0 | 0 | 0 | 0 | 0 |
| Rotamer outliers (%) | 0.84 | 1.38 | 2.22 | 2.47 | 1.66 | 2.47 | 1.38 | 0.83 | 0.73 |
| Clashscore | 0.91 | 0.6 | 1.64 | 1.63 | 3.72 | 1.78 | 1.94 | 1.95 | 1.58 |
| Average B-factor | 18.43 | 16.09 | 19.28 | 21.63 | 18.98 | 19.31 | 19.43 | 18.61 | 10.55 |
| - macromolecules | 16.73 | 14.36 | 17.52 | 20.19 | 17.24 | 17.52 | 17.66 | 16.9 | 9.34 |
| - ligands | 16.3 | 9.08 | 20.39 | 18.21 | 17.89 | 23.04 | 16.4 | 20.65 | 7.92 |
| - solvent | 30.64 | 28.23 | 31.9 | 33.29 | 31.53 | 32.15 | 32.41 | 31.05 | 18.6 |

#### Supplementary Experimental Procedures

In vitro ADME assays were performed by Quintara Discovery (a Frontage Labs company) in Hayward, CA, using the procedures provided below.

**Kinetic solubility**
Compounds dissolved in DMSO (10 mM) were diluted with 190 µL of phosphate buffered saline (pH 7.4) in 96-well solubility filter plates (Millipore, MSSLBPC50). Solutions were mixed by shaking for 1.5 hours at room temperature and samples were filtered by vacuum into fresh 96-well plates. Compound concentration was determined by absorbance (220 nm, 254 nm, and 280 nm) relative to a three-point standard curve prepared in DMSO (500, 50 and 5 µM). Mean solubility was determined from triplicate measurements.

**Bi-directional Transport in Caco-2 Cells**
Caco-2 cell plates were obtained commercially and were maintained for 21 days at 37°C with 5% $CO_2$. Cells were washed with Hank's Balanced Salt Solution (HBSS) with 5 mM HEPES for 30 minutes before starting the experiment. Test compound solutions are prepared by diluting DMSO stock into HBSS buffer, resulting in a final DMSO concentration of 0.1%. Prior to the assay, cell monolayer integrity was verified by transendothelial electrical resistance (TEER) to ensure that wells have resistance above the acceptance cut-off (1 kΩ). Transport experiments were initiated by adding test compounds to the apical (75 µL) or basal (250 µL) side. Transport plates were incubated at 37°C in a humidified incubator with 5% $CO_2$. Samples were taken from the donor and acceptor compartments after one hour and analyzed by liquid chromatography with tandem mass spectrometry (LC/MS/MS) using an AB Sciex API 4000 instrument, coupled to a Shimadzu LC-20AD LC Pump system. Samples were separated using a Waters Atlantis T3 dC18 reverse phase HPLC column (20 mm x 2.1 mm) at a flow rate of 0.5 ml/min. The mobile phase consisted of 0.1% formic acid in acetonitrile (solvent A) and 0.1% formic acid in water (solvent B).

Apparent permeability ($P_{app}$) values were calculated using the following equation:

$$P_{app} = (dQ/dt) / A / C_0$$

where dQ/dt is the initial rate of compound transport across the cell monolayer, A is the surface area of the filter membrane, and $C_0$ is the initial concentration of the test compound, calculated for each direction using a four-point calibration curve by LC/MS/MS.

Net flux ratio between the two directional transports was calculated by the following equation:

$$\text{Efflux Ratio (ER)} = P_{app}\text{B-A} / P_{app}\text{A-B}$$

where $P_{app}$B-A and $P_{app}$A-B represent the apparent permeability of compounds from the basal-to-apical and apical-to-basal side of the cellular monolayer, respectively. Recovery was calculated based on the compound concentration at the end of the experiment, compared to that at the beginning of the experiment, adjusted for volumes.

**Microsomal stability**

The assay was carried out in 96-well plates at 37°C. Reaction mixtures (25 µL) contained a final concentration of 1 µM compound, 0.5 mg/mL liver microsomes protein, and 1 mM NADPH and/or 1 mM UDPGA (with alamethicin) in 100 mM potassium phosphate (pH 7.4) with 3 mM $MgCl_2$. After 0, 15, 30, and 60 minutes, 150 µL of quench solution (100% acetonitrile with 0.1% formic acid) with internal standard was transferred to each well. Reactions containing the same components except the NADPH were prepared as the negative control. Plates were sealed, vortexed, and centrifuged at 4°C for 15 minutes at 4000 rpm. The supernatant was transferred to fresh plates for LC/MS/MS analysis using an AB Sciex API 4000 instrument, coupled to a Shimadzu LC-20AD LC Pump system. Analytical samples are separated using a Waters Atlantis T3 dC18 reverse phase HPLC column (20 mm x 2.1 mm) at a flow rate of 0.5 ml/min. The mobile phase consisted of 0.1% formic acid in water (solvent A) and 0.1% formic acid in 100% acetonitrile (solvent B).

The extent of metabolism was calculated as the disappearance of compounds, compared to the 0-min time incubation. Initial rates were calculated for the compound concentration and used to determine t1/2 values and subsequently, the intrinsic clearance, CLint = (0.693)(1/ t1/2 (min))(g of liver / kg of body weight)(ml incubation/mg of microsomal protein)(45 mg of microsomal protein/g of liver weight).

#### Supplementary Synthetic Schemes and Procedures:

##### 4-chloro-9H-pyrimido[4,5-b]indol-8-amine

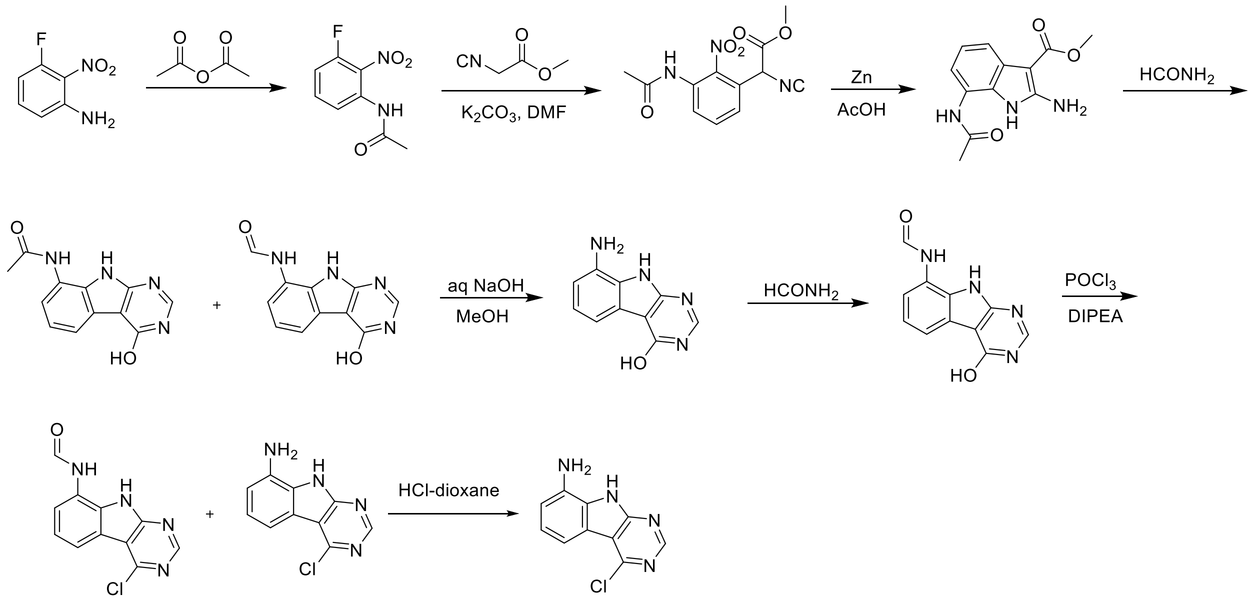


A solution 3-fluoro-2-nitroaniline (11 g, 70.51 mmol) in acetic anhydride (20 mL) was stirred at room temperature for 16 hours. The reaction mixture was filtered and the solids were washed with petroleum ether (100 ml) and dried to obtain 10.7 g (77%) of N-(3-fluoro-2-nitrophenyl)acetamide as a brown solid. LCMS (ESI): m/z= 199.3 $(M+H)^+$

To a solution of N-(3-fluoro-2-nitrophenyl)acetamide (10.7 g, 54.04 mmol) in DMF (100 mL) was added methyl 2-isocyanoacetate (8.02 g, 81.06 mmol) and potassium carbonate (14.92 g, 108.08 mmol). After stirring at 80°C for 2 hours, the reaction mixture was cooled to room temperature, acidified with 2N HCl (ca. 2000 mL), and extracted with ethyl acetate (300 mL *3). The combined organic layers were washed with brine (100 mL), dried over Sodium sulfate and concentrated under reduced pressure. The residue was purified by silica gel chromatography (10: 1 petroleum ether/ethyl acetate) to obtain 11 g (73%) of methyl 2-(3-acetamido-2-nitrophenyl)-2-isocyanoacetate as a yellow solid. LCMS (ESI): m/z= 278.2 $(M+H)^+$

To a solution of methyl 2-(3-acetamido-2-nitrophenyl)-2-isocyanoacetate (11 g, 39.71 mmol) in *glacial* acetic acid (100 ml), was added slowly zinc dust (25.81 g, 397.10 mmol) in two portions. After stirring at 60°C for 2 h, the reaction mixture was cooled to room temperature, filtered and washed with THF. The filtrate was concentrated under reduced pressure and purified by silica gel chromatography (10:1 dichloromethane/methanol) to obtain 6.2 g (63%) of methyl 7-acetamido-2-amino-1H-indole-3-carboxylate as a yellow solid. LCMS (ESI): m/z=248.3 $(M+H)^+$

A solution of methyl 7-acetamido-2-amino-1H-indole-3-carboxylate (6.2 g, 25.10 mmol) in formamide (450 mL) was stirred at 220°C for 2 hours. The reaction mixture was then cooled to room temperature and poured in 100 ml of water. The resulting mixture was allowed to stand for 15 min before the solids were collected by filtration, washed with water, and dried to obtain 4.1 g of a 1:2 mixture of N-(4-hydroxy-9H-pyrimido[4,5-b]indol-8-yl)acetamide and N-(4-hydroxy-9H-pyrimido[4,5-b]indol-8-yl)formamide. This mixture was taken in methanol (25 mL) and aqueous 12 N NaOH (25 ml). After stirring at 60°C for 16 h, the reaction mixture was then cooled to room temperature, concentrated under reduced pressure to remove methanol and the residue was poured into 100 mL of water. The resulting mixture was allowed to stand for 15 min before the solids were collected by filtration, washed with water, and dried to obtain 3.5 g (70%) of 8-amino-9H-pyrimido[4,5-b]indol-4-ol as a brown solid. LCMS (ESI): m/z=201.2 $(M+H)^+$

A solution of 8-amino-9H-pyrimido[4,5-b]indol-4-ol (3.5g, 17.5mmol) in formamide (30 mL) was stirred at 150°C. After 6 h, the reaction mixture was cooled to room temperature and poured into water (200 ml). The resulting mixture was allowed to stand for 15 min before the solids were collected by filtration, washed with water, and dried to obtain 3.5 g (88%) of N-(4-hydroxy-9H-pyrimido[4,5-b]indol-8-yl)formamide as a brown solid. LCMS (ESI): m/z=229.2 $(M+H)^+$

To a solution of N-(4-hydroxy-9H-pyrimido[4,5-b]indol-8-yl)formamide (3.5 g, 15.35 mmol) in phosphorous oxychloride (30 mL) was added N,N-diiisopropylethylamine (5.94 g, 46.05 mmol), After refluxing for 16 hours, the reaction mixture was cooled to room temperature, concentrated and poured into water (20 mL). The resulting solid was filtered to obtain 500 mg of a mixture of N-(4-chloro-9H-pyrimido[4,5-b]indol-8-yl)formamide and 4-chloro-9H-pyrimido[4,5-b]indol-8-amine as a black solid. This mixture was taken in 4 N HCl in dioxane (15 mL). After stirring at room temperature for 4 h, reaction mixture was concentrated under reduced pressure, the residue was adjusted to pH7 with aq.$Na_2CO_3$, and extracted with EA (3 x 30 mL). The organic layers was dried over Sodium sulfate, concentrated under reduced pressure and the residue was purified by reverse phase chromatography (water/acetonitrile /0.1% ammonium bicarbonate) to obtain 320 mg (10%) of 4-chloro-9H-pyrimido[4,5-b]indol-8-amine as a white solid. $^1$H NMR (DMSO-$d_6$, 400 MHz) δ 12.99 (br s, 1H), 8.62 (s, 1H), 7.92 (br d, 1H, *J*=7.5 Hz), 7.27 (t, 1H, *J*=7.9 Hz), 7.05 (br d, 1H, *J*=7.5 Hz), 3.70 (br t, 2H, *J*=6.9 Hz), 2.44-2.53 (m, 2H), 2.20 (br t, 2H, *J*=7.4 Hz). $^{13}$C NMR (METHANOL-$d_4$, 100 MHz) δ 175.9, 155.9, 154.3, 153.2, 132.5, 125.7, 121.9, 119.4, 111.3, 111.1, 97.0, 48.6, 28.5, 15.9

LCMS (ESI): m/z= 283 $(M+H)^+$

**4-chloro-6-(trifluoromethyl)-9H-pyrimido[4,5-b]indole**

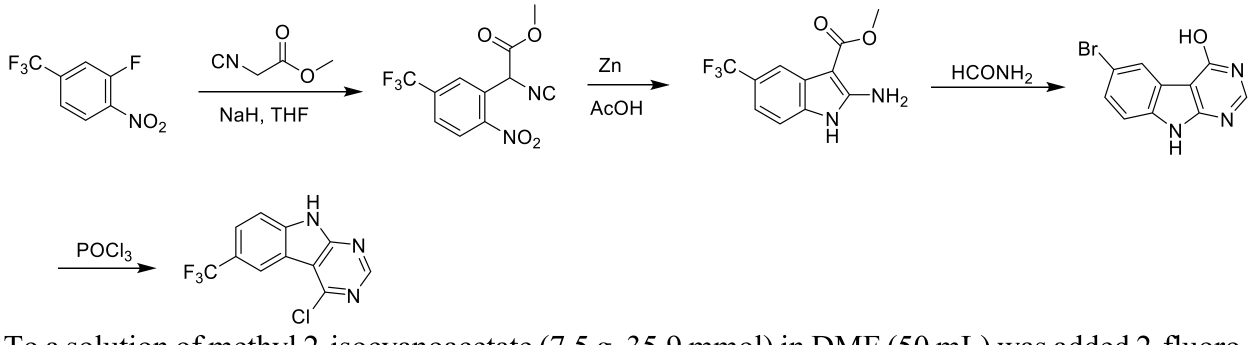


To a solution of methyl 2-isocyanoacetate (7.5 g, 35.9 mmol) in DMF (50 mL) was added 2-fluoro-1-nitro-4-(trifluoromethyl)benzene (4.3 g, 43.1 mmol) and potassium carbonate (14.9 g, 107.7 mmol). After stirring at 80°C for 2 h, the reaction mixture was cooled to room temperature, acidified with 2N HCl (ca. 75 mL), and extracted with ethyl acetate (300 mL *3). The combined organics layers were washed with brine (50 mL), dried over Sodium sulfate, concentrated under reduced [pressure and purified by silica gel chromatography (10: 1 petroleum ether/ethyl acetate) to afford 6.3 g (61%) of methyl 2-isocyano-2-(2-nitro-5-(trifluoromethyl)phenyl)acetate as yellow solid. LCMS (ESI): m/z=306.2 $(M+18)^+$

To a solution of of methyl 2-isocyano-2-(2-nitro-5-(trifluoromethyl)phenyl)acetate (6.3 g, 21.9 mmol) in glacial acetic acid (100 mL), was added zinc dust (14.3 g, 218.8 mmol) in two portions. The mixture was stirred at 60°C for 2 h, cooled to room temperature and filtered through a Celite pad. The residue was washed with THF and the filtrate was concentrated under reduced pressure. The crude product was purified by silica gel column chromatography (10:1 dichloromethane/methanol) to afford 3.1 g (55%) of methyl 2-amino-5-(trifluoromethyl)-1H-indole-3-carboxylate as a yellow solid. LCMS (ESI): m/z=259.0 $(M+H)^+$

A solution of methyl 2-amino-5-(trifluoromethyl)-1H-indole-3-carboxylate (3 g, 11.63 mmol) in formamide (30 mL) was stirred at 220°C for 2 hours. The reaction mixture was then cooled to room temperature and poured into 100 mL of water. The resulting mixture was allowed to stand for 15 min before the solids were collected by filtration, washed with water, and dried to afford 1.9 g (65%) of 6-(trifluoromethyl)-9H-pyrimido[4,5-b]indol-4-ol as a brown solid. LCMS (ESI): m/z=254.2 $(M+H)^+$

A mixture of 6-(trifluoromethyl)-9H-pyrimido[4,5-b]indol-4-ol (1.9 g, 7.5 mmol) in POCl3 (12 mL) was refluxed for two hours. The reaction mixture was then cooled to room temperature, concentrated and poured into 20 mL of water. The resulting solid was filtered, residue was triturated with ethyl acetate and filtered again to afford 0.9 g (45%) of 4-chloro-6-(trifluoromethyl)-9H-pyrimido[4,5-b]indole as a brown solid. 1H NMR (400 MHz, MeOD-*d*4) δ 8.75 (s, 1H), 8.57 (s, 1H), 7.87 (d, 1H, J = 8.6 Hz), 7.78 (d, 1H, J = 8.6 Hz). LCMS (ESI): m/z=272.1 $(M+H)^+$

**8-bromo-4-chloro-9H-pyrimido[4,5-b]indole**

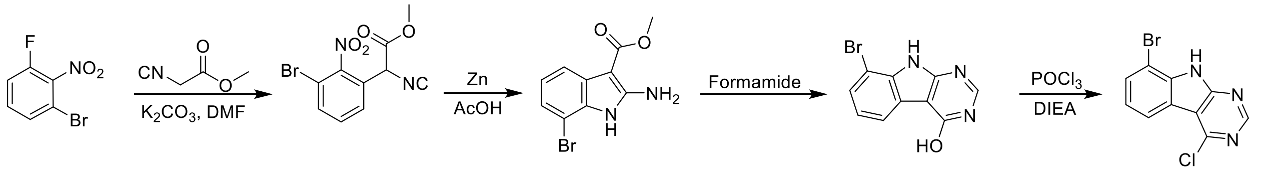


To a solution of 1-bromo-3-fluoro-2-nitrobenzene (13 g, 59.36 mmol) in dry DMF (100 mL) was added methyl 2-isocyanoacetate (11.75 g, 118.72 mmol) and $K_2CO_3$ (24.57 g, 178.08 mmol). The solution was stirred at 90°C for 16 hours. The mixture was adjusted to be weakly acidic with 2N HCl and extracted with ethyl acetate (200 mL *3). The combined organic layers were washed with brine (200 mL), dried over $Na_2SO_4$, filtered and concentrated under reduced pressure. The residue was purified by silica gel chromatography (0-50% ethyl acetate/hexanes) to afford methyl 2-(3-bromo-2-nitrophenyl)-2-isocyanoacetate as yellow solid (13 g, Yield: 73.48%). LCMS (ESI): m/z= 297.1 $(M-H)^-$

A mixture of methyl 2-(3-bromo-2-nitrophenyl)-2-isocyanoacetate (13 g, 43.62 mmol) and acetic acid (150 mL) was heated to 40 °C. Zinc (22.82 g, 348.99 mmol) was then added in portions at a rate such that the reaction temperature did not rise above 60 °C. After the addition was complete, the reaction mixture was stirred at 60°C for 4 h. The reaction mixture was cooled to room temperature and filtered through a celite and the filtrate was concentrated under reduced pressure. The crude product was purified by silica gel column chromatography (10:1 dichloromethane/methanol) to afford methyl 2-amino-7-bromo-1H-indole-3-carboxylate as a purple solid (7.5 g, Yield: 63.91%). $^1$H NMR (500 MHz, DMSO-*d*6) δ 10.77 (s, 1H), 7.55 (d, J = 7.7 Hz, 1H), 7.08 (d, J = 7.8 Hz, 1H), 6.91 (t, J = 7.8 Hz, 1H), 6.48 (s, 2H), 3.76 (s, 3H). LCMS (ESI): m/z=269.1 $(M+H)^+$.

A solution of methyl 2-amino-7-bromo-1H-indole-3-carboxylate (7.5 g, 27.88 mmol) in formamide (40 mL) was stirred at 200°C for 2 hours. The reaction mixture was then cooled to room temperature and poured into 300 mL of water. The resulting mixture was allowed to stand for 15 min, the solids were collected by filtration, washed with water, and dried to obtain 8-bromo-9H-pyrimido[4,5-b]indol-4-ol as a brown solid (6.5 g, Yield: 88.64%). LCMS (ESI): m/z=264.1 $(M+H)^+$

To a solution of 8-bromo-9H-pyrimido[4,5-b]indol-4-ol (6.5 g, 24.71 mmol) in $POCl_3$ (50 mL) was added DIPEA (2 mL). The mixture was heated to reflux for 2 hours. After cooling to room temperature, the mixture concentrated was under reduced pressure, diluted with CAN and adjusted to pH 7.0 with ammonium hydroxide slowly. The resulting solids were filtered, washed with water and dried to obtain 8-bromo-4-chloro-9H-pyrimido[4,5-b]indole (5.6 g, Yield: 80.63%) as a brown solid. $^1H$ NMR (400 MHz, DMSO-*d6*) δ 13.11 (s, 1H), 8.87 (s, 1H), 8.30 (d, J = 7.7 Hz, 1H), 7.87 (d, J = 7.8 Hz, 1H), 7.38 (t, J = 7.9 Hz, 1H). LCMS (ESI): m/z=282.0 $(M+H)^+$.

**4-Chloro-8-cyclopropyl-7-fluoro-9H-pyrimido[4,5-b]indole**

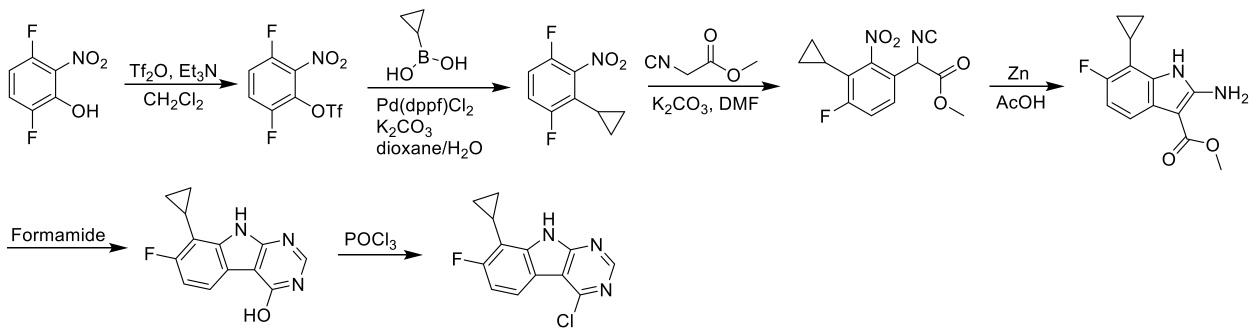


To a solution 3,6-difluoro-2-nitrophenol (15.0 g, 85.7mmol) in dichloromethane (150 mL) was added triethylamine (26g, 257.1mmol) and trifluoromethanesulfonic anhydride (128.6g, 48 mmol). After stirring at room temperature for 1h, the mixture was poured into 500mL of water and extracted with dichloromethane (500 mL *3). The combined organic layers were washed with brine (200 mL), dried over $Na_2SO_4$, filtered and concentrated under reduced pressure. The residue was purified by silica gel column chromatography (0-20% ethyl acetate/hexanes) to afford 3,6-difluoro-2-nitrophenyl trifluoromethanesulfonate as white solid (23.00 g, Yield: 87.4%).$^1H$ NMR (500 MHz, $d_6$-DMSO) δ 8.18 (td, $J$ = 9.7, 4.6 Hz, 1H), 7.97 (td, $J$ = 9.8, 4.1 Hz, 1H). LCMS (ESI): m/z= 306 $(M-H)^-$ .

To a solution of 3,6-difluoro-2-nitrophenyl trifluoromethanesulfonate (23g, 75mmol) in dioxane (250mL) and $H_2O$ (25ml) was added cyclopropylboronic acid (30.7g, 375mmol), Pd(dppf)$Cl_2$(3.45g) and potassium carbonate (31g, 225mmol). The solution was stirred at 100°C for 6 hours. The reaction mixture was cooled to room temperature and filtered through a celite pad. The filtrate was concentrated and the crude product was purified by silica gel column chromatography (20:1, petroleum ether/ethyl acetate) to afford 2-cyclopropyl-1,4-difluoro-3-nitrobenzene as a yellow oil (12 g, Yield: 80.5%).$^1H$ NMR (500 MHz, $d_6$-DMSO) δ 7.62 – 7.50 (m, 2H), 1.88 (tt, $J$ = 8.6, 5.6 Hz, 1H), 1.01 – 0.91 (m, 2H), 0.73 – 0.63 (m, 2H).

To a solution of 2-cyclopropyl-1,4-difluoro-3-nitrobenzene (12g, 60.3mmol) in dry DMF (100 mL) was added methyl 2-isocyanoacetate (7.76 g, 78.4mmol) and potassium carbonate (24.9g, 180.9mmol). The solution was stirred at 90°C for 16 hours. The mixture was adjusted to be weakly acidic with 2N HCl, extracted with ethyl acetate (300 mL *3), the organic layers were washed with brine (200 mL), dried over $Na_2SO_4$, filtered and concentrated under reduced pressure. The residue was purified by silica gel chromatography (0-50% ethyl acetate/hexanes) to afford methyl 2-(3-cyclopropyl-4-fluoro-2-nitrophenyl)-2-isocyanoacetate as a white solid (15 g, Yield: 89.5%). $^1$H NMR (400 MHz, *d$_6$*-DMSO) δ 7.61 (dd, *J* = 9.9, 7.4 Hz, 2H), 6.02 (s, 1H), 3.76 (s, 3H), 1.90 (tt, *J* = 8.6, 5.6 Hz, 1H), 1.03 – 0.84 (m, 2H), 0.69 – 0.46 (m, 2H). LCMS (ESI): m/z= 296 $(M+NH_4)^+$.

A mixture of methyl 2-(3-cyclopropyl-4-fluoro-2-nitrophenyl)-2-isocyanoacetate (15g, 54mmol) and acetic acid (150mL) was heated to 40 °C. Zinc (28g, 432mmol) was then added in portions at a rate such that the reaction temperature did not rise above 60 °C. After the addition was complete, the reaction mixture was stirred at 60°C for 2 h. The reaction mixture was cooled to room temperature and filtered through a celite pad. The filtrate was concentrated and the crude product was purified by silica gel chromatography (10:1 dichloromethane/methanol) to give methyl 2-amino-7-cyclopropyl-6-fluoro-1H-indole-3-carboxylate as a white solid (7g, Yield: 52.3%). $^1$H NMR (400 MHz, DMSO) δ 10.54 (s, 1H), 7.33 (dd, *J* = 8.4, 5.0 Hz, 1H), 6.71 (dd, *J* = 11.7, 8.5 Hz, 1H), 6.39 (s, 2H), 3.73 (s, 3H), 1.86 (tt, *J* = 8.5, 5.4 Hz, 1H), 1.03 – 0.91 (m, 2H), 0.78 – 0.62 (m, 2H). LCMS (ESI): m/z=249$(M+H)^+$.

A solution of methyl 2-amino-7-cyclopropyl-6-fluoro-1H-indole-3-carboxylate (7g, 28.2mmol) in formamide (40 mL) was stirred at 200°C for 2 hours. Upon completion of reaction, the reaction mixture was cooled to room temperature and poured into 300 mL of water. The resulting mixture was allowed to stand for 15 min before the solids were collected by filtration, washed with water, and dried in vacuo to afford 8-cyclopropyl-7-fluoro-9H-pyrimido[4,5-b]indol-4-ol as a brown solid (5g, Yield: 72.9%). $^1$H NMR (500 MHz, DMSO) δ 12.15 (s, 2H), 8.17 (s, 1H), 7.81 (dd, *J* = 8.4, 4.9 Hz, 1H), 7.01 (dd, *J* = 11.8, 8.6 Hz, 1H), 2.12 – 2.03 (m, 1H), 1.06 (t, *J* = 13.9 Hz, 2H), 0.91 (dd, *J* = 19.7, 5.2 Hz, 2H). LCMS (ESI): m/z=244 $(M+H)^+$.

To a solution of 8-cyclopropyl-7-fluoro-9H-pyrimido[4,5-b]indol-4-ol (5g, 20.6mmol) in $POCl_3$ (30 mL) was added DIPEA (0.5 ml). The mixture was heated to reflux for 2 hours. After cooling to room temperature, the mixture was concentrated under reduced pressure and diluted with acetonitrile, adjusted the PH to 7.0 with ammonium hydroxide slowly. The resulting solid was filtered with vacuum filter and washed with water to obtain 4-chloro-8-cyclopropyl-7-fluoro-9H-pyrimido[4,5-b]indole (3.5g, Yield: 65.1%) as a brown solid. $^1$H NMR (500 MHz, DMSO) δ 12.67 (s, 1H), 8.80 (s, 1H), 8.09 (dd, *J* = 8.6, 4.9 Hz, 1H), 7.19 (dd, *J* = 11.6, 8.7 Hz, 1H), 2.09 (tt, *J* = 8.6, 5.4 Hz, 1H), 1.21 – 1.02 (m, 2H), 0.94 – 0.79 (m, 2H). LCMS (ESI): m/z=262.2 $(M+H)^+$.

**4-Chloro-8-cyclopropyl-6-fluoro-9H-pyrimido[4,5-b]indole**

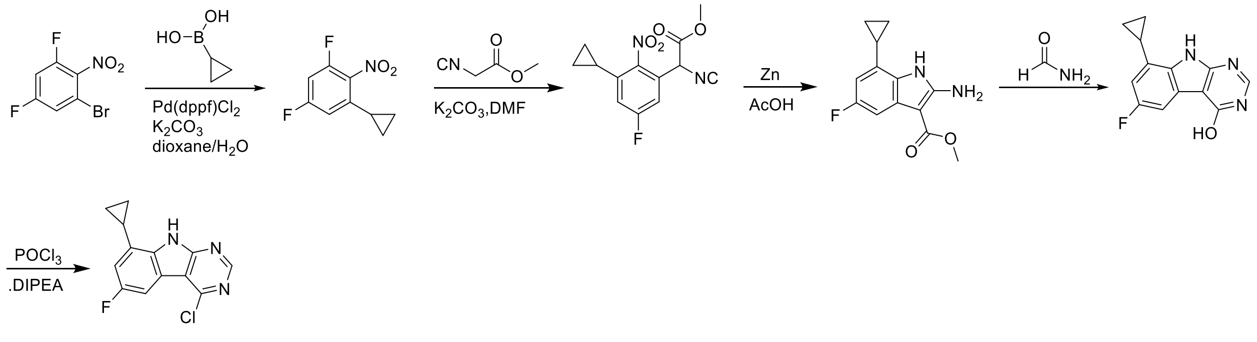

To a solution of 1-bromo-3,5-difluoro-2-nitrobenzene (25 g, 105.0 mmol) in 10:1 Dioxane/ H2O (100.0 mL) was added cyclopropylboronic acid (54.14 g, 630.2 mmol), potassium carbonate (28.99 g, 210.0 mmol) and Pd(dppf)$Cl_2$ (2.04 g, 2.1 mmol). After stirring at 100°C for 16 h, the reaction mixture was cooled to room temperature and poured into water (150 mL) and extracted with ethyl acetate (300 mL *3). The combined organic layers were washed with brine (100 mL), dried over $Na_2SO_4$ and concentrated. The residue was purified by silica gel column chromatography (petroleum ether/ ethyl acetate 10: 1) to obtain 1-cyclopropyl-3,5-difluoro-2-nitrobenzene as a yellow oil (20 g, Yield: 95.67%). 1H NMR (500 MHz, DMSO-d6) δ 7.61 – 7.45 (m, 1H), 7.09 – 6.92 (m, 1H), 2.04 – 1.86 (m, 1H), 1.10 – 1.04 (m, 2H), 0.91 – 0.86 (m, 2H).

To a solution of 1-cyclopropyl-3,5-difluoro-2-nitrobenzene (20 g, 100.5 mmol) in DMF (100 mL) was added methyl 2-isocyanoacetate (9.95 g, 100.5 mmol) and potassium carbonate (41.61 g, 301.5 mmol), the solution was stirred at 80°C for 3 hours. The mixture was cooled to room temperature, acidified with 2N HCl (ca. 130 mL) and extracted with ethyl acetate (300 mL *3). The combined organic extracts were washed with brine (100 mL), dried over $Na_2SO_4$ and concentrated. The residue was purified by silica gel column chromatography (petroleum ether/ ethyl acetate 2: 1) to obtain methyl 2-(3-cyclopropyl-5-fluoro-2-nitrophenyl)-2-isocyanoacetate as a white solid (10 g, Yield: 35.79%). 1H NMR (400 MHz, DMSO-d6) δ 7.42 (dd, J = 8.6, 2.6 Hz, 1H), 7.26 (dd, J = 9.6, 2.5 Hz, 1H), 6.03 (s, 1H), 3.77 (s, 3H), 2.07 – 1.81 (m, 1H), 1.10 – 0.95 (m, 2H), 0.93 – 0.79 (m, 2H). LCMS (ESI): m/z= 296.2 $(M+18)^+$

To a solution of methyl 2-(3-cyclopropyl-5-fluoro-2-nitrophenyl)-2-isocyanoacetate (10 g, 35.9 mmol) in glacial acetic acid (80 ml), was added zinc (18.81 g, 287.7 mmol) in two portions and the reaction mixture was stirred at 60°C for 3 h. The reaction mixture was then cooled to room temperature and filtered. The residue was washed with THF and the filtrate was concentrated. The residue was purified by silica gel column chromatography (dichloromethane/ methanol 10:1) to obtain methyl 2-amino-7-cyclopropyl-5-fluoro-1H-indole-3-carboxylate as an off-white solid (6 g, Yield: 67.25%). 1H NMR (500 MHz, DMSO-d6) δ 10.81 (s, 1H), 7.05 (dd, J = 9.8, 1.8 Hz, 1H), 6.48 (s, 2H), 6.27 (dd, J = 11.0, 2.5 Hz, 1H), 3.75 (s, 3H), 2.20 – 2.03 (m, 1H), 1.00 – 0.93 (m, 2H), 0.74 – 0.68 (m, 2H). LCMS (ESI): m/z=249.1 $(M+H)^+$

A solution of methyl 2-amino-7-cyclopropyl-5-fluoro-1H-indole-3-carboxylate (6 g, 24.2 mmol) in formamide (40 mL) was stirred at 200°C for 4 hours. The reaction mixture was then cooled to room temperature and poured into 80 mL of water. The resulting mixture was allowed to stand for 15 min before the solids were collected by filtration, washed with water, and dried in vacuo to obtain 8-cyclopropyl-6-fluoro-9H-pyrimido[4,5-b]indol-4-ol as a brown solid (5.8 g, Yield: 98.65%). 1H NMR (500 MHz, DMSO-d6) δ 12.42 (s, 1H), 12.23 (s, 1H), 8.16 (s, 1H), 7.43 (dd, J = 8.8, 2.4 Hz, 1H), 6.70 (dd, J = 11.0, 2.4 Hz, 1H), 2.45 – 2.38 (m, 1H), 1.11 – 1.01 (m, 2H), 0.87 – 0.76 (m, 2H). LCMS (ESI): m/z=244.1 $(M+H)^+$

To a solution of obtain 8-cyclopropyl-6-fluoro-9H-pyrimido[4,5-b]indol-4-ol (5.8 g, 23.8 mmol) in $POCl_3$ (40 mL) was added DIPEA (4 mL) and the reaction mixture was stirred at 90°C for 5h. After cooling to room temperature the mixture was concentrated and poured into 30 mL of water. The resulting solid was filtered to obtain the crude product. The residue was triturated with ethyl acetate and filtered to afford 4-chloro-8-cyclopropyl-6-fluoro-9H-pyrimido[4,5-b]indole as brown solid (5.2 g, Yield 83.47%).

**(R)-2-((6-Bromo-7*H*-pyrrolo[2,3-*d*]pyrimidin-4-yl)amino)-3-methylbutan-1-ol**

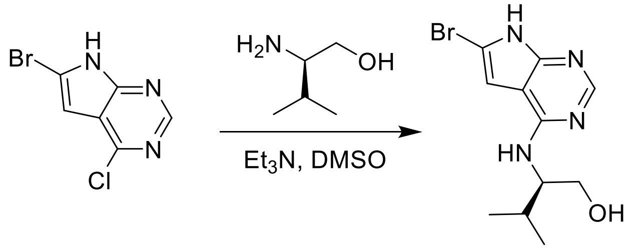

To a solution of 6-bromo-4-chloro-7*H*-pyrrolo[2,3-*d*]pyrimidine (900 mg, 3.9 mmol) in dry DMSO (10 mL) was added (*R*)-2-amino-3-methylbutan-1-ol (602 mg, 5.8 mmol) and TEA (787 mg, 7.8 mmol), the mixture was stirred at 110 °C for 16 hours. The reaction mixture was diluted with ethyl acetate (40.0 mL) and washed with water (5.0 mL), and brine (5.0 mL). The organic layer was dried over $Na_2SO_4$ and concentrated under reduced pressure. The residue was purified by prep-HPLC (0.1% $NH_4HCO_3$ in water, 10%-100% ACN) to give (R)-2-((6-bromo-7*H*-pyrrolo[2,3-*d*]pyrimidin-4-yl)amino)-3-methylbutan-1-ol as a white solid (522 mg, yield: 45%). 1H NMR (500 MHz, DMSO-*d6*) δ 12.22 (s, 1H), 8.03 (s, 1H), 7.00 (d, 1H, *J* = 8.8 Hz), 6.79 (s, 1H), 4.62 (t, 1H, *J* = 5.2 Hz), 4.13 (s, 1H), 3.52 (dd, 2H, *J* = 9.4, 4.0 Hz), 1.98 (dt, 1H, *J* = 13.6, 6.8 Hz), 0.91 (dd, 6H, *J* = 8.6, 6.9 Hz). LCMS (ESI): m/z=299.2 (M+H)+.

**(R)-2-((6-iodo-7H-pyrrolo[2,3-d]pyrimidin-4-yl)amino)-3-methylbutan-1-ol**

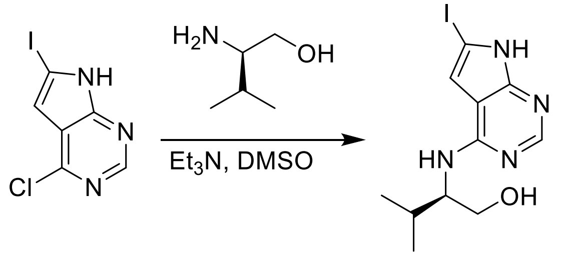

To a solution of 4-chloro-6-iodo-7*H*-pyrrolo[2,3-*d*]pyrimidine (200 mg, 715.6 µmol) in DMSO (1.0 mL) was added (*R*)-2-amino-3-methylbutan-1-ol (88.6 mg, 858.8 µmol) and TEA (362.1 mg, 3.58 mmol). The mixture was stirred at 100 °C for 16 hours, filtered and the residue was purified by prep-HPLC (0%-40% water/acetonitrile) to obtain (R)-2-((6-iodo-7H-pyrrolo[2,3-d]pyrimidin-4-yl)amino)-3-methylbutan-1-ol as a white solid (70.9 mg, yield: 25%). 1H NMR (400 MHz, MeOD) (mixture of rotamers was observed) δ 8.28 (brs, 1H), 8.01 (s, 1H), 6.90 (s, 1H), 4.13-4.09 (m, 1H), 3.81-3.71 (m, 2H), 2.12-2.03 (m, 1H), 1.05-1.00 (m, 6H). LCMS (ESI): m/z=347 $(M+H)^+$

**4-(4-Chloro-7H-pyrrolo[2,3-d]pyrimidin-6-yl)-1-methyl-1H-pyrazole-5-carbonitrile**

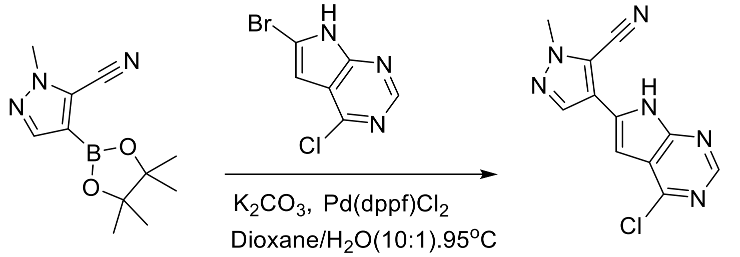


To a solution of 1-methyl-4-(4,4,5,5-tetramethyl-1,3,2-dioxaborolan-2-yl)-1H-pyrazole-5-carbonitrile (241 mg, 1.03 mmol) in 1,4-dioxane (2 mL) and water (0.2 mL) was added 6-bromo-4-chloro-7H-pyrrolo[2,3-d]pyrimidine (200 mg, 0.86 mmol), $K_2CO_3$ (356.04 mg, 2.58 mmol) and Pd(dppf)$Cl_2$ (66 mg, 10%w/w). The mixture was stirred at 95 °C for 2 hours under nitrogen gas. The reaction mixture was diluted with water (40 mL) and extracted with DCM (3 x 80 mL). The organic layers were dried over $Na_2SO_4$ and concentrated under reduced pressure. The residue was purified by column chromatography on silica gel (petroleum ether: ethyl acetate = 2:1) to give 4-(4-chloro-7H-pyrrolo[2,3-d]pyrimidin-6-yl)-1-methyl-1H-pyrazole-5-carbonitrile as a yellow solid (160 mg, yield: 72.30%). LCMS (ESI): m/z= 259.0 $(M+18)^+$

**4-(4-Chloro-7-((2-(trimethylsilyl)ethoxy)methyl)-7H-pyrrolo[2,3-d]pyrimidin-6-yl)-1-methyl-1H-pyrazole-5-carbonitrile**

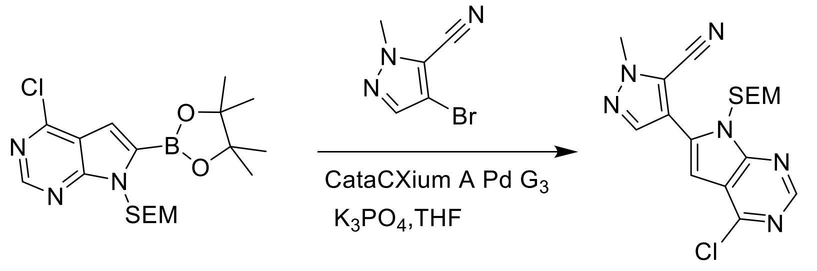


To a solution of 4-bromo-1-methyl-1H-pyrazole-5-carbonitrile (1.06 g, 5.70 mmol) in THF (20 ml) was added 4-chloro-6-(4,4,5,5-tetramethyl-1,3,2-dioxaborolan-2-yl)-7-((2-(trimethylsilyl)ethoxy)methyl)-7H-pyrrolo[2,3-d]pyrimidine (3.50 g, 8.55 mmol), $K_3PO_4$ (2.42 g, 11.40 mmol) and CataCXium A Pd G3 (249 mg, 0.34 mmol). After stirring the mixture at 70°C for 2 hours under nitrogen gas, the reaction mixture was suspended in water (60 mL) and extracted with ethyl acetate (3 x 100 mL). The organic layers were dried over $Na_2SO_4$ and concentrated under reduced pressure. The residue was purified by column chromatography on silica gel (petroleum ether/ ethyl acetate, 2:1) to obtain 4-(4-chloro-7-((2-(trimethylsilyl)ethoxy)methyl)-7H-pyrrolo[2,3-d]pyrimidin-6-yl)-1-methyl-1H-pyrazole-5-carbonitrile as a yellow solid (600 mg, Yield: 27.1%). LCMS (ESI): m/z= 389.0 $(M+H)^+$

**1-Amino-5,5-dimethylpyrrolidin-2-one hydrochloride**

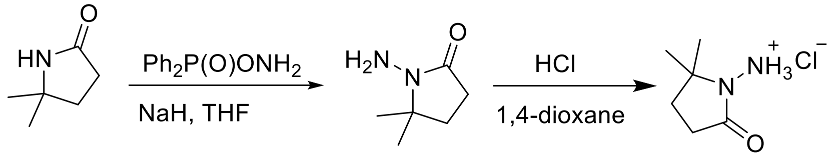


To a cooled (0°C) solution of 5,5-dimethylpyrrolidin-2-one (3 g, 26.54 mmol) in THF (60 mL) was added sodium hydride (2.13 g, 53.09 mmol), followed by addition of O-diphenylphosphinyl hydroxylamine (12.4 g, 53.09 mmol) after 30 min. After stirring the resultant white suspension at 0°C for 2 h, the reaction mixture was filtered through a Celite pad, the filtrate was concentrated and purified by silica gel chromatography (10:1 dichloromethane/methanol) to afford 3 g (75%) of 1-amino-5,5-dimethylpyrrolidin-2-one as yellow oil. LCMS (ESI): m/z= 129.1 $(M+18)^+$;

A solution of 1-amino-5,5-dimethylpyrrolidin-2-one (1.5 g, crude) in 4 N HCl in dioxane (15 mL) was stirred at room temperature for 4h. The mixture was concentrated under reduced pressure, residue was triturated with diethyl ether and filtered to afford 1 g (53%) of 1-amino-5,5-dimethylpyrrolidin-2-one hydrochloride salt as a white solid. 1H NMR (500 MHz, DMSO) δ 9.48 (s, 3H), 2.39 (t, 2H, J = 7.8 Hz), 1.90 (t, 2H, J = 7.8 Hz), 1.30 (s, 6H). LCMS (ESI): m/z= 129.1 $(M+18)^+$

**1-Amino-6,6-dimethylpiperidin-2-one**

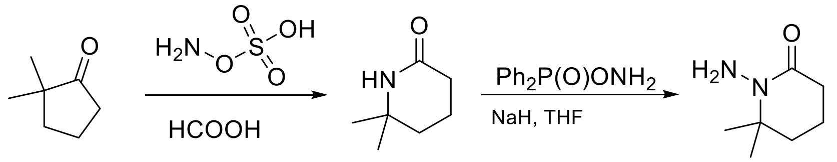


To a cooled (0 °C)solution of 2,2-dimethylcyclopentan-1-one (25 g, 222.87 mmol) in formic acid (140 mL), was added hydroxyamine-*O*-sulfonic acid (37.80 g, 334.30 mmol). After stiring at 100°C for 4 hours, the reaction mixture was concentrated and the light orange residue was taken in dichloromethane (250 mL) and stirred until completely dissolved. Washed with 20% NaOH solution and the aqueous phase was extracted with dichloromethane (300 mL x 3) and the combined organic layers concentrated. The residue was purified by column chromatography on silica gel (ethyl acetate /methanol, 9:1) to obtain 13.5 g of the product as white solid. This was further purified by reverse phase chromatography (0.05% TFA in water, 10%-100% ACN) to obtain 6,6-dimethylpiperidin-2-one (9.1 g, Yield: 32.15%) as an off-white solid. $^1$H NMR (500 MHz, DMSO-d6) δ 7.32 (d, J = 43.8 Hz, 1H), 2.05 (t, J = 6.6 Hz, 2H), 1.70 – 1.65 (m, 2H), 1.54 – 1.50 (m, 2H), 1.14 (s, 6H). LCMS (ESI): m/z=128.4 (M+H)+

A solution of 6,6-dimethylpiperidin-2-one (9.10 g, 71.65 mmol) in THF (300 mL) was cooled to 0°C under nitrogen gas, followed by addition of NaH (60% in oil) (8.60 g, 214.95 mmol). After 30 minutes, O-diphenylphosphinyl hydroxylamine (25.04 g, 107.47 mmol) was added. After stirring the resultant white suspension at 0°C for 5 hours, the mixture was filtered through celite pad, the filtrate was concentrated and the residue was purified by silica gel column chromatography (dichloromethane: methanol, 10: 1) to afford 1-amino-6,6-dimethylpiperidin-2-one as yellow oil (6.50 g, yield: 63.91%). $^1$H NMR (500 MHz, DMSO) δ 4.44 (s, 2H), 2.24 (t, J = 6.5 Hz, 2H), 1.73 – 1.63 (m, 4H), 1.23 (s, 6H). LCMS (ESI): m/z= 143.1 $(M+H)^+$

**3-Amino-4,4-dimethyloxazolidin-2-one**

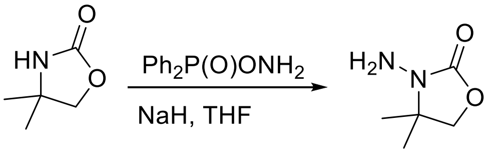


A solution of 4,4-dimethyloxazolidin-2-one (2.5 g, 21.74 mmol) in THF (100 mL) was cooled to 0°C under nitrogen gas, followed by addition of NaH (60% in oil) (2.6 g, 65.22 mmol). After 30 minutes, O-diphenylphosphinyl hydroxylamine (7.60 g, 32.61 mmol) was added. After stirring the resultant white suspension at 0°C for 4 hours, the mixture was filtered through celite pad, the filtrate was concentrated and the residue was purified by silica gel column chromatography (dichloromethane: methanol, 10: 1) to afford 3-Amino-4,4-dimethyloxazolidin-2-one as yellow oil (1.3 g, Yield: 46.00%). $^1$H NMR (500 MHz, DMSO) δ 4.22 (d, $J$ = 104.8 Hz, 2H), 3.96 (s, 2H). LCMS (ESI): m/z= 131.2 $(M+H)^+$

**1-Amino-3,8-dioxa-1-azaspiro[4.5]decan-2-one**

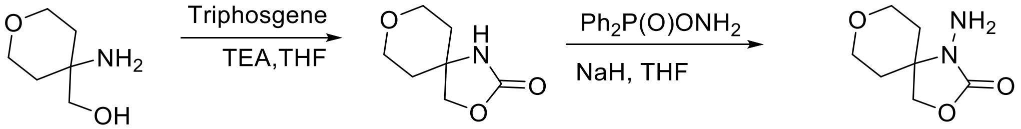


To a cooled (0°C) solution of (4-aminotetrahydro-2H-pyran-4-yl)methanol (4 g, 30.5 mmol) and triethylamine (9.2 g,91.5 mmol) in THF (50 ml) was added triphosgene (5.4 g,18.3 mmol). After stirring the mixture for 2 h, the reaction mixture was concentrated and purified by column chromatography on silica gel (dichloromethane: methanol, 10: 1) to obtain 3,8-dioxa-1-azaspiro[4.5]decan-2-one (3.5 g, Yield:73.1) as a white solid. LCMS (ESI): m/z= 158.1 $(M+H)^+$
To a cooled (0°C) solution of 3,8-dioxa-1-azaspiro[4.5]decan-2-one (1.57 g, 10.00 mmol) in THF (40 mL) under an atmosphere of nitrogen gas was added 60% sodium hydride suspension in mineral oil (1.2 g, 30.00 mmol). After stirring for 30 min, O-diphenylphosphinyl hydroxylamine (3.50 g, 15.00 mmol) was added. The resultant white suspension was stirred at 0°C for 4 hours. The reaction mixture was filtered through celite pad, the filtrate was concentrated and the residue was purified by column chromatography on silica gel (dichloromethane: methanol, 10: 1) to afford 1-amino-3,8-dioxa-1-azaspiro[4.5]decan-2-one as a white solid (700 mg, yield: 40.70%). $^1$H NMR (500 MHz, DMSO) δ 4.41 (s, 2H), 4.19 (s, 2H), 3.85 (dd, $J$ = 11.9, 5.1 Hz, 2H), 3.38 – 3.05 (m, 2H), 1.92 (td, $J$ = 13.1, 5.1 Hz, 2H), 1.37 (dd, $J$ = 12.8, 1.4 Hz, 2H). LCMS (ESI): m/z= 173.1 $(M+H)^+$

**1-Amino-3-oxa-1,8-diazaspiro[4.5]decan-2-one**

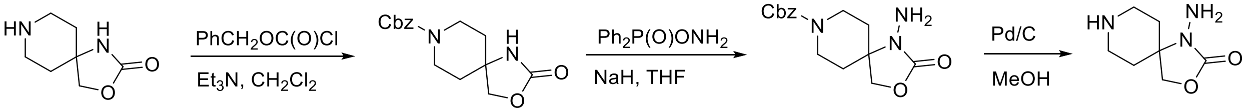


To a cooled (0°C) solution of 3-oxa-1,8-diazaspiro[4.5]decan-2-one (2 g, 12.80 mmol) and triethylamine (3.88 g, 38.40 mmol) in dry dichloromethane (50 mL) was added benzyl chloroformate (1.74 g, 10.24 mmol) under nitrogen gas. After stirring at room temperature for 16

h, the mixture was poured into $H_2O$ (100 mL) and extracted with ethyl acetate (3 x 100 mL ). The combined organic extracts were washed with brine, dried over sodium sulfate, and concentrated in vacuo. The crude product was purified by column chromatography on silica gel (dichloromethane: methanol, 10: 1) to afford benzyl 2-oxo-3-oxa-1,8-diazaspiro[4.5]decane-8-carboxylate (1.70 g, yield: 45.82%) as a white solid. LCMS (ESI): m/z= 291.3 $(M+H)^+$

To a cooled (0°C) solution of benzyl 2-oxo-3-oxa-1,8-diazaspiro[4.5]decane-8-carboxylate (1.70 g, 5.86 mmol) in dry THF (100 mL) was added 60% in oil sodium hydride (703 mg, 17.58 mmol) under nitrogen gas. After stirring for 30 min, O-diphenylphosphinyl hydroxylamine (2.05 g, 8.79 mmol) was added. The resultant white suspension was stirred at 0°C for 5 hours. The mixture was filtered through celite pad, the filtrate was concentrated and the residue was purified by column chromatography on silica gel (dichloromethane: methanol, 10: 1) to afford benzyl 1-amino-2-oxo-3-oxa-1,8-diazaspiro[4.5]decane-8-carboxylate as a yellow oil (1.60 g, yield: 89.88%). LCMS (ESI): m/z= 306.3 $(M+H)^+$

A solution of benzyl 1-amino-2-oxo-3-oxa-1,8-diazaspiro[4.5]decane-8-carboxylate (1.60 g, 3.27 mmol) in methanol (30 mL) was added 10 wt% Pd/C (300 mg). The mixture was stirred at 25°C for 3 hours under hydrogen (1 atm, balloon), filtered and the filtrate was concentrated under vacuum to afford 1-amino-3-oxa-1,8-diazaspiro[4.5]decan-2-one as a white solid (643 mg, yield: 71.63%). $^1H$ NMR (400 MHz, MeOD-*d*4) δ 4.22 (s, 2H), 3.12 – 3.00 (m, 2H), 2.59 (td, J = 13.3, 2.6 Hz, 2H), 2.09 – 1.92 (m, 2H), 1.62 – 1.42 (m, 2H). LCMS (ESI): m/z= 172.2 $(M+H)^+$

**1-Amino-8-(methylsulfonyl)-3-oxa-1,8-diazaspiro[4.5]decan-2-one**

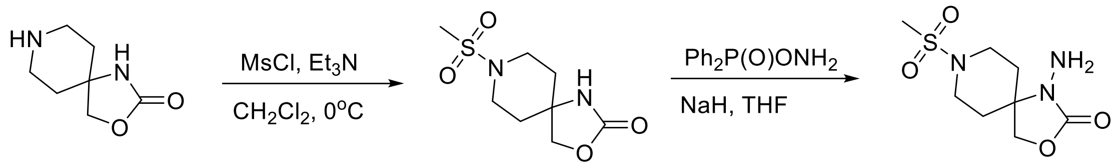


To a cooled (0°C) solutiion of 3-oxa-1,8-diazaspiro[4.5]decan-2-one (2 g, 12.80 mmol, 1 equiv) and triethylamine (1.94 g, 19.20 mmol, 1.5 equiv) in dichloromethane (30 mL) was added methanesulfonyl chloride (1.48 g, 10.24 mmol, 0.8 equiv) under nitrogen gas. After the mixture was stirred at 0°C for 1 hour, the mixture was poured into water (80 mL) and extracted with ethyl acetate (100 mL *3). The combined organic extracts were washed with brine, dried over sodium sulfate, and concentrated. The crude product was purified by silica gel column chromatography (dichloromethane: methanol, 10: 1) to afford 8-(methylsulfonyl)-3-oxa-1,8-diazaspiro[4.5]decan-2-one (1.90 g, Yield: 63.54%) as a white solid. LCMS (ESI): m/z= 234.9 $(M+H)^+$

To a cooled (0°C) solution of 8-(methylsulfonyl)-3-oxa-1,8-diazaspiro[4.5]decan-2-one (1.90 g, 8.11 mmol) in THF (100 mL) was added 60% in oil sodium hydride (973 mg, 24.33 mmol) under nitrogen gas. After stirring for 30 min, O-diphenylphosphinyl hydroxylamine (2.84 g, 12.16 mmol) was added. The resultant white suspension was stirred at 0°C for 5 hours, filtered through celite, the filtrate concentrated and the residue was purified by silica gel column chromatography (dichloromethane: methanol, 10: 1) to afford 1-amino-8-(methylsulfonyl)-3-oxa-1,8-diazaspiro[4.5]decan-2-one as a yellow solid (633 mg, Yield: 31.65%). $^1H$ NMR (500 MHz, DMSO-d6) δ 4.44 (s, 2H), 4.15 (s, 2H), 3.57 (dd, J = 10.1, 2.3 Hz, 2H), 2.87 (s, 3H), 2.78 (td, J = 12.8, 2.1 Hz, 2H), 1.93 (td, J = 13.1, 4.7 Hz, 2H), 1.57 (d, J = 13.3 Hz, 2H). LCMS (ESI): m/z= 250.1 $(M+H)^+$

**1-((8-Amino-9H-pyrimido[4,5-b]indol-4-yl)amino)-5,5-dimethylpyrrolidin-2-one**

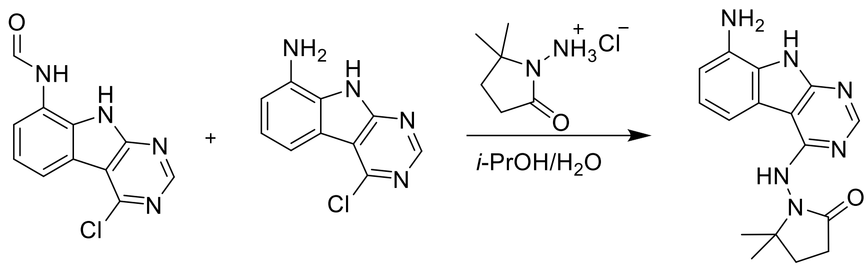


A solution of a mixture of N-(4-chloro-9H-pyrimido[4,5-b]indol-8-yl)formamide and 4-chloro-9H-pyrimido[4,5-b]indol-8-amine (15 g, 60.97 mmol) and 1-amino-5,5-dimethylpyrrolidin-2-one hydrochloride (11.7 g, 91.46 mmol) in isopropanol (150 mL) and water(15 mL) was stirred at 100°C for 16 hours. The reaction mixture was diluted with water (30 mL) and extracted with ethyl acetate (3 x 60 mL). The combined organic layers were dried over Sodium sulfate, concentrated under reduced pressure and the residue was purified by reverse phase chromatography (water/acetonitrile /0.1% ammonium bicarbonate) to obtain 77.3 mg (10.2%) of 1-((8-amino-9H-pyrimido[4,5-b]indol-4-yl)amino)-5,5-dimethylpyrrolidin-2-one as a white solid. 1H NMR (500 MHz, MeOD) δ 8.32 (s, 1H), 7.56 (d, 1H, J = 7.8 Hz), 7.12 (s, 1H), 6.85 (d, 1H, J = 7.6 Hz), 2.56 (t, 2H, J = 7.9 Hz), 2.17 (s, 2H), 1.37 (s, 6H). LCMS (ESI): m/z 311.3 $(M+H)^+$

**Synthesis and Characterization Data for Final Analogs**

**AVI-1499**

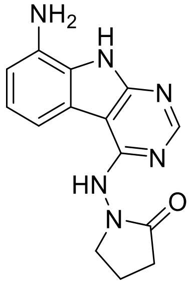


A mixture of 4-chloro-9H-pyrimido[4,5-b]indol-8-amine (28 mg, 0.13 mmol) and 1-aminopyrrolidin-2-one hydrochloride (35 mg, 0.26 mmol) in isopropanol/water (10: 1, 1.1 mL) were heated to 100 °C for 18 h. The reaction mixture was filtered, the residue was washed with ethyl acetate and dried to obtain 28 mg (77%) of 1-((8-amino-9H-pyrimido[4,5-b]indol-4-yl)amino)pyrrolidin-2-one (**AVI-1499**) as brown solid. $^1$H NMR (DMSO-d$_6$, 400 MHz) δ 12.99 (br s, 1H), 8.62 (s, 1H), 7.92 (br d, 1H, *J*=7.5 Hz), 7.27 (t, 1H, *J*=7.9 Hz), 7.05 (br d, 1H, *J*=7.5 Hz), 3.70 (br t, 2H, *J*=6.9 Hz), 2.44-2.53 (m, 2H), 2.20 (br t, 2H, *J*=7.4 Hz). LCMS (ESI): m/z= 283 $(M+H)^+$

**AVI-1501**

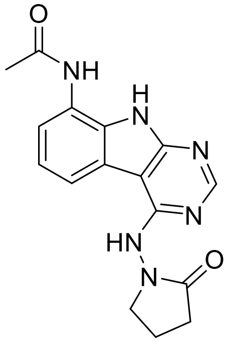


To a solution of 1-((8-amino-9H-pyrimido[4,5-b]indol-4-yl)amino)pyrrolidin-2-one (15 mg, 0.053 mmol) and triethylamine (0.015 mL, 0.11 mmol) in THF (1 mL), was added acetyl chloride (0.004 mL, 0.056 mmol). After stirring at 65 °C for 3 h, the reaction mixture was purified by reverse phase chromatography (water/acetonitrile/0.1% formic acid) to obtain 9 mg (50%) of N-(4-((2-oxopyrrolidin-1-yl)amino)-9H-pyrimido[4,5-b]indol-8-yl)acetamide formic acid salt (**AVI-1501**)as a white solid. $^1$H NMR (METHANOL-d$_4$, 400 MHz) δ 8.39 (s, 1H), 7.90 (d, 1H, *J*=7.8 Hz), 7.45 (d, 1H, *J*=7.8 Hz), 7.19-7.21 (m, 1H), 3.81-3.85 (m, 2H), 2.59-2.63 (m, 2H), 2.27-2.31 (m, 5H). LCMS (ESI): m/z= 325 $(M+H)^+$

**AVI-1500**

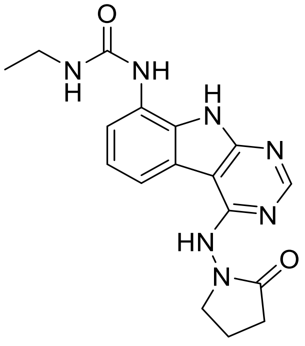

To a solution of 4-chloro-9H-pyrimido[4,5-b]indol-8-amine (50 mg, 0.23 mmol) and triethylamine (0.064 mL, 0.46 mmol) in THF (2 mL), was added ethyl isocyanate (0.018 mL, 0.23 mmol). After stirring at 65 °C for 18 h, the reaction mixture was filtered. The residue was washed with ethyl acetate and dried to obtain 50 mg of 1-(4-chloro-9H-pyrimido[4,5-b]indol-8-yl)-3-ethylurea as a white solid that was used without further purification. $^{1}H$ NMR (DMSO-$d_6$, 400 MHz) δ 12.39 (br s, 1H), 8.80 (s, 1H), 8.43 (s, 1H), 7.96 (d, 1H, *J*=7.6 Hz), 7.72 (d, 1H, *J*=7.8 Hz), 7.35 (t, 1H, *J*=7.9 Hz), 6.38 (s, 1H), 3.18-3.21 (m, 2H), 1.12 (t, 3H, *J*=7.2 Hz). LCMS (ESI): m/z= 290, 292 $(M+H)^+$

A mixture of 1-(4-chloro-9H-pyrimido[4,5-b]indol-8-yl)-3-ethylurea (26 mg, 0.09 mmol) and 1-aminopyrrolidin-2-one hydrochloride (25 mg, 0.18 mmol) in isopropanol/water (10: 1, 1.1 mL) were heated to 100 °C for 18 h. The reaction mixture was purified by reverse phase chromatography (water/acetonitrile/0.1% formic acid) to obtain 8 mg (20%) of 1-ethyl-3-(4-((2-oxopyrrolidin-1-yl)amino)-9H-pyrimido[4,5-b]indol-8-yl)urea formic acid salt (**AVI-1500**) as a white solid. $^{1}H$ NMR (DMSO-$d_6$, 400 MHz) δ 11.75 (br s, 1H), 9.31 (s, 1H), 8.53 (s, 1H), 8.41 (s, 1H), 7.97 (d, 1H, *J*=7.8 Hz), 7.63 (d, 1H, *J*=7.8 Hz), 7.20 (t, 1H, *J*=7.9 Hz), 6.37 (br s, 1H), 3.70 (t, 2H, *J*=7.1 Hz), 3.17-3.20 (m, 2H), 2.39-2.41 (m, 2H), 2.12-2.16 (m, 2H), 1.09-1.13 (m, 3H). $^{13}C$ NMR (DMSO-$d_6$, 100 MHz) δ 173.5, 156.2, 155.9, 155.5, 155.0, 128.5, 125.5, 121.3, 120.3, 117.1, 116.7, 96.4, 48.4, 34.8, 28.9, 16.7, 15.9. LCMS (ESI): m/z= 354 $(M+H)^+$

**AVI-3367**

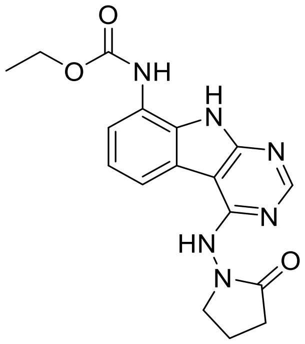

To a solution of 1-((8-amino-9H-pyrimido[4,5-b]indol-4-yl)amino)pyrrolidin-2-one (15 mg, 0.053 mmol) and triethylamine (0.015 mL, 0.11 mmol) in THF (1 mL), was added ethyl chloroformate (0.005 mL, 0.056 mmol). After stirring at 65 °C for 18 h, the reaction mixture was purified by reverse phase chromatography (water/acetonitrile/0.1% formic acid) to obtain 2.7 mg (13%) of ethyl (4-((2-oxopyrrolidin-1-yl)amino)-9H-pyrimido[4,5-b]indol-8-yl)carbamate formic acid salt (**AVI-3367**) as tan solid. $^{1}H$ NMR (METHANOL-$d_4$, 400 MHz) δ 8.42 (s, 1H), 7.94 (d, 1H, *J*=7.8 Hz), 7.59 (br s, 1H), 7.28 (t, 1H, *J*=7.9 Hz), 4.1-4.26-4.30 (m, 2H), 3.84 (t,

2H, *J*=7.1 Hz), 2.60 (t, 2H, *J*=8.0 Hz), 2.30-2.33 (m, 2H), 1.36-1.39 (m, 3H). LCMS (ESI): m/z= 355 $(M+H)^+$

**AVI-3766**

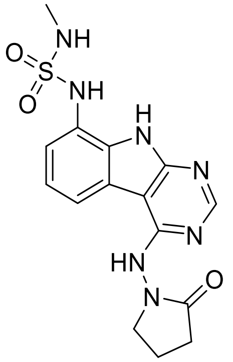


To a solution of 1-((8-amino-9H-pyrimido[4,5-b]indol-4-yl)amino)pyrrolidin-2-one (10 mg, 0.035 mmol), DMAP (2.2 mg, 0.18 mmol) and triethylamine (0.010 mL, 0.071 mmol) in DMA (0.5 mL), was added methylsulfamoyl chloride (15 mg, 0.11 mmol). After stirring at 65 °C for 90 h, the reaction mixture was purified by reverse phase chromatography (water/acetonitrile/0.1% formic acid) to obtain 2.6 mg (17%) of methyl (4-((2-oxopyrrolidin-1-yl)amino)-9H-pyrimido[4,5-b]indol-8-yl)sulfonamide formic acid salt (**AVI-3766**) as brown solid. $^1$H NMR (METHANOL-$d_4$, 400 MHz) δ 8.44 (s, 1H), 8.01 (d, 1H, *J*=7.8 Hz), 7.51 (d, 1H, *J*=7.8 Hz), 7.34 (t, 1H, *J*=7.9 Hz), 3.84 (t, 2H, *J*=7.2 Hz), 2.68 (s, 3H), 2.60 (t, 2H, *J*=8.2 Hz), 2.28-2.32 (m, 2H). LCMS (ESI): m/z= 376 $(M+H)^+$

**AVI-4051**

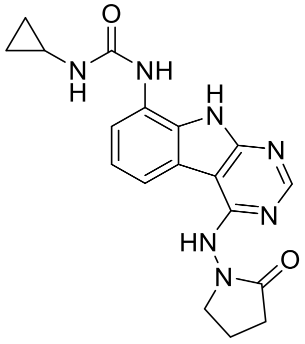


To a solution of 1-((8-amino-9H-pyrimido[4,5-b]indol-4-yl)amino)pyrrolidin-2-one (20 mg, 0.071 mmol) and triethylamine (0.040 mL, 0.28 mmol) in THF (1 mL), was added cyclopropyl isocyanate (24 mg, 0.28 mmol). After stirring at 65 °C for 48 h, the reaction mixture was purified by reverse phase chromatography (water/acetonitrile/0.1% formic acid) to obtain 12 mg (41%) of 1-cyclopropyl-3-(4-((2-oxopyrrolidin-1-yl)amino)-9H-pyrimido[4,5-b]indol-8-yl)urea formic acid salt (**AVI-4051**) as a white solid. $^1$H NMR (DMSO-$d_6$, 400 MHz) δ 11.81 (br s, 1H), 9.31 (s, 1H), 8.50 (br s, 1H), 8.41 (s, 1H), 7.99 (d, 1H, *J*=7.8 Hz), 7.63 (d, 1H, *J*=7.8 Hz), 7.21 (t, 1H, *J*=7.9 Hz), 6.74 (br s, 1H), 3.70 (br t, 2H, *J*=7.1 Hz), 3.12-3.17 (m, 2H), 2.54-2.65 (m, 1H), 2.39-2.43 (m, 2H), 0.98 (t, 2H, *J*=7.1 Hz), 0.68-0.70 (m, 2H). LCMS (ESI): m/z= 366 $(M+H)^+$

**AVI-4057**

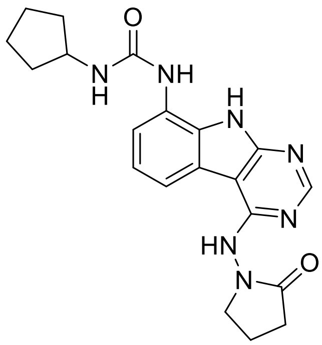

To a solution of 1-((8-amino-9H-pyrimido[4,5-b]indol-4-yl)amino)pyrrolidin-2-one (20 mg, 0.071 mmol) and triethylamine (0.040 mL, 0.28 mmol) in THF (1 mL), was added cyclopentyl isocyanate (0.016 mL, 0.14 mmol). After stirring at 65 ºC for 18 h, the reaction mixture was purified by reverse phase chromatography (water/acetonitrile/0.1% formic acid) to obtain 6 mg (20%) of 1-cyclopentyl-3-(4-((2-oxopyrrolidin-1-yl)amino)-9H-pyrimido[4,5-b]indol-8-yl)urea formic acid salt (**AVI-4057**) as a white solid. $^{1}$H NMR (DMSO-d$_6$, 400 MHz) δ 11.74 (br s, 1H), 9.31 (s, 1H), 8.41 (s, 1H), 7.97 (d, 1H, *J*=7.8 Hz), 7.58 (d, 1H, *J*=7.8 Hz), 7.35 (s, 1H), 7.20 (t, 1H, *J*=7.9 Hz), 6.47 (br d, 1H, *J*=7.1 Hz), 4.03-4.05 (m, 1H), 3.70 (t, 2H, *J*=7.1 Hz), 2.41 (t, 2H, *J*=8.0 Hz), 2.13 (br dd, 2H, *J*=7.8, 15.6 Hz), 1.89 (br dd, 2H, *J*=5.5, 12.1 Hz), 1.67-1.69 (m, 2H), 1.46-1.58 (m, 4H). LCMS (ESI): m/z= 394 $(M+H)^+$

**AVI-4211**

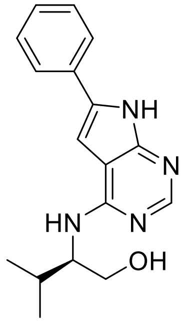

A mixture of (R)-2-((6-bromo-7*H*-pyrrolo[2,3-*d*]pyrimidin-4-yl)amino)-3-methylbutan-1-ol (15.0 mg, 50.1 μmol), phenylboronic acid (12.2 mg, 100.0 μmol), Pd(dppf)$Cl_2$ (3.7 mg, 5.01 μmol) and $Cs_2CO_3$ (40.8 mg, 125 μmol) in 0.22 mL of dioxane/$H_2O$ (10:1) was stirred at 110 °C for 17 hours. The residue was purified by prep-HPLC (0%-70% water/ACN with 0.1% formic acid) to give (R)-3-methyl-2-((6-phenyl-7H-pyrrolo[2,3-d]pyrimidin-4-yl)amino)butan-1-ol, formic acid salt (**AVI-4211**) as a white solid (9.7 mg, yield: 57%). 1H NMR (400 MHz, MeOD) δ 8.41 (brs, 1H), 8.11 (s, 1H), 7.79 (brd, 1H, *J* = 8.0 Hz), 7.45 (brdd, 2H, *J* = 8.0, 7.5 Hz), 7.33 (brt, 1H, *J* = 7.5 Hz), 7.03 (s, 1H), 4.16-4.12 (m, 1H), 3.85-3.75 (m, 2H), 2.15-2.08 (m, 1H), 1.08-1.04 (m, 6H). LCMS (ESI): m/z=297 $(M+H)^+$

**AVI-4678**

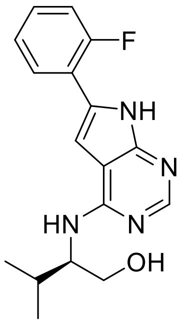

A mixture of (R)-2-((6-bromo-7*H*-pyrrolo[2,3-*d*]pyrimidin-4-yl)amino)-3-methylbutan-1-ol (15.0 mg, 50.1 µmol), (2-fluorophenyl)boronic acid (14.0 mg, 100.0 µmol), Pd(dppf)$Cl_2$ (3.7 mg, 5.01 µmol) and $Cs_2CO_3$ (40.8 mg, 125 µmol) in 0.22 mL of of dioxane/$H_2O$ (10:1) was stirred at 110 °C for 2 hours. The residue was purified by prep-HPLC (0%-70% water/ACN with 0.1% formic acid) to give (R)-2-((6-(2-fluorophenyl)-7H-pyrrolo[2,3-d]pyrimidin-4-yl)amino)-3-methylbutan-1-ol, formic acid salt (**AVI-4678**) as a white solid (10.9 mg, yield: 60%). 1H NMR (400 MHz, MeOD) δ 8.39 (brs, 1H), 8.13 (s, 1H), 7.83-7.78 (m, 1H), 7.37-7.20 (m, 3H), 7.16 (s, 1H), 4.17-4.12 (m, 1H), 3.86-3.75 (m, 2H), 2.17-2.07 (m, 1H), 1.08-1.03 (m, 6H). LCMS (ESI): m/z=315 $(M+H)^+$

**AVI-4335**

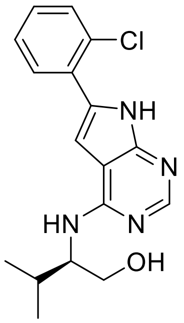

A mixture of (R)-2-((6-bromo-7*H*-pyrrolo[2,3-*d*]pyrimidin-4-yl)amino)-3-methylbutan-1-ol (15.0 mg, 50.1 µmol), (2-chlorophenyl)boronic acid (15.7 mg, 100.0 µmol), Pd(dppf)$Cl_2$ (3.7 mg, 5.01 µmol) and $Cs_2CO_3$ (40.8 mg, 125 µmol) in 0.22 mL of of dioxane/$H_2O$ (10:1) was stirred at 110 °C for 1 hour. The residue was purified by prep-HPLC (0%-50% water/ACN with 0.1% formic acid) to give (R)-2-((6-(2-chlorophenyl)-7H-pyrrolo[2,3-d]pyrimidin-4-yl)amino)-3-methylbutan-1-ol, formic acid salt (**AVI-4335**) as a white solid (8.3 mg, yield: 44%). 1H NMR (400 MHz, MeOD) δ 8.16 (brs, 1H), 7.66 (brd, 1H, *J* = 7.7 Hz), 7.55 (brd, 1H, *J* = 7.9 Hz), 7.45-7.35 (m, 2H), 7.13 (s, 1H), 4.15-4.11 (m, 1H), 3.86-3.75 (m, 2H), 2.15-2.07 (m, 1H), 1.08-1.04 (m, 6H). LCMS (ESI): m/z=331 $(M+H)^+$

**AVI-4370**

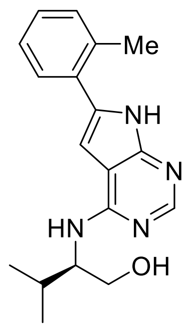


A mixture of (R)-2-((6-iodo-7H-pyrrolo[2,3-d]pyrimidin-4-yl)amino)-3-methylbutan-1-ol (15.0 mg, 43.3 μmol), *o*-tolylboronic acid (11.8 mg, 86.7 μmol), Pd(dppf)$Cl_2$ (3.2 mg, 4.3 μmol) and $Cs_2CO_3$ (35.3 mg, 108 μmol) in 0.22 mL of of dioxane/$H_2O$ (10:1) was stirred at 110 °C for 1 hour. The residue was purified by prep-HPLC (0%-50% water/ACN with 0.1% formic acid) to give (R)-3-methyl-2-((6-(o-tolyl)-7H-pyrrolo[2,3-d]pyrimidin-4-yl)amino)butan-1-ol, formic acid salt (**AVI-4370**) as a white solid (5.4 mg, yield: 35%). 1H NMR (400 MHz, MeOD) δ 8.35 (brs, 1H), 8.13 (s, 1H), 7.51-7.49 (m, 1H), 7.34-7.27 (m, 3H), 7.16-7.14 (brs, 1H), 6.68 (brs, 1H), 4.13-4.11 (m, 1H), 3.86-3.75 (m, 2H), 2.50 (s, 3H), 2.14-2.09 (m, 1H), 1.08-1.04 (m, 6H). LCMS (ESI): m/z=311 $(M+H)^+$

**AVI-4213**

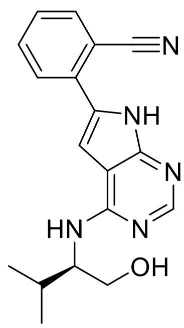


A mixture of (R)-2-((6-bromo-7*H*-pyrrolo[2,3-*d*]pyrimidin-4-yl)amino)-3-methylbutan-1-ol (15.0 mg, 50.1 μmol), (2-cyanophenyl)boronic acid (14.7 mg, 100.0 μmol), Pd(dppf)$Cl_2$ (3.7 mg, 5.01 μmol) and $Cs_2CO_3$ (40.8 mg, 125 μmol) in 0.22 mL of of dioxane/$H_2O$ (10:1) was stirred at 110 °C for 4 hours. The residue was purified by prep-HPLC (0%-50% water/ACN) to give (R)-2-(4-((1-hydroxy-3-methylbutan-2-yl)amino)-7H-pyrrolo[2,3-d]pyrimidin-6-yl)benzonitrile (**AVI-4213**), as a white solid (1.2 mg, yield: 7%). 1H NMR (400 MHz, MeOD) δ 8.52 (brs, 1H), 8.16 (s, 1H), 7.88-7.75 (m, 3H), 7.52 (ddd, 1H, $J$ = 7.6, 7.6, 1.4 Hz), 7.33 (s, 1H), 4.21-4.14 (m, 1H), 3.85-3.75 (m, 2H), 2.14-2.09 (m, 1H), 1.08-1.04 (m, 6H). LCMS (ESI): m/z=322 $(M+H)^+$

**AVI-4271**

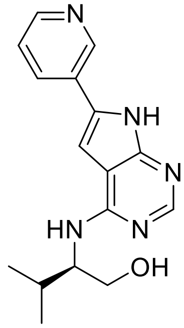


A mixture of (R)-2-((6-bromo-7*H*-pyrrolo[2,3-*d*]pyrimidin-4-yl)amino)-3-methylbutan-1-ol (15.0 mg, 50.1 µmol), pyridin-3-ylboronic acid (18.5 mg, 150.0 µmol), Pd(dppf)$Cl_2$ (7.4 mg, 10.0 µmol) and $Cs_2CO_3$ (40.8 mg, 125 µmol) in 0.22 mL of of dioxane/$H_2O$ (10:1)was stirred at 110 °C for 24 hours. The residue was purified by prep-HPLC (0%-30% water/ACN with 0.1% formic acid) to give (R)-3-methyl-2-((6-(pyridin-3-yl)-7H-pyrrolo[2,3-d]pyrimidin-4-yl)amino)butan-1-ol, formic acid salt (**AVI-4271**) as a white solid (3.7 mg, yield: 22%). 1H NMR (400 MHz, MeOD) δ 8.98 (s, 1H), 8.48 (d, 1H, *J* = 4.8 Hz), 8.20 (d, 1H, *J* = 8.1 Hz), 8.13 (s, 1H), 7.52 (dd, 1H, *J* = 8.1, 4.7 Hz), 7.16 (brs, 1H), 4.18 (ddd, 1H, *J* = 7.6, 5.8, 4.1 Hz), 3.85-3.76 (m, 2H), 2.16-2.07 (m, 1H), 1.08-1.04 (m, 6H). LCMS (ESI): m/z=298 $(M+H)^+$

**AVI-4272**

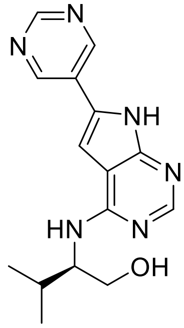


A mixture of (R)-2-((6-bromo-7*H*-pyrrolo[2,3-*d*]pyrimidin-4-yl)amino)-3-methylbutan-1-ol (15.0 mg, 50.1 µmol), pyrimidin-5-ylboronic acid (24.8 mg, 200.0 µmol), Pd(dppf)$Cl_2$ (18.4 mg, 25.1 µmol) and $Cs_2CO_3$ (40.8 mg, 125 µmol) in 0.22 mL of of dioxane/$H_2O$ (10:1)was stirred at 110 °C for 4 days. The residue was purified by prep-HPLC (0%-30% water/ACN with 0.1% formic acid) to give (R)-3-methyl-2-((6-(pyrimidin-5-yl)-7H-pyrrolo[2,3-d]pyrimidin-4-yl)amino)butan-1-ol, formic acid salt (**AVI-4272**) as a white solid (1.0 mg, yield: 6%). 1H NMR (400 MHz, MeOD) δ 9.19 (s, 2H), 9.08 (s, 1H), 8.48 (brs, 2H), 8.16 (s, 1H), 7.26 (s, 1H), 4.22-4.18 (m, 1H), 3.85-3.75 (m, 2H), 2.14-2.07 (m, 1H), 1.08-1.04 (m, 6H). LCMS (ESI): m/z=299 $(M+H)^+$

**AVI-4094**

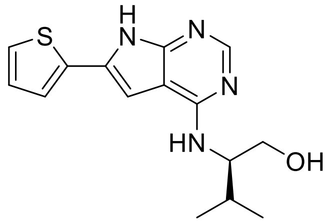

A mixture of (R)-2-((6-bromo-7*H*-pyrrolo[2,3-*d*]pyrimidin-4-yl)amino)-3-methylbutan-1-ol (15 mg, 50.1 µmol), thiophen-2-ylboronic acid (12.8 mg, 100.0 µmol), Pd(dppf)$Cl_2$ (3.7 mg, 5.01 µmol) and $Cs_2CO_3$ (40.8 mg, 125 µmol) in 0.22 mL of dioxane/$H_2O$ (10:1) was stirred at 110 °C for 10 min. The residue was purified by prep-HPLC (0%-50% water/ACN with 0.1% formic acid) to give (R)-3-methyl-2-((6-(thiophen-2-yl)-7H-pyrrolo[2,3-d]pyrimidin-4-yl)amino)butan-1-ol, formic acid salt (**AVI-4094**) as a white solid (13.2 mg, yield: 87%). 1H NMR (400 MHz, MeOD) δ 8.30 (brs, 1H), 8.11 (s, 1H), 7.43 (dd, 1H, *J* = 3.6, 1.4 Hz), 7.39 (dd, 1H, *J* = 5.1, 1.4 Hz), 7.12-7.09 (m, 1H), 6.89 (s, 1H), 4.14-4.04 (m, 1H), 3.85-3.74 (m, 2H), 2.14-2.06 (m, 1H), 1.07-1.03 (m, 6H). LCMS (ESI): m/z=303 $(M+H)^+$

**AVI-4334**

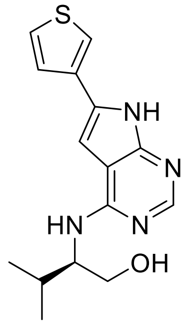

A mixture of (R)-2-((6-bromo-7*H*-pyrrolo[2,3-*d*]pyrimidin-4-yl)amino)-3-methylbutan-1-ol (15 mg, 50.1 µmol), thiophen-3-ylboronic acid (12.8 mg, 100.0 µmol), Pd(dppf)$Cl_2$ (3.7 mg, 5.01 µmol) and $Cs_2CO_3$ (40.8 mg, 125 µmol) in 0.22 mL of dioxane/$H_2O$ (10:1)was stirred at 110 °C for 2 hours. The residue was purified by prep-HPLC (0%-60% water/ACN with 0.1% formic acid) to give (R)-3-methyl-2-((6-(thiophen-3-yl)-7H-pyrrolo[2,3-d]pyrimidin-4-yl)amino)butan-1-ol, formic acid salt (**AVI-4334**) as a white solid (10.5 mg, yield: 60%). 1H NMR (400 MHz, MeOD) δ 8.10 (s, 1H), 7.72-7.70 (m, 1H), 7.52-7.50 (m, 2H), 6.87 (brs, 1H), 4.13 (ddd, 1H, *J* = 7.5, 5.7, 4.1 Hz), 3.84-3.75 (m, 2H), 2.14-2.09 (m, 1H), 1.08-1.03 (m, 6H). LCMS (ESI): m/z=303 $(M+H)^+$

**AVI-4684**

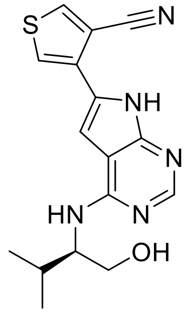


A mixture of (R)-2-((6-iodo-7H-pyrrolo[2,3-d]pyrimidin-4-yl)amino)-3-methylbutan-1-ol (15.0 mg, 43.3 µmol), 4-(4,4,5,5-tetramethyl-1,3,2-dioxaborolan-2-yl)thiophene-3-carbonitrile (16.3 mg, 69.3 µmol), Pd(dppf)$Cl_2$ (3.2 mg, 4.33 µmol) and $Cs_2CO_3$ (35.3 mg, 108 µmol) in 0.22 mL of dioxane/$H_2O$ (10:1) was stirred at 110 °C for 1 hour. The residue was purified by prep-HPLC (0%-80% water/ACN) to give (R)-4-(4-((1-hydroxy-3-methylbutan-2-yl)amino)-7H-pyrrolo[2,3-d]pyrimidin-6-yl)thiophene-3-carbonitrile (**AVI-4684**), as a white solid (4.5 mg, yield: 32%). 1H NMR (400 MHz, *d*-DMSO) δ 12.0 (brs, 1H), 8.71 (d, 1H, $J$ = 3.1 Hz), 8.11 (s, 1H), 8.02 (d, 1H, $J$ = 3.1 Hz), 7.43 (brd, 1H, $J$ = 8.6 Hz), 7.28 (s, 1H), 4.65-4.62 (m, 1H), 4.25-4.18 (m, 1H), 3.60-3.57 (m, 2H), 2.07-1.99 (m, 1H), 0.96-0.92 (m, 6H). LCMS (ESI): m/z=328 $(M+H)^+$

**AVI-4683**

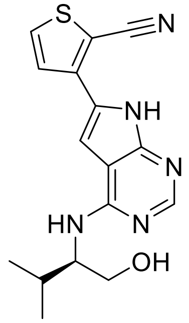


A mixture of (R)-2-((6-iodo-7H-pyrrolo[2,3-d]pyrimidin-4-yl)amino)-3-methylbutan-1-ol (15.0 mg, 43.3 µmol), 3-(4,4,5,5-tetramethyl-1,3,2-dioxaborolan-2-yl)thiophene-2-carbonitrile (16.3 mg, 69.3 µmol), Pd(dppf)$Cl_2$ (3.2 mg, 4.33 µmol) and $Cs_2CO_3$ (35.3 mg, 108 µmol) in 0.22 mL of dioxane/$H_2O$ (10:1) was stirred at 110 °C for 1 hour. The residue was purified by prep-HPLC (0%-60% water/ACN) to give (R)-3-(4-((1-hydroxy-3-methylbutan-2-yl)amino)-7H-pyrrolo[2,3-d]pyrimidin-6-yl)thiophene-2-carbonitrile (**AVI-4683**), as a white solid (5.3 mg, yield: 37%). 1H NMR (400 MHz, *d*-DMSO) δ 12.2 (brs, 1H), 8.15-8.12 (m, 2H), 7.74 (d, 1H, $J$ = 5.2 Hz), 7.57 (brd, 1H, $J$ = 8.7 Hz), 7.50 (s, 1H), 4.65-4.64 (m, 1H), 4.25-4.17 (m, 1H), 3.61-3.58 (m, 2H), 2.06-2.01 (m, 1H), 0.96-0.93 (m, 6H). LCMS (ESI): m/z=328 $(M+H)^+$

**AVI-4267**

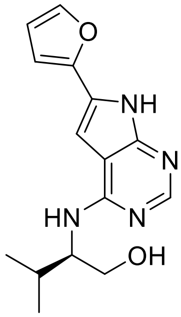

A mixture of (R)-2-((6-bromo-7*H*-pyrrolo[2,3-*d*]pyrimidin-4-yl)amino)-3-methylbutan-1-ol (15.0 mg, 50.1 μmol), 2-(furan-2-yl)-4,4,5,5-tetramethyl-1,3,2-dioxaborolane (19.5 mg, 100.0 μmol), Pd(dppf)$Cl_2$ (3.7 mg, 5.0 μmol) and $Cs_2CO_3$ (40.8 mg, 125 μmol) in 0.22 mL of dioxane/$H_2O$ (10:1) was stirred at 110 °C for 5 minutes. The residue was purified by prep-HPLC (0%-50% water/ACN with 0.1% formic acid) to give (R)-2-((6-(furan-2-yl)-7H-pyrrolo[2,3-d]pyrimidin-4-yl)amino)-3-methylbutan-1-ol, formic acid salt (**AVI-4267**) as a white solid (6.7 mg, yield: 40%). 1H NMR (400 MHz, MeOD) δ 8.11 (brs, 1H), 7.59-7.57 (m, 1H), 6.90 (m, 1H), 6.75 (m, 1H), 6.57-6.54 (m, 1H), 4.15-4.11 (m, 1H), 3.84-3.74 (m, 1H), 2.15-2.06 (m, 1H), 1.07-1.02 (m, 6H). LCMS (ESI): m/z=287 $(M+H)^+$

**AVI-4097**

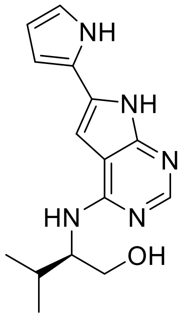

A mixture of (R)-2-((6-bromo-7*H*-pyrrolo[2,3-*d*]pyrimidin-4-yl)amino)-3-methylbutan-1-ol (15.0 mg, 50.1 μmol), (1-(tert-butoxycarbonyl)-1*H*-pyrrol-2-yl)boronic acid (21.2 mg, 100.0 μmol), Pd(dppf)$Cl_2$ (3.67 mg, 5.0 μmol) and $Cs_2CO_3$ (40.8 mg, 125 μmol) in 0.22 mL of dioxane/$H_2O$ (10:1) was stirred at 110 °C for 24 hours. The residue was purified by NP-column (only DCM → DCM/MeOH 95:5 with $NH_3$) to give tert-butyl (*R*)-2-(4-((1-hydroxy-3-methylbutan-2-yl)amino)-7*H*-pyrrolo[2,3-*d*]pyrimidin-6-yl)-1*H*-pyrrole-1-carboxylate as a brown solid (11.9 mg, yield: 62%). LCMS (ESI): m/z=386 $(M+H)^+$
Add solution of tert-butyl (*R*)-2-(4-((1-hydroxy-3-methylbutan-2-yl)amino)-7*H*-pyrrolo[2,3-*d*]pyrimidin-6-yl)-1*H*-pyrrole-1-carboxylate (11.9 mg, 30.9 μmol) in 0.20 mL of methanol dropwise at 0 °C to a solution of acetyl chloride (44.0 μL, 618 mmol). Stir the reaction mixture for 2 days at room temperature. To remove HCl from this residue, it was washed with 7 M $NH_3$ in MeOH and evaporated three times repeatedly. The residue was purified by prep-HPLC (0%-50% water/ACN with 0.1% formic acid) to give (R)-2-((6-(1H-pyrrol-2-yl)-7H-pyrrolo[2,3-

d]pyrimidin-4-yl)amino)-3-methylbutan-1-ol, formic acid salt (**AVI-4097**) as a white solid (5.8 mg, yield: 66%). 1H NMR (400 MHz, MeOD) δ 8.31 (brs, 1H), 8.07 (brs, 1H), 6.87-6.86 (m, 1H), 6.68-6.66 (m, 1H), 6.57-6.55 (m, 1H), 6.21-6.19 (m, 1H), 4.10-4.04 (m, 1H), 3.85-3.75 (m, 2H), 2.17-2.04 (m, 1H), 1.08-1.04 (m, 6H). LCMS (ESI): m/z=286 $(M+H)^+$

**AVI-4212**

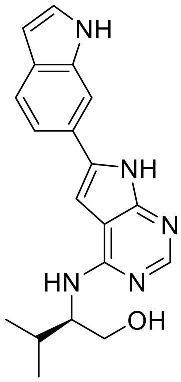

A mixture of (R)-2-((6-bromo-7*H*-pyrrolo[2,3-*d*]pyrimidin-4-yl)amino)-3-methylbutan-1-ol (15.0 mg, 50.1 μmol), (1*H*-indol-6-yl)boronic acid (16.1 mg, 200.0 μmol), Pd(dppf)$Cl_2$ (3.7 mg, 5.01 μmol) and $Cs_2CO_3$ (40.8 mg, 125 μmol) in 0.22 mL of dioxane/$H_2O$ (10:1)was stirred at 110 °C for 17 hours. The residue was purified by prep-HPLC (0%-50% water/ACN with 0.1% formic acid) to give (R)-2-((6-(1H-indol-6-yl)-7H-pyrrolo[2,3-d]pyrimidin-4-yl)amino)-3-methylbutan-1-ol, formic acid salt (**AVI-4212**) as a white solid (8.1 mg, yield: 42%). 1H NMR (400 MHz, MeOD) δ 8.43 (brs, 1H), 8.10 (s, 1H), 7.81 (s, 1H), 7.62 (d, 1H, *J* = 8.3 Hz), 7.48 (d, 1H, *J* = 8.3 Hz), 7.30 (d, 1H, *J* = 3.1 Hz), 6.97 (s, 1H), 6.48 (d, 1H, *J* = 3.1 Hz), 4.14-4.10 (m, 1H), 3.87-3.76 (m, 2H), 2.17-2.08 (m, 1H), 1.09-1.05 (m, 6H). LCMS (ESI): m/z=336 $(M+H)^+$

**AVI-4099**

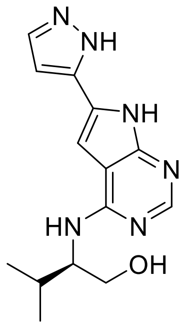

A mixture of (R)-2-((6-bromo-7*H*-pyrrolo[2,3-*d*]pyrimidin-4-yl)amino)-3-methylbutan-1-ol (10.0 mg, 33.4 μmol), 5-(4,4,5,5-tetramethyl-1,3,2-dioxaborolan-2-yl)-1*H*-pyrazole (13.0 mg, 66.9 μmol), Pd(dppf)$Cl_2$ (4.9 mg, 6.7 μmol) and CsOH (12.5 mg, 83.6 μmol) in 0.25 mL of $^n$BuOH/$H_2O$ (4:1) was stirred at 130 °C for 20 minutes with microwave. The residue was purified by prep-HPLC (0%-30% water/ACN with 0.1% formic acid) to give (R)-2-((6-(1H-pyrazol-5-yl)-7H-pyrrolo[2,3-d]pyrimidin-4-yl)amino)-3-methylbutan-1-ol, formic acid salt (**AVI-4099**) as a white solid (3.7 mg, yield: 39%). 1H NMR (400 MHz, MeOD) δ 8.42 (brs, 1H),

8.12 (brs, 1H), 7.73 (d, 1H, *J* = 2.3 Hz), 6.97 (s, 1H), 6.72 (d, 1H, *J* = 2.3 Hz), 4.16-4.11 (m, 1H), 3.84-3.75 (m, 2H), 2.16-2.06 (m, 1H), 1.09-1.02 (m, 6H). LCMS (ESI): m/z=287 $(M+H)^+$

**AVI-4268**

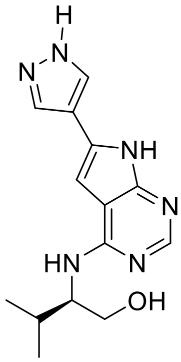


A mixture of (R)-2-((6-bromo-7*H*-pyrrolo[2,3-*d*]pyrimidin-4-yl)amino)-3-methylbutan-1-ol (15.0 mg, 50.1 μmol), (1*H*-pyrazol-4-yl)boronic acid (22.4 mg, 200.0 μmol), $Pd(dppf)Cl_2$ (7.3 mg, 10.0 μmol) and $Cs_2CO_3$ (40.8 mg, 125 μmol) in 0.22 mL of dioxane/$H_2O$ (10:1)was stirred at 110 °C for 4 days. The residue was purified by prep-HPLC (0%-40% water/ACN with 0.1% formic acid) to give (R)-2-((6-(1H-pyrazol-4-yl)-7H-pyrrolo[2,3-d]pyrimidin-4-yl)amino)-3-methylbutan-1-ol, formic acid salt (**AVI-4268**) as a white solid (2.7 mg, yield: 16%). 1H NMR (400 MHz, MeOD) δ 8.48 (brs, 1H), 8.15-8.14 (m, 2H), 7.77 (m, 1H), 6.71 (s, 1H), 6.57 (m, 1H), 4.18-4.14 (m, 1H), 3.84-3.74 (m, 2H), 2.16-2.01 (m, 1H), 1.07-1.03 (m, 6H). LCMS (ESI): m/z=287 $(M+H)^+$

**AVI-4214**

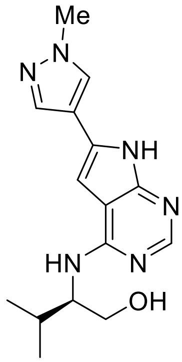


A mixture of (R)-2-((6-bromo-7*H*-pyrrolo[2,3-*d*]pyrimidin-4-yl)amino)-3-methylbutan-1-ol (15.0 mg, 50.1 μmol), (1-methyl-1*H*-pyrazol-4-yl)boronic acid (12.6 mg, 100.0 μmol), $Pd(dppf)Cl_2$ (3.7 mg, 5.01 μmol) and $Cs_2CO_3$ (40.8 mg, 125 μmol) in 0.22 mL of dioxane/$H_2O$ (10:1) was stirred at 110 °C for 4 hours. The residue was purified by prep-HPLC (0%-50% water/ACN with 0.1% formic acid) to give (R)-3-methyl-2-((6-(1-methyl-1H-pyrazol-4-yl)-7H-pyrrolo[2,3-d]pyrimidin-4-yl)amino)butan-1-ol, formic acid salt (**AVI-4214**) as a white solid (11.1 mg, yield: 64%). 1H NMR (400 MHz, MeOD) δ 8.29 (brs, 1H), 8.08 (s, 1H), 7.97 (s, 1H), 7.86 (s, 1H), 6.75 (brs, 1H), 4.10-4.05 (m, 1H), 3.96 (s, 3H), 3.86-3.74 (m, 2H), 2.14-2.06 (m, 1H), 1.09-1.03 (m, 6H). LCMS (ESI): m/z=301 $(M+H)^+$

**AVI-4266**

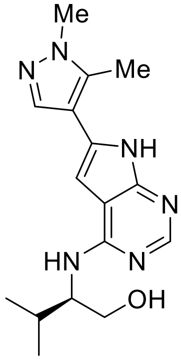


A mixture of (R)-2-((6-bromo-7*H*-pyrrolo[2,3-*d*]pyrimidin-4-yl)amino)-3-methylbutan-1-ol (15.0 mg, 50.1 μmol), 1,5-dimethyl-4-(4,4,5,5-tetramethyl-1,3,2-dioxaborolan-2-yl)-1*H*-pyrazole (44.6 mg, 200.0 μmol), Pd(dppf)$Cl_2$ (7.3 mg, 10.0 μmol) and $Cs_2CO_3$ (40.8 mg, 125 μmol) in 0.22 mL of dioxane/$H_2O$ (10:1) was stirred at 110 °C for 2 days. The residue was purified by prep-HPLC (0%-50% water/ACN with 0.1% formic acid) to give (R)-2-((6-(1,5-dimethyl-1H-pyrazol-4-yl)-7H-pyrrolo[2,3-d]pyrimidin-4-yl)amino)-3-methylbutan-1-ol (**AVI-4266**), formic acid salt as a white solid (4.1 mg, yield: 23%). 1H NMR (400 MHz, MeOD) δ 8.52 (brs, 1H), 8.08 (s, 1H), 7.74 (s, 1H), 6.67 (s, 1H), 4.15-4.10 (m, 1H), 3.88-3.87 (m, 3H), 3.84-3.74 (m, 2H), 2.53-2.52 (m, 3H), 2.14-2.07 (m, 1H), 1.07-1.03 (m, 6H). LCMS (ESI): m/z=315 $(M+H)^+$

**AVI-4054**

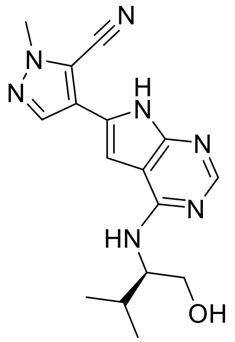


To a solution of 4-(4-chloro-7H-pyrrolo[2,3-d]pyrimidin-6-yl)-1-methyl-1H-pyrazole-5-carbonitrile (70 mg, 0.27 mmol) in dry DMSO (2 mL) was added (*R*)-2-amino-3-methylbutan-1-ol (42.23 mg, 0.41 mmol) and triethylamine (72.4 mg, 0.81 mmol). After stirring at 100$^o$C for 16 h, the mixture was diluted with ethyl acetate (40.0 mL) and washed with water (5.0 mL) and brine (5.0 mL). The organic layer was dried over $Na_2SO_4$ and concentrated under reduced pressure. The residue was purified by prep-HPLC (10%-100% 0.1% $NH_4HCO_3$ in water/ACN) to give (R)-4-(4-((1-hydroxy-3-methylbutan-2-yl)amino)-7H-pyrrolo[2,3-d]pyrimidin-6-yl)-1-methyl-1H-pyrazole-5-carbonitrile (**AVI-4054**) as a white solid (25.4 mg, yield: 28.8%). 1H NMR (500 MHz, MeOD) δ 8.09 (s, 1H), 7.98 (s, 1H), 7.13 (s, 1H), 4.14 (s, 1H), 4.09 (s, 3H), 3.77 (dd, J = 13.4, 5.0 Hz, 2H), 2.08 (dt, J = 13.8, 6.9 Hz, 1H), 1.02 (dd, J = 13.2, 6.8 Hz, 6H). LCMS (ESI): m/z 326.1 $(M+H)^+$

**AVI-5707**

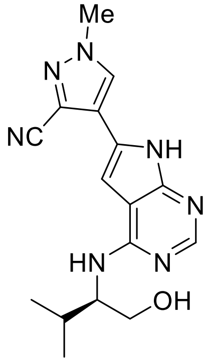

A mixture of (R)-2-((6-iodo-7H-pyrrolo[2,3-d]pyrimidin-4-yl)amino)-3-methylbutan-1-ol (15 mg, 43.3 umol), 1-methyl-4-(4,4,5,5-tetramethyl-1,3,2-dioxaborolan-2-yl)-1H-pyrazole-3-carbonitrile (16.2 mg, 69.3 umol), Pd(dppf)$Cl_2$ (3.2 mg, 4.33 umol) and CsCO3 (35.3 mg, 108 mmol) in 0.22 mL of dioxane/$H_2O$ (10:1) was stirred at 110 °C for 1 hour. Subsequently, the mixture was extracted thrice with EtOAc, the combined organic phase was washed with brine, dried over $Na_2SO_4$ and concentrated in vacuo. The residue was purified by silica gel chromatography (20-40%EtOAc in hexane and prep-HPLC, 2.1 mg (13%) of (R)-4-(4-((1-hydroxy-3-methylbutan-2-yl)amino)-7H-pyrrolo[2,3-d]pyrimidin-6-yl)-1-methyl-1H-pyrazole-3-carbonitrile, formic acid salt (**AVI-5707**)as a white solid. $^1$H NMR (DMSO-$d_6$, 400 MHz) δ 11.96-12.03 (m, 1H), 8.44 (br s, 1H), 8.28-8.30 (m, 1H), 8.08 (s, 1H), 7.40 (br d, 1H, *J*=8.0 Hz), 7.14 (s, 1H), 4.19 (br s, 1H), 4.00 (s, 3H), 3.57-3.61 (m, 2H), 2.02 (qd, 1H, *J*=6.7, 13.6 Hz), 1.24 (s, 1H), 0.94 (t, 6H, *J*=6.5 Hz). LCMS (ESI): m/z=326 $(M+H)^+$

**AVI-4100**

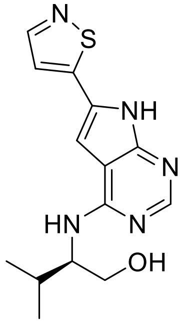

A mixture of (R)-2-((6-bromo-7*H*-pyrrolo[2,3-*d*]pyrimidin-4-yl)amino)-3-methylbutan-1-ol (10.0 mg, 33.4 μmol), 5-(4,4,5,5-tetramethyl-1,3,2-dioxaborolan-2-yl)isothiazole (14.1 mg, 66.9 μmol), $Pd(PPh_3)_4$ (7.7 mg, 6.7 μmol) and $Na_2CO_3$ (7.09 mg, 66.9 μmol) in 0.22 mL of dioxane/$H_2O$ (10:1) was stirred at 110 °C for 24 hours. The residue was purified by prep-HPLC (0%-40% water/ACN with 0.1% formic acid) to give (R)-2-((6-(isothiazol-5-yl)-7H-pyrrolo[2,3-d]pyrimidin-4-yl)amino)-3-methylbutan-1-ol, formic acid salt (**AVI-4100**) as a white solid (1.2 mg, yield: 10%). 1H NMR (400 MHz, MeOD) δ 8.55 (brs, 1H), 8.49 (d, 1H, *J* = 1.9 Hz), 8.15 (s, 1H), 7.63 (d, 1H, *J* = 1.9 Hz), 7.14 (s, 1H), 4.19-4.18 (m, 1H), 3.83-3.74 (m, 2H), 2.13-2.08 (m, 1H), 1.07-1.03 (m, 6H). LCMS (ESI): m/z=304 $(M+H)^+$

**AVI-4052**

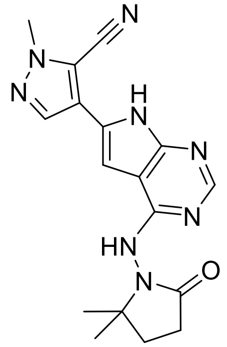


A solution of 4-(4-chloro-7H-pyrrolo[2,3-d]pyrimidin-6-yl)-1-methyl-1H-pyrazole-5-carbonitrile (70 mg, 0.27 mmol) and 1-amino-5,5-dimethylpyrrolidin-2-one (63.76 mg, 0.41 mmol) in iPrOH (2 mL) and HCl (1 drop) was stirred at 100 °C for 16 hours. The reaction mixture was suspended in water (30 mL x 3) and extracted with ethyl acetate (3 x 60 mL). The organic layer was dried over $Na_2SO_4$ and concentrated under reduced pressure. The residue was purified by prep-HPLC (10%-100% 0.1% $NH_4HCO_3$ in water/ACN) to give 4-(4-((2,2-dimethyl-5-oxopyrrolidin-1-yl)amino)-7H-pyrrolo[2,3-d]pyrimidin-6-yl)-1-methyl-1H-pyrazole-5-carbonitrile (**AVI-4052**) as a white solid (25.1 mg, yield: 26.4%). 1H NMR (500 MHz, MeOD) δ 8.32 (s, 1H), 8.04 (s, 1H), 7.34 (s, 1H), 4.12 (s, 3H), 2.60 (d, J = 7.7 Hz, 2H), 2.22 (s, 2H), 1.37 (s, 6H). LCMS (ESI): m/z 351.1 $(M+H)^+$

**AVI-6320**

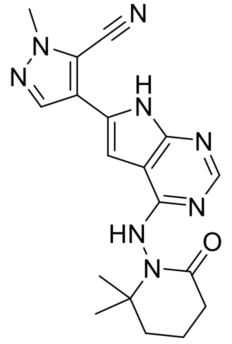


To a solution of 4-(4-chloro-7H-pyrrolo[2,3-d]pyrimidin-6-yl)-1-methyl-1H-pyrazole-5-carbonitrile (120 mg, 0.46 mmol) in IPA (5 mL) was added 1-amino-6,6-dimethylpiperidin-2-one (99 mg, 0.70 mmol) and conc. HCl (0.04 ml). After stirring at 100°C for 24 hours, aqueous, saturated sodium bicarbonate (5 ml) was added to the reaction mixture and extracted with DCM (20 ml×3), organic layers were separated and concentrated in vacuo, and purified by preprarative HPLC (10%-40% 0.1% $NH_4HCO_3$ in water/ACN) to obtain 4-(4-((2,2-dimethyl-6-oxopiperidin-1-yl)amino)-7H-pyrrolo[2,3-d]pyrimidin-6-yl)-1-methyl-1H-pyrazole-5-carbonitrile (**AVI-6320**) as a white solid (24mg, Yield:14.45%). $^1$H NMR (400 MHz, DMSO) δ 12.11 (s, 1H), 9.15 (s, 1H), 8.15 (d, J = 12.7 Hz, 2H), 6.92 (s, 1H), 4.04 (s, 3H), 2.68 – 2.52 (m, 1H), 2.39 – 2.25 (m, 1H), 2.05 (s, 1H), 1.84 (d, J = 5.5 Hz, 3H), 1.30 (d, J = 26.5 Hz, 6H). LCMS (ESI): m/z= 365.3 $(M+H)^+$

**AVI-6318**

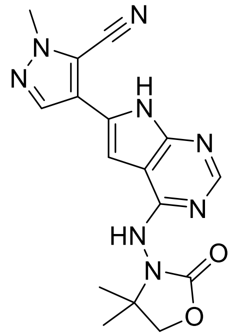

To a solution of 4-(4-chloro-7-((2-(trimethylsilyl)ethoxy)methyl)-7H-pyrrolo[2,3-d]pyrimidin-6-yl)-1-methyl-1H-pyrazole-5-carbonitrile (200 mg, 0.52 mmol) in IPA (5 mL) were added 3-amino-4,4-dimethyloxazolidin-2-one (98 mg, 0.75 mmol) and conc. HCl (0.04 ml). After stirring at 100°C for 24 hours, aqueous saturated sodium bicarbonate (5 ml) was added to the reaction mixture and extracted with DCM (20 ml×3), organic layers were separated and concentrated in vacuo and purified by silica gel chromatography (dichloromethane/methanol, 10:1) to obtain 4-(4-((4,4-dimethyl-2-oxooxazolidin-3-yl)amino)-7-((2-(trimethylsilyl)ethoxy)methyl)-7H-pyrrolo[2,3-d]pyrimidin-6-yl)-1-methyl-1H-pyrazole-5-carbonitrile (55 mg, Yield:21.9%) as a white solid. LCMS (ESI): m/z= 483.2 $(M+H)^+$
To a solution of 4-(4-((4,4-dimethyl-2-oxooxazolidin-3-yl)amino)-7-((2-(trimethylsilyl)ethoxy)methyl)-7H-pyrrolo[2,3-d]pyrimidin-6-yl)-1-methyl-1H-pyrazole-5-carbonitrile (55 mg, 0.11mmol) in DCM (2.0 ml) was added TFA (0.5 ml). After stirring at 20 °C for 4 h, the reaction mixture was concentrated to dryness, the resulting residue was suspended in aqueous saturated NaHCO3 and extracted with DCM (10 ml × 3). The organic layers were washed with brine, concentrated under reduced pressure and purified by preprarative HPLC (10%-50% 0.1% $NH_4HCO_3$ in water/ACN) to obtain 4-(4-((4,4-dimethyl-2-oxooxazolidin-3-yl)amino)-7H-pyrrolo[2,3-d]pyrimidin-6-yl)-1-methyl-1H-pyrazole-5-carbonitrile (**AVI-6318**)as a white solid (20 mg, Yield: 29.8%). $^1$H NMR (500 MHz, DMSO) δ 12.44 (s, 1H), 9.76 (s, 1H), 8.25 (s, 1H), 8.17 (s, 1H), 7.08 (s, 1H), 4.41 – 4.15 (m, 2H), 4.07 (s, 3H), 1.31 (d, *J* = 11.9 Hz, 6H). LCMS (ESI): m/z= 353.1 $(M+H)^+$

**AVI-6319**

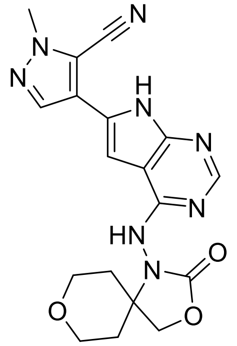

To a solution of 4-(4-chloro-7-((2-(trimethylsilyl)ethoxy)methyl)-7H-pyrrolo[2,3-d]pyrimidin-6-yl)-1-methyl-1H-pyrazole-5-carbonitrile (200 mg, 0.52 mmol) in IPA(5 mL) were added 1-

amino-3,8-dioxa-1-azaspiro[4.5]decan-2-one (129 mg, 0.75 mmol) and conc. HCl (0.04 ml ). After stirring at 100°C for 24 hours, aqueous saturated sodium bicarbonate (5 ml) was added to the reaction mixture and extracted with DCM (20 ml×3), organic layers were separated and concentrated in vacuo and purified by silica gel chromatography (dichloromethane/methanol, 10:1) to obtain 1-methyl-4-(4-((2-oxo-3,8-dioxa-1-azaspiro[4.5]decan-1-yl)amino)-7-((2-(trimethylsilyl)ethoxy)methyl)-7H-pyrrolo[2,3-d]pyrimidin-6-yl)-1H-pyrazole-5-carbonitrile (60 mg, Yield:22%) as a white solid. LCMS (ESI): m/z= 525.3 $(M+H)^+$
To a solution of 1-methyl-4-(4-((2-oxo-3,8-dioxa-1-azaspiro[4.5]decan-1-yl)amino)-7-((2-(trimethylsilyl)ethoxy)methyl)-7H-pyrrolo[2,3-d]pyrimidin-6-yl)-1H-pyrazole-5-carbonitrile (60 mg, 0.11mmol) in DCM (2.0 ml) was added TFA (0.5 ml). After stirring at 20 °C for 4 h, the reaction mixture was concentrated to dryness, the resulting residue was suspended in aqueous saturated NaHCO3 and extracted with DCM (10 ml × 3). The organic layers were washed with brine, concentrated under reduced pressure and purified by preprarative HPLC (10%-50% 0.1% $NH_4HCO_3$ in water/ACN) to obtain 1-methyl-4-(4-((2-oxo-3,8-dioxa-1-azaspiro[4.5]decan-1-yl)amino)-7H-pyrrolo[2,3-d]pyrimidin-6-yl)-1H-pyrazole-5-carbonitrile (**AVI-6319**) as a white solid (27 mg, yield: 62.3%). $^1$H NMR (500 MHz, DMSO) δ 12.47 (s, 1H), 9.88 (s, 1H), 8.25 (s, 1H), 8.18 (s, 1H), 7.13 (s, 1H), 4.50 (d, *J* = 76.4 Hz, 2H), 4.07 (s, 3H), 3.91 (dd, *J* = 11.8, 4.2 Hz, 1H), 3.78 (d, *J* = 9.5 Hz, 1H), 3.46 (s, 1H), 3.26 (d, *J* = 12.2 Hz, 1H), 1.96 (td, *J* = 12.9, 4.7 Hz, 1H), 1.85 (s, 1H), 1.76 – 1.56 (m, 2H). LCMS (ESI): m/z= 395.1 $(M+H)^+$

**AVI-6345**

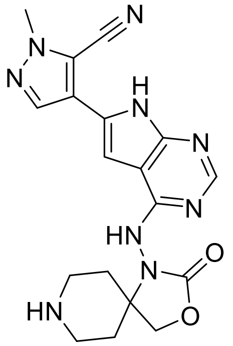


To a solution of 4-(4-chloro-7H-pyrrolo[2,3-d]pyrimidin-6-yl)-1-methyl-1H-pyrazole-5-carbonitrile (500 mg, 1.93 mmol) in IPA (15 mL) were added benzyl 1-amino-2-oxo-3-oxa-1,8-diazaspiro[4.5]decane-8-carboxylate (707 mg, 2.32 mmol) and conc. HCl (0.06 ml). After stirring at 100°C for 24 hours, aqueous saturated sodium bicarbonate (5 ml) was added to the reaction mixture and extracted with DCM (20 ml×3), organic layers were separated and concentrated in vacuo and purified by silica gel chromatography (dichloromethane/methanol, 10:1) to obtain benzyl 1-((6-(5-cyano-1-methyl-1H-pyrazol-4-yl)-7H-pyrrolo[2,3-d]pyrimidin-4-yl)amino)-2-oxo-3-oxa-1,8-diazaspiro[4.5]decane-8-carboxylate (30 mg, Yield: 2.9%) as a white solid. LCMS (ESI): m/z= 528.1 $(M+H)^+$
To a solution of benzyl 1-((6-(5-cyano-1-methyl-1H-pyrazol-4-yl)-7H-pyrrolo[2,3-d]pyrimidin-4-yl)amino)-2-oxo-3-oxa-1,8-diazaspiro[4.5]decane-8-carboxylate (30 mg, 0.057mmol) in acetonitrile (2.0 ml) was added TMSI (23 mg, 0.114 mmol). After stirring at 20 °C for 4 h, the reaction mixture was concentrated to dryness, the resulting residue was suspended in aqueous saturated NaHCO3 and extracted with DCM (10 ml × 3). The organic layers were washed with brine, concentrated under reduced pressure and purified by prep-TLC (DCM/MeOH=10/1) to

obtain 1-methyl-4-(4-((2-oxo-3-oxa-1,8-diazaspiro[4.5]decan-1-yl)amino)-7H-pyrrolo[2,3-d]pyrimidin-6-yl)-1H-pyrazole-5-carbonitrile (**AVI-6345**) as a white solid (6 mg, Yield: 26.8%). 1H NMR (500 MHz, DMSO) δ 12.44 (s, 1H), 9.85 (s, 1H), 8.24 (s, 1H), 8.17 (s, 1H), 7.13 (s, 1H), 4.39 (d, J = 69.5 Hz, 2H), 4.06 (s, 3H), 2.95 (d, J = 12.4 Hz, 1H), 2.82 (d, J = 12.5 Hz, 1H), 2.64-2.62 (m, 1H), 2.54-2.32 (m, 2H), 1.92 – 1.70 (m, 2H), 1.60-1.52 (m, 2H). LCMS (ESI): m/z= 394.1 $(M+H)^+$

**AVI-6344**

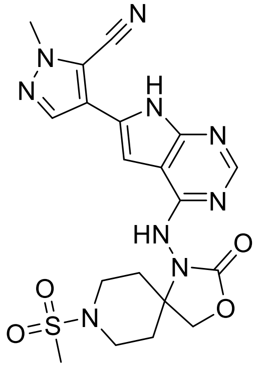

To a solution of 4-(4-chloro-7H-pyrrolo[2,3-d]pyrimidin-6-yl)-1-methyl-1H-pyrazole-5-carbonitrile (100 mg, 0.39 mmol) in IPA (5 mL) were added 1-amino-8-(methylsulfonyl)-3-oxa-1,8-diazaspiro[4.5]decan-2-one (149mg, 0.60 mmol) and conc. HCl (0.04 ml). After stirring at 100°C for 24 hours, aqueous saturated sodium bicarbonate (5 ml) was added to the reaction mixture and extracted with DCM (20 ml×3), organic layers were separated and concentrated in vacuo and purified by prep-HPLC (10%-50% 0.1% $NH_4HCO_3$ in water/ACN) to obtain 1-methyl-4-(4-((8-(methylsulfonyl)-2-oxo-3-oxa-1,8-diazaspiro[4.5]decan-1-yl)amino)-7H-pyrrolo[2,3-d]pyrimidin-6-yl)-1H-pyrazole-5-carbonitrile (**AVI-6344**) as a white solid (22 mg, yield: 12.0%). $^1$H NMR (500 MHz, DMSO) δ 12.47 (s, 1H), 9.88 (s, 1H), 8.25 (s, 1H), 8.18 (s, 1H), 7.02 (d, *J* = 100.1 Hz, 1H), 4.78 – 4.20 (m, 2H), 4.07 (s, 3H), 3.63 (d, *J* = 12.0 Hz, 1H), 3.50 (d, *J* = 12.2 Hz, 1H), 2.93 (s, 1H), 2.86 (s, 3H), 2.76 (dd, *J* = 22.8, 12.0 Hz, 1H), 2.16 – 1.89 (m, 3H), 1.76 (d, *J* = 65.1 Hz, 2H). LCMS (ESI): m/z= 472.0 $(M+H)^+$

**AVI-6315** & **AVI-6316**

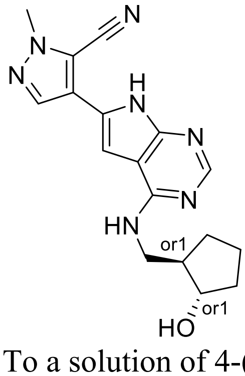

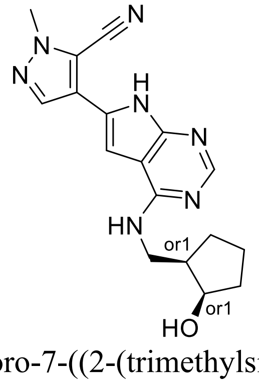

To a solution of 4-(4-chloro-7-((2-(trimethylsilyl)ethoxy)methyl)-7H-pyrrolo[2,3-d]pyrimidin-6-yl)-1-methyl-1H-pyrazole-5-carbonitrile (500 mg, 1.29 mmol) in dry DMSO (10 mL) were added 2-(aminomethyl)cyclopentan-1-ol (223 mg, 1.93 mmol) and TEA (651mg, 6.44mmol)

**AVI-6317**

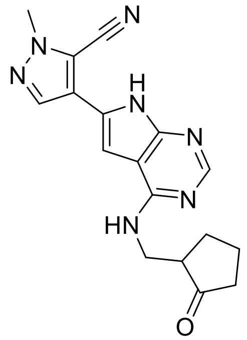

To a cooled (-78°C) solution of oxalyl chloride (82mg, 0.64mmol) in DCM (3mL) was added DMSO (82 mg, 1.07 mmol) in DCM (1mL) dropwise while stirring at under nitrogen gas. After 30 min, 4-(4-((((1R,2S)-2-hydroxycyclopentyl)methyl)amino)-7-((2-(trimethylsilyl)ethoxy)methyl)-7H-pyrrolo[2,3-d]pyrimidin-6-yl)-1-methyl-1H-pyrazole-5-carbonitrile (200 mg, 0.43 mmol) in DCM (1mL) was added dropwise and the mixture was stirred for an additional 90 min at -78°C . Triethylamine (433mg, 4.3mmol) was then added dropwise and the reaction mixture was allowed to warm to room temperature. To this mixture was added water (10 mL) and extracted with DCM (10 ml × 3). The organic layers were washed with water, dried over Na2SO4, filtered, and concentrated under reduced pressure. The residue was purified by prep-HPLC (10mmol $NH_4HCO_3$ in water, 10%-60% MeCN) to obtain 1-methyl-4-(4-(((2-oxocyclopentyl)methyl)amino)-7-((2-(trimethylsilyl)ethoxy)methyl)-7H-pyrrolo[2,3-d]pyrimidin-6-yl)-1H-pyrazole-5-carbonitrile (90 mg, Yield: 45.2%) as a white solid. LCMS (ESI): m/z= 466 $(M+H)^+$

To a solution of 1-methyl-4-(4-(((2-oxocyclopentyl)methyl)amino)-7-((2-(trimethylsilyl)ethoxy)methyl)-7H-pyrrolo[2,3-d]pyrimidin-6-yl)-1H-pyrazole-5-carbonitrile (80 mg, 0.19 mmol) in DCM (3 ml) was added TFA (1 ml). After stirring at 20°C for 4h, the reaction mixture was concentrated to dryness. To the resulting residue was added saturated aqueous NaHCO3 (5 ml) and extracted with DCM (30 ml × 2). The organic layers were washed with brine and concentrated under reduced pressure. The crude was purified by prep-HPLC (10%-100% 10mmol $NH_4HCO_3$ in water/ ACN) to obtain 1-methyl-4-(4-(((2-oxocyclopentyl)methyl)amino)-7H-pyrrolo[2,3-d]pyrimidin-6-yl)-1H-pyrazole-5-carbonitrile (**AVI-6317**) as a white solid (28 mg, yield: 40.7%). $^1$H NMR (500 MHz, DMSO) δ 12.07 (s, 1H), 8.14 (d, *J* = 3.9 Hz, 2H), 7.89 – 7.74 (m, 1H), 7.06 (s, 1H), 4.05 (s, 3H), 3.85 (dt, *J* = 13.2, 5.4 Hz, 1H), 3.37 (dd, *J* = 8.1, 5.7 Hz, 1H), 2.67 – 2.52 (m, 1H), 2.28 – 2.05 (m, 3H), 1.98 – 1.85 (m, 1H), 1.83 – 1.55 (m, 2H). LCMS (ESI): m/z= 336.5 $(M+H)^+$

**AVI-6254**

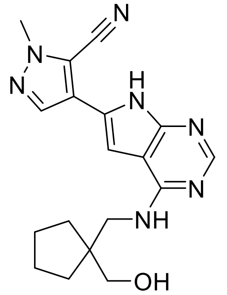

To a solution of 4-(4-chloro-7-((2-(trimethylsilyl)ethoxy)methyl)-7H-pyrrolo[2,3-d]pyrimidin-6-yl)-1-methyl-1H-pyrazole-5-carbonitrile (100.0 mg, 0.25 mmol) in DMSO (2 mL) was added (1-(aminomethyl)cyclopentyl)methanol (49.9 mg, 0.38 mmol) and TEA (52.1 mg, 0.51 mmol). After stirred at 100°C for 1 hour, the reaction mixture was extracted with ethyl acetate (3 x 20 mL), washed with brine (10 mL) and dried over $Na_2SO_4$. The organic layers were concentrated and the residue was purified by silica gel column (DCM/MeOH, 20：1) to obtain 4-(4-(((1-(hydroxymethyl)cyclopentyl)methyl)amino)-7-((2-(trimethylsilyl)ethoxy)methyl)-7H-pyrrolo[2,3-d]pyrimidin-6-yl)-1-methyl-1H-pyrazole-5-carbonitrile as a yellow solid (120 mg, Yield: 96.79%). LCMS (ESI): m/z= 482.4 $(M+H)^+$
To a solution of 4-(4-chloro-7-((2-(trimethylsilyl)ethoxy)methyl)-7H-pyrrolo[2,3-d]pyrimidin-6-yl)-1-methyl-1H-pyrazole-5-carbonitrile (120 mg, 0.25 mmol) in DCM (2 ml) was added TFA (1.0 mL). After stirring at 20°C for 3 h, the reaction mixture was concentrated to dryness. To the resulting residue was added saturated aqueous NaHCO3 (5 ml) and extracted with DCM (30 ml × 2). The organic layers were washed with brine and concentrated under reduced pressure. The crude was purified by prep-HPLC (10%-50% 0.1% $NH_4HCO_3$ in water/ ACN) to obtain 4-(4-(((1-(hydroxymethyl)cyclopentyl)methyl)amino)-7H-pyrrolo[2,3-d]pyrimidin-6-yl)-1-methyl-1H-pyrazole-5-carbonitrile (**AVI-6254**) as a white solid (29 mg, Yield: 33.33%). 1H NMR (500 MHz, DMSO-d6) δ 12.19 (s, 1H), 8.12 (d, J = 12.0 Hz, 2H), 7.85 (s, 1H), 7.11 (s, 1H), 5.41 (s, 1H), 4.05 (s, 3H), 3.44 (d, J = 6.4 Hz, 2H), 3.14 (d, J = 6.6 Hz, 2H), 1.58 (dd, J = 17.5, 6.9 Hz, 4H), 1.49 – 1.43 (m, 2H), 1.40 – 1.34 (m, 2H). LCMS (ESI): m/z= 352.4 $(M+H)^+$

**AVI-6253**

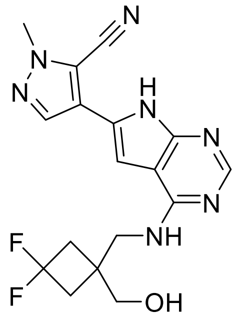

To a solution of 4-(4-chloro-7-((2-(trimethylsilyl)ethoxy)methyl)-7H-pyrrolo[2,3-d]pyrimidin-6-yl)-1-methyl-1H-pyrazole-5-carbonitrile (120 mg, 0.31 mmol) in dry DMSO (5 mL) was added (1-(aminomethyl)-3,3-difluorocyclobutyl)methanol (70 mg, 0.46 mmol) and TEA (156 mg, 1.55 mmol). After stirring at 110°C for 3 h, the reaction mixture was diluted with ethyl acetate (50.0

mL), washed with water (10.0 mL) and brine (10.0 mL). The organic layer was dried over $Na_2SO_4$ and concentrated under reduced pressure. The residue was purified by column chromatography on silica gel (DCM/MeOH, 10:1)) to obtain 4-(4-(((3,3-difluoro-1-(hydroxymethyl)cyclobutyl)methyl)amino)-7-((2-(trimethylsilyl)ethoxy)methyl)-7H-pyrrolo[2,3-d]pyrimidin-6-yl)-1-methyl-1H-pyrazole-5-carbonitrile (100 mg, Yield: 64.3%） as a white solid. LCMS (ESI): m/z= 504.3 $(M+H)^+$
To a solution of 4-(4-(((3,3-difluoro-1-(hydroxymethyl)cyclobutyl)methyl)amino)-7-((2-(trimethylsilyl)ethoxy)methyl)-7H-pyrrolo[2,3-d]pyrimidin-6-yl)-1-methyl-1H-pyrazole-5-carbonitrile (100 mg, 0.2 mmol) in DCM (3 ml) was added TFA (1 ml). After stirring at 20°C for 4 h, the reaction mixture was concentrated to dryness. To the resulting residue was added saturated aqueous NaHCO3 (5 ml) and extracted with DCM (30 ml × 2). The organic layers were washed with brine and concentrated under reduced pressure. The crude was purified by prep-HPLC (10%-100% 10mmol $NH_4HCO_3$ in water/ACN) to obtain 4-(4-(((3,3-difluoro-1-(hydroxymethyl)cyclobutyl)methyl)amino)-7H-pyrrolo[2,3-d]pyrimidin-6-yl)-1-methyl-1H-pyrazole-5-carbonitrile (**AVI-6253**)as a white solid (27 mg, yield: 36.4%).

**AVI-6252**

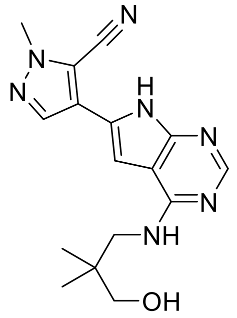


To a solution of 4-(4-chloro-7-((2-(trimethylsilyl)ethoxy)methyl)-7H-pyrrolo[2,3-d]pyrimidin-6-yl)-1-methyl-1H-pyrazole-5-carbonitrile (200 mg, 0.52 mmol) in DMSO (2 mL) was added 3-amino-2,2-dimethylpropan-1-ol (77 mg, 0.75 mmol) and TEA (158 mg ,1.56 mmol). After stirring at 100°C for 4 h, the reaction mixture was diluted with dichloromethane (5 mL) and washed with water (10.0 mL x 3). The organic layer was dried over $Na_2SO_4$ and concentrated under reduced pressure. The residue was purified by silica gel chromatography (DCM/MeOH, 10:1) to obtain 4-(4-((3-hydroxy-2,2-dimethylpropyl)amino)-7-((2-(trimethylsilyl)ethoxy)methyl)-7H-pyrrolo[2,3-d]pyrimidin-6-yl)-1-methyl-1H-pyrazole-5-carbonitrile (90 mg, Yield:38.0%） as a white solid. LCMS (ESI): m/z= 456.2 $(M+H)^+$
To a solution of 4-(4-((3-hydroxy-2,2-dimethylpropyl)amino)-7-((2-(trimethylsilyl)ethoxy)methyl)-7H-pyrrolo[2,3-d]pyrimidin-6-yl)-1-methyl-1H-pyrazole-5-carbonitrile (90 mg, 0.20 mmol) in DCM(2.0 ml) was added TFA (0.5 ml). After stirring at 20°C for 4 h, the reaction mixture was concentrated to dryness. To the resulting residue was added saturated aqueous NaHCO3 (5 ml) and extracted with DCM (30 ml × 2). The organic layers were washed with brine and concentrated under reduced pressure. The crude was purified by prep-HPLC (10%-50% 0.1% $NH_4HCO_3$ in water/MeCN) to obtain 4-(4-((3-hydroxy-2,2-dimethylpropyl)amino)-7H-pyrrolo[2,3-d]pyrimidin-6-yl)-1-methyl-1H-pyrazole-5-carbonitrile (**AVI-6252**) as a white solid (29 mg, yield: 44.6%). $^1H$ NMR (500 MHz, DMSO) δ 12.19 (s,

1H), 8.12 (d, $J$ = 11.9 Hz, 2H), 7.83 (s, 1H), 7.14 (d, $J$ = 1.4 Hz, 1H), 5.27 (s, 1H), 4.05 (s, 3H), 3.34 (d, $J$ = 6.4 Hz, 2H), 3.07 (s, 2H), 0.87 (s, 6H). LCMS (ESI): m/z= 325.3 $(M+H)^+$

**AVI-4206**

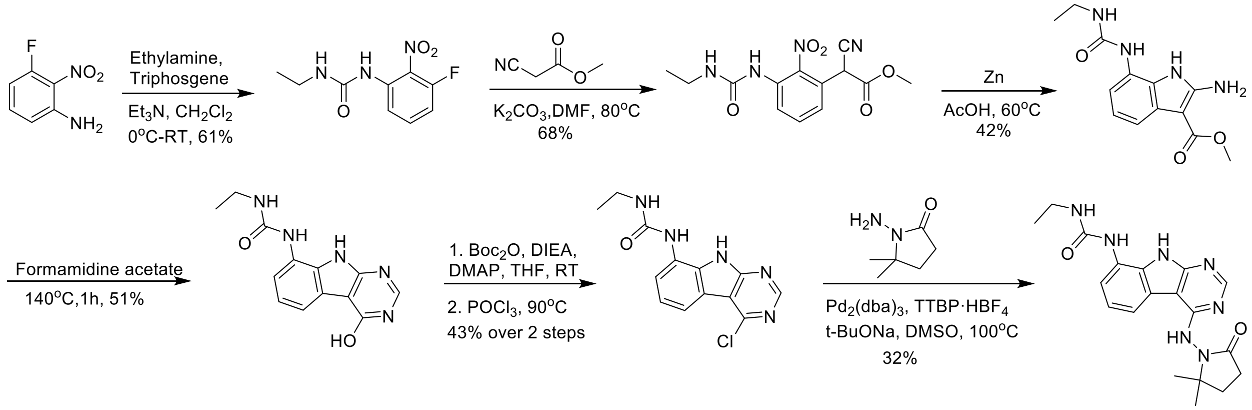


A solution of 3-fluoro-2-nitroaniline (25.00 g, 160 mmol) in THF (500 mL) were added triethylamine (48 g, 480 mmol) and triphosgene (14.2 g, 48 mmol) at 0ºC. After stirring for an hour, ethylamine as 2.0 M solution in THF (200mL) was added. Upon completion of reaction, the mixture was poured into 500mL of water, extracted with ethyl acetate (3 x 500 mL), the combined organic layers were washed with brine (500 mL), dried over $Na_2SO_4$, filtered, concentrated under reduced pressure and the residue was purified by silica gel column chromatography (0-20% ethyl acetate/hexanes) to afford 1-ethyl-3-(3-fluoro-2-nitrophenyl)urea as yellow solid (22.00 g, yield: 60.57%). LCMS (ESI): m/z= 228.1 $(M+H)^+$

To a solution of 1-ethyl-3-(3-fluoro-2-nitrophenyl)urea (48 g, 211.45 mmol) in DMF (300 mL) were added methyl 2-isocyanoacetate (41.86 g, 422.90 mmol) and $K_2CO_3$ (87.54 g, 634.36 mmol). The solution was stirred at 80°C for 16 hours. The mixture was adjusted to be weakly acidic by 2N HCl, extracted with EA (500 mL *3), the combined organic layers were washed with brine (300 mL), dried over $Na_2SO_4$, filtered and concentrated under reduced pressure. The residue was purified by silica gel column chromatography (0-20% ethyl acetate/hexanes) to afford methyl 2-cyano-2-(3-(3-ethylureido)-2-nitrophenyl)acetate as yellow solid (44.3 g, yield: 68.46%). LCMS (ESI): m/z= 307.2 $(M+H)^+$.

A mixture of methyl 2-cyano-2-(3-(3-ethylureido)-2-nitrophenyl)acetate (42 g, 137.25 mmol) and acetic acid (250mL) was heated to 40 °C. Zinc (89.75 g, 1372.54 mmol) was then added in portions at a rate such that the reaction temperature did not rise above 60 °C. After the addition was complete, the reaction mixture was stirred at 60°C for 2 h. The reaction mixture was cooled to room temperature and filtered through a celite pad. The filtrate was concentrated under vacuum. The crude product was purified by silica gel column chromatography (dicholoromethane/methanol 10:1) to obtain methyl 2-amino-7-(3-ethylureido)-1H-indole-3-carboxylate as a white solid (16 g, yield: 42.23%). LCMS (ESI): m/z=277.2 $(M+H)^+$

Methyl 2-amino-7-(3-ethylureido)-1H-indole-3-carboxylate (2.0 g, 7.25 mmol) and formamidine acetate (4.53 g, 43.48 mmol) were heated to 140 °C for 1 h. The mixture was cooled to room temperature and diluted with approximately 100 mL of water. The resulting mixture was stirred for 15 min before the solid was collected by filtration. The residue was triturated with DMSO and filtered to afford 1-ethyl-3-(4-hydroxy-9H-pyrimido[4,5-b]indol-8-yl)urea as an off-white

solid (1.0 g, yield: 50.8%). 1H NMR (500 MHz, DMSO) δ 12.21 (s, 1H), 11.76 (s, 1H), 8.34 (s, 1H), 8.13 (d, J = 3.5 Hz, 1H), 7.64 (d, J = 7.7 Hz, 1H), 7.40 (d, J = 7.3 Hz, 1H), 7.13 (t, J = 7.8 Hz, 1H), 6.28 (t, J = 5.5 Hz, 1H), 3.26 – 3.11 (m, 2H), 1.10 (t, J = 7.2 Hz, 3H). LCMS (ESI): m/z=272.3 $(M+H)^+$
To a solution of 1-ethyl-3-(4-hydroxy-9H-pyrimido[4,5-b]indol-8-yl)urea (500 mg, 1.85 mmol) in THF (20 mL) were added di-*tert*-butyl dicarbonate (1.21 g, 5.54 mmol), DIPEA (955 mg, 7.4 mmol) and DMAP (226 mg, 1.85 mmol). The mixture was stirred at room temperature for 16 hours. The mixture was then concentrated under reduced pressure to give crude *tert*-butyl 4-((*tert*-butoxycarbonyl)oxy)-8-(3-ethylureido)-9H-pyrimido[4,5-b]indole-9-carboxylate as a yellow oil. It was used in the next step without any purification.
A solution of *tert*-butyl 4-((*tert*-butoxycarbonyl)oxy)-8-(3-ethylureido)-9H-pyrimido[4,5-b]indole-9-carboxylate (crude) in POCl3 (10 mL) was stirred at 90°C for 30min. The solution was concentrated under reduced pressure and diluted with acetonitrile, then adjusted the pH to 7.0 with ammonium hydroxide slowly. The resulting solid was filtered with vacuum filter and washed with water to obtain 1-(4-chloro-9H-pyrimido[4,5-b]indol-8-yl)-3-ethylurea (230 mg, two steps yield: 43.1%) as a light yellow solid. LCMS (ESI): m/z=290.2 $(M+H)^+$
To a solution of 1-(4-chloro-9H-pyrimido[4,5-b]indol-8-yl)-3-ethylurea (290 mg,1.0 mmol) in dry DMSO(6.0 ml) was added 1-amino-5,5-dimethylpyrrolidin-2-one (192 mg,1.5 mmol) , $Pd_2(dba)_3$ (92 mg, 0.1 mmol), Tri-*tert*-butylphosphine tetrafluoroborate (44 mg, 0.15 mmol) and t-BuONa(240 mg, 2.5 mmol). After stirring at 100 ºC for 8h, the reaction mixture was filtered and the filtrate was purified by reversed phase chromatography (water /acetonitrile/0.1%TFA). Further purification by silica gel column chromatography (dicholoromethane/methanol 10:1) afforded 1-(4-((2,2-dimethyl-5-oxopyrrolidin-1-yl)amino)-9H-pyrimido[4,5-b]indol-8-yl)-3-ethylurea (**AVI-4206**) (120 mg, yield:31.5%) as a white solid.
1H NMR (400 MHz, DMSO) δ 11.67 (s, 1H), 9.06 (s, 1H), 8.37 (d, J = 15.7 Hz, 2H), 8.07 (d, J = 7.6 Hz, 1H), 7.59 (d, J = 7.8 Hz, 1H), 7.20 (t, J = 7.9 Hz, 1H), 6.28 (t, J = 5.4 Hz, 1H), 3.27 – 3.10 (m, 2H), 2.42 (t, J = 7.8 Hz, 2H), 2.03 (t, J = 7.8 Hz, 2H), 1.26 (d, J = 21.1 Hz, 6H), 1.11 (t, J = 7.2 Hz, 3H).
13C NMR (DMSO-d6, 151 MHz) δ 171.8, 157.9, 155.9, 155.5, 154.9, 128.6, 125.3, 121.1, 120.4, 117.2,117.1, 96.7, 61.1, 34.8, 32.4, 27.8, 26.6, 15.9. LCMS (ESI): m/z= 382 $(M+H)^+$

**AVI-6347**

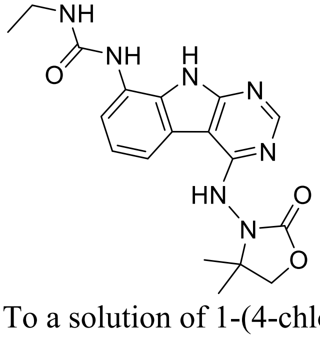


To a solution of 1-(4-chloro-9H-pyrimido[4,5-b]indol-8-yl)-3-ethylurea (100 mg, 0.35 mmol) in dry DMSO (3.0 ml) was added 3-amino-4,4-dimethyloxazolidin-2-one (67 mg, 0.52 mmol) , $Pd_2(dba)_3$ (32 mg, 0.03 mmol), tri-*tert*-butylphosphine tetrafluoroborate (15 mg, 0.05 mmol) and t-BuONa(83 mg, 0.86 mmol). After stirring at 100 ºC for 2 h, the reaction mixture was filtered and the filtrate was purified by reversed phase chromatography (10-32%

acetonitrile/water/0.1%TFA). Further purification by silica gel column chromatography (dicholoromethane/methanol 10:1) afforded 1-(4-((2,2-dimethyl-5-oxopyrrolidin-1-yl)amino)-9H-pyrimido[4,5-b]indol-8-yl)-3-ethylurea (**AVI-6347**) (20 mg, yield:15%) as a white solid. $^{1}$H NMR (500 MHz, DMSO-*d6*) δ 11.74 (s, 1H), 9.28 (s, 1H), 8.45 (d, J = 10.6 Hz, 1H), 8.38 (s, 1H), 8.07 (d, J = 7.8 Hz, 1H), 7.61 (d, J = 7.8 Hz, 1H), 7.22 (t, J = 7.9 Hz, 1H), 6.29 (t, J = 5.5 Hz, 1H), 4.29 (s, 2H), 3.22 – 3.16 (m, 2H), 1.34 (s, 6H), 1.11 (t, J = 7.2 Hz, 3H). LCMS (ESI): m/z=384.3 $(M+H)^+$

**AVI-6187**

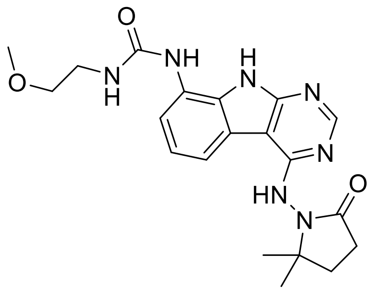

To a solution of 1-((8-amino-9H-pyrido[2,3-b]indol-4-yl)amino)-5,5-dimethylpyrrolidin-2-one (20 mg, 0.064 mmol) and triethylamine (0.026 mL, 0.26 mmol) in THF (1 mL), was added 1-isocyanato-2-methoxyethane (0.013 mg, 0.13 mmol). After stirring at 55 °C for 18 h, the reaction mixture was purified by reverse phase chromatography (water/acetonitrile) to obtain 4.3 mg (16%) 1-(4-((2,2-dimethyl-5-oxopyrrolidin-1-yl)amino)-9H-pyrimido[4,5-b]indol-8-yl)-3-(2-methoxyethyl)urea (**AVI-6187**) as a white solid. 1H NMR (METHANOL-d4, 400 MHz) δ 8.27 (s, 1H), 7.86 (d, 1H, J=8.0 Hz), 7.29 (d, 1H, J=7.8 Hz), 7.14 (t, 1H, J=7.5 Hz), 3.57-3.60 (m, 2H), 3.49-3.51 (m, 2H), 3.44 (s, 3H), 2.61 (t, 2H, J=8.0 Hz), 2.21 (br s, 2H), 1.37 (s, 6H). LCMS (ESI): m/z= 412 $(M+H)^+$

**AVI-6188**

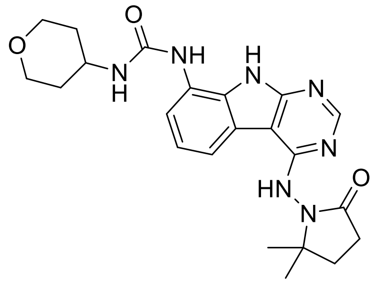

To a solution of 1-((8-amino-9H-pyrido[2,3-b]indol-4-yl)amino)-5,5-dimethylpyrrolidin-2-one (20 mg, 0.064 mmol) and triethylamine (0.026 mL, 0.26 mmol) in THF (1 mL), was added 4-isocyanatotetrahydro-2H-pyran (0.016 mg, 0.13 mmol). After stirring at 55 °C for 18 h, the reaction mixture was purified by reverse phase chromatography (water/acetonitrile) to obtain 3.6 mg (13%) 1-(4-((2,2-dimethyl-5-oxopyrrolidin-1-yl)amino)-9H-pyrimido[4,5-b]indol-8-yl)-3-(tetrahydro-2H-pyran-4-yl)urea (**AVI-6188**) as a white solid. $^{1}$H NMR (METHANOL-$d_4$, 400 MHz) δ 8.33 (s, 1H), 7.94 (dd, 1H, *J*=1.0, 7.8 Hz), 7.32 (dd, 1H, *J*=0.7, 7.8 Hz), 7.23 (t, 1H, *J*=7.8 Hz), 3.88-4.01 (m, 3H), 3.52-3.59 (m, 2H), 2.60 (t, 2H, *J*=7.9 Hz), 2.20 (br s, 2H), 1.98 (br d, 2H, *J*=2.2 Hz), 1.60 (br s, 2H), 1.39 (s, 6H). LCMS (ESI): m/z= 438 $(M+H)^+$

#### AVI-6414

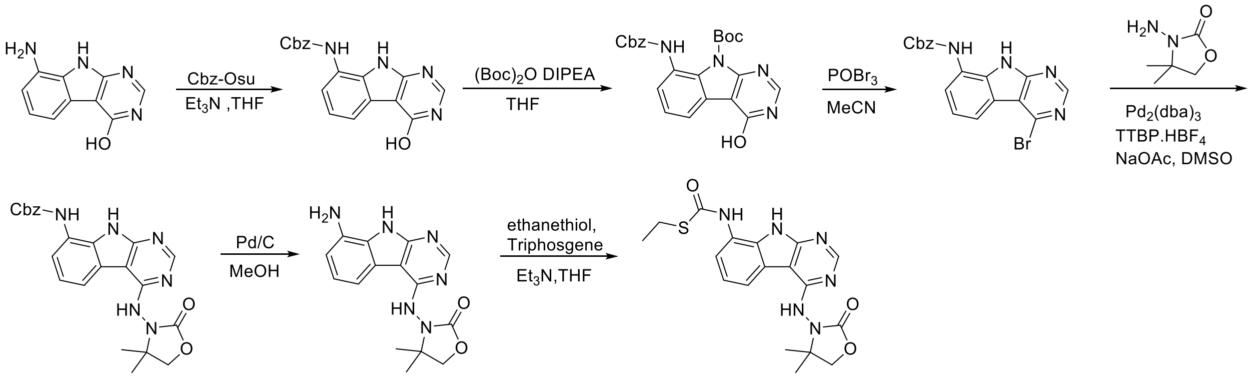


To a suspension of 8-amino-9H-pyrimido[4,5-b]indol-4-ol (5.0 g, 25 mmol) in THF (60 mL) were added triethylamine (7.6 g, 75 mmol) and N-(benzyloxycarbonyloxy)succinimide (9.3 g, 37.5 mmol). After stirring at 80°C for 16 hours, reaction mixture was concentrated under reduced pressure, the residue was poured in 50 ml of MeCN. The resulting mixture was allowed to stand for 15 min before the solids were collected by filtration, washed with MeCN, and dried in vacuo to afford benzyl (4-hydroxy-9H-pyrimido[4,5-b]indol-8-yl)carbamate as an off-white solid (7.5 g, yield: 89.8%). $^{1}$H NMR (500 MHz, DMSO) δ 11.98 (s, 1H), 12.04 – 11.32 (m, 1H), 9.54 (d, *J* = 82.4 Hz, 1H), 8.14 (s, 1H), 7.74 (d, *J* = 7.7 Hz, 1H), 7.61 (dd, *J* = 65.6, 13.5 Hz, 1H), 7.52 – 7.44 (m, 2H), 7.41 (t, *J* = 7.2 Hz, 2H), 7.35 (t, *J* = 7.1 Hz, 1H), 7.20 (t, *J* = 7.8 Hz, 1H), 5.21 (s, 2H). LCMS (ESI): m/z= 335.3 $(M+H)^+$

To a solution of benzyl (4-hydroxy-9H-pyrimido[4,5-b]indol-8-yl)carbamate (2.5 g, 7.5 mmol) in THF (40 mL) were added di-*tert*-butyl dicarbonate (4.9 g, 22.5mmol), DIPEA (4.9 g, 37.5mmol) and DMAP (460 mg, 3.75 mmol). The mixture was stirred at room temperature for 2 h and concentrated under reduced pressure to afford crude *tert*-butyl 8-(((benzyloxy)carbonyl)amino)-4-hydroxy-9H-pyrimido[4,5-b]indole-9-carboxylate as a yellow oil (4g, yield: 100%) which was used in the next step without any purification. LCMS (ESI): m/z=434.1$(M+H)^+$

A solution of *tert*-butyl 8-(((benzyloxy)carbonyl)amino)-4-hydroxy-9H-pyrimido[4,5-b]indole-9-carboxylate (crude) in MeCN (50 mL) were added phosphoryl tribromide (10.6 g, 37.5 mmol) and DIPEA (1 ml). The mixture was stirred at 90°C for 30min, concentrated under reduced pressure and diluted with MeCN, then adjusted the pH 7.0 with ammonium hydroxide slowly. The resulting solid was filtered and washed with water to afford benzyl (4-bromo-9H-pyrimido[4,5-b]indol-8-yl)carbamate (1.1g, two steps of yield: 33.7%) as a light yellow solid. $^{1}$H NMR (500 MHz, DMSO) δ 12.67 – 12.09 (m, 1H), 9.56 (s, 1H), 8.76 (d, *J* = 13.3 Hz, 1H), 8.29 – 8.12 (m, 1H), 7.83 (d, *J* = 61.0 Hz, 1H), 7.53 – 7.32 (m, 6H), 5.22 (d, *J* = 7.9 Hz, 2H). LCMS (ESI): m/z=397 $(M+H)^+$

To a solution of benzyl (4-bromo-9H-pyrimido[4,5-b]indol-8-yl)carbamate (600 mg, 1.52 mmol) in dry DMSO (10 ml) were added 3-amino-4,4-dimethyloxazolidin-2-one (394 mg, 3.04 mmol), $Pd_2(dba)_3$ (138 mg, 0.15 mmol), tri-*tert*-butylphosphine tetrafluoroborate (66 mg, 0.22 mmol) and AcONa (370 mg, 4.56 mmol). After stirring at 100°C for 30min, the reaction mixture was filtered and the filtrate purified by reversed phase chromatography (10mM $NH_4HCO_3$ in water, 5%-60% ACN) to obtain benzyl (4-((4,4-dimethyl-2-oxooxazolidin-3-yl)amino)-9H-pyrimido[4,5-b]indol-8-yl)carbamate as a white solid (100 mg, yield: 14.8%). LCMS (ESI): m/z=447.3 $(M+H)^+$

To a solution of benzyl (4-((4,4-dimethyl-2-oxooxazolidin-3-yl)amino)-9H-pyrimido[4,5-b]indol-8-yl)carbamate (100 mg, 0.22mmol) in THF (5 ml) in MeOH (5 mL) was added 10% Pd/C (30 mg), The mixture was stirred at room temperature for 3 h under $H_2$ gas. The solution was filtered through a celite and the filtrate was concentrated under reduced pressure to obtain 3-((8-amino-9H-pyrimido[4,5-b]indol-4-yl)amino)-4,4-dimethyloxazolidin-2-one as a white solid (60 mg, yield: 86%) which was used in the next step without any purification. LCMS (ESI): m/z=313.1 $(M-56+H)^+$

To a cooled (0°C) solution of 3-((8-amino-9H-pyrimido[4,5-b]indol-4-yl)amino)-4,4-dimethyloxazolidin-2-one (60 mg, 0.19 mmol) in THF (8 mL) were added ethanethiol (24 mg, 0.38 mmol), Et3N (58 mg, 0.58 mmol) and triphosgene (34 mg, 0.12 mmol). After stirring 1h, the mixture was poured into 20mL of water and extracted with ethyl acetate (20mL *3), the combined organic layers were washed with brine (20mL), dried over $Na_2SO_4$, filtered and concentrated under reduced pressure, the residue was purified by silica gel column chromatography (10:1 dichloromethane/methanol) to afford S-ethyl (4-((4,4-dimethyl-2-oxooxazolidin-3-yl)amino)-9H-pyrimido[4,5-b]indol-8-yl)carbamothioate (**AVI-6414**) as a white solid (30 mg, yield: 39%). $^1$H NMR (500 MHz, DMSO) δ 11.94 (s, 1H), 9.93 (s, 1H), 9.35 (s, 1H), 8.47 (s, 1H), 8.23 (d, *J* = 7.7 Hz, 1H), 7.76 (s, 1H), 7.29 (t, *J* = 7.9 Hz, 1H), 4.29 (s, 2H), 2.93 (q, *J* = 7.2 Hz, 2H), 1.49 – 1.19 (m, 9H). LCMS (ESI): m/z=401.1 $(M+H)^+$

**AVI-6249**

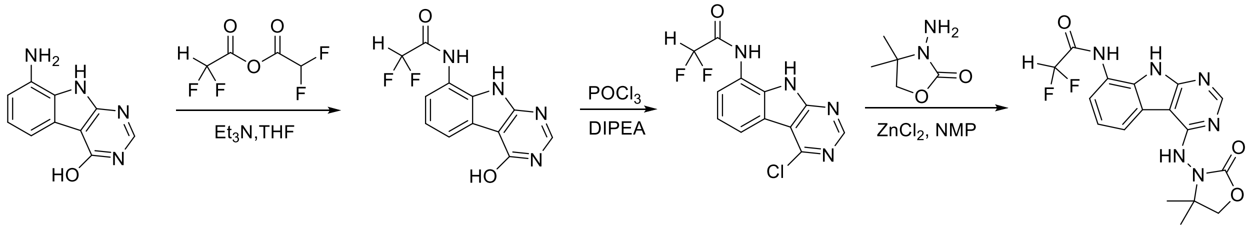


To a suspension of 8-amino-9H-pyrimido[4,5-b]indol-4-ol (2.0 g, 10 mmol) in THF (20 mL) were added $Et_3N$ (3.03 g, 30 mmol) and 2,2-difluoroacetic anhydride (2.61 g, 15 mmol). The reaction was stirred at rt for 3 h, then concentrated under reduced pressure and the residue was poured in 30 ml of water. The resulting mixture was allowed to stand for 15 min before the solids were collected by filtration, washed with water, and dried in vacuo to afford 2,2-difluoro-N-(4-hydroxy-9H-pyrimido[4,5-b]indol-8-yl)acetamide as an off-white solid (2.0 g, yield: 72%). $^1$H NMR (500 MHz, DMSO) δ 12.30 (s, 1H), 12.11 (s, 1H), 10.64 (s, 1H), 8.18 (s, 1H), 7.91 (d, *J* = 7.7 Hz, 1H), 7.42 (t, *J* = 10.4 Hz, 1H), 7.25 (t, *J* = 7.8 Hz, 1H), 6.58 – 6.34 (m, 1H). LCMS (ESI): m/z= 279.3 $(M+H)^+$

A solution of 2,2-difluoro-N-(4-hydroxy-9H-pyrimido[4,5-b]indol-8-yl)acetamide (1g,3.6mmol) in $POCl_3$ (10 mL) and DIPEA (1 ml) was stirred at 90°C for 30min. The solution was concentrated under reduced pressure and diluted with MeCN, then adjusted to pH 7.0 with ammonium hydroxide slowly. The resulting solid was filtered and washed with water to afford N-(4-chloro-9H-pyrimido[4,5-b]indol-8-yl)-2,2-difluoroacetamide (500mg, yield:46.9%) as a brown solid. LCMS (ESI): m/z=297.1 $(M+H)^+$

To a solution of N-(4-chloro-9H-pyrimido[4,5-b]indol-8-yl)-2,2-difluoroacetamide (200 mg, 0.68 mmol) in dry NMP (3 ml) were added 3-amino-4,4-dimethyloxazolidin-2-one (175 mg, 1.35 mmol) and zinc(II) chloride (200mg). After stirring at 150°C for 7h, the reaction was filtered and the filtrate was purified by reversed phase chromatography (10mM $NH_4HCO_3$ in water, 5%-90%

ACN) to obtain N-(4-((4,4-dimethyl-2-oxooxazolidin-3-yl)amino)-9H-pyrimido[4,5-b]indol-8-yl)-2,2-difluoroacetamide (**AVI-6249**) as a white solid (25 mg,  yield: 10%). $^1$H NMR (500 MHz, DMSO) δ 12.12 (s, 1H), 10.66 (s, 1H), 9.41 (s, 1H), 8.49 (s, 1H), 8.33 (t, *J* = 17.9 Hz, 1H), 7.51 (d, *J* = 7.7 Hz, 1H), 7.34 (t, *J* = 7.8 Hz, 1H), 6.46 (t, *J* = 53.8 Hz, 1H), 4.30 (s, 2H), 1.39 (t, *J* = 45.9 Hz, 6H). LCMS (ESI): m/z= 391.2 $(M+H)^+$

**AVI-6425**

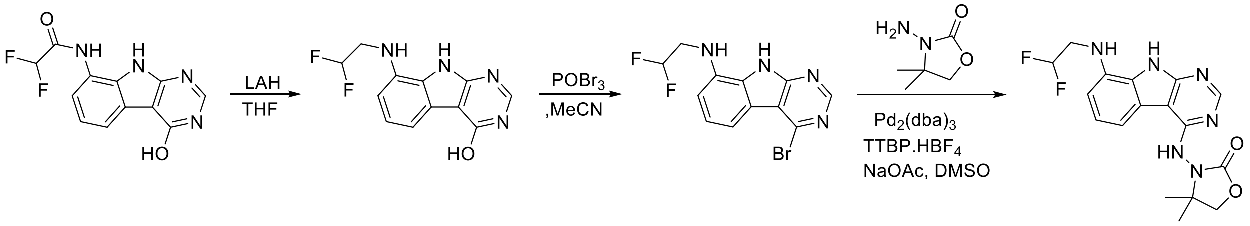


To a cooled (0$^o$C) solution of 2,2-difluoro-N-(4-hydroxy-9H-pyrimido[4,5-b]indol-8-yl)acetamide (1.5 g, 5.4 mmol) in THF (20 mL) was added LAH (6.4 ml,16 mmol). The reaction was warmed up to room temperature & stirred for 3 h. The reaction mixture was cooled the cooled to 0 °C and was quenched by addition of solid $Na_2SO_4$.10H2O (10 g). The resulting mixture was warmed to room temperature and stirred for 0.5 h, filtered through Celite and washed with EtOAc (100mL). The filtrate was concentrated in vacuo to afford 8-((2,2-difluoroethyl)amino)-9H-pyrimido[4,5-b]indol-4-ol as a brown solid (1.0 g, 70%). $^1$H NMR (500 MHz, DMSO) δ 12.12 (s, 1H), 11.75 (d, *J* = 33.5 Hz, 1H), 8.08 (s, 1H), 7.36 – 7.31 (m, 1H), 7.08 – 7.04 (m, 1H), 6.66 (d, *J* = 7.8 Hz, 1H), 6.23 (tt, *J* = 55.8, 3.9 Hz, 1H), 5.70 (t, *J* = 6.3 Hz, 1H), 3.71 (tdd, *J* = 15.8, 6.3, 4.0 Hz, 2H). LCMS (ESI): m/z=265.2 $(M+H)^+$
To a  solution of 8-((2,2-difluoroethyl)amino)-9H-pyrimido[4,5-b]indol-4-ol (1 g, 3.79 mmol) in MeCN (10 mL) was added phosphoryl tribromide (837 g, 30.3 mmol) and DIPEA (1 ml). The mixture was stirred at 90°C for 30min. The reaction mixture was then concentrated under reduced pressure, diluted with ACN, then adjusted to pH 7.0 with ammonium hydroxide slowly. The resulting solid was filtered and the filtrate was concentrated under reduced pressure. The crude product was purified by silica gel column chromatography (10:1 dichloromethane/methanol) to afford 4-bromo-N-(2,2-difluoroethyl)-9H-pyrimido[4,5-b]indol-8-amine as a white solid (600 mg, yield: 48.6%). LCMS (ESI): m/z=327 $(M+H)^+$
To a solution of 4-bromo-N-(2,2-difluoroethyl)-9H-pyrimido[4,5-b]indol-8-amine (300 mg, 0.61 mmol) in dry DMSO (8 ml) were added 3-amino-4,4-dimethyloxazolidin-2-one (239 mg, 1.84 mmol), $Pd_2(dba)_3$ (30 mg, 0.06 mmol), tri-*tert*-butylphosphine tetrafluoroborate (60 mg, 0.09 mmol) and AcONa (252mg, 3.07mmol) and stirred at 100$^o$C for 1h. The reaction mixture was filtered and the filtrate was purified by reversed phase chromatography (0.1%TFA/ACN=0-50%). Further purification by silica gel column chromatography (dicholoromethane/methanol 10:1) afforded 3-((8-((2,2-difluoroethyl)amino)-9H-pyrimido[4,5-b]indol-4-yl)amino)-4,4-dimethyloxazolidin-2-one (**AVI-6425**) as a white solid (20 mg, yield:8.6%). $^1$H NMR (500 MHz, DMSO) δ 11.79 (s, 1H), 9.19 (s, 1H), 8.41 (s, 1H), 7.75 (d, *J* = 7.8 Hz, 1H), 7.15 (t, *J* = 7.9 Hz, 1H), 6.81 (d, *J* = 7.9 Hz, 1H), 6.26 (tt, *J* = 55.7, 3.7 Hz, 1H), 5.77 (t, *J* = 6.3 Hz, 1H), 4.28 (s, 2H), 3.86 – 3.59 (m, 2H), 1.30 (d, *J* = 39.2 Hz, 6H). LCMS (ESI): m/z=377.3 $(M+H)^+$

**AVI-6355**

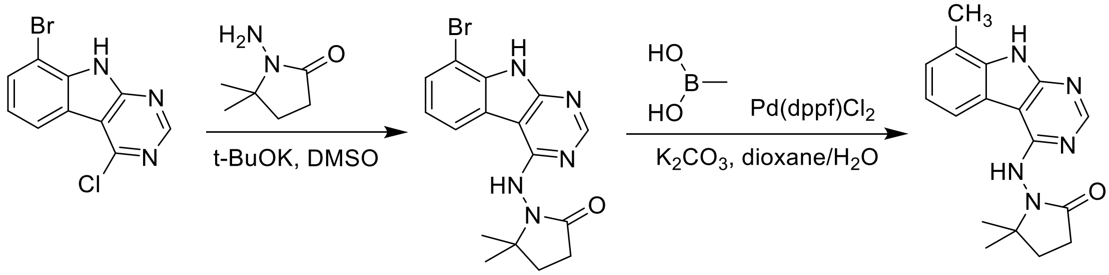

To a solution of 8-bromo-4-chloro-9H-pyrimido[4,5-b]indole (4 g, 14.23 mmol) in dry DMSO (50 mL) was added 1-amino-5,5-dimethylpyrrolidin-2-one (2.73 g, 21.35 mmol) and t-BuOK (3.99 g, 35.58 mmol). The mixture was stirred at 100ºC for 45 minutes, diluted with ethyl acetate (80.0 mL) and washed with water (100.0 mL), brine (100.0 mL). The organic layers were dried over $Na_2SO_4$ and concentrated under reduced pressure. The crude product was purified by silica gel chromatography (dicholoromethane/methanol 10:1) to obtain 1-((8-bromo-9H-pyrimido[4,5-b]indol-4-yl)amino)-5,5-dimethylpyrrolidin-2-one as a brown solid (2 g, yield: 37.66%). $^1$H NMR (500 MHz, DMSO) δ 12.65 (s, 1H), 9.55 (br s, 1H), 8.50 (s, 1H), 8.46 (d, J = 7.8 Hz, 1H), 7.69 (d, J = 7.7 Hz, 1H), 7.29 (t, J = 7.9 Hz, 1H), 2.45 (t, J = 7.9 Hz, 2H), 2.05 (t, J = 7.8 Hz, 2H), 1.29 (s, 6H). LCMS (ESI): m/z=374.1 $(M+H)^+$.

To a solution of 1-((8-bromo-9H-pyrimido[4,5-b]indol-4-yl)amino)-5,5-dimethylpyrrolidin-2-one (120 mg, 0.32 mmol) in dioxane (2 mL) and $H_2O$ (0.2 mL) was added methylboronic acid (38 mg, 0.64 mmol), $K_2CO_3$ (110 mg, 0.80 mmol) and Pd(dppf)$Cl_2$ (35 mg, 0.05 mmol). The mixture was stirred at 95 °C for 16 hours under $N_2$. The reaction mixture was diluted with water (50 mL) and extracted with DCM (3 x 50 mL). The organic layer was dried over $Na_2SO_4$ and concentrated under reduced pressure. The residue was purified by reverse phase chromatography (10mM $NH_4HCO_3$ in water, 5%-35% MeCN) to obtain 5,5-dimethyl-1-((8-methyl-9H-pyrimido[4,5-b]indol-4-yl)amino)pyrrolidin-2-one (**AVI-6355**) as a white solid (25 mg, yield: 14.63%). $^1$H NMR (500 MHz, DMSO-*d6*) δ 12.08 (s, 1H), 9.05 (s, 1H), 8.39 (s, 1H), 8.23 (t, J = 16.0 Hz, 1H), 7.32 – 7.10 (m, 2H), 2.56 (s, 3H), 2.42 (t, J = 7.8 Hz, 2H), 2.03 (t, J = 7.8 Hz, 2H), 1.27 (d, J = 21.2 Hz, 6H). LCMS (ESI): m/z=310.4 $(M+H)^+$.

**AVI-6371**

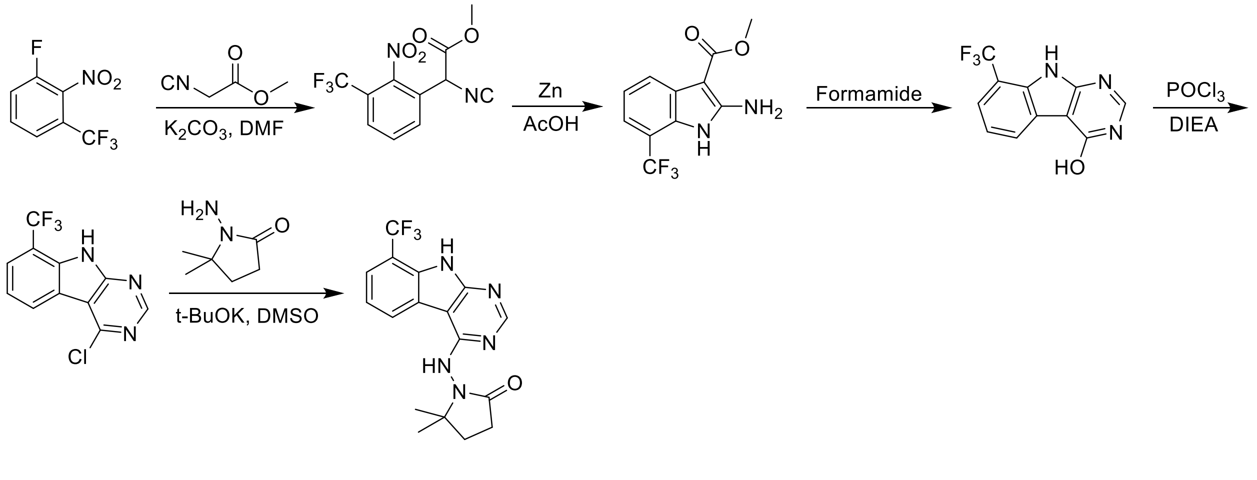

To a solution of 1-fluoro-2-nitro-3-(trifluoromethyl)benzene (10 g, 47.84 mmol) in dry DMF (80 mL) was added methyl 2-isocyanoacetate (4.73 g, 47.84 mmol) and $K_2CO_3$ (19.81 g, 143.54 mmol). The solution was stirred at 90°C for 16 hours. The mixture was adjusted to be weakly acidic pH with 2N HCl and extracted with ethyl acetate (150 mL *3). The combined organic layers were washed with brine (200 mL), dried over $Na_2SO_4$, filtered and concentrated under reduced pressure. The residue was purified by silica gel chromatography (0-50% ethyl acetate/hexanes) to afford methyl 2-isocyano-2-(2-nitro-3-(trifluoromethyl)phenyl)acetate as yellow solid (11 g, Yield: 79.82%). LCMS (ESI): m/z= 306.1 $(M+18)^+$.
A mixture of methyl 2-isocyano-2-(2-nitro-3-(trifluoromethyl)phenyl)acetate (11 g, 38.19 mmol) and acetic acid (150 mL) was heated to 40 °C. Zinc (19.98 g, 305.55 mmol) was then added in portions at a rate such that the reaction temperature did not rise above 60 °C. After stirring at 60°C for 2 h, the reaction mixture was cooled to room temperature and filtered through celite. The filtrate was concentrated and the crude product was purified by silica gel column chromatography (dicholoromethane/methanol 10:1) to give methyl 2-amino-7-(trifluoromethyl)-1H-indole-3-carboxylate as a white solid (6.3 g, Yield: 63.93%). $^1$H NMR (500 MHz, DMSO-*d*6) δ 10.89 (s, 1H), 7.84 (d, J = 7.6 Hz, 1H), 7.20 (d, J = 7.6 Hz, 1H), 7.13 (t, J = 7.7 Hz, 1H), 6.54 (s, 2H), 3.79 (s, 3H). LCMS (ESI): m/z=259.2 $(M+H)^+$.
A solution of methyl 2-amino-7-(trifluoromethyl)-1H-indole-3-carboxylate (6.3 g, 24.41 mmol) in formamide (40 mL) was stirred at 200°C for 2 hours. The reaction mixture was then cooled to room temperature and poured into 300 mL of water. The resulting mixture was allowed to stand for 15 min, the solids were collected by filtration, washed with water, and dried to afford 8-(trifluoromethyl)-9H-pyrimido[4,5-b]indol-4-ol as a brown solid (6 g, Yield: 97.12%). $^1$H NMR (500 MHz, DMSO-*d*6) δ 12.65 (s, 1H), 12.47 (s, 1H), 8.30 (d, J = 7.8 Hz, 1H), 8.28 (d, J = 6.8 Hz, 1H), 7.68 (d, J = 7.6 Hz, 1H), 7.43 (t, J = 7.7 Hz, 1H). LCMS (ESI): m/z=254.2 $(M+H)^+$.
To a solution of 8-(trifluoromethyl)-9H-pyrimido[4,5-b]indol-4-ol (3 g, 11.85 mmol) in $POCl_3$ (30 mL) was added DIEA (0.5 mL). The mixture was heated to reflux for 2 hours. After cooling to room temperature, the mixture was concentrated under reduced pressure and diluted with ACN, then adjusted slowly to pH 7.0 with ammonium hydroxide. The resulting solids were filtered, washed with water and dried to obtain 4-chloro-8-(trifluoromethyl)-9H-pyrimido[4,5-b]indole (1.7 g, Yield: 52.90%) as a brown solid.$^1$H NMR (500 MHz, DMSO-*d*6) δ 13.25 (s, 1H), 8.92 (s, 1H), 8.59 (d, J = 7.9 Hz, 1H), 7.98 (d, J = 7.7 Hz, 1H), 7.75 – 7.46 (m, 1H). LCMS (ESI): m/z=272.1 $(M+H)^+$.
To a solution of 4-chloro-8-(trifluoromethyl)-9H-pyrimido[4,5-b]indole (200 mg, 0.74 mmol) in dry DMSO (4 mL) was added 1-amino-5,5-dimethylpyrrolidin-2-one (142 mg, 1.1 mmol) and t-BuOK (414 mg, 3.69 mmol). The mixture was stirred at 100°C for 45 minutes, diluted with ethyl acetate (50.0 mL) and washed with water (10.0 mL), brine (10.0 mL). The organic layers was dried over $Na_2SO_4$ and concentrated under reduced pressure. The residue was purified by reverse phase chromatography (10mM $NH_4HCO_3$ in water, 5%-60% ACN) to give 5,5-dimethyl-1-((8-(trifluoromethyl)-9H-pyrimido[4,5-b]indol-4-yl)amino)pyrrolidin-2-one (**AVI-6371**) as a white solid (25 mg, yield: 9.33%). $^1$H NMR (500 MHz, DMSO-*d*6) δ 12.57 (s, 1H), 9.33 (s, 1H), 8.73 (d, J = 7.8 Hz, 1H), 8.49 (s, 1H), 7.77 (d, J = 7.7 Hz, 1H), 7.48 (t, J = 7.8 Hz, 1H), 2.45 (t, J = 7.8 Hz, 2H), 2.04 (t, J = 7.7 Hz, 2H), 1.30 (s, 6H). LCMS (ESI): m/z=364.3 $(M+H)^+$.

**AVI-6354**

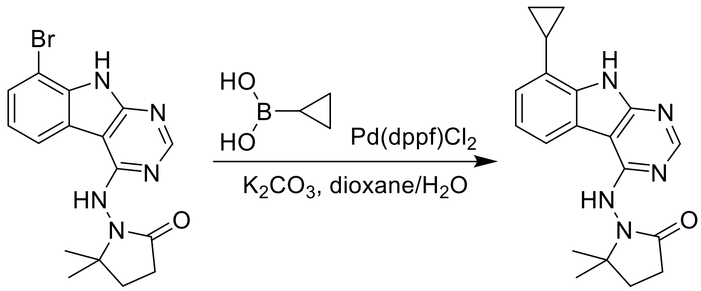

To a solution of 1-((8-bromo-9H-pyrimido[4,5-b]indol-4-yl)amino)-5,5-dimethylpyrrolidin-2-one (200 mg, 0.53 mmol) in dioxane (5 mL) and water (0.5 mL) was added cyclopropylboronic acid (184 mg, 2.14 mmol), $K_2CO_3$ (185 mg, 1.34 mmol) and Pd(dppf)$Cl_2$ (58 mg, 0.08 mmol). The mixture was stirred at 95 °C for 16 hours under $N_2$. The reaction mixture was then diluted with water (50 mL) and extracted with DCM (3 x 50 mL). The organic layers were dried over $Na_2SO_4$ and concentrated under reduced pressure. The residue was purified by reverse phase chromatography (10mM $NH_4HCO_3$ in water, 5%-50% ACN) to give 1-((8-cyclopropyl-9H-pyrimido[4,5-b]indol-4-yl)amino)-5,5-dimethylpyrrolidin-2-one (**AVI-6354**) as a white solid (25 mg, yield: 13.91%). $^1$H NMR (500 MHz, DMSO-*d6*) δ 12.22 (s, 1H), 9.05 (s, 1H), 8.40 (s, 1H), 8.22 (d, J = 7.8 Hz, 1H), 7.19 (t, J = 7.7 Hz, 1H), 6.97 (d, J = 7.5 Hz, 1H), 2.43-2.39 (m, 3H), 2.03 (t, J = 7.8 Hz, 2H), 1.29 (s, 6H), 1.08 – 1.02 (m, 2H), 0.80 – 0.75 (m, 2H). LCMS (ESI): m/z=336.3 $(M+H)^+$.

**AVI-6357**

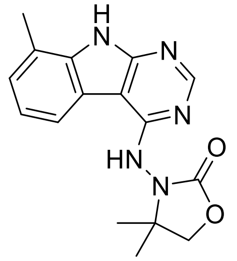

To a solution of 4-chloro-8-methyl-9H-pyrimido[4,5-b]indole(50 mg, 0.23 mmol) and 3-amino-4,4-dimethyloxazolidin-2-one (45 mg, 0.34 mmol) in i-PrOH (0.8 mL) was added 1N aqueous HCl (0.2 mL). The reaction mixture was stirred at100 ºC for 18 h, concentrated and the residue was purified by reverse phase chromatography (water/10-100% MeCN) to obtain 18 mg (25%) 4,4-dimethyl-3-((8-methyl-9H-pyrimido[4,5-b]indol-4-yl)amino)oxazolidin-2-one (**AVI-6357**) as a cream colored solid. $^1$H NMR (DMSO-$d_6$, 400 MHz) δ 12.15 (s, 1H), 9.27 (s, 1H), 8.46 (s, 1H), 8.24 (br d, 1H, *J*=7.3 Hz), 7.22-7.27 (m, 2H), 4.29 (s, 2H), 2.57 (s, 3H), 1.35 (br s, 6H). LCMS (ESI): m/z= 312 $(M+H)^+$

**AVI-6372**

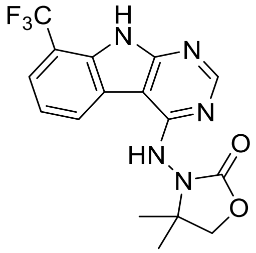


To a solution of 4-chloro-8-(trifluoromethyl)-9H-pyrimido[4,5-b]indole (50 mg, 0.18 mmol) and 3-amino-4,4-dimethyloxazolidin-2-one (36 mg, 0.27 mmol) in i-PrOH (0.8 mL) was added 1N aqueous HCl (0.2 mL). The reaction mixture was stirred at100 °C for 18 h, concentrated and the residue was purified by reverse phase chromatography (water/10-100% MeCN) to obtain 2.1 mg (3%) of 4,4-dimethyl-3-((8-(trifluoromethyl)-9H-pyrimido[4,5-b]indol-4-yl)amino)oxazolidin-2-one (**AVI-6372**) as a white colored solid. 1H NMR (METHANOL-d4, 400 MHz) δ 8.50-8.57 (m, 2H), 7.79 (d, 1H, *J*=7.8 Hz), 7.49 (t, 1H, *J*=7.8 Hz), 4.41 (br s, 2H), 1.47 (br s, 6H). LCMS (ESI): m/z= 366 $(M+H)^+$

**AVI-6373**

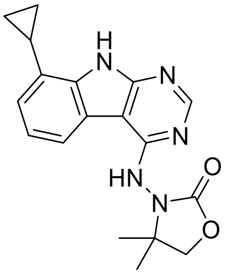


To a solution of 4-chloro-8-(cyclopropyl)-9H-pyrimido[4,5-b]indole (50 mg, 0.21 mmol) and 3-amino-4,4-dimethyloxazolidin-2-one (40 mg, 0.31 mmol) in i-PrOH (0.8 mL) was added 1N aqueous HCl (0.2 mL). The reaction mixture was stirred at100 °C for 18 h, concentrated and the residue was purified by reverse phase chromatography (water/10-100% MeCN) to obtain 2.2 mg (3%) of 4,4-dimethyl-3-((8-(cyclopropyl)-9H-pyrimido[4,5-b]indol-4-yl)amino)oxazolidin-2-one (**AVI-6373**) as a white colored solid. $^{1}$H NMR (METHANOL-$d_4$, 400 MHz) δ 8.45 (s, 1H), 8.05 (d, 1H, *J*=7.8 Hz), 7.27 (t, 1H, *J*=7.7 Hz), 7.12 (d, 1H, *J*=7.3 Hz), 4.40 (br s, 2H), 2.28-2.35 (m, 1H), 1.45 (br s, 6H), 1.12 (dd, 2H, *J*=1.8, 8.4 Hz), 0.83 (dd, 2H, *J*=1.7, 5.1 Hz). LCMS (ESI): m/z= 338 $(M+H)^+$

**AVI-3769**

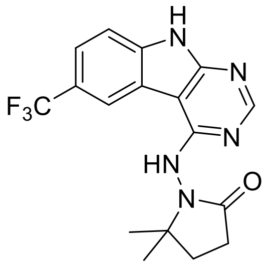

A solution of 4-chloro-6-(trifluoromethyl)-9H-pyrimido[4,5-b]indole (900 mg, 3.3 mmol) and 1-amino-5,5-dimethylpyrrolidin-2-one hydrochloride (637.6 mg, 4.98 mmol,) in isopropanol (10 mL) and concentrated HCl (1 drop) was stirred at 100°C for 16 hours. The reaction mixture was concentrated under reduced pressure and the residue was purified by reverse phase chromatography ((0.1% NH4HCO3 in water, 10-100%ACN) to obtain 102 mg (9.9%) of 5,5-dimethyl-1-((6-(trifluoromethyl)-9H-pyrimido[4,5-b]indol-4-yl)amino)pyrrolidin-2-one (**AVI-3769**) as a white solid. 1H NMR (500 MHz, DMSO) δ 12.55 (s, 1H), 9.47 (s, 1H), 8.93 (s, 1H), 8.46 (s, 1H), 7.75 (d, 1H, J = 8.5 Hz), 7.68 (d, 1H, J = 8.5 Hz), 2.45 (t, 2H, J = 7.8 Hz), 2.04 (d, 2H, J = 7.5 Hz), 1.30 (s, 6H). LCMS (ESI): m/z 364 $(M+H)^+$

**AVI-3865**

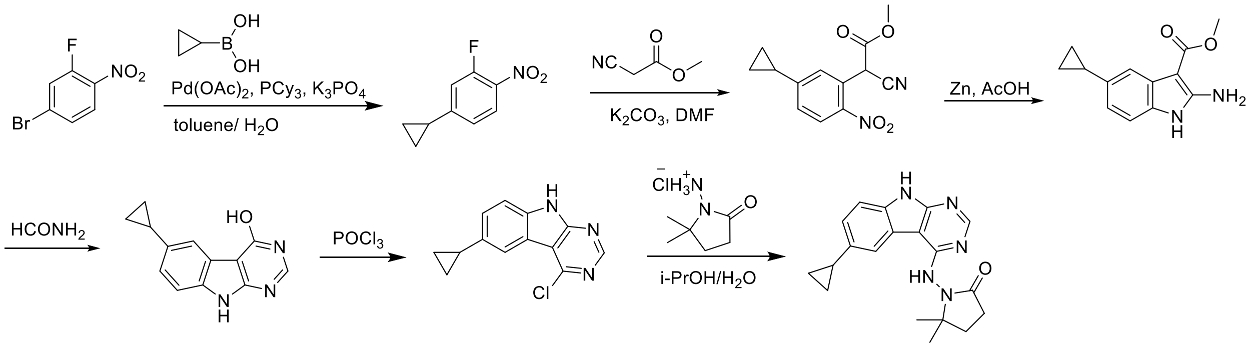


To a solution of 4-bromo-2-fluoro-1-nitrobenzene (24 g, 109.1 mmol) in toluene (240 mL) and water (24 mL) was added cyclopropylboronic acid (14.05 g, 163.63 mmol), potassium phosphate (69.387 g, 327.3 mmol), tricyclohexylphosphine (10 g, 54.55 mmol) an palladium acetate(2.55 g, 10.91 mmol). After stirring at 95 °C for 16 h, the reaction mixture was poured into water (400 mL) and extracted with dichloromethane (3 x 800 mL). The organic layer was dried over $Na_2SO_4$, concentrated under reduced pressure and the residue was purified by silica gel chromatography (2:1 petroleum ether/ethyl acetate) to obtain 19.8 g (99.4%) of 4-cyclopropyl-2-fluoro-1-nitrobenzene as a yellow solid. LCMS (ESI): m/z= 182 $(M+H)^+$

To a solution of 4-cyclopropyl-2-fluoro-1-nitrobenzene (19 g, 104.97 mmol) in DMF (100 mL) was added methyl cyanoacetate (20.78g, 209.9 mmol) and potassium carbonate (43.5 g, 314.9 mmol). After stirring at 80°C for 2 h, the reaction mixture was cooled to room temperature, acidified with 2N HCl (ca. 2000 mL), and extracted with ethyl acetate (300 mL *3). The organic layers were washed with brine (100 mL), dried over $Na_2SO_4$, concentrated under reduced pressure and the residue was purified by silica gel chromatography (10:1 petroleum ether/ethyl acetate) to obtain 20 g (73.5%) of methyl 2-cyano-2-(5-cyclopropyl-2-nitrophenyl)acetate as a yellow solid. LCMS (ESI): m/z= 278 $(M+18)^+$

To a solution of methyl 2-cyano-2-(5-cyclopropyl-2-nitrophenyl)acetate (20 g, 76.9 mmol) in glacial acetic acid (200 ml), was added in two portions, zinc dust (50.3 g, 769 mmol) After stirring at 60°C for 2 h, the reaction mixture was cooled to room temperature, filtered and the residue was washed with with THF. The filtrate was concentrated and the residue was purified by silica gel (10:1 dichloromethane /methanol) to obtain 15 g (85.2%) of methyl 2-amino-5-cyclopropyl-1H-indole-3-carboxylate as a yellow solid.LCMS (ESI): m/z=231 $(M+H)^+$
A solution of methyl 2-amino-5-cyclopropyl-1H-indole-3-carboxylate (15g, 65.21 mmol) in formamide (150 mL) was stirred at 220°C for 2 hours. The reaction mixture was then cooled to room temperature and poured in 75 ml of water. The resulting mixture was allowed to stand for 15 min and precipitate formed were collected by filtration, washed with water, and dried in vacuo to obtain 11 g (75.0%) of 6-cyclopropyl-9H-pyrimido[4,5-b]indol-4-ol as a brown solid. LCMS (ESI): m/z=226 $(M+H)^+$
To a solution of 6-cyclopropyl-9H-pyrimido[4,5-b]indol-4-ol (11 g, 48.88 mmol) in POCl3 (50 mL) was added DIPEA (18.7 g, 146.6 mmol) and refluxed 16 hours. The reaction mixture was then cooled to room temperature, concentrated and poured into 20 mL of water. The resulting precipitate was filtered, the residue was triturated with ethyl acetate and filtered to afford 5.4 g (45.7%) of 4-chloro-6-cyclopropyl-9H-pyrimido[4,5-b]indole as black solid. LCMS (ESI): m/z=244.1 $(M+H)^+$
A neat mixture of 4-chloro-6-cyclopropyl-9H-pyrimido[4,5-b]indole (260 mg, 1.07 mmol) and 1-amino-5,5-dimethylpyrrolidin-2-one hydrochloride (351 mg, 2.14 mmol) was heated at 115°C for 16 h. The reaction mixture was taken in water (10 mL) and extracted with ethyl acetate (3 x 20 mL). The organic layer was dried over $Na_2SO_4$ and concentrated. The residue was purified by reverse phase chromatography (0.1% $NH_4HCO_3$ in water/MeCN) to obtain 26 mg (7.26%) of 1-((6-cyclopropyl-9H-pyrimido[4,5-b]indol-4-yl)amino)-5,5-dimethylpyrrolidin-2-one (**AVI-3865**) as a white solid. $^1$H NMR (500 MHz, DMSO) δ 11.94 (s, 1H), 9.10 (s, 1H), 8.34 (s, 1H), 8.08 (s, 1H), 7.38 (d, J = 8.3 Hz, 1H), 7.20 (d, J = 8.3 Hz, 1H), 2.47 – 2.35 (m, 2H), 2.06 (ddd, J = 16.3, 11.1, 6.6 Hz, 3H), 1.27 (d, J = 19.4 Hz, 6H), 1.01 – 0.93 (m, 2H), 0.85 – 0.74 (m, 2H). LCMS (ESI): m/z= 336 $(M+H)^+$

**AVI-6412**

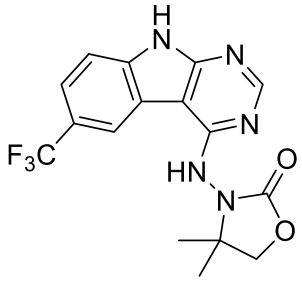


To a solution of 4-chloro-6-(trifluoromethyl)-9H-pyrimido[4,5-b]indole (50 mg, 0.18 mmol) and 3-amino-4,4-dimethyloxazolidin-2-one (36 mg, 0.27 mmol) in i-PrOH (0.8 mL) was added 1N aqueous HCl (0.2 mL). The reaction mixture was stirred at100 ºC for 18 h, concentrated and the residue was purified by reverse phase chromatography (water/10-100% MeCN) to obtain 6 mg (9%) of 4,4-dimethyl-3-((6-(trifluoromethyl)-9H-pyrimido[4,5-b]indol-4-yl)amino)oxazolidin-2-one (**AVI-6412**) as a white colored solid. $^1$H NMR (METHANOL-$d_4$, 400 MHz) δ 8.68 (s, 1H), 8.47 (s, 1H), 7.69-7.77 (m, 2H), 4.41 (br s, 2H), 1.46 (br s, 6H). LCMS (ESI): m/z= 366 $(M+H)^+$

**AVI-6451**

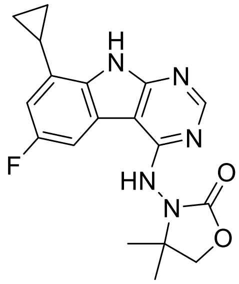

To a solution of 4-chloro-8-cyclopropyl-6-fluoro-9H-pyrimido[4,5-b]indole (900 mg, 3.45 mmol) in i-PrOH (20 mL) was added 3-amino-4,4-dimethyloxazolidin-2-one (897mg, 6.9mmol) and conc.HCl (0.1 mL). After stirring at 100°C for 72 h, the reaction mixture was concentrated and the residue was purified by reverse phase chromatography (0.1% NH4HCO3 in water, 10-100%ACN) to obtain 3-((8-cyclopropyl-6-fluoro-9H-pyrimido[4,5-b]indol-4-yl)amino)-4,4-dimethyloxazolidin-2-one (**AVI-6451**) as a white solid (220 mg, Yield: 17.9%). 1H NMR (500 MHz, DMSO-d6) δ 12.39 (s, 1H), 9.31 (s, 1H), 8.47 (s, 1H), 8.13 (d, J = 8.2 Hz, 1H), 6.84 (dd, J = 10.8, 2.2 Hz, 1H), 4.29 (s, 2H), 2.48 – 2.42 (m, 1H), 1.46 – 1.27 (m, 6H), 1.09 (dd, J = 8.3, 1.7 Hz, 2H), 0.85 (d, J = 4.1 Hz, 2H). LCMS (ESI): m/z 356.1 $(M+H)^+$

**AVI-6452**

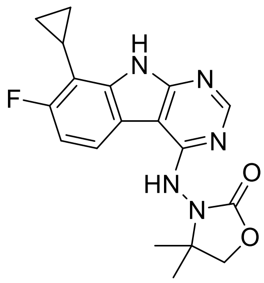

To solution of 4-chloro-8-cyclopropyl-7-fluoro-9H-pyrimido[4,5-b]indole (900mg, 3.45mmol) in isopropanol (20 mL) was added 3-amino-4,4-dimethyloxazolidin-2-one (897mg, 6.9mmol) and conc. HCl (0.1 ml). The mixture was stirred at 100°C for 72 hours, concentrated under reduced pressure and the residue was purified by reverse phase chromatography (0.1% NH4HCO3 in water, 10-100% MeCN) to obtain 3-((8-cyclopropyl-7-fluoro-9H-pyrimido[4,5-b]indol-4-yl)amino)-4,4-dimethyloxazolidin-2-one (**AVI-6452**) as a white solid (240mg, yield: 19.6%). $^1$H NMR (400 MHz, DMSO) δ 12.31 (s, 1H), 9.32 (s, 1H), 8.46 (s, 1H), 8.24 (dt, *J* = 13.8, 6.9 Hz, 1H), 7.10 (dd, *J* = 11.7, 8.7 Hz, 1H), 4.29 (s, 2H), 2.08 (tt, *J* = 8.6, 5.4 Hz, 1H), 1.30 (t, *J* = 26.8 Hz, 6H), 1.18 – 0.98 (m, 2H), 0.92 – 0.76 (m, 2H). LCMS (ESI): m/z 356.2 $(M+H)^+$.

#### AVI-6612

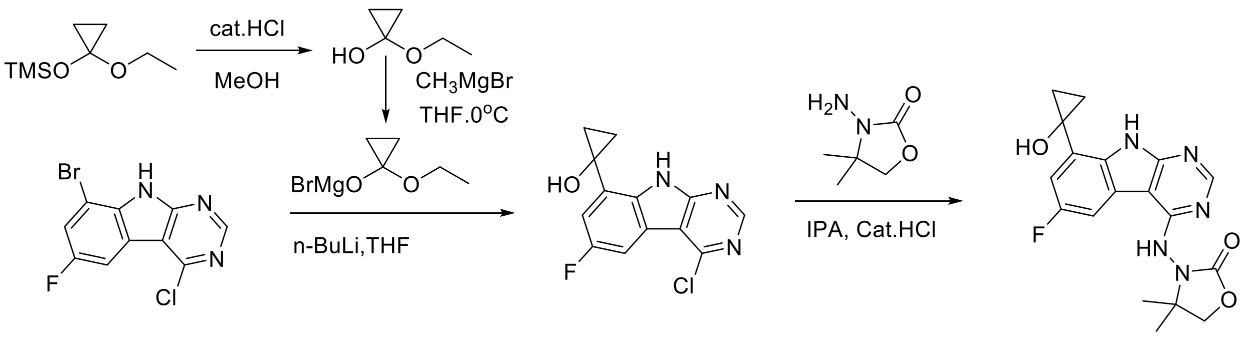


To a solution of (1-ethoxycyclopropoxy)trimethylsilane (1.0 g, 5.57 mmol) in MeOH (20 mL) was added conc. HCl (0.1 ml), The mixture was stirred at room temperature for 16 hours and concentrated under reduced pressure to give crude 1-ethoxycyclopropan-1-ol as a colorless oil. It was used in the next step without any purification. LCMS (ESI): m/z=103 $(M+H)^+$
A 2.5 M solution of *n*-BuLi in pentane (4ml, 10mmol.) was slowly added to a solution of 8-bromo-4-chloro-6-fluoro-9H-pyrimido[4,5-b]indole (1.0 g, 3.34 mmol) in anhydrous THF (15 ml) at -78 °C over a period of 10 min. The resultant blood red solution was stirred at -78 °C for 0.5 h. Concurrently, to a cooled (0 °C, ice/water bath) solution of 1-ethoxycyclopropan-1-ol (374 mg, 3.68 mmol) in anhydrous THF(8 ml) was added 2.5 M solution of MeMgBr in diethyl ether (4.0 ml, 10mmol,) dropwise via syringe. The resultant white suspension was stirred at 0 °C for 10 min. The above organic lithium solution was then cannulated into this suspension. The resultant reaction mixture was stirred at ambient temperature for 30 min, followed by stirring at 40 °C overnight. The reaction mixture was then cooled down to 0 °C (ice/water bath) and quenched with saturated aqueous $NH_4Cl$. The separated aqueous layer was extracted with ethyl acetate (80 ml*3), The combined organic layers were dried over anhydrous $Na_2SO_4$, filtered and concentrated under reduced pressure. The crude residue was then purified by flash chromatography on silica gel (2:1 petroleum ether/ ethyl acetate) to obtain 1-(4-chloro-6-fluoro-9H-pyrimido[4,5-b]indol-8-yl)cyclopropan-1-ol as a white solid (150mg, Yield: 16.2%). LCMS (ESI): m/z=287.0 $(M+H)^+$.
To solution of -(4-chloro-6-fluoro-9H-pyrimido[4,5-b]indol-8-yl)cyclopropan-1-ol (100 mg, 0.36 mmol) in *i*-PrOH (6 mL) was added 3-amino-4,4-dimethyloxazolidin-2-one (94 mg, 0.72 mmol) and conc. HCl (0.1 ml). After stirring at 100°C for 16 hours, the reaction mixture was concentrated and the residue was purified by reverse phase chromatography (10 mmol $NH_4HCO_3$ in water, 10-100% ACN) to obtain 3-((6-fluoro-8-(1-hydroxycyclopropyl)-9H-pyrimido[4,5-b]indol-4-yl)amino)-4,4-dimethyloxazolidin-2-one (**AVI-6612**) as a white solid (15 mg, yield: 11.2%). $^1$H NMR (400 MHz, DMSO) δ 11.73 (d, *J* = 24.3 Hz, 1H), 9.36 (s, 1H), 8.49 (s, 1H), 8.25 (dd, *J* = 9.7, 1.9 Hz, 1H), 7.15 (dd, *J* = 10.2, 2.4 Hz, 1H), 6.10 (d, *J* = 5.1 Hz, 1H), 4.29 (s, 2H), 1.30 (dd, *J* = 31.3, 27.0 Hz, 6H), 1.14 (d, *J* = 13.9 Hz, 2H), 1.08 – 0.89 (m, 2H). LCMS (ESI): m/z 372.1 $(M+H)^+$.

**AVI-6613**

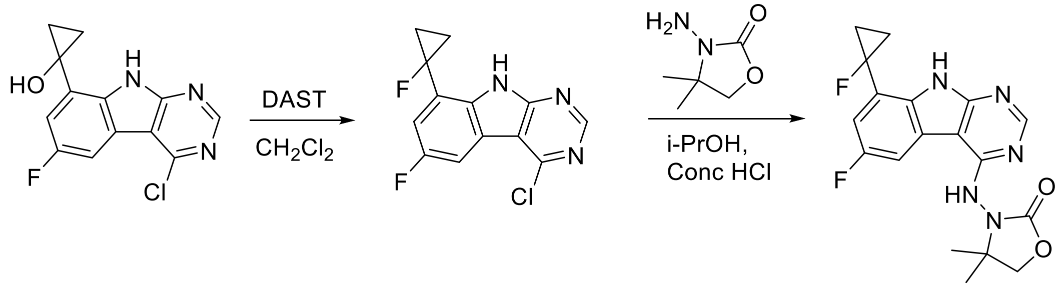


To a solution of 1-(4-chloro-6-fluoro-9H-pyrimido[4,5-b]indol-8-yl)cyclopropan-1-ol (200 mg, 0.72 mmol) in dichloromethane (10 mL) was added DAST (581 mg, 3.61 mmol) and the mixture was stirred at room temperature for an hour under nitrogen gas. The mixture was then quenched with saturated aqueous NaHCO3. The layers were separated and aqueous layer was extracted with ethyl acetate (30 ml*3), The combined organic layers were dried over anhydrous $Na_2SO_4$, filtered and concentrated under reduced pressure. The crude residue was then purified by silica gel chromatography (3:1 petroleum ether/ ethyl acetate) to afford 4-chloro-6-fluoro-8-(1-fluorocyclopropyl)-9H-pyrimido[4,5-b]indole as a white solid (150 mg, Yield: 74.5%). LCMS (ESI): m/z=280.2 $(M+H)^+$.

To solution of 4-chloro-6-fluoro-8-(1-fluorocyclopropyl)-9H-pyrimido[4,5-b]indole (100 mg, 0.36 mmol) in *i*-PrOH (6 mL) was added 3-amino-4,4-dimethyloxazolidin-2-one (94 mg, 0.72 mmol) and conc. HCl (0.1 ml). The mixture was stirred at 100°C for 16 h, concentrated and the residue was purified by reverse phase chromatography (10 mmol $NH_4HCO_3$ in water, 10-100% ACN) to obtain 3-((6-fluoro-8-(1-fluorocyclopropyl)-9H-pyrimido[4,5-b]indol-4-yl)amino)-4,4-dimethyloxazolidin-2-one (**AVI-6613**) as a white solid (13 mg, yield: 9.76%). $^1$H NMR (400 MHz, DMSO) δ 12.41 (d, *J* = 36.2 Hz, 1H), 9.42 (s, 1H), 8.51 (s, 1H), 8.42 (d, *J* = 9.5 Hz, 1H), 7.40 (d, *J* = 9.7 Hz, 1H), 4.30 (s, 2H), 1.55 (d, *J* = 18.6 Hz, 2H), 1.34 (d, *J* = 33.5 Hz, 6H), 1.20 (t, *J* = 15.1 Hz, 2H). LCMS (ESI): m/z 374.2 $(M+H)^+$.
